# The low complexity motif of cytoplasmic polyadenylation element binding protein 3 (CPEB3) is critical for the trafficking of its targets in neurons

**DOI:** 10.1101/2020.05.16.100032

**Authors:** Lenzie Ford, Arun Asok, Arielle D. Tripp, Cameron Parro, Michelle Fitzpatrick, Christopher A. de Solis, Neeva Shafiian, Luana Fioriti, Rajesh Kumar Soni, Eric R. Kandel

## Abstract

Biomolecular condensates, membraneless organelles found throughout the cell, play critical roles in many aspects of cellular function. Ribonucleoprotein granules (RNPs), a type of biomolecular condensate found in neurons that are necessary for local protein synthesis and are involved in long-term potentiation (LTP). Several RNA-binding proteins present in RNPs are necessary for the synaptic plasticity involved in LTP and long-term memory. Most of these proteins possess low complexity motifs, allowing for increased promiscuity. We explore the role the low complexity motif plays for RNA binding protein cytoplasmic polyadenylation element binding protein 3 (CPEB3), a protein necessary for long-term memory persistence. We found that RNA binding and SUMOylation are necessary for CPEB3 localization to the P body, thereby having functional implications on translation. Here, we investigate the role of the low complexity motif of CPEB3 and find that it is necessary for P body localization and downstream targeting for local protein synthesis.

## Introduction

Local protein synthesis is a critical component of long-term memory, allowing for the morphological changes observed in activated neurons, such as synapse enlarging, spine formation, and AMPA-receptor insertion (for review, see Hanus and Schuman, 2013). A key component of local protein synthesis near active synapses is the presence of ribosomes, mRNAs, and translation-associated proteins (for review, see Hanus and Schuman, 2013). Interestingly, these key components are often compartmentalized into membraneless organelles known as ribonucleoprotein granules (RNPs). RNPs contain a high concentration of mRNAs and proteins involved in translation, and are shuttled from the soma to distal processes by motor protein movement along the cytoskeleton (for review, see Di Liegro et al., 2014).

Of the proteins commonly found in RNPs, RNA-binding proteins are highly prevalent. Our lab has characterized the RNA-binding protein cytoplasmic polyadenylation element binding protein 3 (CPEB3), and found that it is necessary for long-term memory maintenance (Fioriti et al., 2015). This multi-faceted polyQ-containing RNA-binding protein with prion-like properties is regulated by what appears to be a complex relationship between the cellular environment and the protein’s structure. From our previous work, we know that CPEB3 is soluble in the basal state and functions to inhibit translation of mRNA targets (Fioriti et al., 2015; Drisaldi et al., 2015; Stephan et al., 2015). Upon neuronal activation, CPEB3 undergoes a structural change that makes the protein semi-insoluble and switches CPEB3 to a state that promotes translation of mRNA targets (Fioriti et al., 2015; Stephan et al., 2015).

We have developed a working model of CPEB3 regulation in the neuron **(Figure 1)**: According to this model, CPEB3 exits the nucleus via nuclear export signals located in the C-terminal zinc finger (Ford et al., 2019) and in the second prion domain (Chao et al., 2012). Once in the cytoplasm, CPEB3 is SUMOylated and functionally inhibits translation of mRNA targets (Drisaldi et al., 2015). SUMOylated CPEB3 is then localized to RNP processing bodies (P bodies), in part through its RNA Recognition Motif 1 (Ford et al., 2019). However, 30 minutes following chemical long-term potentiation (cLTP) CPEB3 is deSUMOylated (Drisaldi et al., 2015) and leaves P bodies; by 1 hour post-cLTP, it relocates to the polysome (Ford et al., 2019). Presumably, this is where oligomeric CPEB3 promotes translation of its mRNA targets, in part through its N-terminal prion domain 1 (Fioriti et al., 2015; Stephan et al., 2015). After neuronal stimulation, CPEB3 is also ubiquitinated by Neuralized 1 (Pavlopolous et al., 2011).

**Figure 1.**
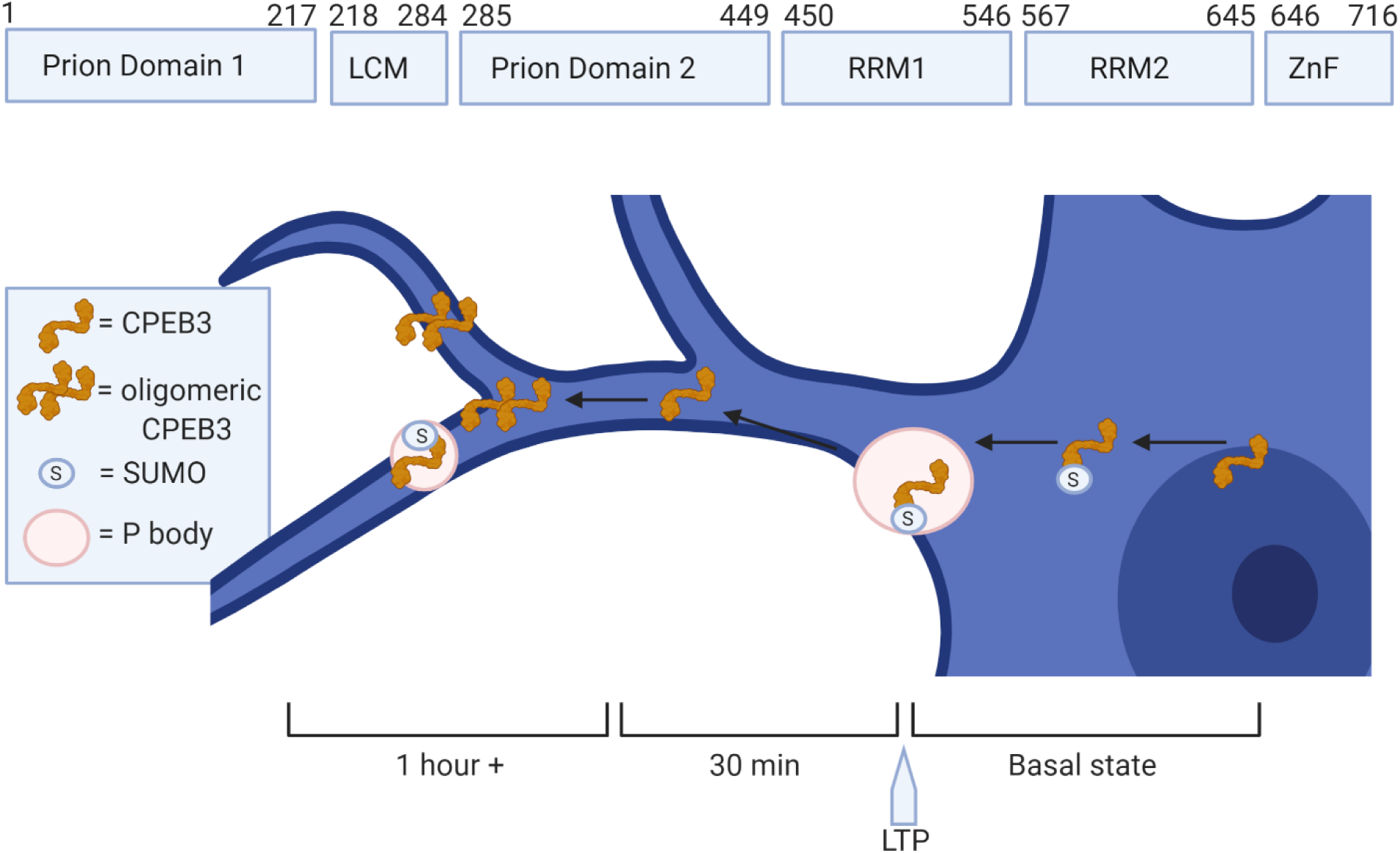
CPEB3 protein regulation in the neuron. A sequence diagram and spatiotemporal cellular diagram of CPEB3. CPEB3 exits the nucleus because of sequences in the zinc-finger domain (ZnF) and Prion Domain 2. CPEB3 is SUMOylated (circle with S) and moves to the P body in the basal state and early timepoints of long-term potentiation (LTP). This is dependent upon sequences in the RNA Recognition Motif (RRM) 1. 30 minutes post-LTP, CPEB3 is deSUMOylated and leaves the P body. By 1 hour post-LTP, CPEB3 is located in the polysome, as well as again in P bodies. CPEB3 at the polysome is oligomerized, due to sequences in Prion Domain 1. Low complexity motif (LCM).

Within the past decade, P bodies have been implicated in localized neuronal translation (for review, see Kiebler and Bassell, 2006). Seminal work conducted in non-mammalian and somatic cells, found that P bodies contain mRNA destined for degradation through the RNA-induced silencing complex (for review, see Decker and Parker, 2012). Paradoxically, P bodies are known to play a dual role in both degradation and translation within neurons. Indeed, a number of RNA-binding proteins involved in synaptic plasticity, such as Staufen, Fragile X Mental Retardation protein (FMRP), Zipcode binding protein 1, and La, have been found in neuronal P bodies (Barbee et al., 2006; Tiruchinapalli et al., 2003; Cougot et al., 2008; Kota et al., 2016). Some of the proteins within P bodies move to distal processes in order to promote translation in an NMDA-dependent manner (Tiruchinapalli et al., 2003). This is similar to CPEB3, which appears to promote translation, at least partially, in an NMDA-dependent manner (Chao et al., 2013; Huang et al., 2014; Chao et al., 2012; Wang and Huang, 2012). This investigation into well-characterized CPEB3 provides crucial insight into P body-regulated local protein synthesis involved in memory. For this reason, we delve further into mechanistic understanding of neuronal regulation of CPEB3 in P bodies. Specifically, we explore the promiscuous low complexity motif and find that it is necessary for P body localization and for downstream targeting for local protein synthesis.

## Results

### CPEB3 inhibits translation of mRNA targets via its low complexity motif

CPEB3 has two distinct functions; in the basal state, CPEB3 inhibits mRNA targets (Fioriti et al., 2015; Stephan et al., 2015; Ford et al., 2019). Upon neuronal stimulation, CPEB3 promotes the translation of its mRNA targets (Fioriti et al., 2015; Stephan et al., 2015; Ford et al., 2019). Recent work from our lab has identified that CPEB3 inhibits translation by localization to P bodies (Ford et al., 2019). SUMOylation of CPEB3 drives this localization, and encourages CPEB3 inhibitory function and inhibits spine growth (Ford et al., 2019; Drisaldi et al., 2015). We decided to delve further into understanding the mechanism behind CPEB3 inhibitory function, and ultimately, its importance for long-term memory.

We utilized the SUMOylation algorithm GPS-SUMO (Zhao et al., 2014) and found that a SUMO interaction motif (SIM) is predicted to span a segment of the low complexity motif at amino acids 220-242 (Score= 1.922, p value 0.058). SIMs do not require a lysine, and bond non-covalently. Thus, we screened a combination of mutants within the predicted SIM for effect on translation. Of the 16 mutants screened, the serine to alanine mutation of amino acids 240, 241, 242 (identified as S240-242A) was disruptive to function **(Figure 2; Figure S1)**.

**Figure 2.**
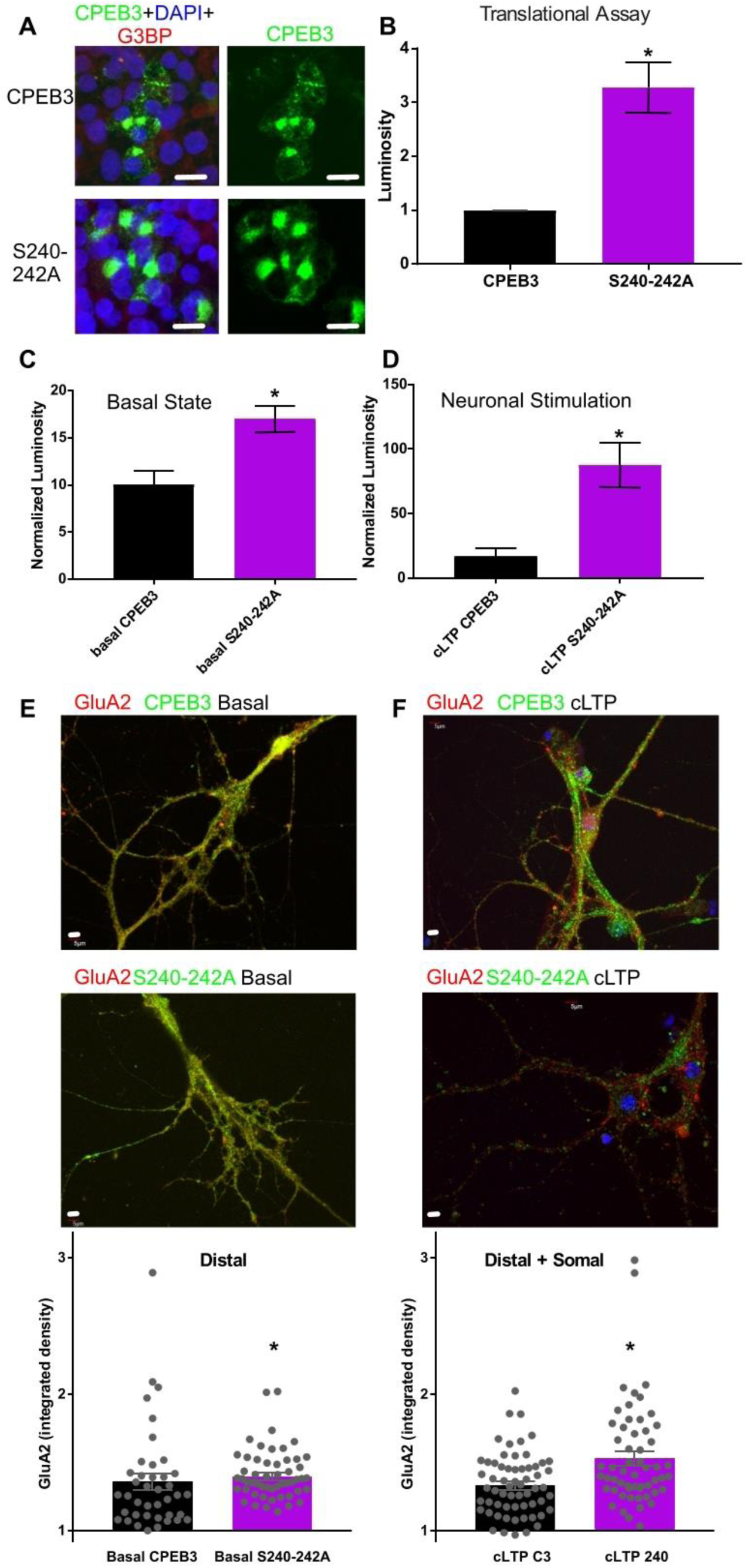
A mutation in the low complexity motif disrupts CPEB3 morphology and function. A. CPEB3 or the S240-242A mutant (green) was expressed in HEK cells, which were co-labeled for DAPI (blue) and G3BP (red). Scale bars = 10 μm. B. Mutant and full length CPEB3 were expressed in confluent HEK cells along with an actin-3’UTR-Renilla reporter plasmid. Translation of the reporter was measured using a luciferase assay, and luminosity was measured as a readout of translation. The S240-242A mutant had significantly increased luminosity from CPEB3 levels, *= p<0.05, as measured with a Student’s t-test. n=9. Error bars = +SEM. C, D. Full-length CPEB3 or the S240-242A mutant were expressed in primary neuronal cultures along with a SUMO2-3’UTR-Renilla reporter plasmid. Translation of the reporter was measured using a luciferase assay, and luminosity was measured as a readout of translation. The S240-242A mutant (purple) has significantly increased luminosity in a translational assay compared to full-length CPEB3 (black) in basal (C; p=0.0042, n=8) and stimulated (D; p=0.0004, n=9) conditions. E, F. CPEB3 or S240-242A (green) were expressed in primary neuronal cultures and GluA2 (red) was measured under basal and cLTP stimulated conditions; DAPI in blue. Neurons expressing the S240-242A mutant have significantly increased GluA2 levels in distal sites compared to neurons expressing CPEB3 in the basal (E; p=0.0006, n=40) and stimulated (F; p=0.0004, n=27) condition. Student’s t-test were used to determine significance, where *=p<0.05. Images were enhanced for the figure; images were not altered for data analysis. See also Figures S1-4.

We identified 14 additional mutations outside of the low complexity motif that could possibly lead to morphological and functional differences. We identified these mutations using various bioinformatic approaches (**Figure S2, S3**). The stress granule marker G3BP was used to identify non-specific CPEB3 puncta, likely an artifact of protein overexpression. The puncta with G3BP co-label were removed from analysis. S240-242A was the only mutant screened that had a significant difference in morphology and function compared to the full-length protein. We therefore hypothesized that the S240-242A mutation plays an important role in CPEB3 inhibitory function.

To ensure that mutant disruption of inhibitory function is relevant in neurons, we measured the influence of the S240-242A mutant on translation in primary hippocampal cultures. Neurons were transfected with either full-length CPEB3 or the S240-242A mutant, and a SUMO2 3’-UTR Renilla reporter or mutant control (Drisaldi et al., 2015). Neurons were stimulated using chemical long-term potentiation (cLTP) and lysate was collected immediately or at 1 hour. We found that the S240-242A mutant displayed significantly increased translation compared to full-length CPEB3 (**Figure 2C, D**). Mutated reporters failed to express, as expected (**Figure S4**). We verified these results by measuring levels of an endogenous reporter, GluA2, in CPEB3- or S240-242A-expressing neurons. GluA2 levels increase after cLTP in neurons, and CPEB3 is known to regulate GluA2 protein levels (Fioriti et al., 2015). In the basal state, distal GluA2 expression was increased in S240-242A-expressing neurons compared to CPEB3-expressing neurons (**Figure 2E, F**). 1 hour after neuronal stimulation, GluA2 expression increased in both the soma and distal processes of S240-242A-expressing neurons compared to CPEB3-expressing neurons (**Figure 2E, F**). Regardless of stimulation state or method for assaying translation, we found that the S240-242A mutant had significantly decreased inhibitory function compared to full-length CPEB3.

### The CPEB3 low complexity motif drives P body localization

Because the S240-242A mutation is necessary for CPEB3 inhibitory function, we hypothesized that modification of the low complexity motif would also disrupt CPEB3 localization to P bodies. To test this hypothesis, we measured the colocalization of the S240-242A mutant with P bodies (**Figure 3**) using the Colocalization Test in Fiji (Dunn et al., 2011). When the S240-242A mutant is expressed in HeLa cells, the mutant fails to localize to the P body marker DCP1, compared to the full-length protein (**Figure 3A**). When full length CPEB3 or the S240-242A mutant was expressed in primary hippocampal cultures, the mutant failed to localize to DCP1 in the basal state (not shown). Using cLTP, we stimulated the primary hippocampal cultures and measured colocalization to P bodies at the 1 hour timepoint. We found that the mutant also failed to localize to DCP1 in the stimulated state (**Figure 3B**).

**Figure 3.**
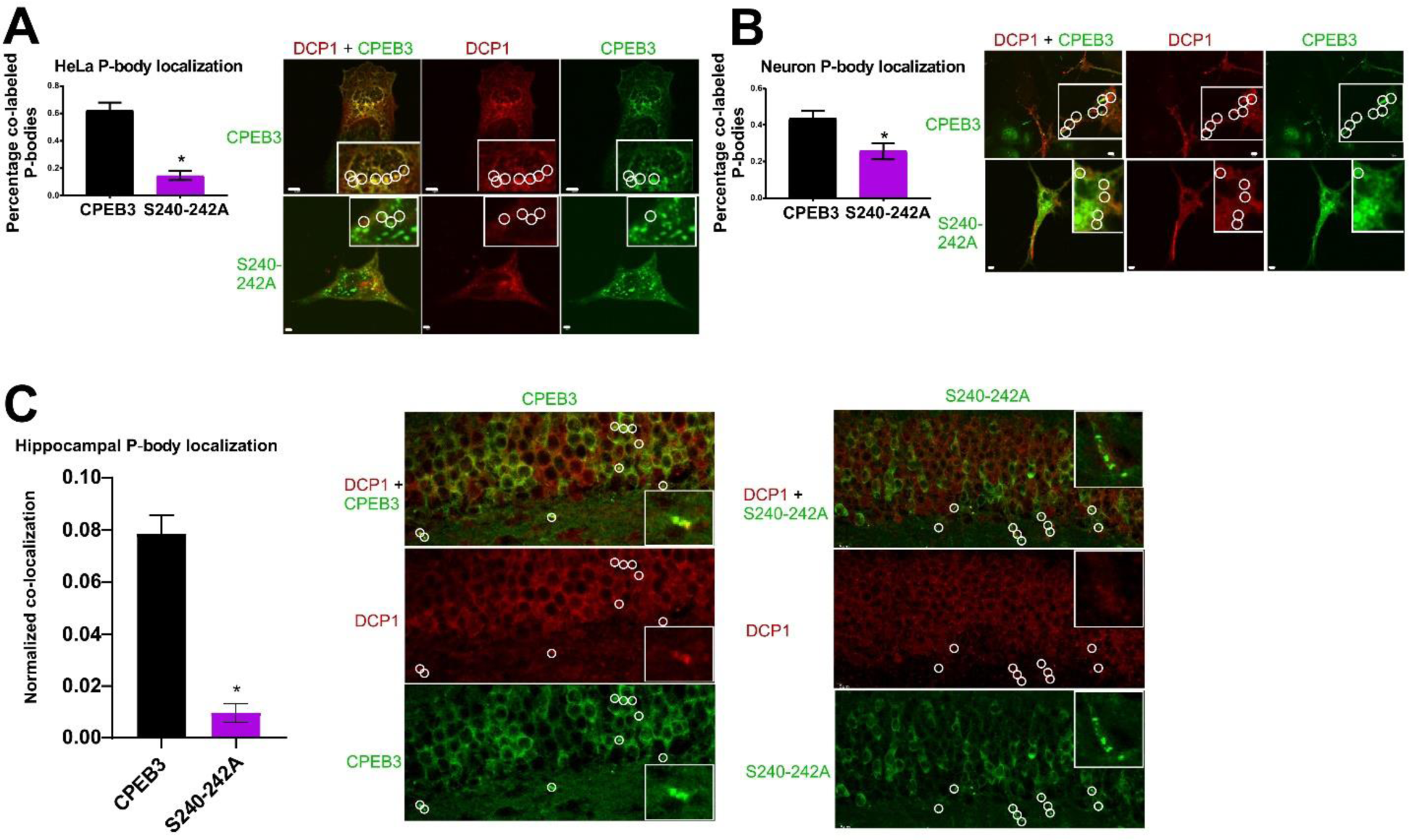
S240-242A does not localize to P bodies. Graphical representation of co-label of P body marker DCP1 and either CPEB3 or the S240-242A mutant in HeLa cells (A; p<0.0001, n=30), stimulated state neurons (B; p=0.0072, n=21), or mouse forebrain (C; p=0.0021, n=8; technical repeats=5). Student’s t-test used, *= significance = p<0.05. Colocalization of DCP1 (red) and CPEB3 or S240-242A (green) in HeLa (A; top scale bars = 5 μm, bottom scale bars = 2 μm), stimulated neurons (B; scale bars = 2 μm), or mouse forebrain (C; scale bars=5 μm). Inserts offer a magnified region of interest. White circles indicate P body location to indicate presence or lack of colocalization. Images were enhanced for the figure; images were not altered for data analysis. See also Figure S5.

Although our findings provide strong support for the necessity of the S240-242A site in localizing CPEB3 to P bodies *in vitro*, we wanted to determine if this was also true *in vivo*. We thus generated two viral constructs under the control of the alpha calcium calmodulin-dependent protein kinase 2 (aCamKII) promoter that contained either the mutated CPEB3 (AAV-DJ8-(0.4)aCamKII-S240-242A-HA-P2A-eGFP) or full-length CPEB3. We injected these constructs bilaterally into the dorsal hippocampus of wild-type mice and sacrificed the animals 2 weeks later. We sectioned and probed hippocampal slices for HA and DCP1. Viral expression was verified with GFP (**Figure S5**). We measured colocalization of HA and DCP1 using the Spots function in Imaris. Consistent with our *in vitro* work, mice expressing full-length CPEB3 exhibited significantly more HA-DCP1 colocalization than mice injected with S240-242A mutant (**Figure 3**).

### The CPEB3 low complexity motif does not influence SUMOylation in neurons

A number of RNA-binding proteins involved in synaptic plasticity (e.g. La and heterogeneous nuclear ribonucleoprotein M and C) are SUMOylated, which encourages mRNA binding in the cytoplasm (Kota et al., 2016; Vertegaal et al., 2004; Vassileva and Matunis, 2004). Since mutation of the low-complexity domain predicted-SIM caused disruption to translation inhibition and to P body localization, we hypothesized that the S240-242A mutant also disrupts SUMOylation of CPEB3. We therefore expressed full-length CPEB3 or the S240-242A mutant in primary hippocampal cultures and collected lysate in the basal or cLTP-stimulated state. CPEB3 was immunoprecipitated from the lysate via its HA tag, and eluate was further immunoprecipitated via SUMO1-, SUMO2-, or SUMO3-binding. We observed no obvious change in SUMOylation across samples (**Figure 4**).

**Figure 4.**
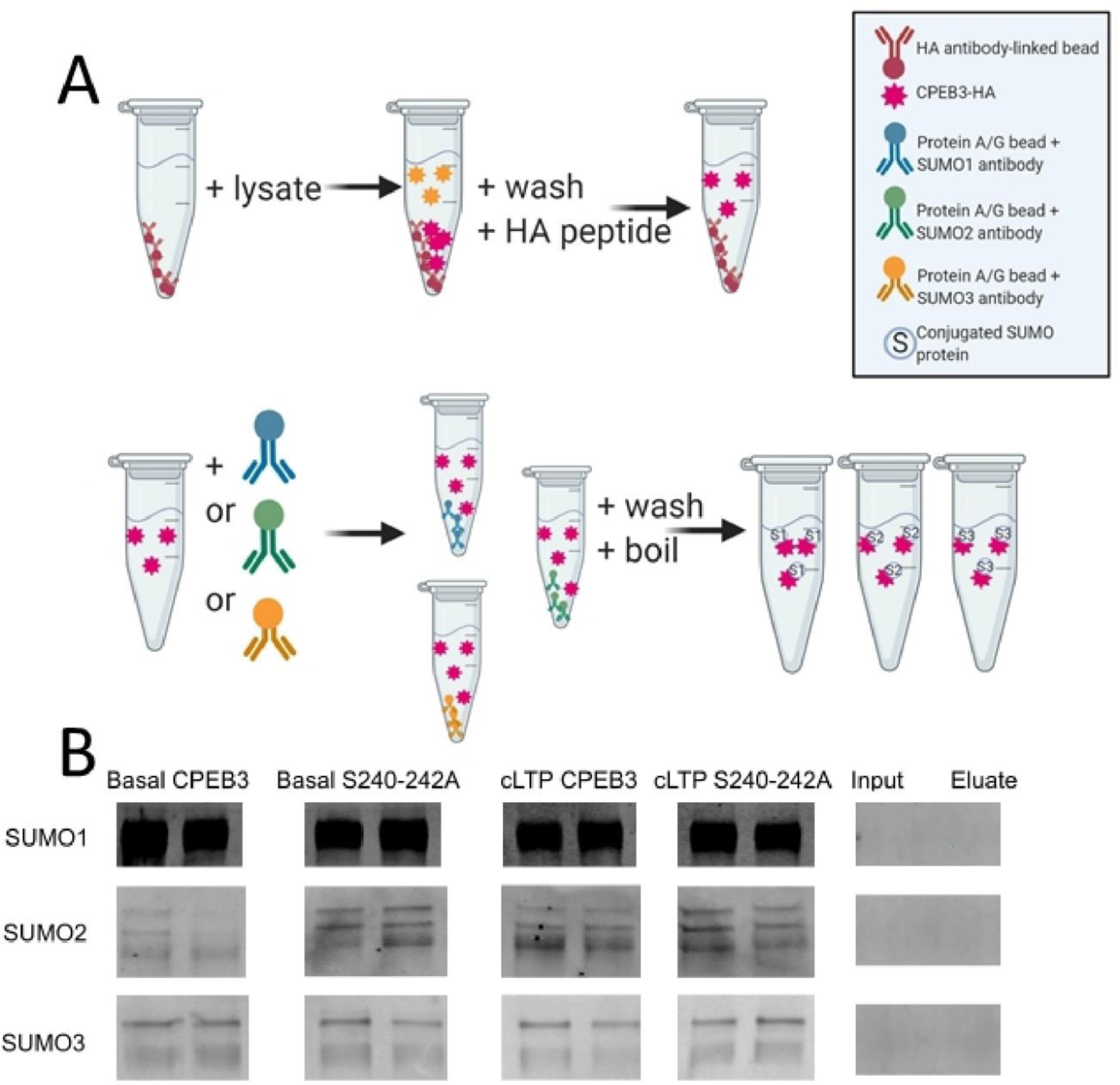
The S240-242A mutant has similar SUMOylation patterns to full-length CPEB3. A. Immunoprecipitation schematic. B. Western blots of SUMO1-, SUMO2-, and SUMO3-immunoprecipitation of CPEB3- or S240-242A-expressing neurons, under basal and stimulated conditions. Samples are noted atop the blots. Molecular weight ∼ 250 kDa.

Thus, the increased translation in cells expressing the S240-242A mutant is not due to SUMOylation modifications. Taken together, these data suggested that a key, unknown mechanism regulates CPEB3 function.

### The CPEB3 interactome identifies potential regulatory binding partners

The question remains: what occurs downstream of CPEB3 SUMOylation before localizing to the P body? We utilized an array of bioinformatic tools (**Figure S3**) and observed high probability of protein-protein interaction in the low complexity motif (**Figure S6**). To investigate the role that protein-protein interactions play in CPEB3-related mRNA transport, we examined the CPEB3 interactome (**Figure 5**). To build an interactome, we expressed CPEB3-HA and CPEB3-GFP in HEK293T cells under a CMV promoter, and immunoprecipitated via the respective tag. Presence of CPEB3 was validated by western blot. We performed on-bead trypsin digestion and measured peptides using mass spectrometry. 1492 proteins were identified in both the CPEB3-HA and CPEB3-GFP samples (**Supplementary Data 1**). CPEB3-HA was also expressed in mouse primary cortical cultures under a human synapsin promoter, immunoprecipitated via the HA tag, and digested on-bead with trypsin before being measured for peptides using mass spectrometry. Across three samples, 2294 proteins were identified (**Supplementary Data 2**). To enrich for the most conserved interactors, we identified 293 proteins from the HEK293T and neuron data combined (**Supplementary Data 3**). We analyzed the data using Strings (Szklarczyk et al., 2019) and identified groups that were mRNA processing or transport-related (red), ribosomal (yellow), actin-related (green), metabolic/ ATP-regulated (cyan), and microtubule-related (blue) (**Figure 5A; Supplementary Table 1**).

**Table 1.** Antibodies used in these studies.

|  | Immunocytochemistry | Immunohistochemistry | Immunoprecipitation |
| --- | --- | --- | --- |
| GFP<br>(abcam<br>cat#ab13970) | 1/1000 | 1/1000 | x |
| HA<br>(Biolegend<br>cat#16B12) | 1/500 | 1/1000 | x |
| G3BP<br>(abcam<br>cat#ab56574) | 1/1000 | x | x |
| DCP1a<br>(abcam<br>cat#ab47811) | 1/1000 | 1/1000 | x |
| GluA2<br>(abcam<br>cat#ab20673) | 1/750 | x | x |
| SUMO1<br>(ENZO cat#BML-PW8330) | x | x | 20 μL |
| SUMO2<br>(Sigma<br>cat#S9571-200UL) | X | X | 16.6 μL |
| SUMO3<br>(Fisher<br>cat#AF2959) | X | X | 26 μL |
| FMRP<br>(abcam<br>cat#ab17722) | 1/1000 | X | X |
| Extracellular<br>GluA2 (alomone<br>cat#AGC-005) | 1/500 | X | X |

**Figure 5.**
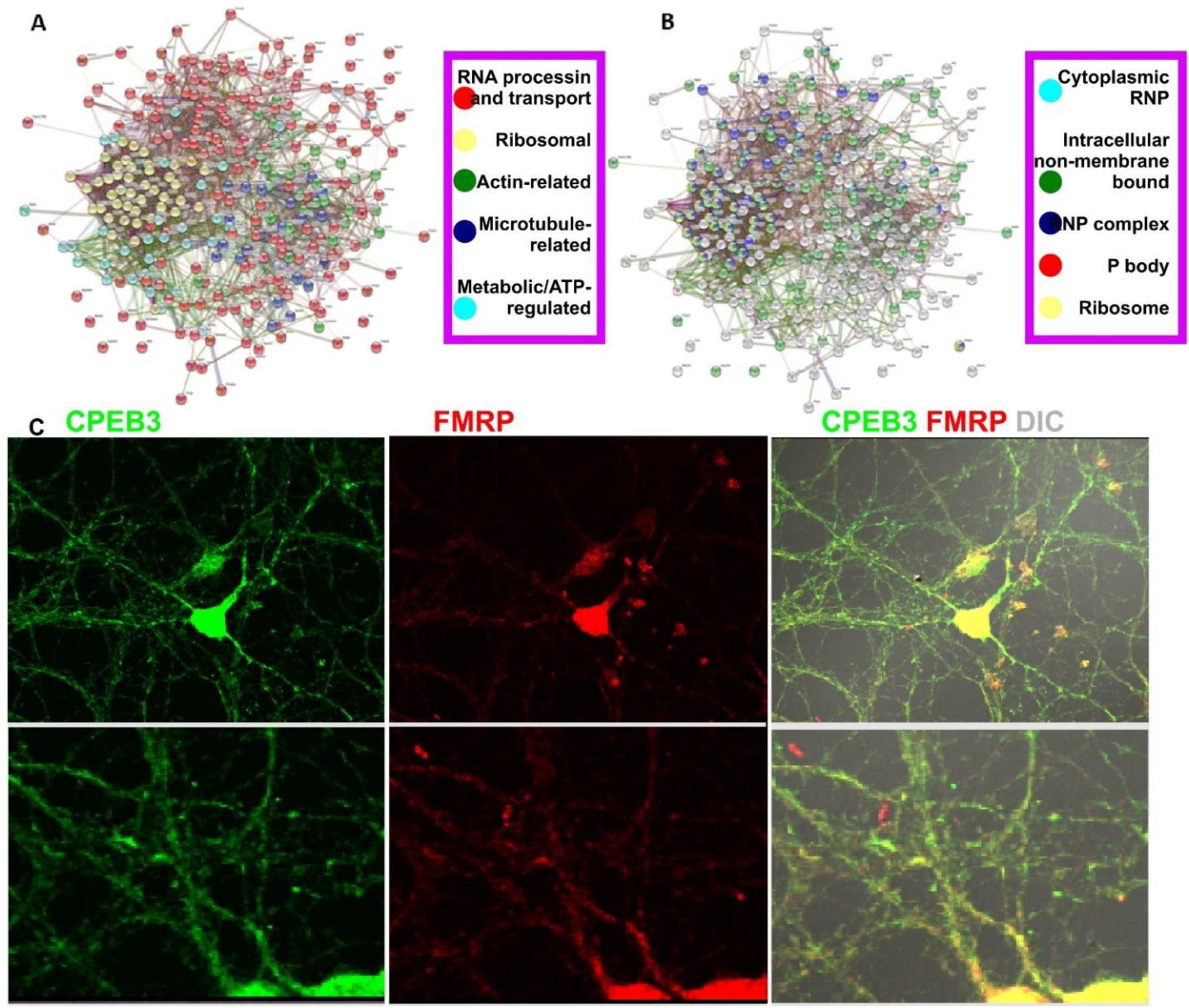
Clustering of the CPEB3 interactome. A. CPEB3 interactome data was clustered using k means with 5 clusters, which identified groups that were mRNA processing or transport-related (red), ribosomal (yellow), actin-related (green), metabolic/ ATP-regulated (cyan), and microtubule-related (blue). B. CPEB3 interactome was clustered using Cellular Component Gene Ontology, and found that CPEB3 interacts with proteins in various RNPs: cytoplasmic ribonucleoprotein granules (cyan), intracellular non-membrane-bounded organelles (green), ribonucleoprotein complexes (blue), and P bodies (red). CPEB3 also interacts with ribosomal proteins (yellow). C. FMRP and CPEB3 colocalization in mouse primary neuronal cultures. Inserts below provide magnified view of FMRP and CPEB3 colocalized puncta. See also Figures S3 and S6.

Gene ontology (GO) analysis using PantherGO (Mi et al., 2013; Thomas et al., 2003) revealed that CPEB3 interacts with proteins in various RNPs: cytoplasmic ribonucleoprotein granules (cyan), intracellular non-membrane-bounded organelles (green), ribonucleoprotein complexes (blue), and P bodies (red). CPEB3 also interacts with ribosomal proteins (yellow) (**Figure 5B; Supplementary Table 1**). Interestingly, P body-related proteins do not overlap with ribosomal proteins in the GO clustering. Previously, our lab discovered that CPEB3 is stored in P bodies in the basal state and leaves the P body by 30 minutes post-cLTP, relocating to the polysome by 1 hour post-cLTP (Ford et al., 2019). Thus, we would not expect to see overlap of CPEB3 interactors in the P body and ribosome clusters, and our interactome data confirms this predication.

From the clustering, it became apparent that CPEB3 interacts with well-defined complexes involved in neuronal mRNA trafficking. We considered four transport methods: 1) Staufen-mediated, 2) zipcode-binding protein-mediated, 3) exon junction complex protein-mediated, and 4) locasome-mediated (Martin and Ephrussi, 2009). The Staufen-mediated method of neuronal mRNA trafficking is the most commonly studied, and the field considers Staufen to be an essential core component of neuronal RNP granule transport (Kiebler and Bassell, 2006). We found that Staufen2 is part of the CPEB3 interactome (**Figure 5; Supplementary Data 3**). Furthermore, the exon junction complex (EJC) protein-mediated method of neuronal mRNA trafficking occurs when EJC proteins bind to mRNAs during splicing; these complexes are found in dendrites and RNP granules (Glanzer et al., 2005; Macchi et al., 2003; Giorgi et al., 2007). We found that EJC proteins eIF4A3 and Mago Nashi are part of the CPEB3 interactome (**Figure 5; Supplementary Data 3**). We did not find zipcode binding proteins or locasome proteins in the CPEB3 interactome. Taken together, the interactome data indicates that CPEB3 is involved in neuronal mRNA trafficking through defined RNP mechanisms.

Finally, we considered the long-term potentiation-relevant CYFIP1-eIF4e-FMR1 complex, which inhibits mRNA translation through the RNA-binding protein FMRP (Napoli et al., 2008). We found that CPEB3 binds all three components, CYFIP1, eIF4e, and FXR1 (**Figure 5; Supplementary Data 3**). We verified the mass spectrometry data by observing FMRP and CPEB3 colocalization in mouse primary neuronal cultures (**Figure 5C**). These data point to an LTP-specific RNP mechanism in which CPEB3 traffics mRNA in neurons.

### The low complexity motif of CPEB3 is necessary for targeted local protein synthesis

The S240-242A mutant no longer localizes to P bodies and no longer exhibits an inhibitory function. We therefore hypothesized that the S240-242A mutant is no longer capable of targeting to specific sites of local protein synthesis, which is a critical component of synaptic plasticity. To investigate this further, we compared the spatial expression of membrane-bound GluA2 in CPEB3- and S240-242A-expressing neurons (**Figure 6A**). We analyzed data using the “spots” and “surface” functions in Imaris, with Matlab XTensions. We found that membrane-bound GluA2 in CPEB3-expressing neurons were located throughout the neuron, where distance was measured as GluA2 spot location from a nuclear surface (**Figure 6B**). However, the S240-242A mutant-expressing neurons contained membrane-bound GluA2 localized much closer to the nuclear surface, suggesting that mutated CPEB3 disrupts appropriate GluA2 trafficking and/or insertion into the membrane (**Figure 6B**). We were able to glean additional information about morphological differences of membrane-bound GluA2 in CPEB3 and the S240-242A mutant expressing neurons. Overall, CPEB3 appears to influence membrane-bound GluA2 expression throughout the neurons, with more intense membrane-bound GluA2 expression (likely from the presence of more GluA2) than the mutant (**Figure 6A-D**). However, the S240-242A mutant severely disrupted membrane-bound GluA2 expression: expression was largely somal, with large elliptical “rafts” of membrane-bound GluA2 forming (**Figure 6A-C,E-F**). Interestingly, total GluA2 expression is increased in S240-242A-expressing neurons (**Figure 2E-F**), but more of the GluA2 appears to be membrane-bound in full-length CPEB3-expressing neurons than in the mutant-expressing neurons (**Figure 6D**). Because of the disruption in expression pattern of membrane-bound GluA2, we find that mutation of the low complexity motif of CPEB3 disrupts appropriate local protein synthesis of mRNA targets.

**Figure 6.**
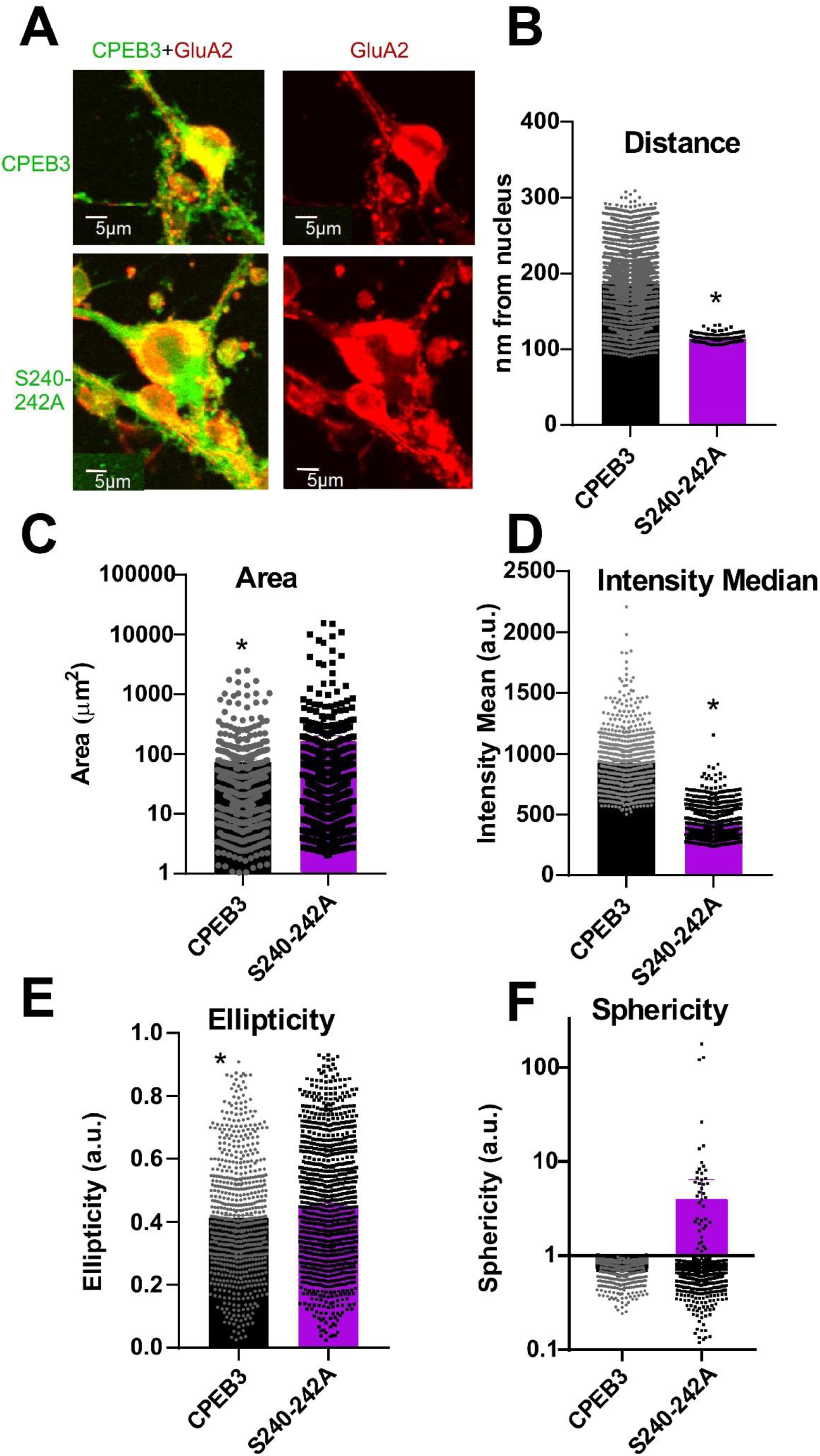
CPEB3 low complexity motif plays a role in targeted local protein synthesis. A. Membrane-bound GluA2 (red) expression in CPEB3 or S240-242A (green)-expressing neurons. B. Distance of membrane-bound GluA2 from nucleus (p<0.005; n= 5; technical replicate= 5). C. Area of membrane-bound GluA2 (p=0.015; n=5; technical replicate =5). D. Median intensity value of membrane-bound GluA2 (p<0.005; n=5; technical replicate= 5). E. Ellipticity of membrane-bound GluA2 (p<0.005; n=5; technical replicate = 5). F. Sphericity of membrane-bound GluA2 (p=0.2462; n=5; technical replicate=5). Student’s t-test determined statistical significance. *= statistical significance if p<0.05.

These data suggest that there is an issue in appropriate trafficking of CPEB3 and its mRNA targets when the S240-242A mutant is present.

## Discussion

Here, we utilize the RNA-binding protein CPEB3 to understand the role that low complexity motifs play in RNP localization and local protein synthesis. This has implications for memory, as CPEB3 is necessary for long-term memory maintenance. Our lab has previously obtained a good spatiotemporal understanding of how neurons regulate CPEB3 function during cLTP. We have now included a critical component of CPEB3 function: protein-protein interaction within the low complexity motif.

We have found that soluble CPEB3 binds mRNA targets and inhibits translation (Fioriti et al., 2015). Soluble, inhibitory CPEB3 is found in P bodies (Ford et al., 2019). The P body-contained CPEB3 is SUMOylated, which is necessary for the inhibitory function of the protein (Drisaldi et al., 2015). Upon cLTP, CPEB3 is deSUMOylated, leaves the P body at a timepoint coinciding with peak SUMO Specific Peptidase I (SENP1) activity (Schorova et al., 2019), and localizes to the polysome (Drisaldi et al., 2015; Ford et al., 2019). This activated CPEB3 promotes translation of its mRNA targets, and is semi-insoluble and oligomeric (Fioriti et al., 2015). We have now found that SUMOylated CPEB3 needs an intact low complexity motif to localize to P bodies. Likely, the low complexity motif is important for dynamic protein-protein interactions.

P bodies have similar structural and functional characteristics as neuronal RNPs (Barbee et al., 2006; Rosario et al., 2016). Thus, P bodies likely act as a “basal state” neuronal RNP in neurons. This provides a great tool for studies of low complexity motif – RNP – local protein synthesis, because P bodies have well-defined components compared to the less well-defined neuronal RNP. Additionally, the neuronal regulation of CPEB3 function is well-defined. Thus, CPEB3 studies in P bodies will help address outstanding questions on the role of local protein synthesis in synaptic plasticity. A comparative protein is FMRP, which is necessary for local protein synthesis, is located in P bodies and RNPs, inhibits translation of mRNA targets, exhibits a reciprocal function when SUMOylated in distal processes, and is required for long-term memory (for review, see Hanus and Schuman, 2013 and Di Liegro et al., 2014; Sudhakaran et al., 2014; Khayachi et al., 2018). Indeed, CPEB4, GIGYF2, Caprin 1, and FXR2 are believed to follow the RNP – mRNA trafficking – local protein synthesis model as well (Tsang et al., 2018).

The CPEB3 interactome contains several RNP, mRNA trafficking, and cytoskeletal proteins. Because of the quantity of interactors that are involved in mRNA trafficking, the low complexity motif is likely involved in binding these families of proteins. To investigate the low complexity motif – RNP – local protein synthesis mechanism, we measured the spatial morphology of membrane-inserted GluA2. GluA2 is a known CPEB3 target, and membrane-bound GluA2 is known to increase in synaptic sites as a result of LTP (for review, see Hanus and Schuman, 2013). We found that disrupting the low complexity motif of CPEB3 resulted in disrupted membrane-bound GluA2 expression. We therefore conclude that CPEB3 localization to P bodies is necessary for appropriate local protein synthesis, likely due to an mRNA-trafficking-based protein interaction with the low complexity motif.

## Supporting information

Key Resources Table

## Acknowledgements

We thank the Howard Hughes Medical Institute for funding this work. We thank the Cellular Imaging Facility of the Zuckerman Mind, Brain, Behavior Institute of Columbia University, and the Proteomics and Macromolecular Crystallography Shared Resource of Columbia University Irving Medical Center.

## Author Contributions

Conceptualization, L.F. Methodology, L.F, A.A. Validation, L.F. Formal analysis, L.F. Investigation, L.F., A.A, A.D.T, C.P., M.F., N.S., L.Fioriti, R.K.S. Resources, L.F., L. Fioriti, R.K.S., E.R.K. Writing-Original Draft, L.F. Writing-Review and Editing, all authors. Visualization, L.F. Supervision, L.F. Project Administration, L.F. Funding Acquisition, R.K.S, E.R.K.

## The authors declare no competing interests

## STAR Methods

### Lead Contact

Further information and requests for resources and reagents should be directed to and will be fulfilled by the Lead Contact, Eric R. Kandel.

### Materials Availability

Materials are available with an appropriate MTA, please contact the Lead Contact. hSyn-tdTomato and aCaMKII-eGFP are not available, due to MTA limitations.

### Data and Code Availability

The published article includes all datasets generated or analyzed during this study. Method Details

## Antibodies

See **Table 1**

### Animals: *Animal maintenance*

Wild-type (C57BL/6J background) male mice were obtained from Jackson Laboratory (Bar Harbor, ME) at approximately 9-10 weeks of age. Animals were housed in the Zuckerman Institute vivarium at Columbia University and maintained on a standard 12-h 12-h light-dark cycle with *ad libitum* access to food and water. All animal procedures were conducted in accordance with the Institutional Animal Care and Use Committee (IACUC) at Columbia University.

### Animals: *Viral Surgery*

For all surgeries, mice were first anesthetized with isoflurane and administered a weight-appropriate dose of Bupivacaine, a local anesthetic, subcutaneously at the surgical site and an intraperitoneal dose of Carprofen for pain. A bilateral craniotomy was made directly above the dorsal hippocampus. Next, a glass micropipette attached to a Nanoject III (Drummond Scientific, Bromall, PA) was lowered to a region just below the dorsal blade of CA1 in the dorsal hippocampus (AP = ±1.4, ML = -2.3, and DV = -2.1). 500 nL of either AAV-DJ8-(0.4)aCamKII-CPEB3-HA-P2A-eGFP or AAV-DJ8-(0.4)aCamKII-S240-242A-HA-P2A-eGFP was injected at a rate of 100 nL/min for 5 min. Following surgery, mice were allowed to recover for two weeks.

### Animals: *Immunohistochemistry*

Animals were anesthetized with a mixture of ketamine and xylazine (200/20 mg/mL). Animals were then transcardially perfused with 10mL 1x PBS and 10mL 4% paraformaldehyde. Brains were extracted and post-fixed for 24h in 4% paraformaldehyde. Brains were washed twice for 30-minutes in 1x PBS, before being stored in 1x PBS + 0.02% NaN_3_.

Brains were sliced coronally at 40 μm on a vibratome (Leica VT1200s). Forebrain tissue was collected in 1x PBS.

Tissue was washed for 10 min in 300 μl of 1x TBS, and then washed 3 times in 300 μl of TBST (TBS + 0.05% Triton-x). Tissue was placed in 300 μl of 1x Antigen Unmasking Solution (Citric Acid Based, H-3300, (Vector Laboratories)) and loaded into an oven at 70°;C for 1h. Samples cooled to room temperature before being washed 3 times in 300 μl of TBST. Tissue was placed in a blocking buffer consisting of 2.5% of Normal Goat Serum, 2.5% of Normal Donkey Serum, 1% BSA and 94% TBST overnight. Tissue was stained with a combination of primary antibodies in blocking buffer, again, overnight. Finally, tissue was washed 4 times for 10min with 300 μl TBST. Tissue was incubated in blocking buffer and secondary antibodies for 2h at room temperature. Tissue was then washed 4 last times for 10minutes in TBST. Tissue was rinsed briefly with 70% EtOH.

Tissue was mounted using Prolong Gold Antifade Reagent on Superplus Frost Slides and let dry in the dark for 24h before being imaged.

### Cell culture: *Cell lines*

HEK293T, HeLa, and N2A cells were cultured in Dulbecco’s modified Eagle’s medium with high glucose and L-glutamine (Gibco) with 10% FBS (Sigma) and 1% penicillin/streptomycin (Gibco) in a tissue-culture incubator at 37°C and 5% CO_2_. Cells were plated in T25 or T75 flasks and grown to 40-60% confluency before transfection. Additionally, HEK293T were plated on 12-mm glass coverslips coated with poly-L-lysine (Sigma) in 24-well flat-bottom cell-culture plates.

HEK293T and HeLa cells were transfected with TransFast Transfection Reagent (Promega) – 500 μg DNA per 24-well culture plate. N2A cells were transfected with 2 μL Lipofectamine 3000 (ThermoFisher) + 800 ng DNA per 24-well in Opti-MEM (ThermoFisher). N2As incubated in transfection media for 1 hour and then changed to fresh complete DMEM (ThermoFisher). Cells expressed for 24 hours before lysate was collected.

Cells were transfected with: CPEB3-GFP (Fioriti et al., 2015), S240-242A-GFP [a mutation of CPEB3-GFP], CPEB3-HA (Fioriti et al., 2015), S240-242A-HA [a mutation of CPEB3-HA], Actin 3’UTR-Renilla (Stephan et al., 2015), Sumo 2 3’UTR-Renilla (Drisaldi et al., 2015), mutant Sumo 2 3’UTR-Renilla (Drisaldi et al., 2015).

### Cell culture: *Primary neuronal culture*

The primary neuron isolation and culture methods were adapted from Kaech and Banker, 2006. Hippocampi and neocortices were dissected from P0-P2 mouse pups, and isolated from meningeal tissue in an ice-cold dish of Hank’s Balanced Salt Solution. In a 15 mL Falcon tube prepared with warm papain dissociation media (Worthington Biochemical Corporation), large pieces of tissue were mechanically dissociated with forceps and then incubated in the media for 40-50 minutes at 37°C and 5% CO_2_. The tubes were agitated every 15 minutes. Tissue was allowed to settle to the bottom of the tube and then transferred with a 1 mL pipette into another 15 mL tube prepared with ovomucoid protease inhibitor solution (Worthington) and incubated for 10 minutes at room temperature. Tissue was again allowed to settle and transferred to a third tube prepared with warm Dulbecco’s modified Eagle’s medium (Gibco) containing 10% horse serum (Gibco) and 1.6% glucose (Sigma). Tissue was gently triturated in this media with a Pasteur pipette 15 times, allowed to settle, and then the supernatant was transferred to a final Falcon tube for dissociated cell collection. The trituration process was repeated three times by adding warm MEM to the remaining tissue, triturating, and transferring the supernatant into the collection tube. At the end of the process, dissociated cells in the collection tube were centrifuged at 1000 g for 5 minutes, resuspended in warm MEM and plated at 300k cells/mL in 24- and 12-well cell-culture plates pretreated with poly-L-lysine (1mg/mL, Sigma). For imaging samples, isolated neurons were plated on cleaned and poly-L-lysine pretreated coverslips in multi-well cell-culture plates. Plated cells were incubated in the MEM for 4 hours at 37°C and 5% CO_2_. After this incubation the MEM was carefully aspirated from each well and immediately replaced with warm Neurobasal media containing B27 supplement and GlutaMax (Gibco). One third of the media was removed and replaced with fresh Neurobasal media every 4-5 days. Neurons were cultured for up to 21 days at 37°C and 5% CO_2_. Cells were transfected at DIV 5 with Lipofectamine 3000 (ThermoFisher) and 1 μg DNA per 2-cm^2^ culture well. Or, cells were transduced with a lentivirus (modified from Malcolm Moore, Addgene plasmid #48687, RRID: Addgene_48687) and LentiX Accelerator (Takara Bio). Cells expressed for at least 24 hours before being collected.

Cells were transfected with: CPEB3-GFP (Fioriti et al., 2015), S240-242A-GFP [a mutation of CPEB3-GFP], CPEB3-HA (Fioriti et al., 2015), S240-242A-HA [a mutation of CPEB3-HA], synapsin-CPEB3-HA-tdTomato, synapsin-S240-242A-HA-tdTomato, Actin 3’UTR-Renilla (Stephan et al., 2015), Sumo 2 3’UTR-Renilla (Drisaldi et al., 2015), mutant Sumo 2 3’UTR-Renilla (Drisaldi et al., 2015).

### Cell culture: *Chemical long-term potentiation*

DIV 15 neurons were placed in fresh Neurobasal media for 3 minutes and then moved to stimulation media for 5 minutes. Stimulation media: 50 μM forskolin, 0.1 μM rolipram, 50 μM picrotoxin in ACSF without MgCl. Neurons were moved back to conditioned media.

### Cell culture: *Immunocytochemistry*

Cells were washed with cold PBS and fixed in-plate with 4% PFA in PBS (neuronal cultures additional 5% sucrose). Cells were permeabilized for 2 minutes with 0.1% Triton-X in PBS, blocked for 1 hour at room temperature in 5% FBS in PBS, probed overnight at 4°C with primary antibody in block, and probed for 1.5 hours at room temperature with secondary antibody in block. Coverslips were mounted using Fluoroshield (Sigma).

### Mutagenesis

A plasmid containing mCPEB3 in the pAcGFP1-N1 vector (Fioriti et al., 2015) was used as a template for site-directed mutagenesis using the Q5® Site-Directed Mutagenesis Kit (NEB). Complementary primers containing the desired deletion or substitution were annealed to the template plasmid and the Q5 Hot Start High-Fidelity DNA Polymerase (NEB) replicated and incorporated the mutant primers into the template. The PCR mix was digested with Dpn1 to fragment the template DNA, and then transformed into XL10-Gold Ultracompetent Cells (Agilent) to be grown overnight and have single colonies picked for DNA extraction using the QIAprep Spin Miniprep Kit. The correct DNA sequence of all mutant constructs was confirmed by DNA sequencing. Mutants were transformed into NEB^®^ 5-alpha Competent E. coli (High Efficiency) and then the DNA extracted using the QIAGEN Plasmid Maxi Kit.

Splice C:

AAATTTATTCCCATGCGACCTTTCAGAGGCTCATGGTC GACCATGAGCCTCTGAAAGGTCGCATGGGAATAAATTT

Splice B: C

TTCAAAGGGAAAGAGAGAAGAGGTGTTAAGTGCCATAATGTTA TAACATTATGGCACTTAACACCTCTTCTCTCTTTCCCTTTGAAG

S194A:

GCTGGCGGGCGCGCGGCGTTGCT

AGCAACGCCGCGCGCCCGCCAGC

S197A:

CCTGGCTGGGGGCGGCGGGCGAGC

GCTCGCCCGCCGCCCCCAGCCAGG

S237-238A:

GGACGAGGCCGCAGCGGCAGCGGCCG

CGGCCGCTGCCGCTGCGGCCTCGTCC

S240-242A:

GCTGCCTCTTCGGCCGCGGCCGCCTGGAACACGCACCA

TGGTGCGTGTTCCAGGCGGCCGCGGCCGAAGAGGCAGC

S349A:

CATTAAGGAGTTCTCCAAGGCGTGCAAGTTAAAAGTGTCAT

ATGACACTTTTAACTTGCACGCCTTGGAGAACTCCTTAATG

Del_N:

Published previously, see Fioritit et al., 2015 Del_RBD:

GAACGAGTGGAACGCTACTCTCCGTACGTGCT

AGCACGTACGGAGAGTAGCGTTCCACTCGTTC

S419-420A:

CAAAGGGAAAGAGAGCAGCCCGACCTCGTCTCCG.

CGGAGACGAGGTCGGGCTGCTCTCTTTCCCTTTG

S444A:

AGGAGCAGCTAAGCC

GGCTTAGCTGCTCCT

Y457A/S458A:

CCAACAAACACCTTTCTAGCGGCGCGTTCCACTCGTTCCCCA

TGGGGAACGAGTGGAACGCGCCGCTAGAAAGGTGTTTGTTGG

K459A/R460A:

GGCCTCCAACAAACACCGCTGCAGAGTAGCGTTCCACTCGTTCCC

GGGAACGAGTGGAACGCTACTCTGCAGCGGTGTTTGTTGGAGGCC

S222A:

GCCGAGGATGAGGCCGGCTTGCTGGTC

GACCAGCAAGCCGGCCTCATCCTCGGC

S223A:

CCGCCGAGGATGCGGACGGCTTGCT

AGCAAGCCGTCCGCATCCTCGGCGG

S224A:

CCAGCAAGCCGTCCTCAGCCTCGGCGG

CCGCCGAGGCTGAGGACGGCTTGCTGG

S225A:

CGTCCTCATCCGCGGCGGTCGCG

CGCGACCGCCGCGGATGAGGACG

S237A:

CGGCCGCTGCCGCTTCGGCCTCG

CGAGGCCGAAGCGGCAGCGGCCG

S238A:

CGAGGCCGCAGAGGCAGCGGCCGC

GCGGCCGCTGCCTCTGCGGCCTCG

S240A:

GCCTCTTCGGCCGCGTCCAGCTGGA

TCCAGCTGGACGCGGCCGAAGAGGC

S242A:

GTGTTCCAGCTGGCCGAGGCCGAAGAG

CTCTTCGGCCTCGGCCAGCTGGAACAC

### Translational assay

Double transfected cells (a CPEB3 and Renilla transfection) were washed in cold PBS, scraped, and lysed in the lysis buffer provided in the Dual-Glo Luciferase Assay System (Promega). Lysate luminosity was measured with a luminometer. Background luminosity was measured with Luciferase substrate (Firefly; Promega). Experimental luminosity was measured with Stop & Glo substrate (Renilla; Promega). Luminosity was normalized to protein concentration.

### Confocal imaging and analysis

All images were acquired on an Olympus IX81 laser-scanning confocal microscope using the FluoView FV1000 Microscopy System. Cell culture images were quantified using SyNPanal (Danielson et al., 2014) or, ImageJ/Fiji integrated density measurements and Colocalization Test (Dunn et al., 2011). Tissue images and spatial GluA2 images were quantified using the “spots” and “surfaces” functions, and Matlab XTension “find spots near surface”, in Imaris. Each spot/surface selection was verified by eye. Access to Imaris software was provided by Cellular Imaging at Zuckerman Mind, Brain, Behavior Institute.

### Biochemistry: *Sample preparation and immunoprecipitation*

Cells were washed with cold PBS and lysed in RIPA + NEM (n-ethylmaleimide) buffer. Lysate was added to either 50 μL GFP-Trap magnetic bead (Chromotek) slurry or 50 μL HA-linked magnetic bead (Pierce) slurry. Lysate bound to magnetic beads overnight at 4°C with end-over-end mixing. Beads were washed 3x with GFP-Trap Wash Buffer (Chromotek). Washed GFP-Trap beads were boiled in 2x Laemelli Buffer (Bio-Rad) and immediately loaded on to a 4-20% Tris-glycine SDS-PAGE gel (Bio-Rad). Washed HA beads were mixed with a 4:1 column volume HA peptide (Sigma Aldrich) for 15 min at 37°C. Eluate was further processed for western blotting (addition of 4x Laemmeli [Bio-Rad] and boiling) or additional immunoprecipitation.

### Biochemistry: *SUMO immunoprecipitation*

Protein A/G magnetic beads (ThermoFisher) were incubated with either SUMO1 antibody (ENZO; 26 μL), SUMO2 antibody (Sigma Aldrich; 16.6 μL), or SUMO3 antibody (LifeTechnologies; 20 μL) and bound for 1 hr in GFP-Trap Wash Buffer (Chromotek) at 37°C with mixing. Unbound antibody was washed from the beads 3x with GFP-Trap Wash Buffer. SUMO-conjugated beads were then bound to HA-immunoprecipitation eluate overnight at 4°C with end-over-end mixing. Beads were washed 3x with GFP-Trap Wash Buffer (Chromotek). Washed GFP-Trap beads were boiled in 2x Laemelli Buffer (Bio-Rad) and immediately loaded on to a 4-20% Tris-glycine SDS-PAGE gel (Bio-Rad).

### Biochemistry: *Western blotting*

Boiled samples were loaded on to a 4-20% Tris-glycine SDS-PAGE gel (Bio-Rad) and run at 120 V until the dye face reached the bottom of the gel. Gels were then wet transferred to PVDF 0.45 μm membrane overnight at 20 V, 200 mA / gel. Membranes were blocked in Odyssey Blocking Buffer (Li-Cor) 1:1 TBST, and probed with either a GFP or HA antibody in block overnight at 4°C with mixing. Membranes were washed in TBST, probed with an appropriate IR680 or IR800-conjugated secondary antibody (1/5000; Li-Cor), and imaged on a Li-Cor Odyssey Infrared Imager.

### Mass Spectrometry: *Preparation of samples*

Proteins bound to streptavidin beads were washed five times with 200 µl of 50 mM ammonium bicarbonate and subjected to disulfide bond reduction with 5 mM TECP (room temperature, 30 min) and alkylation with 10 mM iodoacetamide (room temperature, 30 min in the dark). Excess iodoacetamide was quenched with 5 mM DTT (room temperature, 15 min). Proteins bound on beads were digested overnight at 37°C with 1 µg of trypsin/LysC mix. The next day, digested peptides were collected in a new microfuge tube and digestion was stopped by the addition of 1% TFA (final v/v), and centrifuged at 14,000 g for 10 min at room temperature. Cleared digested peptides were desalted on SDB-RP Stage-Tip and dried in a speed-vac. Peptides were dissolved in 3% acetonitrile/0.1% formic acid.

### Mass Spectrometry: *LC-MS/MS analysis*

LC-MS/MS was performed using a Waters NanoAcquity M-class system coupled to a Thermo Scientific Q Exactive HF mass spectrometer. Thermo Scientific EASY-Spray 50 cm × 75 µm ID length C18 column were used to separate desalted peptides with a 5-30% acetonitrile gradient in 0.1% formic acid over 70 min at a flow rate of 300 nL/min. After each gradient, the column was washed with 90% buffer B (0.1% formic acid, 100% HPLC-grade acetonitrile) for 5 min and re-equilibrated with 98% buffer A (0.1% formic acid, 100% HPLC-grade water) for 40 min.

MS data were acquired in data dependent acquisition (DDA) mode with an automatic switch between a full scan and 15 data-dependent MS/MS scans (TopN method). Target value for the full scan MS spectra was 3 x 10^6^ ions in the 375-1600 m/z range with a maximum injection time of 100 ms and resolution of 60,000 at 200 m/z with data collected in profile mode. Precursors were selected using a 1.6 m/z isolation width. Precursors were fragmented by higher-energy C-trap dissociation (HCD) with normalized collision energy of 27 eV. MS/MS scans were acquired at a resolution of 15,000 at 200 m/z with an ion target value of 5 x 10^5^, maximum injection time of 50 ms, dynamic exclusion for 15 s and data collected in centroid mode.

### Mass Spectrometry: *Data analysis*

Raw mass spectrometric data were analyzed using MaxQuant (Cox and Mann, 2008) v.1.6.1.0 and employed Andromeda for database search (Cox et al., 2011) at default settings with a few modifications. The default was used for first search tolerance and main search tolerance: 20 ppm and 6 ppm, respectively. MaxQuant was set up to search the reference mouse proteome database downloaded from Uniprot. MaxQuant performed the search trypsin digestion with up to 2 missed cleavages. Peptide, Site and Protein FDR were all set to 1% with a minimum of 2 peptide needed for Identification but 2 peptides needed to calculate a protein level ratio. The following modifications were used as fixed carbamidomethyl modification of cysteine, and oxidation of methionine (M), Deamination for asparagine or glutamine (NQ) and acetylation on N-terminal of protein were used as variable modifications. MaxQuant combined folder was uploaded in scaffold for data visualization.

### Quantification and Statistical Analysis

Specific statistical analyses are indicated in figure legends. Mass spectrometry data analysis is indicated in the appropriate methods section. Graphs represent mean +/- SEM. Statistical analyses were performed using GraphPad Prism 7. All data were analyzed using Student’s t-test or one-way ANOVA, with Dunn’s multiple comparison tests. Statistical significance was designated as p<0.05.

### Supplemental titles and legends

Supplementary Data 3. CPEB3 interactome components

