## Supplementary material for "The low complexity motif of cytoplasmic polyadenylation element binding protein 3 (CPEB3) is critical for the trafficking of its targets in neurons": Key Resources Table

| REAGENT or RESOURCE | SOURCE | IDENTIFIER |
|---|---|---|
| Antibodies | | |
| GFP | Abcam | cat#ab13970 |
| HA | Biolegend | cat#16B12 |
| G3BP | Abcam | cat#ab56574 |
| DCP1a | Abcam | cat#ab47811 |
| GluA2 | Abcam | cat#ab20673 |
| SUMO1 | ENZO | cat#BML-PW8330 |
| SUMO2 | Sigma | cat#S9571-200UL |
| SUMO3 | Fisher Scientific | cat#AF2959 |
| FMRP | abcam | cat#ab17722 |
| Extracellular GluA2 | Alomone | cat#AGC-005 |
| Bacterial and Virus Strains | | |
| AAV-DJ8-(0.4)aCamKII-CPEB3-HA-P2A-eGFP | This manuscript | N/A |
| AAV-DJ8-(0.4)aCamKII-S240-242A-HA-P2A-eGFP | This manuscript | N/A |
| Critical Commercial Assays | | |
| Q5® Site-Directed Mutagenesis Kit | NEB | E0554 |
| Dual-Glo Luciferase Assay System | Promega | E2920 |
| Deposited Data | | |
| Supplementary Data 1 | This manuscript | N/A |
| Supplementary Data 2 | This manuscript | N/A |
| Supplementary Data 3 | This manuscript | N/A |
| Experimental Models: Cell Lines | | |
| HEK293T | ATCC | CRL-11268 |

| | | |
|---|---|---|
| HeLa | ATCC | CRL-2 |
| N2A | ATCC | CCL-131 |
| Experimental Models: Organisms/Strains | | |
| C57BL/6J mice | Jackson Lab | 000664 |
| Oligonucleotides | | |
| Splice C:<br>AAATTTATTCCCATGCGACCTTTCAGAGGCTCATG GTC<br>GACCATGAGCCTCTGAAAGGTCGCATGGGAATAA ATTT | This manuscript | N/A |
| Splice B: C<br>TTCAAAGGGAAAGAGAGAAGAGGTGTTAAGTGC CATAATGTTA<br>TAACATTATGGCACTTAACACCTCTTCTCTCTTTCC CTTTGAAG | This manuscript | N/A |
| S194A:<br>GCTGGCGGGCGCGCGGCGTTGCT<br>AGCAACGCCGCGCGCCCGCCAGC | This manuscript | N/A |
| S197A:<br>CCTGGCTGGGGGCGGCGGGCGAGC<br>GCTCGCCCGCCGCCCCCAGCCAGG | This manuscript | N/A |
| S237-238A:<br>GGACGAGGCCGCAGCGGCAGCGGCCG<br>CGGCCGCTGCCGCTGCGGCCTCGTCC | This manuscript | N/A |
| S240-242A:<br>GCTGCCTCTTCGGCCGCGGCCGCCTGGAACACGC ACCA<br>TGGTGCGTGTTCCAGGCGGCCGCGGCCGAAGAG GCAGC | This manuscript | N/A |
| S349A:<br>CATTAAGGAGTTCTCCAAGGCGTGCAAGTTAAAA GTGTCAT<br>ATGACACTTTTAACTTGCACGCCTTGGAGAACTCCTTAATG | This manuscript | N/A |
| Del_N | Fioriti et al., 2015 (Kandel Lab) | N/A |
| Del_RBD:<br>GAACGAGTGGAACGCTACTCTCCGTACGTGCT<br>AGCACGTACGGAGAGTAGCGTTCCACTCGTTC | This manuscript | N/A |

| | | |
|---|---|---|
| S419-420A:<br>CAAAGGGAAAGAGAGCAGCCCGACCTCGTCTCCG<br>.<br>CGGAGACGAGGTCGGGCTGCTCTCTTTCCCTTTG | This manuscript | N/A |
| S444A:<br>AGGAGCAGCTAAGCC<br>GGCTTAGCTGCTCCT | This manuscript | N/A |
| Y457A/S458A:<br>CCAACAAACACCTTTCTAGCGGCGCGTTCCACTCG<br>TTCCCCA<br><br>TGGGGAACGAGTGGAACGCGCCGCTAGAAAGGTGTTTGT<br>TGG | This manuscript | N/A |
| K459A/R460A:<br>GGCCTCCAACAAACACCGCTGCAGAGTAGCGTTC<br>CACTCGTTCCC<br><br>GGGAACGAGTGGAACGCTACTCTGCAGCGGTGTTTGTTGG<br>AGGCC | This manuscript | N/A |
| S222A:<br>GCCGAGGATGAGGCCGGCTTGCTGGTC<br>GACCAGCAAGCCGGCCTCATCCTCGGC | This manuscript | N/A |
| S223A:<br>CCGCCGAGGATGCGGACGGCTTGCT<br>AGCAAGCCGTCCGCATCCTCGGCGG | This manuscript | N/A |
| S224A:<br>CCAGCAAGCCGTCCTCAGCCTCGGCGG<br>CCGCCGAGGCTGAGGACGGCTTGCTGG | This manuscript | N/A |
| S225A:<br>CGTCCTCATCCGCGGCGGTCGCG<br>CGCGACCGCCGCGGATGAGGACG | This manuscript | N/A |
| S237A:<br>CGGCCGCTGCCGCTTCGGCCTCG<br>CGAGGCCGAAGCGGCAGCGGCCG | This manuscript | N/A |
| S238A:<br>CGAGGCCGCAGAGGCAGCGGCCGC<br>GCGGCCGCTGCCTCTGCGGCCTCG | This manuscript | N/A |
| S240A:<br>GCCTCTTCGGCCGCGTCCAGCTGGA<br>TCCAGCTGGACGCGGCCGAAGAGGC | This manuscript | N/A |
| S242A:<br>GTGTTCCAGCTGGCCGAGGCCGAAGAG<br>CTCTTCGGCCTCGGCCAGCTGGAACAC | This manuscript | N/A |
| Recombinant DNA | | |

| | | |
|---|---|---|
| CPEB3-GFP | Fioriti et al., 2015 (Kandel Lab) | N/A |
| S240-242A-GFP | This Manuscript | N/A |
| CPEB3-HA | Fioriti et al., 2015 (Kandel Lab) | N/A |
| S240-242A-GFP | This Manuscript | N/A |
| Actin 3’ UTR-Renilla | Stephan et al., 2015 (Kandel Lab) | N/A |
| Sumo 2 3’ UTR- Renilla | Drisaldi et al., 2015 (Kandel Lab) | N/A |
| Mutant Sumo 2 3’ UTR-Renilla | Drisaldi et al., 2015 (Kandel lab) | N/A |
| synapsin-CPEB3-HA-tdTomato | This Manuscript | N/A |
| synapsin-S240-242A-HA-tdTomato | This Manuscript | N/A |
| Software and Algorithms | | |
| SyNPAnal | Danielson et al., 2014 | N/A |
| Fiji/ImageJ | NIH.gov | N/A |
| Imaris | Cellular Imaging at Zuckerman Institute/ Oxford Instruments | 9.5 |
| Scaffold | Proteome Software | 7 |
| GraphPad Prism | GraphPad | 7 |
| Strings | Strings-db | 11.0 |
| PantherGO | Pantherdb | 15.0 |
