## Supplementary Table 1 for "The low complexity motif of cytoplasmic polyadenylation element binding protein 3 (CPEB3) is critical for the trafficking of its targets in neurons"

Supplementary Table 1. Detailed Strings data.

| Protein ID | A color code | B color code |
|---|---|---|
| Magohb | red | blue |
| Ktn1 | red | |
| Bzw1 | red | |
| Tardbp | red | |
| Clasp1 | blue | green |
| Naa15 | red | |
| Atp6v1h | red | |
| Cnn3 | red | green |
| Fxr1 | red | green, blue |
| Eif4e | yellow | cyan, green, blue, red |
| Ubap2l | red | |
| Ctbp1 | red | |
| Lrp1 | red | |
| Cyfip1 | red | |
| Cpne2 | red | |
| Dbn1 | red | |
| Tmem33 | cyan | |
| Etfb | red | |
| Atxn2l | red | cyan, blue, green |
| Tjp1 | red | green |
| Rpl18a | yellow | yellow, green, blue |
| Tln2 | red | green |

| | | |
|---|---|---|
| Rps25 | yellow | yellow, green, blue |
| Idh3a | cyan | |
| Chordc1 | red | |
| Rps12 | yellow | yellow, green, blue |
| Atp2b1 | red | green |
| Psmd6 | green | |
| Anxa7 | red | |
| Actn1 | red | green |
| Gatad2b | red | |
| Kpna2 | red | cyan, blue, green |
| Sept9 | red | green |
| Eif3m | yellow | |
| Srp68 | yellow | green, blue |
| Rpn2 | cyan | |
| Eif4a3 | yellow | blue, green |
| Hsd17b10 | red | green, blue |
| Agps | red | green |
| Nckap1 | red | green |
| Rrbp1 | red | yellow, green, blue |
| Dync1i2 | blue | |
| Xpo5 | red | green |
| Clasp2 | blue | green |
| Ywhaq | red | |
| Acaca | red | green |
| Acly | yellow | |

| | | |
|---|---|---|
| Actr1a | blue | green |
| Slc25a4 | red | |
| Ago2 | red | cyan, green, blue, red |
| Ina | red | green, blue, cyan |
| Rnpep | red | |
| Apob | red | |
| Aprt | cyan | |
| Araf | red | |
| Arf5 | blue | |
| Arl1 | red | |
| Atrx | red | green |
| Acadvl | red | green |
| Macf1 | red | green |
| Lrrc47 | cyan | |
| Prdx1 | red | green |
| Mylk | red | green |
| Gmps | cyan | |
| Actb | red | green, blue, cyan |
| Parp14 | red | |
| Strap | red | |
| Wdr82 | red | green |
| Cnot1 | red | green, blue, cyan, red |
| Cbr1 | red | |
| Ddx3x | red | blue, cyan, green |
| Eprs | cyan | blue |

| | | |
|---|---|---|
| Bag6 | red | |
| Mthfd1l | red | |
| Chd4 | red | green |
| C3 | red | |
| Csk | red | |
| Cul1 | red | |
| Cul3 | red | green |
| Cul5 | red | |
| Glyr1 | red | green |
| Prdx6 | red | |
| Hspd1 | cyan | |
| Ctnnd1 | red | |
| Dip2b | red | |
| Ddx6 | green | cyan, green, blue, red |
| Dld | red | |
| Dpp3 | red | |
| Drg2 | red | |
| Dnah5 | blue | green |
| Nae1 | red | |
| Xpo7 | red | |
| Usp24 | red | |
| Gcc2 | red | |
| Iars2 | cyan | |
| Dock7 | red | |
| Cul4b | red | blue |

| | | |
|---|---|---|
| Npepps | red | |
| Camk2d | red | |
| Itsn1 | red | |
| Sec22b | red | |
| Luc7l2 | red | blue |
| Farp1 | red | green |
| Atp2b4 | red | |
| Scamp3 | red | |
| Cad | red | |
| Mars | cyan | green |
| Rab3gap2 | blue | |
| Dhx9 | green | blue, cyan, green, yellow |
| Sacs | red | |
| Hadha | red | green |
| Echs1 | red | |
| Eef1g | cyan | |
| Eef2 | yellow | yellow, blue, reen |
| Etf1 | yellow | |
| Gspt1 | yellow | |
| Hnrnpd | green | blue |
| Ube2m | green | |
| Cfl1 | red | green |
| Ddx46 | green | green |
| Rps15a | yellow | yellow, green, blue |
| Fasn | red | |

| | | |
|---|---|---|
| Flnb | red | green |
| Fxr2 | red | green |
| Hk1 | red | |
| Gps1 | red | |
| Qrich1 | red | |
| Sec23ip | red | |
| Supt16 | red | green |
| Mink1 | red | |
| Dnm2 | red | green |
| Sf3b1 | green | |
| Hnrnpl | green | blue, green |
| Gars | cyan | |
| Gdi1 | red | |
| Gpc4 | red | |
| Cpsf6 | red | cyan, green |
| Cct8 | yellow | green |
| Hat1 | red | green |
| Hnrnpf | green | blue |
| Larp7 | red | blue |
| Eif2s2 | yellow | |
| Ilf2 | red | green, blue |
| Ipo11 | red | green |
| Ipo7 | cyan | |
| Ckb | red | |
| Kif5b | blue | green |

| | | |
|---|---|---|
| Mta1 | red | green |
| Mecp2 | red | green |
| Msn | red | green |
| Mt2 | red | |
| Mta2 | red | green |
| Mug1 | red | |
| Myh9 | red | green |
| Por | red | |
| Nsf | red | |
| rps14 | yellow | yellow, green, blue |
| Pes1 | cyan | green, blue |
| Pgm1 | red | |
| Phb | red | |
| Pls3 | red | green |
| Prep | red | |
| Psma4 | green | green, blue, cyan, red |
| Ptpn11 | red | |
| Adsl | cyan | |
| Atic | yellow | |
| Ssrp1 | red | green |
| Uchl1 | red | |
| Coro1c | red | green |
| Pgm3 | red | |
| Pelo | yellow | |
| Eef1a1 | yellow | green |

| | | |
|---|---|---|
| Clic1 | red | |
| Parp1 | red | green |
| Gdi2 | red | |
| Rtcb | cyan | |
| Vps35 | red | |
| Ola1 | cyan | green |
| Pafah1b1 | blue | green |
| Tnpo1 | green | |
| Map2k1 | red | green |
| G6pdx | red | |
| Gnb1 | red | green |
| Ap1b1 | red | |
| Psmd2 | green | |
| Dnajb1 | red | green |
| Hspa4 | green | green |
| Cct3 | cyan | green |
| Ugp2 | red | |
| Ppia | red | |
| Mcm6 | red | |
| Ddost | red | |
| Sptbn2 | blue | |
| Myh10 | red | green, blue |
| Ugdh | red | |
| Mcm3 | red | green |
| Stat3 | red | green |

| | | |
|---|---|---|
| Actn | red | |
| Tubb6 | blue | green |
| Gna11 | red | |
| Ruvbl2 | red | green, blue |
| Eif2b1 | red | |
| Rpl30 | yellow | yellow, green, blue |
| Wars | cyan | |
| Arpc2 | red | green |
| Rpl21 | yellow | yellow, green, blue |
| Rpl22 | yellow | yellow, green, blue |
| Tuba1c | blue | green |
| Kpna2 | red | cyan |
| Alb | red | |
| Ube2l3 | red | |
| Psmc4 | green | |
| Hnrnpm | green | blue |
| Rpl7a | yellow | yellow, green, blue |
| Rpl5 | yellow | yellow, green, blue |
| Matr3 | red | |
| Rps6 | yellow | cyan, yellow, green, blue |
| Rps16 | yellow | yellow, green, blue |
| Ube3a | red | |
| Hnrnpa1 | green | blue |
| Aldoa | red | green |
| Vim | red | green, blue |

| | | |
|---|---|---|
| Rpl9 | yellow | yellow, green, blue |
| Rpl28 | yellow | yellow, cyan, blue, green |
| Rps8 | yellow | yellow, green, blue |
| Abcf1 | cyan | yellow, blue, green |
| Actr2 | red | green |
| Ap2b1 | red | |
| Prpsap2 | cyan | yellow, green, blue |
| Rpl10a | yellow | yellow, green, blue |
| Copa | blue | |
| Eif3a | yellow | blue, green |
| Ap2m1 | red | |
| Eif3f | yellow | |
| Abce1 | yellow | |
| Snx1 | red | |
| Rcc1 | red | green |
| Rpl17 | yellow | yellow, green, blue |
| Aars | cyan | |
| Iqgap1 | red | green, blue, cyan |
| Hsp90ab1 | blue | |
| Rangap1 | blue | green |
| Prmt1 | red | |
| Nup155 | red | |
| Pdcd6ip | red | green |
| H2afy2 | red | green |
| Nap1l1 | red | |

| | | |
|---|---|---|
| Vamp4 | red | |
| Elavl1 | green | green, cyan, blue |
| Eif4a2 | yellow | green |
| Psmc2 | green | red, green, blue, cyan |
| Kars | cyan | |
| Klc1 | blue | green |
| Eif3b | yellow | |
| Qars | cyan | |
| Abcd3 | red | |
| Snx2 | blue | |
| Ddb1 | red | |
| Anxa6 | red | |
| Tubb4b | blue | green |
| Anxa2 | red | green |
| Mtch2 | red | |
| Arpc4 | red | green |
| Dynll1 | blue | green |
| Uqcrc1 | red | |
| Rcc2 | blue | green |
| Upf1 | yellow | cyan, red, blue, green |
| Rpl11 | yellow | yellow, green, blue |
| Rpl32 | yellow | yellow, green, blue |
| Rpl4 | yellow | yellow, green, blue |
| Rpn1 | cyan | |
| Rps11 | yellow | yellow, green, blue |

| | | |
|---|---|---|
| Rps17 | yellow | yellow, green, blue |
| Rps20 | yellow | yellow, green, blue |
| Rps27 | yellow | yellow, green, blue |
| Rps9 | yellow | yellow, green, blue |
| Xpnpep1 | red | |
| Ank2 | blue | green |
| Sae1 | cyan | |
| Sec31a | green | |
| Sdha | red | green |
| Sfpq | green | green |
| Snx27 | red | |
| Ubr4 | red | green |
| Gls | cyan | |
| Gcn1 | red | blue |
