## Supplementary Figures for "The low complexity motif of cytoplasmic polyadenylation element binding protein 3 (CPEB3) is critical for the trafficking of its targets in neurons"

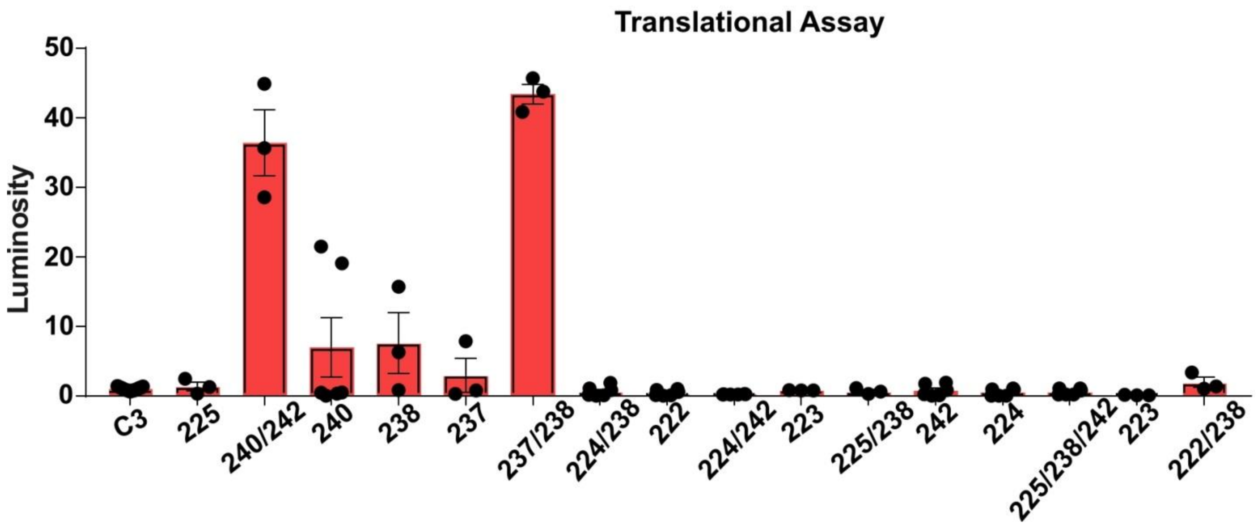


**Figure S1. CPEB3 low-complexity domain mutants disrupt inhibitory function.** The 16 mutants were created using mutagenesis of wild-type CPEB3-GFP, targeting the serines within the low-complexity domain. Our 16 mutants and full length CPEB3 were expressed in confluent HEK cells along with an actin-3'UTR-Renilla reporter plasmid. Translation of the reporter was measured using a luciferase assay, and luminosity was measured as a readout of translation. S240-242A, S240A, S238A, S237-238A are significantly different from CPEB3, $p<0.05$, as measured with One-Way ANOVA with Dunn's Multiple Comparison. n=3- 23. Error bars = +-SEM.

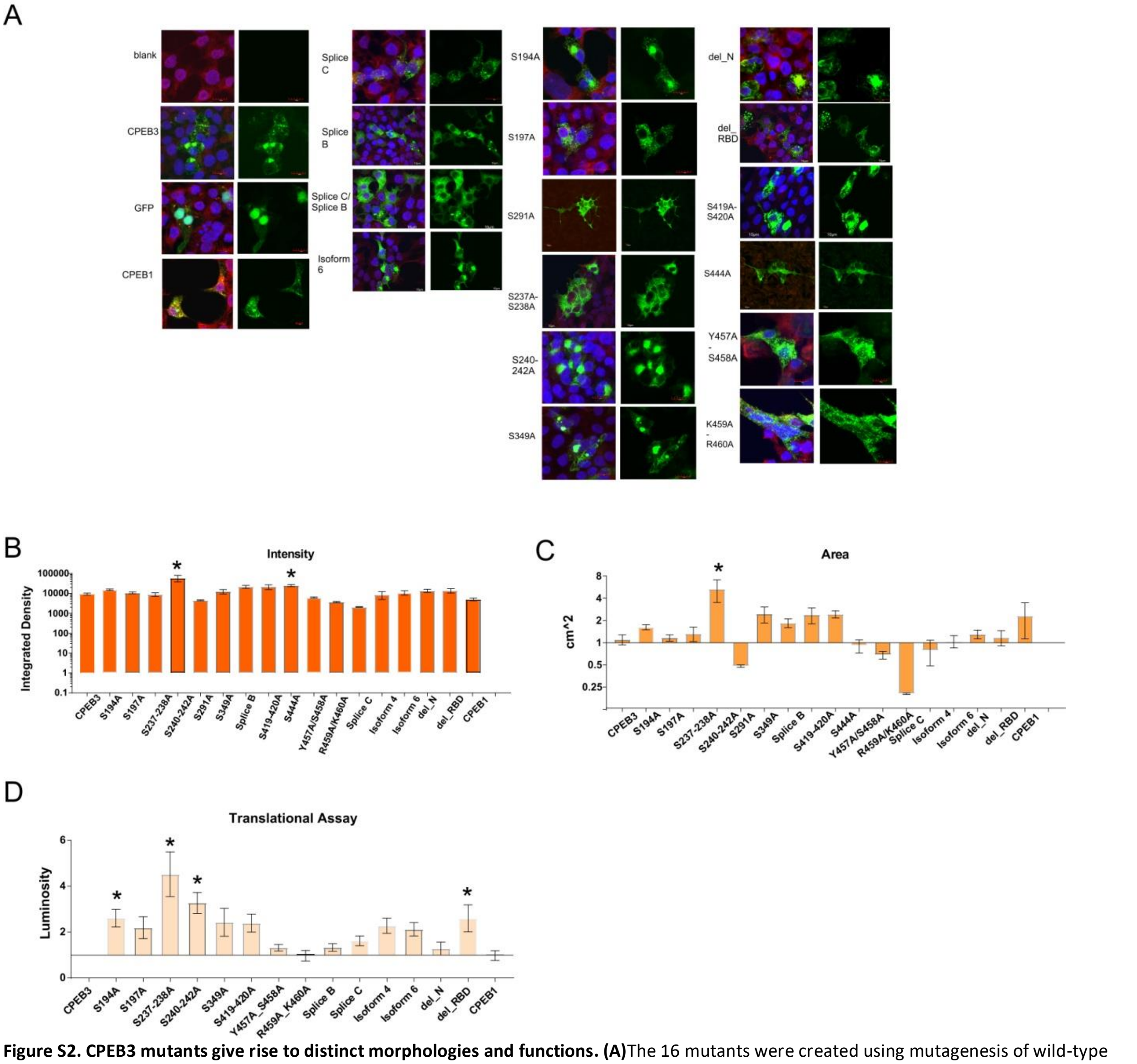


**Figure S2. CPEB3 mutants give rise to distinct morphologies and functions. (A)**The 16 mutants were created using mutagenesis of wild-type CPEB3-GFP. S194A, S197A, S291A, and S444A were created to disrupt the ability of CPEB3 to be phosphorylated by mutating serine to alanine (Creixell et al., 2012). delN and delRBD were created to delete entire domains that had previously been identified as critical for structure and function (Fioriti et al., 2015; Stephan et al., 2015; Ford et al., 2019), as well as to remove mass spectroscopy-identified modification sites. S237-238A, S240-242A, S349A, S419/420A, Y457A/S458A, and R459A/K460A was created to disrupt sites that are critically situated in areas of predicated structure, protein-binding, and modification, as well as to disrupt S/T/Y repeat sequences, which exhibit high phosphorylation probability in RNA-binding proteins (Monahan et. al, 2017). Finally, isoforms 2b, 3c, 4d, and 6f were created to avoid bias towards overexpressing isoform 1a. Expressed protein (green), G3BP control (red), and DAPI. **(B and C)** Our 16 mutants and full length CPEB3 were expressed in confluent HEK cells and visualized via their GFP tag using fluorescent microscopy. We hypothesized that disrupting regions important for function and/or structure would result in distinct morphologies *in vivo*. To test this we compared both the punctae area of the mutants to full length CPEB3, as well as signal intensity of punctae. We also measured these characteristics on our method control proteins: non-oligomerizing CPEB1 (Fioriti et al., 2015) and GFP. Critically, we included labeling of G3BP, a protein with high localization to stress granules (Kedersha and Anderson, 2007), to ensure that any CPEB3-GFP morphology wasn't an overexpression artifact. Co-localization of G3BP in red with CPEB3-GFP can be considered an artifact of overexpression, as CPEB3 does not naturally occur in stress granules (Kedersha and Anderson, 2007; Ford et al., 2019), and so all quantitation avoided overlapping fluorescence. The data were analyzed using punctae analysis software SyNPanal (Danielson et al., 2014). Two mutants had a significantly increased intensity of the punctae compared to control CPEB3, S240-242A and S444A. S240-242A was the only mutant to have significantly increased punctae area from control CPEB3. **(D)** Our 16 mutants and full length CPEB3 were expressed in confluent HEK cells along with an actin-3'UTR-Renilla reporter plasmid. Translation of the reporter was measured using a luciferase assay, and luminosity was measured as a readout of translation. S194, S237-238A, S240-242A, and del_RBD mutant were significantly different from CPEB3 levels. Scale bars represent 10 $\mu$m. *= $p<0.05$, as measured with One-Way ANOVA with Dunn's Multiple Comparison. n=10. Error bars = +-SEM.

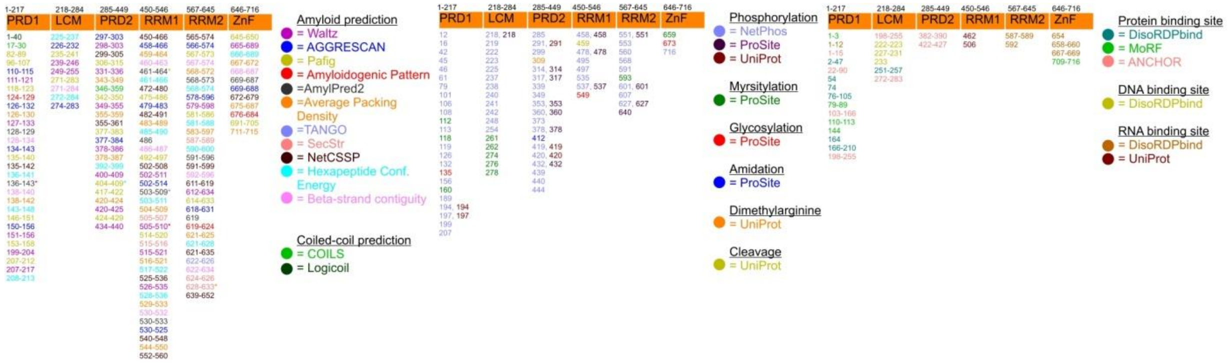

**Figure S3. Predictions of super-secondary structures, post-translational modifications, and binding domains of CPEB3.** Amyloid prediction algorithms AmylPred2 (Tsolis et al., 2013) (which includes Aggrescan (Conchillo-Sole et al., 2007), AmyloidMutants (O'Donnell et al., 2011), Amyloidogenic Pattern (Lopez de la Paz and Serrano, 2004), Average Packing Density (Galzitskaya and Lobanov, 2006), Beta-strand contiguity (Zibaee et al., 2007), Hexapeptide Conformational Energy (Zhang et al., 2007), NetCSSP (Kim et al., 2009), Pafig (Tian et al., 2009), SecStr (Hamodrakas et al., 2007), Tango (Fernandez-Escamilla et al., 2004), and Watlz (Maurer-Stroh et al., 2010)); and coiled-coil prediction algorithms COILS (Lupas et al., 1991) and LogiCoil (Vincent et al., 2013) were utilized to pinpoint regions of aggregation to expand our focus from areas previously studied (Fiumara et al., 2010; Raveendra et al., 2013; and Stephan et al., 2015). We then included post-translational modifications to our search, which have been identified as important for RNA-binding protein self-association (Nielsen et al., 2015; Calabretta and Richard, 2015; Araki et al., 2015; Monahan et al., 2017; Larson et al., 2017; Ambadipudi et al., 2017). From NetPhos (Blom et al., 1999), UniProt (The UniProt Consortium, 2017), and ProSite (Sigrist et al., 2009), we collected a variety of potential post-translational modification sites with an emphasis on phosphorylation, again expanding our scope from previous work (Theis et al., 2003; Fiumara et al., 2015; Drisaldi et al., 2015; and Kaczmarczyk et al., 2016). Finally, we utilized DisoRDPbind (Peng and Kurgan, 2015), MoRF (Disfani et al., 2012), UniProt (The UniProt Consortium, 2017), and ANCHOR (Meszaros et al., 2009; Dosztanyi et al., 2009) to discriminate areas that bind protein, RNA, and DNA (Figure 1), all of which are believed to be critical components of conformation change and/or RNA-binding function (Zanchetta et al., 2008; Nielsen et al., 2015; Calabretta and Richard, 2015; Stephan et al., 2015; Chen et al., 2016; Hult et al., 2017; Watanabe et al., 2017).

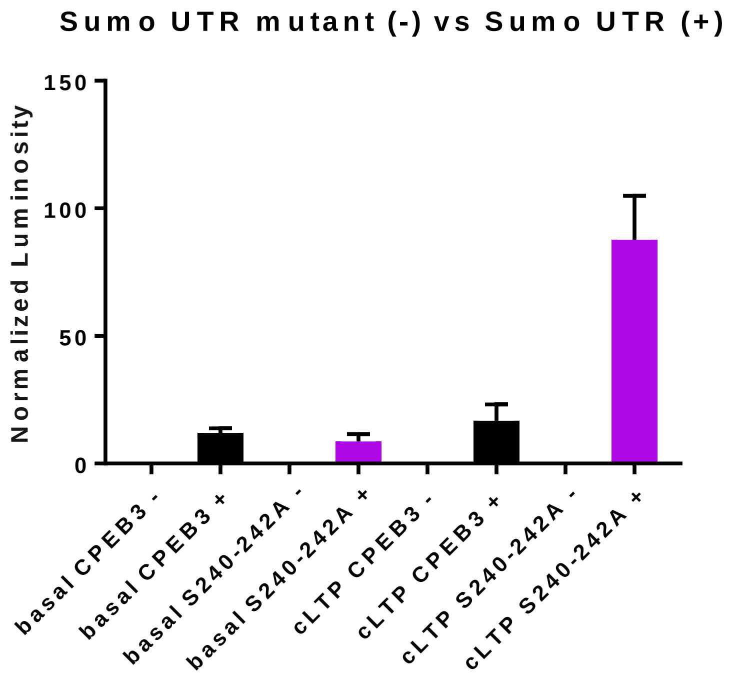


**Figure S4.** CPEB3 influences translation of SUMO2 via 3'UTR cytoplasmic polyadenylation element (CPE) binding. The full-length CPEB3 (black) or S240-242A mutant (purple) were expressed in primary neuronal cultures with a SUMO2 3'UTR-Renilla reporter (+) or a SUMO2 3'UTR mutant-Renilla reporter (-) that has a mutation in the CPE region of the 3'UTR. Both the CPEB3 and the S240-242A proteins can bind and influence translation of the reporter. When the CPE region is mutated, CPEB3 and S240-242A do not influence translation. Mutant reporter levels are negligible after background luminosity normalization. $p<0.001$ One-Way ANOVA with Dunn's Multiple Comparisons; all reporter + samples are significantly increased from the relevant mutant - samples.

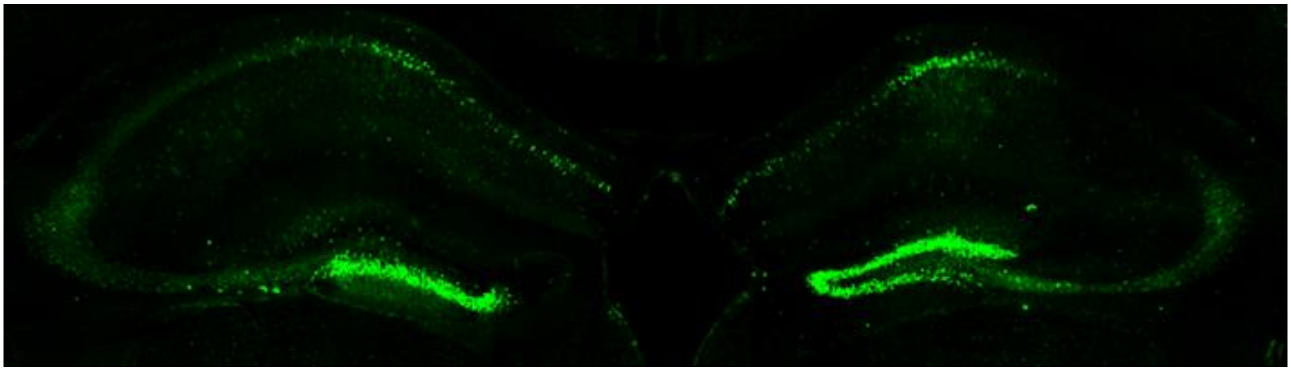
**Supp Figure 5**. CPEB3 *in vivo* viral expression.

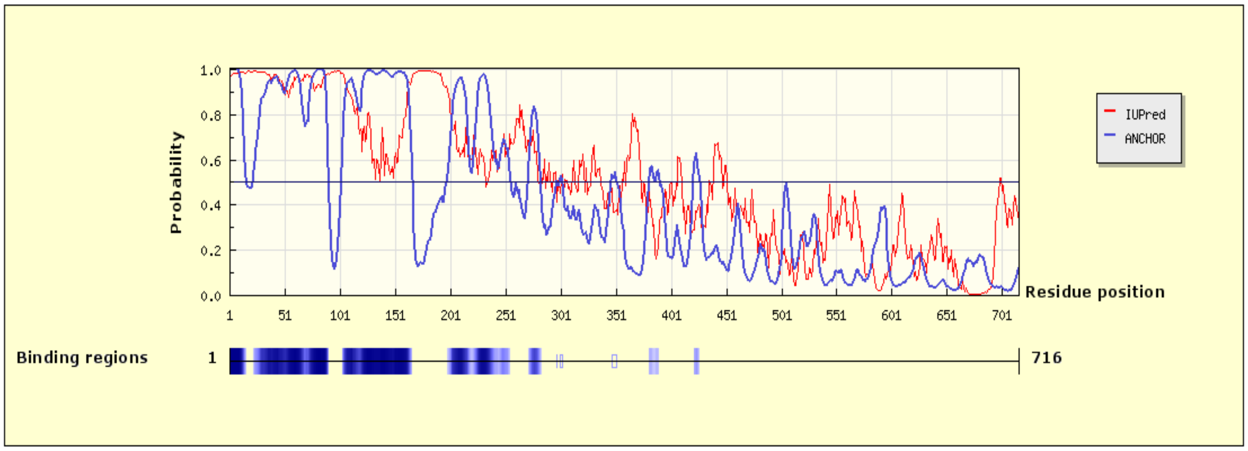

**Figure S6.** ANCHOR protein-protein interaction analysis of CPEB3
