## Supplementary Data 2 for "The low complexity motif of cytoplasmic polyadenylation element binding protein 3 (CPEB3) is critical for the trafficking of its targets in neurons"

Supplementary Data 2. Proteomics results, neurons

| | |
|---|---|
| ACTG_BOVIN | Actin, cytoplasmic 2 OS=Bos taurus OX=9913 GN=ACTG1 PE=1 SV=1 |
| TBB4B_MOUSE | Tubulin beta-4B chain OS=Mus musculus OX=10090 GN=Tubb4b PE=1 SV=1 |
| SPTN1_MOUSE | Spectrin alpha chain, non-erythrocytic 1 OS=Mus musculus OX=10090 GN=Sptan1 PE=1 SV=4 |
| AT1A3_MOUSE | Sodium/potassium-transporting ATPase subunit alpha-3 OS=Mus musculus OX=10090 GN=Atp1a3 PE=1 SV=1 |
| SPTB2_MOUSE | Spectrin beta chain, non-erythrocytic 1 OS=Mus musculus OX=10090 GN=Sptbn1 PE=1 SV=2 |
| HS90B_MOUSE | Heat shock protein HSP 90-beta OS=Mus musculus OX=10090 GN=Hsp90ab1 PE=1 SV=3 |
| MYH10_MOUSE | Myosin-10 OS=Mus musculus OX=10090 GN=Myh10 PE=1 SV=2 |
| ATPB_MOUSE | ATP synthase subunit beta, mitochondrial OS=Mus musculus OX=10090 GN=Atp5f1b PE=1 SV=2 |
| HSP7C_MOUSE | Heat shock cognate 71 kDa protein OS=Mus musculus OX=10090 GN=Hspa8 PE=1 SV=1 |
| G3P_MOUSE | Glyceraldehyde-3-phosphate dehydrogenase OS=Mus musculus OX=10090 GN=Gapdh PE=1 SV=2 |
| EF1A1_MOUSE | Elongation factor 1-alpha 1 OS=Mus musculus OX=10090 GN=Eef1a1 PE=1 SV=3 |
| UBA1_MOUSE | Ubiquitin-like modifier-activating enzyme 1 OS=Mus musculus OX=10090 GN=Uba1 PE=1 SV=1 |
| MAP1B_MOUSE | Microtubule-associated protein 1B OS=Mus musculus OX=10090 GN=Map1b PE=1 SV=2 |
| TERA_HUMAN | Transitional endoplasmic reticulum ATPase OS=Homo sapiens OX=9606 GN=VCP PE=1 SV=4 |
| EF2_MOUSE | Elongation factor 2 OS=Mus musculus OX=10090 GN=Eef2 PE=1 SV=2 |
| ATPA_MOUSE | ATP synthase subunit alpha, mitochondrial OS=Mus musculus OX=10090 GN=Atp5f1a PE=1 SV=1 |
| FAS_MOUSE | Fatty acid synthase OS=Mus musculus OX=10090 GN=Fasn PE=1 SV=2 |

| | |
|---|---|
| CH60_MOUSE | 60 kDa heat shock protein, mitochondrial OS=Mus musculus OX=10090 GN=Hspd1 PE=1 SV=1 |
| DPYL2_MOUSE | Dihydropyrimidinase-related protein 2 OS=Mus musculus OX=10090 GN=Dpysl2 PE=1 SV=2 |
| VIME_MOUSE | Vimentin OS=Mus musculus OX=10090 GN=Vim PE=1 SV=3 |
| MYH9_MOUSE | Myosin-9 OS=Mus musculus OX=10090 GN=Myh9 PE=1 SV=4 |
| KPYM_MOUSE | Pyruvate kinase PKM OS=Mus musculus OX=10090 GN=Pkm PE=1 SV=4 |
| MTAP2_MOUSE | Microtubule-associated protein 2 OS=Mus musculus OX=10090 GN=Map2 PE=1 SV=2 |
| KCRB_MOUSE | Creatine kinase B-type OS=Mus musculus OX=10090 GN=Ckb PE=1 SV=1 |
| ACON_MOUSE | Aconitate hydratase, mitochondrial OS=Mus musculus OX=10090 GN=Aco2 PE=1 SV=1 |
| PZP_MOUSE | Pregnancy zone protein OS=Mus musculus OX=10090 GN=Pzp PE=1 SV=3 |
| HS90A_MOUSE | Heat shock protein HSP 90-alpha OS=Mus musculus OX=10090 GN=Hsp90aa1 PE=1 SV=4 |
| DPYL1_MOUSE | Dihydropyrimidinase-related protein 1 OS=Mus musculus OX=10090 GN=Crmp1 PE=1 SV=1 |
| ENPL_MOUSE | Endoplasmin OS=Mus musculus OX=10090 GN=Hsp90b1 PE=1 SV=2 |
| GRP75_MOUSE | Stress-70 protein, mitochondrial OS=Mus musculus OX=10090 GN=Hspa9 PE=1 SV=3 |
| HSP74_MOUSE | Heat shock 70 kDa protein 4 OS=Mus musculus OX=10090 GN=Hspa4 PE=1 SV=1 |
| STXB1_MOUSE | Syntaxin-binding protein 1 OS=Mus musculus OX=10090 GN=Stxbp1 PE=1 SV=2 |
| NCAM1_MOUSE | Neural cell adhesion molecule 1 OS=Mus musculus OX=10090 GN=Ncam1 PE=1 SV=3 |
| FLNA_MOUSE | Filamin-A OS=Mus musculus OX=10090 GN=Flna PE=1 SV=5 |
| CO3_MOUSE | Complement C3 OS=Mus musculus OX=10090 GN=C3 PE=1 SV=3 |
| VATA_MOUSE | V-type proton ATPase catalytic subunit A OS=Mus musculus OX=10090 GN=Atp6v1a PE=1 SV=2 |

| | |
|---|---|
| GCAM_MOUSE | Ig gamma-2A chain C region, membrane-bound form OS=Mus musculus OX=10090 GN=Igh-1a PE=1 SV=3 |
| NSF_MOUSE | Vesicle-fusing ATPase OS=Mus musculus OX=10090 GN=Nsf PE=1 SV=2 |
| HXK1_MOUSE | Hexokinase-1 OS=Mus musculus OX=10090 GN=Hk1 PE=1 SV=3 |
| H2B1B_MOUSE | Histone H2B type 1-B OS=Mus musculus OX=10090 GN=Hist1h2bb PE=1 SV=3 |
| K2C1_HUMAN | Keratin, type II cytoskeletal 1 OS=Homo sapiens OX=9606 GN=KRT1 PE=1 SV=6 |
| DHX9_MOUSE | ATP-dependent RNA helicase A OS=Mus musculus OX=10090 GN=Dhx9 PE=1 SV=2 |
| ALDOA_MOUSE | Fructose-bisphosphate aldolase A OS=Mus musculus OX=10090 GN=Aldoa PE=1 SV=2 |
| AP2B1_MOUSE | AP-2 complex subunit beta OS=Mus musculus OX=10090 GN=Ap2b1 PE=1 SV=1 |
| MATR3_MOUSE | Matrin-3 OS=Mus musculus OX=10090 GN=Matr3 PE=1 SV=1 |
| HNRPU_MOUSE | Heterogeneous nuclear ribonucleoprotein U OS=Mus musculus OX=10090 GN=Hnrnpu PE=1 SV=1 |
| 1433E_MOUSE | 14-3-3 protein epsilon OS=Mus musculus OX=10090 GN=Ywhae PE=1 SV=1 |
| HNRPK_MOUSE | Heterogeneous nuclear ribonucleoprotein K OS=Mus musculus OX=10090 GN=Hnrnpk PE=1 SV=1 |
| BIP_MOUSE | Endoplasmic reticulum chaperone BiP OS=Mus musculus OX=10090 GN=Hspa5 PE=1 SV=3 |
| ENOA_MOUSE | Alpha-enolase OS=Mus musculus OX=10090 GN=Eno1 PE=1 SV=3 |
| DPYL5_MOUSE | Dihydropyrimidinase-related protein 5 OS=Mus musculus OX=10090 GN=Dpysl5 PE=1 SV=1 |
| CALX_MOUSE | Calnexin OS=Mus musculus OX=10090 GN=Canx PE=1 SV=1 |
| 2AAA_MOUSE | Serine/threonine-protein phosphatase 2A 65 kDa regulatory subunit A alpha isoform OS=Mus musculus OX=10090 GN=Ppp2r1a PE=1 SV=3 |
| SCOT1_MOUSE | Succinyl-CoA:3-ketoacid coenzyme A transferase 1, mitochondrial OS=Mus musculus OX=10090 GN=Oxct1 PE=1 SV=1 |

| | |
|---|---|
| AP2A1_MOUSE | AP-2 complex subunit alpha-1 OS=Mus musculus OX=10090 GN=Ap2a1 PE=1 SV=1 |
| 1433Z_MOUSE | 14-3-3 protein zeta/delta OS=Mus musculus OX=10090 GN=Ywhaz PE=1 SV=1 |
| LRP1_MOUSE | Prolow-density lipoprotein receptor-related protein 1 OS=Mus musculus OX=10090 GN=Lrp1 PE=1 SV=1 |
| GDIB_MOUSE | Rab GDP dissociation inhibitor beta OS=Mus musculus OX=10090 GN=Gdi2 PE=1 SV=1 |
| PGK1_MOUSE | Phosphoglycerate kinase 1 OS=Mus musculus OX=10090 GN=Pgk1 PE=1 SV=4 |
| ACLY_MOUSE | ATP-citrate synthase OS=Mus musculus OX=10090 GN=Acly PE=1 SV=1 |
| DPYL3_MOUSE | Dihydropyrimidinase-related protein 3 OS=Mus musculus OX=10090 GN=Dpysl3 PE=1 SV=1 |
| HNRPL_MOUSE | Heterogeneous nuclear ribonucleoprotein L OS=Mus musculus OX=10090 GN=Hnrnpl PE=1 SV=2 |
| PDIA3_MOUSE | Protein disulfide-isomerase A3 OS=Mus musculus OX=10090 GN=Pdia3 PE=1 SV=2 |
| ADT1_MOUSE | ADP/ATP translocase 1 OS=Mus musculus OX=10090 GN=Slc25a4 PE=1 SV=4 |
| LPPRC_MOUSE | Leucine-rich PPR motif-containing protein, mitochondrial OS=Mus musculus OX=10090 GN=Lrpprc PE=1 SV=2 |
| MIC60_MOUSE | MICOS complex subunit Mic60 OS=Mus musculus OX=10090 GN=Immt PE=1 SV=1 |
| TCPB_MOUSE | T-complex protein 1 subunit beta OS=Mus musculus OX=10090 GN=Cct2 PE=1 SV=4 |
| TCPQ_MOUSE | T-complex protein 1 subunit theta OS=Mus musculus OX=10090 GN=Cct8 PE=1 SV=3 |
| TKT_MOUSE | Transketolase OS=Mus musculus OX=10090 GN=Tkt PE=1 SV=1 |
| KCC2B_MOUSE | Calcium/calmodulin-dependent protein kinase type II subunit beta OS=Mus musculus OX=10090 GN=Camk2b PE=1 SV=2 |
| GNAO_MOUSE | Guanine nucleotide-binding protein G(o) subunit alpha OS=Mus musculus OX=10090 GN=Gnao1 PE=1 SV=3 |
| AL1L1_MOUSE | Cytosolic 10-formyltetrahydrofolate dehydrogenase OS=Mus musculus OX=10090 GN=Aldh1l1 PE=1 SV=1 |

| | |
|---|---|
| AT1A1_MOUSE | Sodium/potassium-transporting ATPase subunit alpha-1 OS=Mus musculus OX=10090 GN=Atp1a1 PE=1 SV=1 |
| AT2B1_MOUSE | Plasma membrane calcium-transporting ATPase 1 OS=Mus musculus OX=10090 GN=Atp2b1 PE=1 SV=1 |
| DHE3_MOUSE | Glutamate dehydrogenase 1, mitochondrial OS=Mus musculus OX=10090 GN=Glud1 PE=1 SV=1 |
| DNM1L_MOUSE | Dynamin-1-like protein OS=Mus musculus OX=10090 GN=Dnm1l PE=1 SV=2 |
| GABT_MOUSE | 4-aminobutyrate aminotransferase, mitochondrial OS=Mus musculus OX=10090 GN=Abat PE=1 SV=1 |
| LMNB1_MOUSE | Lamin-B1 OS=Mus musculus OX=10090 GN=Lmnb1 PE=1 SV=3 |
| VATB2_MOUSE | V-type proton ATPase subunit B, brain isoform OS=Mus musculus OX=10090 GN=Atp6v1b2 PE=1 SV=1 |
| AATM_MOUSE | Aspartate aminotransferase, mitochondrial OS=Mus musculus OX=10090 GN=Got2 PE=1 SV=1 |
| CON__P13645 | CON__P13645 |
| MAP1A_MOUSE | Microtubule-associated protein 1A OS=Mus musculus OX=10090 GN=Map1a PE=1 SV=2 |
| MDHM_MOUSE | Malate dehydrogenase, mitochondrial OS=Mus musculus OX=10090 GN=Mdh2 PE=1 SV=3 |
| FSCN1_MOUSE | Fascin OS=Mus musculus OX=10090 GN=Fscn1 PE=1 SV=4 |
| MARCS_MOUSE | Myristoylated alanine-rich C-kinase substrate OS=Mus musculus OX=10090 GN=Marcks PE=1 SV=2 |
| PPIA_MOUSE | Peptidyl-prolyl cis-trans isomerase A OS=Mus musculus OX=10090 GN=Ppia PE=1 SV=2 |
| ROA2_MOUSE | Heterogeneous nuclear ribonucleoproteins A2/B1 OS=Mus musculus OX=10090 GN=Hnrnpa2b1 PE=1 SV=2 |
| SFPQ_MOUSE | Splicing factor, proline- and glutamine-rich OS=Mus musculus OX=10090 GN=Sfpq PE=1 SV=1 |
| VPP1_MOUSE | V-type proton ATPase 116 kDa subunit a isoform 1 OS=Mus musculus OX=10090 GN=Atp6v0a1 PE=1 SV=3 |

| | |
|---|---|
| HNRPM_MOUSE | Heterogeneous nuclear ribonucleoprotein M OS=Mus musculus OX=10090 GN=Hnrnpm PE=1 SV=3 |
| SYN1_MOUSE | Synapsin-1 OS=Mus musculus OX=10090 GN=Syn1 PE=1 SV=2 |
| TBB2A_MOUSE | Tubulin beta-2A chain OS=Mus musculus OX=10090 GN=Tubb2a PE=1 SV=1 |
| UBP5_MOUSE | Ubiquitin carboxyl-terminal hydrolase 5 OS=Mus musculus OX=10090 GN=Usp5 PE=1 SV=1 |
| VPS35_MOUSE | Vacuolar protein sorting-associated protein 35 OS=Mus musculus OX=10090 GN=Vps35 PE=1 SV=1 |
| GDIA_MOUSE | Rab GDP dissociation inhibitor alpha OS=Mus musculus OX=10090 GN=Gdi1 PE=1 SV=3 |
| TBB3_MOUSE | Tubulin beta-3 chain OS=Mus musculus OX=10090 GN=Tubb3 PE=1 SV=1 |
| VDAC1_MOUSE | Voltage-dependent anion-selective channel protein 1 OS=Mus musculus OX=10090 GN=Vdac1 PE=1 SV=3 |
| MYO5A_MOUSE | Unconventional myosin-Va OS=Mus musculus OX=10090 GN=Myo5a PE=1 SV=2 |
| ODO1_MOUSE | 2-oxoglutarate dehydrogenase, mitochondrial OS=Mus musculus OX=10090 GN=Ogdh PE=1 SV=3 |
| HNRH1_MOUSE | Heterogeneous nuclear ribonucleoprotein H OS=Mus musculus OX=10090 GN=Hnrnph1 PE=1 SV=3 |
| IF4A1_MOUSE | Eukaryotic initiation factor 4A-I OS=Mus musculus OX=10090 GN=Eif4a1 PE=1 SV=1 |
| TCPA_MOUSE | T-complex protein 1 subunit alpha OS=Mus musculus OX=10090 GN=Tcp1 PE=1 SV=3 |
| ACTN4_MOUSE | Alpha-actinin-4 OS=Mus musculus OX=10090 GN=Actn4 PE=1 SV=1 |
| CMC1_MOUSE | Calcium-binding mitochondrial carrier protein Aralar1 OS=Mus musculus OX=10090 GN=Slc25a12 PE=1 SV=1 |
| ROA1_BOVIN | Heterogeneous nuclear ribonucleoprotein A1 OS=Bos taurus OX=9913 GN=HNRNPA1 PE=1 SV=2 |
| CALR_MOUSE | Calreticulin OS=Mus musculus OX=10090 GN=Calr PE=1 SV=1 |
| COF1_MOUSE | Cofilin-1 OS=Mus musculus OX=10090 GN=Cfl1 PE=1 SV=3 |

| | |
|---|---|
| H4_HUMAN | Histone H4 OS=Homo sapiens OX=9606 GN=H4C1 PE=1 SV=2 |
| LDHB_MOUSE | L-lactate dehydrogenase B chain OS=Mus musculus OX=10090 GN=Ldhb PE=1 SV=2 |
| SYAC_MOUSE | Alanine--tRNA ligase, cytoplasmic OS=Mus musculus OX=10090 GN=Aars PE=1 SV=1 |
| MACF1_MOUSE | Microtubule-actin cross-linking factor 1 OS=Mus musculus OX=10090 GN=Macf1 PE=1 SV=2 |
| MPCP_MOUSE | Phosphate carrier protein, mitochondrial OS=Mus musculus OX=10090 GN=Slc25a3 PE=1 SV=1 |
| NUCL_MOUSE | Nucleolin OS=Mus musculus OX=10090 GN=Ncl PE=1 SV=2 |
| QCR2_MOUSE | Cytochrome b-c1 complex subunit 2, mitochondrial OS=Mus musculus OX=10090 GN=Uqcrc2 PE=1 SV=1 |
| RTN4_MOUSE | Reticulon-4 OS=Mus musculus OX=10090 GN=Rtn4 PE=1 SV=2 |
| TOP2B_MOUSE | DNA topoisomerase 2-beta OS=Mus musculus OX=10090 GN=Top2b PE=1 SV=2 |
| DREB_MOUSE | Drebrin OS=Mus musculus OX=10090 GN=Dbn1 PE=1 SV=4 |
| ANXA6_MOUSE | Annexin A6 OS=Mus musculus OX=10090 GN=Anxa6 PE=1 SV=3 |
| IMB1_MOUSE | Importin subunit beta-1 OS=Mus musculus OX=10090 GN=Kpnb1 PE=1 SV=2 |
| RPN2_MOUSE | Dolichyl-diphosphooligosaccharide--protein glycosyltransferase subunit 2 OS=Mus musculus OX=10090 GN=Rpn2 PE=1 SV=1 |
| CISY_MOUSE | Citrate synthase, mitochondrial OS=Mus musculus OX=10090 GN=Cs PE=1 SV=1 |
| G6PI_MOUSE | Glucose-6-phosphate isomerase OS=Mus musculus OX=10090 GN=Gpi PE=1 SV=4 |
| GUAD_MOUSE | Guanine deaminase OS=Mus musculus OX=10090 GN=Gda PE=1 SV=1 |
| MAP6_MOUSE | Microtubule-associated protein 6 OS=Mus musculus OX=10090 GN=Map6 PE=1 SV=2 |
| QCR1_MOUSE | Cytochrome b-c1 complex subunit 1, mitochondrial OS=Mus musculus OX=10090 GN=Uqcrc1 PE=1 SV=2 |
| TCPD_MOUSE | T-complex protein 1 subunit delta OS=Mus musculus OX=10090 GN=Cct4 PE=1 SV=3 |

| | |
|---|---|
| CAND1_MOUSE | Cullin-associated NEDD8-dissociated protein 1 OS=Mus musculus OX=10090 GN=Cand1 PE=1 SV=2 |
| DDX3X_MOUSE | ATP-dependent RNA helicase DDX3X OS=Mus musculus OX=10090 GN=Ddx3x PE=1 SV=3 |
| GBB1_MOUSE | Guanine nucleotide-binding protein G(I)/G(S)/G(T) subunit beta-1 OS=Mus musculus OX=10090 GN=Gnb1 PE=1 SV=3 |
| NDUS1_MOUSE | NADH-ubiquinone oxidoreductase 75 kDa subunit, mitochondrial OS=Mus musculus OX=10090 GN=Ndufs1 PE=1 SV=2 |
| PYC_MOUSE | Pyruvate carboxylase, mitochondrial OS=Mus musculus OX=10090 GN=Pc PE=1 SV=1 |
| SYT1_MOUSE | Synaptotagmin-1 OS=Mus musculus OX=10090 GN=Syt1 PE=1 SV=1 |
| TCPG_MOUSE | T-complex protein 1 subunit gamma OS=Mus musculus OX=10090 GN=Cct3 PE=1 SV=1 |
| ALBU_MOUSE | Serum albumin OS=Mus musculus OX=10090 GN=Alb PE=1 SV=3 |
| CALM1_MOUSE | Calmodulin-1 OS=Mus musculus OX=10090 GN=Calm1 PE=1 SV=1 |
| KAP3_MOUSE | cAMP-dependent protein kinase type II-beta regulatory subunit OS=Mus musculus OX=10090 GN=Prkar2b PE=1 SV=3 |
| CAP1_MOUSE | Adenylyl cyclase-associated protein 1 OS=Mus musculus OX=10090 GN=Cap1 PE=1 SV=4 |
| SDHA_MOUSE | Succinate dehydrogenase [ubiquinone] flavoprotein subunit, mitochondrial OS=Mus musculus OX=10090 GN=Sdha PE=1 SV=1 |
| DYN1_MOUSE | Dynamin-1 OS=Mus musculus OX=10090 GN=Dnm1 PE=1 SV=2 |
| PRP8_MOUSE | Pre-mRNA-processing-splicing factor 8 OS=Mus musculus OX=10090 GN=Prpf8 PE=1 SV=2 |
| STIP1_MOUSE | Stress-induced-phosphoprotein 1 OS=Mus musculus OX=10090 GN=Stip1 PE=1 SV=1 |
| RPN1_MOUSE | Dolichyl-diphosphooligosaccharide--protein glycosyltransferase subunit 1 OS=Mus musculus OX=10090 GN=Rpn1 PE=1 SV=1 |
| PGAM1_MOUSE | Phosphoglycerate mutase 1 OS=Mus musculus OX=10090 GN=Pgam1 PE=1 SV=3 |
| PSA_MOUSE | Puromycin-sensitive aminopeptidase OS=Mus musculus OX=10090 GN=Npepps PE=1 SV=2 |

| | |
|---|---|
| PSMD2_MOUSE | 26S proteasome non-ATPase regulatory subunit 2 OS=Mus musculus OX=10090 GN=Psmd2 PE=1 SV=1 |
| TCPE_MOUSE | T-complex protein 1 subunit epsilon OS=Mus musculus OX=10090 GN=Cct5 PE=1 SV=1 |
| WDR1_MOUSE | WD repeat-containing protein 1 OS=Mus musculus OX=10090 GN=Wdr1 PE=1 SV=3 |
| CON__P35908 | CON__P35908 |
| TBB5_MOUSE | Tubulin beta-5 chain OS=Mus musculus OX=10090 GN=Tubb5 PE=1 SV=1 |
| CAPS1_MOUSE | Calcium-dependent secretion activator 1 OS=Mus musculus OX=10090 GN=Cadps PE=1 SV=3 |
| GANAB_MOUSE | Neutral alpha-glucosidase AB OS=Mus musculus OX=10090 GN=Ganab PE=1 SV=1 |
| IDHC_MOUSE | Isocitrate dehydrogenase [NADP] cytoplasmic OS=Mus musculus OX=10090 GN=Idh1 PE=1 SV=2 |
| KIF5C_MOUSE | Kinesin heavy chain isoform 5C OS=Mus musculus OX=10090 GN=Kif5c PE=1 SV=3 |
| THIL_MOUSE | Acetyl-CoA acetyltransferase, mitochondrial OS=Mus musculus OX=10090 GN=Acat1 PE=1 SV=1 |
| SYEP_MOUSE | Bifunctional glutamate/proline--tRNA ligase OS=Mus musculus OX=10090 GN=Eprs PE=1 SV=4 |
| IGG2B_MOUSE | Ig gamma-2B chain C region OS=Mus musculus OX=10090 GN=Igh-3 PE=1 SV=3 |
| ARP3_MOUSE | Actin-related protein 3 OS=Mus musculus OX=10090 GN=Actr3 PE=1 SV=3 |
| HYOU1_MOUSE | Hypoxia up-regulated protein 1 OS=Mus musculus OX=10090 GN=Hyou1 PE=1 SV=1 |
| XPO2_MOUSE | Exportin-2 OS=Mus musculus OX=10090 GN=Cse1l PE=1 SV=1 |
| CYFP2_MOUSE | Cytoplasmic FMR1-interacting protein 2 OS=Mus musculus OX=10090 GN=Cyfip2 PE=1 SV=2 |
| EF1G_MOUSE | Elongation factor 1-gamma OS=Mus musculus OX=10090 GN=Eef1g PE=1 SV=3 |
| FLNB_MOUSE | Filamin-B OS=Mus musculus OX=10090 GN=Flnb PE=1 SV=3 |
| H2A1C_HUMAN | Histone H2A type 1-C OS=Homo sapiens OX=9606 GN=HIST1H2AC PE=1 SV=3 |

| | |
|---|---|
| CATA_MOUSE | Catalase OS=Mus musculus OX=10090 GN=Cat PE=1 SV=4 |
| GPDM_MOUSE | Glycerol-3-phosphate dehydrogenase, mitochondrial OS=Mus musculus OX=10090 GN=Gpd2 PE=1 SV=2 |
| NCDN_MOUSE | Neurochondrin OS=Mus musculus OX=10090 GN=Ncdn PE=1 SV=1 |
| ROA3_MOUSE | Heterogeneous nuclear ribonucleoprotein A3 OS=Mus musculus OX=10090 GN=Hnrnpa3 PE=1 SV=1 |
| USP9X_MOUSE | Probable ubiquitin carboxyl-terminal hydrolase FAF-X OS=Mus musculus OX=10090 GN=Usp9x PE=1 SV=2 |
| DDX5_MOUSE | Probable ATP-dependent RNA helicase DDX5 OS=Mus musculus OX=10090 GN=Ddx5 PE=1 SV=2 |
| PDIA1_MOUSE | Protein disulfide-isomerase OS=Mus musculus OX=10090 GN=P4hb PE=1 SV=2 |
| PHB2_MOUSE | Prohibitin-2 OS=Mus musculus OX=10090 GN=Phb2 PE=1 SV=1 |
| SYYC_MOUSE | Tyrosine--tRNA ligase, cytoplasmic OS=Mus musculus OX=10090 GN=Yars PE=1 SV=3 |
| ECHA_MOUSE | Trifunctional enzyme subunit alpha, mitochondrial OS=Mus musculus OX=10090 GN=Hadha PE=1 SV=1 |
| MOES_MOUSE | Moesin OS=Mus musculus OX=10090 GN=Msn PE=1 SV=3 |
| RACK1_MOUSE | Receptor of activated protein C kinase 1 OS=Mus musculus OX=10090 GN=Rack1 PE=1 SV=3 |
| 1433G_MOUSE | 14-3-3 protein gamma OS=Mus musculus OX=10090 GN=Ywhag PE=1 SV=2 |
| AP2A2_MOUSE | AP-2 complex subunit alpha-2 OS=Mus musculus OX=10090 GN=Ap2a2 PE=1 SV=2 |
| DX39B_MOUSE | Spliceosome RNA helicase Ddx39b OS=Mus musculus OX=10090 GN=Ddx39b PE=1 SV=1 |
| PSMD1_MOUSE | 26S proteasome non-ATPase regulatory subunit 1 OS=Mus musculus OX=10090 GN=Psmd1 PE=1 SV=1 |
| TCPH_MOUSE | T-complex protein 1 subunit eta OS=Mus musculus OX=10090 GN=Cct7 PE=1 SV=1 |
| AT2A2_MOUSE | Sarcoplasmic/endoplasmic reticulum calcium ATPase 2 OS=Mus musculus OX=10090 GN=Atp2a2 PE=1 SV=2 |

| | |
|---|---|
| PYGB_MOUSE | Glycogen phosphorylase, brain form OS=Mus musculus OX=10090 GN=Pygb PE=1 SV=3 |
| TCPZ_MOUSE | T-complex protein 1 subunit zeta OS=Mus musculus OX=10090 GN=Cct6a PE=1 SV=3 |
| IDHP_MOUSE | Isocitrate dehydrogenase [NADP], mitochondrial OS=Mus musculus OX=10090 GN=Idh2 PE=1 SV=3 |
| PP2AA_MOUSE | Serine/threonine-protein phosphatase 2A catalytic subunit alpha isoform OS=Mus musculus OX=10090 GN=Ppp2ca PE=1 SV=1 |
| ODPB_MOUSE | Pyruvate dehydrogenase E1 component subunit beta, mitochondrial OS=Mus musculus OX=10090 GN=Pdhb PE=1 SV=1 |
| CNTN1_MOUSE | Contactin-1 OS=Mus musculus OX=10090 GN=Cntn1 PE=1 SV=1 |
| PABP1_MOUSE | Polyadenylate-binding protein 1 OS=Mus musculus OX=10090 GN=Pabpc1 PE=1 SV=2 |
| RS4X_MOUSE | 40S ribosomal protein S4, X isoform OS=Mus musculus OX=10090 GN=Rps4x PE=1 SV=2 |
| ANXA2_MOUSE | Annexin A2 OS=Mus musculus OX=10090 GN=Anxa2 PE=1 SV=2 |
| CKAP4_MOUSE | Cytoskeleton-associated protein 4 OS=Mus musculus OX=10090 GN=Ckap4 PE=1 SV=2 |
| ANK2_MOUSE | Ankyrin-2 OS=Mus musculus OX=10090 GN=Ank2 PE=1 SV=2 |
| AT1B1_MOUSE | Sodium/potassium-transporting ATPase subunit beta-1 OS=Mus musculus OX=10090 GN=Atp1b1 PE=1 SV=1 |
| PFKAM_MOUSE | ATP-dependent 6-phosphofructokinase, muscle type OS=Mus musculus OX=10090 GN=Pfkm PE=1 SV=3 |
| HS105_MOUSE | Heat shock protein 105 kDa OS=Mus musculus OX=10090 GN=Hsph1 PE=1 SV=2 |
| LDHA_MOUSE | L-lactate dehydrogenase A chain OS=Mus musculus OX=10090 GN=Ldha PE=1 SV=3 |
| 4F2_MOUSE | 4F2 cell-surface antigen heavy chain OS=Mus musculus OX=10090 GN=Slc3a2 PE=1 SV=1 |
| RS3_MOUSE | 40S ribosomal protein S3 OS=Mus musculus OX=10090 GN=Rps3 PE=1 SV=1 |
| ANXA5_MOUSE | Annexin A5 OS=Mus musculus OX=10090 GN=Anxa5 PE=1 SV=1 |

| | |
|---|---|
| AP2M1_MOUSE | AP-2 complex subunit mu OS=Mus musculus OX=10090 GN=Ap2m1 PE=1 SV=1 |
| C1QA_MOUSE | Complement C1q subcomponent subunit A OS=Mus musculus OX=10090 GN=C1qa PE=1 SV=2 |
| DCTN1_MOUSE | Dynactin subunit 1 OS=Mus musculus OX=10090 GN=Dctn1 PE=1 SV=3 |
| FKBP4_MOUSE | Peptidyl-prolyl cis-trans isomerase FKBP4 OS=Mus musculus OX=10090 GN=Fkbp4 PE=1 SV=5 |
| GARS_MOUSE | Glycine--tRNA ligase OS=Mus musculus OX=10090 GN=Gars PE=1 SV=1 |
| RL4_MOUSE | 60S ribosomal protein L4 OS=Mus musculus OX=10090 GN=Rpl4 PE=1 SV=3 |
| ODPA_MOUSE | Pyruvate dehydrogenase E1 component subunit alpha, somatic form, mitochondrial OS=Mus musculus OX=10090 GN=Pdha1 PE=1 SV=1 |
| SUCB1_MOUSE | Succinate--CoA ligase [ADP-forming] subunit beta, mitochondrial OS=Mus musculus OX=10090 GN=Sucla2 PE=1 SV=2 |
| U520_MOUSE | U5 small nuclear ribonucleoprotein 200 kDa helicase OS=Mus musculus OX=10090 GN=Snrnp200 PE=1 SV=1 |
| ACTA_MOUSE | Actin, aortic smooth muscle OS=Mus musculus OX=10090 GN=Acta2 PE=1 SV=1 |
| SERPH_MOUSE | Serpin H1 OS=Mus musculus OX=10090 GN=Serpinh1 PE=1 SV=3 |
| APOE_MOUSE | Apolipoprotein E OS=Mus musculus OX=10090 GN=Apoe PE=1 SV=2 |
| C1QB_MOUSE | Complement C1q subcomponent subunit B OS=Mus musculus OX=10090 GN=C1qb PE=1 SV=2 |
| PCBP2_MOUSE | Poly(rC)-binding protein 2 OS=Mus musculus OX=10090 GN=Pcbp2 PE=1 SV=1 |
| PRDX6_MOUSE | Peroxiredoxin-6 OS=Mus musculus OX=10090 GN=Prdx6 PE=1 SV=3 |
| UCHL1_MOUSE | Ubiquitin carboxyl-terminal hydrolase isozyme L1 OS=Mus musculus OX=10090 GN=Uchl1 PE=1 SV=1 |
| XPO1_MOUSE | Exportin-1 OS=Mus musculus OX=10090 GN=Xpo1 PE=1 SV=1 |
| BASP1_MOUSE | Brain acid soluble protein 1 OS=Mus musculus OX=10090 GN=Basp1 PE=1 SV=3 |

| | |
|---|---|
| DHB4_MOUSE | Peroxisomal multifunctional enzyme type 2 OS=Mus musculus OX=10090 GN=Hsd17b4 PE=1 SV=3 |
| PLCB1_MOUSE | 1-phosphatidylinositol 4,5-bisphosphate phosphodiesterase beta-1 OS=Mus musculus OX=10090 GN=Plcb1 PE=1 SV=2 |
| AP180_MOUSE | Clathrin coat assembly protein AP180 OS=Mus musculus OX=10090 GN=Snap91 PE=1 SV=1 |
| EIF3A_MOUSE | Eukaryotic translation initiation factor 3 subunit A OS=Mus musculus OX=10090 GN=Eif3a PE=1 SV=5 |
| MDHC_MOUSE | Malate dehydrogenase, cytoplasmic OS=Mus musculus OX=10090 GN=Mdh1 PE=1 SV=3 |
| FABP7_MOUSE | Fatty acid-binding protein, brain OS=Mus musculus OX=10090 GN=Fabp7 PE=1 SV=2 |
| GNAI2_MOUSE | Guanine nucleotide-binding protein G(i) subunit alpha-2 OS=Mus musculus OX=10090 GN=Gnai2 PE=1 SV=5 |
| GSTM1_MOUSE | Glutathione S-transferase Mu 1 OS=Mus musculus OX=10090 GN=Gstm1 PE=1 SV=2 |
| PHB_MOUSE | Prohibitin OS=Mus musculus OX=10090 GN=Phb PE=1 SV=1 |
| AKA12_MOUSE | A-kinase anchor protein 12 OS=Mus musculus OX=10090 GN=Akap12 PE=1 SV=1 |
| DCLK1_MOUSE | Serine/threonine-protein kinase DCLK1 OS=Mus musculus OX=10090 GN=Dclk1 PE=1 SV=1 |
| DDX1_MOUSE | ATP-dependent RNA helicase DDX1 OS=Mus musculus OX=10090 GN=Ddx1 PE=1 SV=1 |
| EFTU_MOUSE | Elongation factor Tu, mitochondrial OS=Mus musculus OX=10090 GN=Tufm PE=1 SV=1 |
| PGBM_MOUSE | Basement membrane-specific heparan sulfate proteoglycan core protein OS=Mus musculus OX=10090 GN=Hspg2 PE=1 SV=1 |
| TAU_MOUSE | Microtubule-associated protein tau OS=Mus musculus OX=10090 GN=Mapt PE=1 SV=3 |
| DLDH_MOUSE | Dihydrolipoyl dehydrogenase, mitochondrial OS=Mus musculus OX=10090 GN=Dld PE=1 SV=2 |
| HNRL2_MOUSE | Heterogeneous nuclear ribonucleoprotein U-like protein 2 OS=Mus musculus OX=10090 GN=Hnrnpul2 PE=1 SV=2 |

| | |
|---|---|
| NONO_MOUSE | Non-POU domain-containing octamer-binding protein OS=Mus musculus OX=10090 GN=Nono PE=1 SV=3 |
| ODP2_MOUSE | Dihydrolipoyllysine-residue acetyltransferase component of pyruvate dehydrogenase complex, mitochondrial OS=Mus musculus OX=10090 GN=Dlat PE=1 SV=2 |
| EIF3B_MOUSE | Eukaryotic translation initiation factor 3 subunit B OS=Mus musculus OX=10090 GN=Eif3b PE=1 SV=1 |
| H2AY_MOUSE | Core histone macro-H2A.1 OS=Mus musculus OX=10090 GN=H2afy PE=1 SV=3 |
| IF4G1_MOUSE | Eukaryotic translation initiation factor 4 gamma 1 OS=Mus musculus OX=10090 GN=Eif4g1 PE=1 SV=1 |
| TPIS_MOUSE | Triosephosphate isomerase OS=Mus musculus OX=10090 GN=Tpi1 PE=1 SV=4 |
| C1TC_MOUSE | C-1-tetrahydrofolate synthase, cytoplasmic OS=Mus musculus OX=10090 GN=Mthfd1 PE=1 SV=4 |
| HNRPQ_MOUSE | Heterogeneous nuclear ribonucleoprotein Q OS=Mus musculus OX=10090 GN=Syncrip PE=1 SV=2 |
| LIS1_MOUSE | Platelet-activating factor acetylhydrolase IB subunit alpha OS=Mus musculus OX=10090 GN=Pafah1b1 PE=1 SV=2 |
| NEST_MOUSE | Nestin OS=Mus musculus OX=10090 GN=Nes PE=1 SV=1 |
| RAB14_MOUSE | Ras-related protein Rab-14 OS=Mus musculus OX=10090 GN=Rab14 PE=1 SV=3 |
| SAHH_MOUSE | Adenosylhomocysteinase OS=Mus musculus OX=10090 GN=Ahcy PE=1 SV=3 |
| ATPG_MOUSE | ATP synthase subunit gamma, mitochondrial OS=Mus musculus OX=10090 GN=Atp5f1c PE=1 SV=1 |
| ATX10_MOUSE | Ataxin-10 OS=Mus musculus OX=10090 GN=Atxn10 PE=1 SV=2 |
| DEST_MOUSE | Destrin OS=Mus musculus OX=10090 GN=Dstn PE=1 SV=3 |
| IGHG3_MOUSE | Ig gamma-3 chain C region OS=Mus musculus OX=10090 PE=1 SV=2 |
| MK01_MOUSE | Mitogen-activated protein kinase 1 OS=Mus musculus OX=10090 GN=Mapk1 PE=1 SV=3 |

| | |
|---|---|
| TFR1_MOUSE | Transferrin receptor protein 1 OS=Mus musculus OX=10090 GN=Tfrc PE=1 SV=1 |
| GHC1_MOUSE | Mitochondrial glutamate carrier 1 OS=Mus musculus OX=10090 GN=Slc25a22 PE=1 SV=1 |
| RAB2A_MOUSE | Ras-related protein Rab-2A OS=Mus musculus OX=10090 GN=Rab2a PE=1 SV=1 |
| RL3_MOUSE | 60S ribosomal protein L3 OS=Mus musculus OX=10090 GN=Rpl3 PE=1 SV=3 |
| RL6_MOUSE | 60S ribosomal protein L6 OS=Mus musculus OX=10090 GN=Rpl6 PE=1 SV=3 |
| DCTN2_MOUSE | Dynactin subunit 2 OS=Mus musculus OX=10090 GN=Dctn2 PE=1 SV=3 |
| M2OM_MOUSE | Mitochondrial 2-oxoglutarate/malate carrier protein OS=Mus musculus OX=10090 GN=Slc25a11 PE=1 SV=3 |
| PEBP1_MOUSE | Phosphatidylethanolamine-binding protein 1 OS=Mus musculus OX=10090 GN=Pebp1 PE=1 SV=3 |
| PRS7_MOUSE | 26S proteasome regulatory subunit 7 OS=Mus musculus OX=10090 GN=Psmc2 PE=1 SV=5 |
| 1433T_MOUSE | 14-3-3 protein theta OS=Mus musculus OX=10090 GN=Ywhaq PE=1 SV=1 |
| CAMKV_MOUSE | CaM kinase-like vesicle-associated protein OS=Mus musculus OX=10090 GN=Camkv PE=1 SV=2 |
| COPA_MOUSE | Coatomer subunit alpha OS=Mus musculus OX=10090 GN=Copa PE=1 SV=2 |
| CTNA2_MOUSE | Catenin alpha-2 OS=Mus musculus OX=10090 GN=Ctnna2 PE=1 SV=3 |
| PGAP1_MOUSE | GPI inositol-deacylase OS=Mus musculus OX=10090 GN=Pgap1 PE=1 SV=3 |
| SYNJ1_MOUSE | Synaptojanin-1 OS=Mus musculus OX=10090 GN=Synj1 PE=1 SV=3 |
| BACH_MOUSE | Cytosolic acyl coenzyme A thioester hydrolase OS=Mus musculus OX=10090 GN=Acot7 PE=1 SV=2 |
| DPYL4_MOUSE | Dihydropyrimidinase-related protein 4 OS=Mus musculus OX=10090 GN=Dpysl4 PE=1 SV=1 |
| E41L3_MOUSE | Band 4.1-like protein 3 OS=Mus musculus OX=10090 GN=Epb41l3 PE=1 SV=1 |
| SYNC_MOUSE | Asparagine--tRNA ligase, cytoplasmic OS=Mus musculus OX=10090 GN=Nars PE=1 SV=2 |

| | |
|---|---|
| VATH_MOUSE | V-type proton ATPase subunit H OS=Mus musculus OX=10090 GN=Atp6v1h PE=1 SV=1 |
| E41L1_MOUSE | Band 4.1-like protein 1 OS=Mus musculus OX=10090 GN=Epb41l1 PE=1 SV=2 |
| IPO5_MOUSE | Importin-5 OS=Mus musculus OX=10090 GN=Ipo5 PE=1 SV=3 |
| SEPT7_MOUSE | Septin-7 OS=Mus musculus OX=10090 GN=Septin7 PE=1 SV=1 |
| SND1_MOUSE | Staphylococcal nuclease domain-containing protein 1 OS=Mus musculus OX=10090 GN=Snd1 PE=1 SV=1 |
| TOM70_MOUSE | Mitochondrial import receptor subunit TOM70 OS=Mus musculus OX=10090 GN=Tomm70 PE=1 SV=2 |
| PP1B_MOUSE | Serine/threonine-protein phosphatase PP1-beta catalytic subunit OS=Mus musculus OX=10090 GN=Ppp1cb PE=1 SV=3 |
| PRDX1_MOUSE | Peroxiredoxin-1 OS=Mus musculus OX=10090 GN=Prdx1 PE=1 SV=1 |
| VA0D1_MOUSE | V-type proton ATPase subunit d 1 OS=Mus musculus OX=10090 GN=Atp6v0d1 PE=1 SV=2 |
| ALDH2_MOUSE | Aldehyde dehydrogenase, mitochondrial OS=Mus musculus OX=10090 GN=Aldh2 PE=1 SV=1 |
| IDH3A_MOUSE | Isocitrate dehydrogenase [NAD] subunit alpha, mitochondrial OS=Mus musculus OX=10090 GN=Idh3a PE=1 SV=1 |
| PSMD3_MOUSE | 26S proteasome non-ATPase regulatory subunit 3 OS=Mus musculus OX=10090 GN=Psmd3 PE=1 SV=3 |
| RL7_MOUSE | 60S ribosomal protein L7 OS=Mus musculus OX=10090 GN=Rpl7 PE=1 SV=2 |
| VATC1_MOUSE | V-type proton ATPase subunit C 1 OS=Mus musculus OX=10090 GN=Atp6v1c1 PE=1 SV=4 |
| CLAP2_MOUSE | CLIP-associating protein 2 OS=Mus musculus OX=10090 GN=Clasp2 PE=1 SV=1 |
| ENOG_MOUSE | Gamma-enolase OS=Mus musculus OX=10090 GN=Eno2 PE=1 SV=2 |
| MYEF2_MOUSE | Myelin expression factor 2 OS=Mus musculus OX=10090 GN=Myef2 PE=1 SV=1 |
| OAT_MOUSE | Ornithine aminotransferase, mitochondrial OS=Mus musculus OX=10090 GN=Oat PE=1 SV=1 |

| | |
|---|---|
| PRDX2_MOUSE | Peroxiredoxin-2 OS=Mus musculus OX=10090 GN=Prdx2 PE=1 SV=3 |
| PROF1_MOUSE | Profilin-1 OS=Mus musculus OX=10090 GN=Pfn1 PE=1 SV=2 |
| STX1B_MOUSE | Syntaxin-1B OS=Mus musculus OX=10090 GN=Stx1b PE=1 SV=1 |
| AP3B2_MOUSE | AP-3 complex subunit beta-2 OS=Mus musculus OX=10090 GN=Ap3b2 PE=1 SV=2 |
| ARF1_MOUSE | ADP-ribosylation factor 1 OS=Mus musculus OX=10090 GN=Arf1 PE=1 SV=2 |
| CTNB1_HUMAN | Catenin beta-1 OS=Homo sapiens OX=9606 GN=CTNNB1 PE=1 SV=1 |
| NCKP1_MOUSE | Nck-associated protein 1 OS=Mus musculus OX=10090 GN=Nckap1 PE=1 SV=2 |
| NEUM_MOUSE | Neuromodulin OS=Mus musculus OX=10090 GN=Gap43 PE=1 SV=1 |
| RS3A_MOUSE | 40S ribosomal protein S3a OS=Mus musculus OX=10090 GN=Rps3a PE=1 SV=3 |
| RTN1_MOUSE | Reticulon-1 OS=Mus musculus OX=10090 GN=Rtn1 PE=1 SV=1 |
| TIF1B_MOUSE | Transcription intermediary factor 1-beta OS=Mus musculus OX=10090 GN=Trim28 PE=1 SV=3 |
| 1433B_MOUSE | 14-3-3 protein beta/alpha OS=Mus musculus OX=10090 GN=Ywhab PE=1 SV=3 |
| AT5F1_MOUSE | ATP synthase F(0) complex subunit B1, mitochondrial OS=Mus musculus OX=10090 GN=Atp5pb PE=1 SV=1 |
| CAPZB_MOUSE | F-actin-capping protein subunit beta OS=Mus musculus OX=10090 GN=Capzb PE=1 SV=3 |
| HMCS1_MOUSE | Hydroxymethylglutaryl-CoA synthase, cytoplasmic OS=Mus musculus OX=10090 GN=Hmgcs1 PE=1 SV=1 |
| PRS8_MOUSE | 26S proteasome regulatory subunit 8 OS=Mus musculus OX=10090 GN=Psmc5 PE=1 SV=1 |
| RL7A_MOUSE | 60S ribosomal protein L7a OS=Mus musculus OX=10090 GN=Rpl7a PE=1 SV=2 |
| ACTZ_MOUSE | Alpha-centractin OS=Mus musculus OX=10090 GN=Actr1a PE=1 SV=1 |
| MTCH2_MOUSE | Mitochondrial carrier homolog 2 OS=Mus musculus OX=10090 GN=Mtch2 PE=1 SV=1 |
| PSA1_MOUSE | Proteasome subunit alpha type-1 OS=Mus musculus OX=10090 GN=Psma1 PE=1 SV=1 |

| | |
|---|---|
| SRSF1_MOUSE | Serine/arginine-rich splicing factor 1 OS=Mus musculus OX=10090 GN=Srsf1 PE=1 SV=3 |
| SYDC_MOUSE | Aspartate--tRNA ligase, cytoplasmic OS=Mus musculus OX=10090 GN=Dars PE=1 SV=2 |
| AMPH_MOUSE | Amphiphysin OS=Mus musculus OX=10090 GN=Amph PE=1 SV=1 |
| ATPO_MOUSE | ATP synthase subunit O, mitochondrial OS=Mus musculus OX=10090 GN=Atp5po PE=1 SV=1 |
| PP2BA_MOUSE | Serine/threonine-protein phosphatase 2B catalytic subunit alpha isoform OS=Mus musculus OX=10090 GN=Ppp3ca PE=1 SV=1 |
| RTCB_MOUSE | tRNA-splicing ligase RtcB homolog OS=Mus musculus OX=10090 GN=Rtcb PE=1 SV=1 |
| SCRN1_MOUSE | Secernin-1 OS=Mus musculus OX=10090 GN=Scrn1 PE=1 SV=1 |
| ADDA_MOUSE | Alpha-adducin OS=Mus musculus OX=10090 GN=Add1 PE=1 SV=2 |
| BDH_MOUSE | D-beta-hydroxybutyrate dehydrogenase, mitochondrial OS=Mus musculus OX=10090 GN=Bdh1 PE=1 SV=2 |
| GFAP_MOUSE | Glial fibrillary acidic protein OS=Mus musculus OX=10090 GN=Gfap PE=1 SV=4 |
| RS8_MOUSE | 40S ribosomal protein S8 OS=Mus musculus OX=10090 GN=Rps8 PE=1 SV=2 |
| SNAB_MOUSE | Beta-soluble NSF attachment protein OS=Mus musculus OX=10090 GN=Napb PE=1 SV=2 |
| NB5R3_MOUSE | NADH-cytochrome b5 reductase 3 OS=Mus musculus OX=10090 GN=Cyb5r3 PE=1 SV=3 |
| RS2_MOUSE | 40S ribosomal protein S2 OS=Mus musculus OX=10090 GN=Rps2 PE=1 SV=3 |
| ATP5H_MOUSE | ATP synthase subunit d, mitochondrial OS=Mus musculus OX=10090 GN=Atp5pd PE=1 SV=3 |
| IF5A1_MOUSE | Eukaryotic translation initiation factor 5A-1 OS=Mus musculus OX=10090 GN=Eif5a PE=1 SV=2 |
| PSIP1_MOUSE | PC4 and SFRS1-interacting protein OS=Mus musculus OX=10090 GN=Psip1 PE=1 SV=1 |
| SERC_MOUSE | Phosphoserine aminotransferase OS=Mus musculus OX=10090 GN=Psat1 PE=1 SV=1 |
| TAGL_MOUSE | Transgelin OS=Mus musculus OX=10090 GN=Tagln PE=1 SV=3 |

| | |
|---|---|
| 1433F_MOUSE | 14-3-3 protein eta OS=Mus musculus OX=10090 GN=Ywhah PE=1 SV=2 |
| AATC_MOUSE | Aspartate aminotransferase, cytoplasmic OS=Mus musculus OX=10090 GN=Got1 PE=1 SV=3 |
| CON__P15497 | CON__P15497 |
| EAA1_MOUSE | Excitatory amino acid transporter 1 OS=Mus musculus OX=10090 GN=Slc1a3 PE=1 SV=2 |
| PURB_MOUSE | Transcriptional activator protein Pur-beta OS=Mus musculus OX=10090 GN=Purb PE=1 SV=3 |
| SSDH_MOUSE | Succinate-semialdehyde dehydrogenase, mitochondrial OS=Mus musculus OX=10090 GN=Aldh5a1 PE=1 SV=1 |
| VATE1_MOUSE | V-type proton ATPase subunit E 1 OS=Mus musculus OX=10090 GN=Atp6v1e1 PE=1 SV=2 |
| ANXA3_MOUSE | Annexin A3 OS=Mus musculus OX=10090 GN=Anxa3 PE=1 SV=4 |
| ETFA_MOUSE | Electron transfer flavoprotein subunit alpha, mitochondrial OS=Mus musculus OX=10090 GN=Etfa PE=1 SV=2 |
| OST48_MOUSE | Dolichyl-diphosphooligosaccharide--protein glycosyltransferase 48 kDa subunit OS=Mus musculus OX=10090 GN=Ddost PE=1 SV=2 |
| OTUB1_MOUSE | Ubiquitin thioesterase OTUB1 OS=Mus musculus OX=10090 GN=Otub1 PE=1 SV=2 |
| PSD12_MOUSE | 26S proteasome non-ATPase regulatory subunit 12 OS=Mus musculus OX=10090 GN=Psmd12 PE=1 SV=4 |
| UCRI_MOUSE | Cytochrome b-c1 complex subunit Rieske, mitochondrial OS=Mus musculus OX=10090 GN=Uqcrfs1 PE=1 SV=1 |
| CCAR2_MOUSE | Cell cycle and apoptosis regulator protein 2 OS=Mus musculus OX=10090 GN=Ccar2 PE=1 SV=2 |
| CNN3_MOUSE | Calponin-3 OS=Mus musculus OX=10090 GN=Cnn3 PE=1 SV=1 |
| RSSA_MOUSE | 40S ribosomal protein SA OS=Mus musculus OX=10090 GN=Rpsa PE=1 SV=4 |
| SFXN3_MOUSE | Sideroflexin-3 OS=Mus musculus OX=10090 GN=Sfxn3 PE=1 SV=1 |
| TOP1_MOUSE | DNA topoisomerase 1 OS=Mus musculus OX=10090 GN=Top1 PE=1 SV=2 |

| | |
|---|---|
| DHX15_MOUSE | Pre-mRNA-splicing factor ATP-dependent RNA helicase DHX15 OS=Mus musculus OX=10090 GN=Dhx15 PE=1 SV=2 |
| GNAQ_MOUSE | Guanine nucleotide-binding protein G(q) subunit alpha OS=Mus musculus OX=10090 GN=Gnaq PE=1 SV=4 |
| GPC4_MOUSE | Glypican-4 OS=Mus musculus OX=10090 GN=Gpc4 PE=1 SV=2 |
| RAB7A_MOUSE | Ras-related protein Rab-7a OS=Mus musculus OX=10090 GN=Rab7a PE=1 SV=2 |
| RL5_MOUSE | 60S ribosomal protein L5 OS=Mus musculus OX=10090 GN=Rpl5 PE=1 SV=3 |
| SV2A_MOUSE | Synaptic vesicle glycoprotein 2A OS=Mus musculus OX=10090 GN=Sv2a PE=1 SV=1 |
| SYSC_MOUSE | Serine--tRNA ligase, cytoplasmic OS=Mus musculus OX=10090 GN=Sars PE=1 SV=3 |
| SYVC_MOUSE | Valine--tRNA ligase OS=Mus musculus OX=10090 GN=Vars PE=1 SV=1 |
| AT2B2_MOUSE | Plasma membrane calcium-transporting ATPase 2 OS=Mus musculus OX=10090 GN=Atp2b2 PE=1 SV=2 |
| IGHM_MOUSE | Immunoglobulin heavy constant mu OS=Mus musculus OX=10090 GN=Ighm PE=1 SV=2 |
| IPYR_MOUSE | Inorganic pyrophosphatase OS=Mus musculus OX=10090 GN=Ppa1 PE=1 SV=1 |
| LA_MOUSE | Lupus La protein homolog OS=Mus musculus OX=10090 GN=Ssb PE=1 SV=1 |
| SEP11_MOUSE | Septin-11 OS=Mus musculus OX=10090 GN=Septin11 PE=1 SV=4 |
| SYN2_MOUSE | Synapsin-2 OS=Mus musculus OX=10090 GN=Syn2 PE=1 SV=2 |
| U5S1_MOUSE | 116 kDa U5 small nuclear ribonucleoprotein component OS=Mus musculus OX=10090 GN=Eftud2 PE=1 SV=1 |
| CBPE_MOUSE | Carboxypeptidase E OS=Mus musculus OX=10090 GN=Cpe PE=1 SV=2 |
| FUBP2_MOUSE | Far upstream element-binding protein 2 OS=Mus musculus OX=10090 GN=Khsrp PE=1 SV=2 |
| FUMH_MOUSE | Fumarate hydratase, mitochondrial OS=Mus musculus OX=10090 GN=Fh PE=1 SV=3 |
| HYEP_MOUSE | Epoxide hydrolase 1 OS=Mus musculus OX=10090 GN=Ephx1 PE=1 SV=2 |

| | |
|---|---|
| KLC1_MOUSE | Kinesin light chain 1 OS=Mus musculus OX=10090 GN=Klc1 PE=1 SV=3 |
| NDKA_MOUSE | Nucleoside diphosphate kinase A OS=Mus musculus OX=10090 GN=Nme1 PE=1 SV=1 |
| PDIA4_MOUSE | Protein disulfide-isomerase A4 OS=Mus musculus OX=10090 GN=Pdia4 PE=1 SV=3 |
| ADT2_MOUSE | ADP/ATP translocase 2 OS=Mus musculus OX=10090 GN=Slc25a5 PE=1 SV=3 |
| IF4A3_MOUSE | Eukaryotic initiation factor 4A-III OS=Mus musculus OX=10090 GN=Eif4a3 PE=1 SV=3 |
| MFGM_MOUSE | Lactadherin OS=Mus musculus OX=10090 GN=Mfge8 PE=1 SV=3 |
| PDIA6_MOUSE | Protein disulfide-isomerase A6 OS=Mus musculus OX=10090 GN=Pdia6 PE=1 SV=3 |
| RAB6A_MOUSE | Ras-related protein Rab-6A OS=Mus musculus OX=10090 GN=Rab6a PE=1 SV=4 |
| SYLC_MOUSE | Leucine--tRNA ligase, cytoplasmic OS=Mus musculus OX=10090 GN=Lars PE=1 SV=2 |
| VDAC2_MOUSE | Voltage-dependent anion-selective channel protein 2 OS=Mus musculus OX=10090 GN=Vdac2 PE=1 SV=2 |
| HVM02_MOUSE | Ig heavy chain V region 93G7 OS=Mus musculus OX=10090 PE=2 SV=1 |
| ACAD9_MOUSE | Complex I assembly factor ACAD9, mitochondrial OS=Mus musculus OX=10090 GN=Acad9 PE=1 SV=2 |
| DDAH1_MOUSE | N(G),N(G)-dimethylarginine dimethylaminohydrolase 1 OS=Mus musculus OX=10090 GN=Ddah1 PE=1 SV=3 |
| H2AW_MOUSE | Core histone macro-H2A.2 OS=Mus musculus OX=10090 GN=H2afy2 PE=1 SV=3 |
| IF2G_MOUSE | Eukaryotic translation initiation factor 2 subunit 3, X-linked OS=Mus musculus OX=10090 GN=Eif2s3x PE=1 SV=2 |
| SODM_MOUSE | Superoxide dismutase [Mn], mitochondrial OS=Mus musculus OX=10090 GN=Sod2 PE=1 SV=3 |
| TICN2_MOUSE | Testican-2 OS=Mus musculus OX=10090 GN=Spock2 PE=1 SV=1 |
| AT1A2_MOUSE | Sodium/potassium-transporting ATPase subunit alpha-2 OS=Mus musculus OX=10090 GN=Atp1a2 PE=1 SV=1 |

| | |
|---|---|
| FABP5_MOUSE | Fatty acid-binding protein 5 OS=Mus musculus OX=10090 GN=Fabp5 PE=1 SV=3 |
| KINH_MOUSE | Kinesin-1 heavy chain OS=Mus musculus OX=10090 GN=Kif5b PE=1 SV=3 |
| OPA1_MOUSE | Dynamin-like 120 kDa protein, mitochondrial OS=Mus musculus OX=10090 GN=Opa1 PE=1 SV=1 |
| PSD13_MOUSE | 26S proteasome non-ATPase regulatory subunit 13 OS=Mus musculus OX=10090 GN=Psmd13 PE=1 SV=1 |
| RUVB2_MOUSE | RuvB-like 2 OS=Mus musculus OX=10090 GN=Ruvbl2 PE=1 SV=3 |
| SNP25_MOUSE | Synaptosomal-associated protein 25 OS=Mus musculus OX=10090 GN=Snap25 PE=1 SV=1 |
| TADBP_MOUSE | TAR DNA-binding protein 43 OS=Mus musculus OX=10090 GN=Tardbp PE=1 SV=1 |
| ALDR_MOUSE | Aldo-keto reductase family 1 member B1 OS=Mus musculus OX=10090 GN=Akr1b1 PE=1 SV=3 |
| ARC1A_MOUSE | Actin-related protein 2/3 complex subunit 1A OS=Mus musculus OX=10090 GN=Arpc1a PE=1 SV=1 |
| C1QC_MOUSE | Complement C1q subcomponent subunit C OS=Mus musculus OX=10090 GN=C1qc PE=1 SV=2 |
| COPG1_MOUSE | Coatomer subunit gamma-1 OS=Mus musculus OX=10090 GN=Copg1 PE=1 SV=1 |
| PRDX3_MOUSE | Thioredoxin-dependent peroxide reductase, mitochondrial OS=Mus musculus OX=10090 GN=Prdx3 PE=1 SV=1 |
| PUR6_MOUSE | Multifunctional protein ADE2 OS=Mus musculus OX=10090 GN=Paics PE=1 SV=4 |
| RAB3C_MOUSE | Ras-related protein Rab-3C OS=Mus musculus OX=10090 GN=Rab3c PE=1 SV=1 |
| RMXL1_MOUSE | RNA binding motif protein, X-linked-like-1 OS=Mus musculus OX=10090 GN=Rbmxl1 PE=2 SV=1 |
| S12A5_MOUSE | Solute carrier family 12 member 5 OS=Mus musculus OX=10090 GN=Slc12a5 PE=1 SV=2 |
| YBOX1_MOUSE | Y-box-binding protein 1 OS=Mus musculus OX=10090 GN=Ybx1 PE=1 SV=3 |

| | |
|---|---|
| ETFB_MOUSE | Electron transfer flavoprotein subunit beta OS=Mus musculus OX=10090 GN=Etfb PE=1 SV=3 |
| GBB2_MOUSE | Guanine nucleotide-binding protein G(I)/G(S)/G(T) subunit beta-2 OS=Mus musculus OX=10090 GN=Gnb2 PE=1 SV=3 |
| HP1B3_MOUSE | Heterochromatin protein 1-binding protein 3 OS=Mus musculus OX=10090 GN=Hp1bp3 PE=1 SV=1 |
| ILF3_MOUSE | Interleukin enhancer-binding factor 3 OS=Mus musculus OX=10090 GN=Ilf3 PE=1 SV=2 |
| PA2G4_MOUSE | Proliferation-associated protein 2G4 OS=Mus musculus OX=10090 GN=Pa2g4 PE=1 SV=3 |
| VDAC3_MOUSE | Voltage-dependent anion-selective channel protein 3 OS=Mus musculus OX=10090 GN=Vdac3 PE=1 SV=1 |
| 6PGD_MOUSE | 6-phosphogluconate dehydrogenase, decarboxylating OS=Mus musculus OX=10090 GN=Pgd PE=1 SV=3 |
| CY1_MOUSE | Cytochrome c1, heme protein, mitochondrial OS=Mus musculus OX=10090 GN=Cyc1 PE=1 SV=1 |
| PRS6A_MOUSE | 26S proteasome regulatory subunit 6A OS=Mus musculus OX=10090 GN=Psmc3 PE=1 SV=2 |
| RAB5C_MOUSE | Ras-related protein Rab-5C OS=Mus musculus OX=10090 GN=Rab5c PE=1 SV=2 |
| RS27A_MOUSE | Ubiquitin-40S ribosomal protein S27a OS=Mus musculus OX=10090 GN=Rps27a PE=1 SV=2 |
| VISL1_MOUSE | Visinin-like protein 1 OS=Mus musculus OX=10090 GN=Vsnl1 PE=1 SV=2 |
| AGRIN_MOUSE | Agrin OS=Mus musculus OX=10090 GN=Agrn PE=1 SV=1 |
| EIF3C_MOUSE | Eukaryotic translation initiation factor 3 subunit C OS=Mus musculus OX=10090 GN=Eif3c PE=1 SV=1 |
| ESTD_MOUSE | S-formylglutathione hydrolase OS=Mus musculus OX=10090 GN=Esd PE=1 SV=1 |
| KAD1_MOUSE | Adenylate kinase isoenzyme 1 OS=Mus musculus OX=10090 GN=Ak1 PE=1 SV=1 |
| PPCE_MOUSE | Prolyl endopeptidase OS=Mus musculus OX=10090 GN=Prep PE=1 SV=1 |
| PRS10_MOUSE | 26S proteasome regulatory subunit 10B OS=Mus musculus OX=10090 GN=Psmc6 PE=1 SV=1 |

| | |
|---|---|
| ARP2_MOUSE | Actin-related protein 2 OS=Mus musculus OX=10090 GN=Actr2 PE=1 SV=1 |
| ARPC2_MOUSE | Actin-related protein 2/3 complex subunit 2 OS=Mus musculus OX=10090 GN=Arpc2 PE=1 SV=3 |
| COPB_MOUSE | Coatomer subunit beta OS=Mus musculus OX=10090 GN=Copb1 PE=1 SV=1 |
| ECHM_MOUSE | Enoyl-CoA hydratase, mitochondrial OS=Mus musculus OX=10090 GN=Echs1 PE=1 SV=1 |
| GDIR1_MOUSE | Rho GDP-dissociation inhibitor 1 OS=Mus musculus OX=10090 GN=Arhgdia PE=1 SV=3 |
| GRIN1_MOUSE | G protein-regulated inducer of neurite outgrowth 1 OS=Mus musculus OX=10090 GN=Gprin1 PE=1 SV=2 |
| HS12A_MOUSE | Heat shock 70 kDa protein 12A OS=Mus musculus OX=10090 GN=Hspa12a PE=1 SV=1 |
| P5CS_MOUSE | Delta-1-pyrroline-5-carboxylate synthase OS=Mus musculus OX=10090 GN=Aldh18a1 PE=1 SV=2 |
| PRS6B_MOUSE | 26S proteasome regulatory subunit 6B OS=Mus musculus OX=10090 GN=Psmc4 PE=1 SV=2 |
| RLA0_MOUSE | 60S acidic ribosomal protein P0 OS=Mus musculus OX=10090 GN=Rplp0 PE=1 SV=3 |
| RUFY3_MOUSE | Protein RUFY3 OS=Mus musculus OX=10090 GN=Rufy3 PE=1 SV=1 |
| SERA_MOUSE | D-3-phosphoglycerate dehydrogenase OS=Mus musculus OX=10090 GN=Phgdh PE=1 SV=3 |
| AP3D1_MOUSE | AP-3 complex subunit delta-1 OS=Mus musculus OX=10090 GN=Ap3d1 PE=1 SV=1 |
| CBR1_MOUSE | Carbonyl reductase [NADPH] 1 OS=Mus musculus OX=10090 GN=Cbr1 PE=1 SV=3 |
| CTBP1_MOUSE | C-terminal-binding protein 1 OS=Mus musculus OX=10090 GN=Ctbp1 PE=1 SV=2 |
| DCX_MOUSE | Neuronal migration protein doublecortin OS=Mus musculus OX=10090 GN=Dcx PE=1 SV=1 |
| DHB12_MOUSE | Very-long-chain 3-oxoacyl-CoA reductase OS=Mus musculus OX=10090 GN=Hsd17b12 PE=1 SV=1 |
| FPPS_MOUSE | Farnesyl pyrophosphate synthase OS=Mus musculus OX=10090 GN=Fdps PE=1 SV=1 |

| | |
|---|---|
| HNRPC_MOUSE | Heterogeneous nuclear ribonucleoproteins C1/C2 OS=Mus musculus OX=10090 GN=Hnrnpc PE=1 SV=1 |
| MRP_MOUSE | MARCKS-related protein OS=Mus musculus OX=10090 GN=Marcksl1 PE=1 SV=2 |
| NDUA9_MOUSE | NADH dehydrogenase [ubiquinone] 1 alpha subcomplex subunit 9, mitochondrial OS=Mus musculus OX=10090 GN=Ndufa9 PE=1 SV=2 |
| TPP2_MOUSE | Tripeptidyl-peptidase 2 OS=Mus musculus OX=10090 GN=Tpp2 PE=1 SV=3 |
| MAP4_MOUSE | Microtubule-associated protein 4 OS=Mus musculus OX=10090 GN=Map4 PE=1 SV=3 |
| PGRC1_MOUSE | Membrane-associated progesterone receptor component 1 OS=Mus musculus OX=10090 GN=Pgrmc1 PE=1 SV=4 |
| PHIPL_MOUSE | Phytanoyl-CoA hydroxylase-interacting protein-like OS=Mus musculus OX=10090 GN=Phyhipl PE=1 SV=1 |
| PSB2_MOUSE | Proteasome subunit beta type-2 OS=Mus musculus OX=10090 GN=Psmb2 PE=1 SV=1 |
| RL12_MOUSE | 60S ribosomal protein L12 OS=Mus musculus OX=10090 GN=Rpl12 PE=1 SV=2 |
| TRAP1_MOUSE | Heat shock protein 75 kDa, mitochondrial OS=Mus musculus OX=10090 GN=Trap1 PE=1 SV=1 |
| CAZA2_MOUSE | F-actin-capping protein subunit alpha-2 OS=Mus musculus OX=10090 GN=Capza2 PE=1 SV=3 |
| CLUS_MOUSE | Clusterin OS=Mus musculus OX=10090 GN=Clu PE=1 SV=1 |
| COPB2_MOUSE | Coatomer subunit beta' OS=Mus musculus OX=10090 GN=Copb2 PE=1 SV=2 |
| IGG2A_RAT | Ig gamma-2A chain C region OS=Rattus norvegicus OX=10116 GN=Igg-2a PE=1 SV=1 |
| IPO7_HUMAN | Importin-7 OS=Homo sapiens OX=9606 GN=IPO7 PE=1 SV=1 |
| LMNA_MOUSE | Prelamin-A/C OS=Mus musculus OX=10090 GN=Lmna PE=1 SV=2 |
| RAN_MOUSE | GTP-binding nuclear protein Ran OS=Mus musculus OX=10090 GN=Ran PE=1 SV=3 |
| ANM1_MOUSE | Protein arginine N-methyltransferase 1 OS=Mus musculus OX=10090 GN=Prmt1 PE=1 SV=1 |

| | |
|---|---|
| COX41_MOUSE | Cytochrome c oxidase subunit 4 isoform 1, mitochondrial OS=Mus musculus OX=10090 GN=Cox4i1 PE=1 SV=2 |
| GSTP1_MOUSE | Glutathione S-transferase P 1 OS=Mus musculus OX=10090 GN=Gstp1 PE=1 SV=2 |
| KAPCB_MOUSE | cAMP-dependent protein kinase catalytic subunit beta OS=Mus musculus OX=10090 GN=Prkacb PE=1 SV=2 |
| NDRG3_MOUSE | Protein NDRG3 OS=Mus musculus OX=10090 GN=Ndrg3 PE=1 SV=1 |
| NDUS2_MOUSE | NADH dehydrogenase [ubiquinone] iron-sulfur protein 2, mitochondrial OS=Mus musculus OX=10090 GN=Ndufs2 PE=1 SV=1 |
| NPM_MOUSE | Nucleophosmin OS=Mus musculus OX=10090 GN=Npm1 PE=1 SV=1 |
| PACN1_MOUSE | Protein kinase C and casein kinase substrate in neurons protein 1 OS=Mus musculus OX=10090 GN=Pacsin1 PE=1 SV=1 |
| SYRC_MOUSE | Arginine--tRNA ligase, cytoplasmic OS=Mus musculus OX=10090 GN=Rars PE=1 SV=2 |
| ALDOC_MOUSE | Fructose-bisphosphate aldolase C OS=Mus musculus OX=10090 GN=Aldoc PE=1 SV=4 |
| HMGB1_MOUSE | High mobility group protein B1 OS=Mus musculus OX=10090 GN=Hmgb1 PE=1 SV=2 |
| HNRH2_MOUSE | Heterogeneous nuclear ribonucleoprotein H2 OS=Mus musculus OX=10090 GN=Hnrnph2 PE=1 SV=1 |
| KCY_MOUSE | UMP-CMP kinase OS=Mus musculus OX=10090 GN=Cmpk1 PE=1 SV=1 |
| NCALD_MOUSE | Neurocalcin-delta OS=Mus musculus OX=10090 GN=Ncald PE=1 SV=4 |
| RS18_MOUSE | 40S ribosomal protein S18 OS=Mus musculus OX=10090 GN=Rps18 PE=1 SV=3 |
| SF3B1_MOUSE | Splicing factor 3B subunit 1 OS=Mus musculus OX=10090 GN=Sf3b1 PE=1 SV=1 |
| THOP1_MOUSE | Thimet oligopeptidase OS=Mus musculus OX=10090 GN=Thop1 PE=1 SV=1 |
| TOM22_MOUSE | Mitochondrial import receptor subunit TOM22 homolog OS=Mus musculus OX=10090 GN=Tomm22 PE=1 SV=3 |
| AKAP5_MOUSE | A-kinase anchor protein 5 OS=Mus musculus OX=10090 GN=Akap5 PE=1 SV=2 |

| | |
|---|---|
| ASNS_MOUSE | Asparagine synthetase [glutamine-hydrolyzing] OS=Mus musculus OX=10090 GN=Asns PE=1 SV=3 |
| AT8A1_MOUSE | Phospholipid-transporting ATPase IA OS=Mus musculus OX=10090 GN=Atp8a1 PE=1 SV=2 |
| CBX3_MOUSE | Chromobox protein homolog 3 OS=Mus musculus OX=10090 GN=Cbx3 PE=1 SV=2 |
| CDC42_CHICK | Cell division control protein 42 homolog OS=Gallus gallus OX=9031 GN=CDC42 PE=2 SV=1 |
| DDX6_MOUSE | Probable ATP-dependent RNA helicase DDX6 OS=Mus musculus OX=10090 GN=Ddx6 PE=1 SV=1 |
| DPP6_MOUSE | Dipeptidyl aminopeptidase-like protein 6 OS=Mus musculus OX=10090 GN=Dpp6 PE=1 SV=1 |
| HCD2_MOUSE | 3-hydroxyacyl-CoA dehydrogenase type-2 OS=Mus musculus OX=10090 GN=Hsd17b10 PE=1 SV=4 |
| HNRPD_MOUSE | Heterogeneous nuclear ribonucleoprotein D0 OS=Mus musculus OX=10090 GN=Hnrnpd PE=1 SV=2 |
| KCC2D_MOUSE | Calcium/calmodulin-dependent protein kinase type II subunit delta OS=Mus musculus OX=10090 GN=Camk2d PE=1 SV=1 |
| NOP56_MOUSE | Nucleolar protein 56 OS=Mus musculus OX=10090 GN=Nop56 PE=1 SV=2 |
| PGM1_MOUSE | Phosphoglucomutase-1 OS=Mus musculus OX=10090 GN=Pgm1 PE=1 SV=4 |
| PSB5_MOUSE | Proteasome subunit beta type-5 OS=Mus musculus OX=10090 GN=Psmb5 PE=1 SV=3 |
| RAB10_MOUSE | Ras-related protein Rab-10 OS=Mus musculus OX=10090 GN=Rab10 PE=1 SV=1 |
| RL9_MOUSE | 60S ribosomal protein L9 OS=Mus musculus OX=10090 GN=Rpl9 PE=2 SV=2 |
| VGF_MOUSE | Neurosecretory protein VGF OS=Mus musculus OX=10090 GN=Vgf PE=1 SV=1 |
| ACADV_MOUSE | Very long-chain specific acyl-CoA dehydrogenase, mitochondrial OS=Mus musculus OX=10090 GN=Acadvl PE=1 SV=3 |
| DDX17_MOUSE | Probable ATP-dependent RNA helicase DDX17 OS=Mus musculus OX=10090 GN=Ddx17 PE=1 SV=1 |

| | |
|---|---|
| EF1A2_MOUSE | Elongation factor 1-alpha 2 OS=Mus musculus OX=10090 GN=Eef1a2 PE=1 SV=1 |
| EIF3E_HUMAN | Eukaryotic translation initiation factor 3 subunit E OS=Homo sapiens OX=9606 GN=EIF3E PE=1 SV=1 |
| MARE1_MOUSE | Microtubule-associated protein RP/EB family member 1 OS=Mus musculus OX=10090 GN=Mapre1 PE=1 SV=3 |
| MMSA_MOUSE | Methylmalonate-semialdehyde dehydrogenase [acylating], mitochondrial OS=Mus musculus OX=10090 GN=Aldh6a1 PE=1 SV=1 |
| NDUV1_MOUSE | NADH dehydrogenase [ubiquinone] flavoprotein 1, mitochondrial OS=Mus musculus OX=10090 GN=Ndufv1 PE=1 SV=1 |
| NOMO1_MOUSE | Nodal modulator 1 OS=Mus musculus OX=10090 GN=Nomo1 PE=1 SV=1 |
| PEA15_MOUSE | Astrocytic phosphoprotein PEA-15 OS=Mus musculus OX=10090 GN=Pea15 PE=1 SV=1 |
| PRS4_MOUSE | 26S proteasome regulatory subunit 4 OS=Mus musculus OX=10090 GN=Psmc1 PE=1 SV=1 |
| PSMD6_MOUSE | 26S proteasome non-ATPase regulatory subunit 6 OS=Mus musculus OX=10090 GN=Psmd6 PE=1 SV=1 |
| RB11B_MOUSE | Ras-related protein Rab-11B OS=Mus musculus OX=10090 GN=Rab11b PE=1 SV=3 |
| RL15_MOUSE | 60S ribosomal protein L15 OS=Mus musculus OX=10090 GN=Rpl15 PE=1 SV=4 |
| RS7_MOUSE | 40S ribosomal protein S7 OS=Mus musculus OX=10090 GN=Rps7 PE=2 SV=1 |
| THIC_MOUSE | Acetyl-CoA acetyltransferase, cytosolic OS=Mus musculus OX=10090 GN=Acat2 PE=1 SV=2 |
| VAT1_MOUSE | Synaptic vesicle membrane protein VAT-1 homolog OS=Mus musculus OX=10090 GN=Vat1 PE=1 SV=3 |
| MUG1_MOUSE | Murinoglobulin-1 OS=Mus musculus OX=10090 GN=Mug1 PE=1 SV=3 |
| NBEA_MOUSE | Neurobeachin OS=Mus musculus OX=10090 GN=Nbea PE=1 SV=1 |
| ACTN1_MOUSE | Alpha-actinin-1 OS=Mus musculus OX=10090 GN=Actn1 PE=1 SV=1 |
| AN32A_MOUSE | Acidic leucine-rich nuclear phosphoprotein 32 family member A OS=Mus musculus OX=10090 GN=Anp32a PE=1 SV=1 |

| | |
|---|---|
| CERU_MOUSE | Ceruloplasmin OS=Mus musculus OX=10090 GN=Cp PE=1 SV=2 |
| COR1C_MOUSE | Coronin-1C OS=Mus musculus OX=10090 GN=Coro1c PE=1 SV=2 |
| ERLN2_MOUSE | Erlin-2 OS=Mus musculus OX=10090 GN=Erlin2 PE=1 SV=1 |
| FUS_MOUSE | RNA-binding protein FUS OS=Mus musculus OX=10090 GN=Fus PE=1 SV=1 |
| GUAA_MOUSE | GMP synthase [glutamine-hydrolyzing] OS=Mus musculus OX=10090 GN=Gmps PE=1 SV=2 |
| HS74L_MOUSE | Heat shock 70 kDa protein 4L OS=Mus musculus OX=10090 GN=Hspa4l PE=1 SV=2 |
| NOP58_MOUSE | Nucleolar protein 58 OS=Mus musculus OX=10090 GN=Nop58 PE=1 SV=1 |
| PRP19_MOUSE | Pre-mRNA-processing factor 19 OS=Mus musculus OX=10090 GN=Prpf19 PE=1 SV=1 |
| PSA6_MOUSE | Proteasome subunit alpha type-6 OS=Mus musculus OX=10090 GN=Psma6 PE=1 SV=1 |
| RS11_HUMAN | 40S ribosomal protein S11 OS=Homo sapiens OX=9606 GN=RPS11 PE=1 SV=3 |
| SF3B3_MOUSE | Splicing factor 3B subunit 3 OS=Mus musculus OX=10090 GN=Sf3b3 PE=1 SV=1 |
| CAPR1_MOUSE | Caprin-1 OS=Mus musculus OX=10090 GN=Caprin1 PE=1 SV=2 |
| KHDR1_MOUSE | KH domain-containing, RNA-binding, signal transduction-associated protein 1 OS=Mus musculus OX=10090 GN=Khdrbs1 PE=1 SV=2 |
| NDUAA_MOUSE | NADH dehydrogenase [ubiquinone] 1 alpha subcomplex subunit 10, mitochondrial OS=Mus musculus OX=10090 GN=Ndufa10 PE=1 SV=1 |
| PDXK_MOUSE | Pyridoxal kinase OS=Mus musculus OX=10090 GN=Pdxk PE=1 SV=1 |
| SC23A_MOUSE | Protein transport protein Sec23A OS=Mus musculus OX=10090 GN=Sec23a PE=1 SV=2 |
| GCN1_MOUSE | eIF-2-alpha kinase activator GCN1 OS=Mus musculus OX=10090 GN=Gcn1 PE=1 SV=1 |
| DC1L1_MOUSE | Cytoplasmic dynein 1 light intermediate chain 1 OS=Mus musculus OX=10090 GN=Dync1li1 PE=1 SV=1 |
| DNJA1_MOUSE | DnaJ homolog subfamily A member 1 OS=Mus musculus OX=10090 GN=Dnaja1 PE=1 SV=1 |

| | |
|---|---|
| FINC_MOUSE | Fibronectin OS=Mus musculus OX=10090 GN=Fn1 PE=1 SV=4 |
| H15_MOUSE | Histone H1.5 OS=Mus musculus OX=10090 GN=Hist1h1b PE=1 SV=2 |
| PSD11_MOUSE | 26S proteasome non-ATPase regulatory subunit 11 OS=Mus musculus OX=10090 GN=Psmd11 PE=1 SV=3 |
| RUVB1_MOUSE | RuvB-like 1 OS=Mus musculus OX=10090 GN=Ruvbl1 PE=1 SV=1 |
| SAHH2_MOUSE | S-adenosylhomocysteine hydrolase-like protein 1 OS=Mus musculus OX=10090 GN=Ahcyl1 PE=1 SV=1 |
| SPEE_MOUSE | Spermidine synthase OS=Mus musculus OX=10090 GN=Srm PE=1 SV=1 |
| UBP14_MOUSE | Ubiquitin carboxyl-terminal hydrolase 14 OS=Mus musculus OX=10090 GN=Usp14 PE=1 SV=3 |
| C1QBP_RAT | Complement component 1 Q subcomponent-binding protein, mitochondrial OS=Rattus norvegicus OX=10116 GN=C1qbp PE=1 SV=2 |
| COR1A_MOUSE | Coronin-1A OS=Mus musculus OX=10090 GN=Coro1a PE=1 SV=5 |
| GLYM_MOUSE | Serine hydroxymethyltransferase, mitochondrial OS=Mus musculus OX=10090 GN=Shmt2 PE=1 SV=1 |
| ILF2_MOUSE | Interleukin enhancer-binding factor 2 OS=Mus musculus OX=10090 GN=Ilf2 PE=1 SV=1 |
| PRPS1_MOUSE | Ribose-phosphate pyrophosphokinase 1 OS=Mus musculus OX=10090 GN=Prps1 PE=1 SV=4 |
| RAB1B_MOUSE | Ras-related protein Rab-1B OS=Mus musculus OX=10090 GN=Rab1b PE=1 SV=1 |
| RAB3A_MOUSE | Ras-related protein Rab-3A OS=Mus musculus OX=10090 GN=Rab3a PE=1 SV=1 |
| RL30_MOUSE | 60S ribosomal protein L30 OS=Mus musculus OX=10090 GN=Rpl30 PE=1 SV=2 |
| ALG2_MOUSE | Alpha-1,3/1,6-mannosyltransferase ALG2 OS=Mus musculus OX=10090 GN=Alg2 PE=1 SV=2 |
| COTL1_MOUSE | Coactosin-like protein OS=Mus musculus OX=10090 GN=Cotl1 PE=1 SV=3 |
| DBNL_MOUSE | Drebrin-like protein OS=Mus musculus OX=10090 GN=Dbnl PE=1 SV=2 |

| | |
|---|---|
| DCE1_MOUSE | Glutamate decarboxylase 1 OS=Mus musculus OX=10090 GN=Gad1 PE=1 SV=2 |
| IF2A_MOUSE | Eukaryotic translation initiation factor 2 subunit 1 OS=Mus musculus OX=10090 GN=Eif2s1 PE=1 SV=3 |
| ODPX_MOUSE | Pyruvate dehydrogenase protein X component, mitochondrial OS=Mus musculus OX=10090 GN=Pdhx PE=1 SV=1 |
| PCBP1_MOUSE | Poly(rC)-binding protein 1 OS=Mus musculus OX=10090 GN=Pcbp1 PE=1 SV=1 |
| PFKAP_MOUSE | ATP-dependent 6-phosphofructokinase, platelet type OS=Mus musculus OX=10090 GN=Pfkp PE=1 SV=1 |
| PSA7_MOUSE | Proteasome subunit alpha type-7 OS=Mus musculus OX=10090 GN=Psma7 PE=1 SV=1 |
| TBB4A_MOUSE | Tubulin beta-4A chain OS=Mus musculus OX=10090 GN=Tubb4a PE=1 SV=3 |
| VAPA_MOUSE | Vesicle-associated membrane protein-associated protein A OS=Mus musculus OX=10090 GN=Vapa PE=1 SV=2 |
| VIAAT_MOUSE | Vesicular inhibitory amino acid transporter OS=Mus musculus OX=10090 GN=Slc32a1 PE=1 SV=3 |
| PUR2_MOUSE | Trifunctional purine biosynthetic protein adenosine-3 OS=Mus musculus OX=10090 GN=Gart PE=1 SV=3 |
| PPID_MOUSE | Peptidyl-prolyl cis-trans isomerase D OS=Mus musculus OX=10090 GN=Ppid PE=1 SV=3 |
| ARL8B_MOUSE | ADP-ribosylation factor-like protein 8B OS=Mus musculus OX=10090 GN=Arl8b PE=1 SV=1 |
| DDB1_MOUSE | DNA damage-binding protein 1 OS=Mus musculus OX=10090 GN=Ddb1 PE=1 SV=2 |
| EF1D_MOUSE | Elongation factor 1-delta OS=Mus musculus OX=10090 GN=Eef1d PE=1 SV=3 |
| GLSK_MOUSE | Glutaminase kidney isoform, mitochondrial OS=Mus musculus OX=10090 GN=Gls PE=1 SV=1 |
| GNAS1_MOUSE | Guanine nucleotide-binding protein G(s) subunit alpha isoforms XLas OS=Mus musculus OX=10090 GN=Gnas PE=1 SV=1 |
| KIF2A_MOUSE | Kinesin-like protein KIF2A OS=Mus musculus OX=10090 GN=Kif2a PE=1 SV=2 |

| | |
|---|---|
| RHOA_MOUSE | Transforming protein RhoA OS=Mus musculus OX=10090 GN=Rhoa PE=1 SV=1 |
| SET_MOUSE | Protein SET OS=Mus musculus OX=10090 GN=Set PE=1 SV=1 |
| LAC1_MOUSE | Ig lambda-1 chain C region OS=Mus musculus OX=10090 PE=1 SV=1 |
| VINC_MOUSE | Vinculin OS=Mus musculus OX=10090 GN=Vcl PE=1 SV=4 |
| CSK21_MOUSE | Casein kinase II subunit alpha OS=Mus musculus OX=10090 GN=Csnk2a1 PE=1 SV=2 |
| BLMH_MOUSE | Bleomycin hydrolase OS=Mus musculus OX=10090 GN=Blmh PE=1 SV=1 |
| ECHB_MOUSE | Trifunctional enzyme subunit beta, mitochondrial OS=Mus musculus OX=10090 GN=Hadhb PE=1 SV=1 |
| HNRDL_MOUSE | Heterogeneous nuclear ribonucleoprotein D-like OS=Mus musculus OX=10090 GN=Hnrnpdl PE=1 SV=1 |
| RL23A_MOUSE | 60S ribosomal protein L23a OS=Mus musculus OX=10090 GN=Rpl23a PE=1 SV=1 |
| RS12_MOUSE | 40S ribosomal protein S12 OS=Mus musculus OX=10090 GN=Rps12 PE=1 SV=2 |
| U2AF2_MOUSE | Splicing factor U2AF 65 kDa subunit OS=Mus musculus OX=10090 GN=U2af2 PE=1 SV=3 |
| EMC1_MOUSE | ER membrane protein complex subunit 1 OS=Mus musculus OX=10090 GN=Emc1 PE=1 SV=1 |
| H33_MOUSE | Histone H3.3 OS=Mus musculus OX=10090 GN=H3-3a PE=1 SV=2 |
| HNRPF_MOUSE | Heterogeneous nuclear ribonucleoprotein F OS=Mus musculus OX=10090 GN=Hnrnpf PE=1 SV=3 |
| CYBP_MOUSE | Calcyclin-binding protein OS=Mus musculus OX=10090 GN=Cacybp PE=1 SV=1 |
| ANXA1_MOUSE | Annexin A1 OS=Mus musculus OX=10090 GN=Anxa1 PE=1 SV=2 |
| COX2_LEGFO | Cytochrome c oxidase subunit 2 OS=Leggadina forresti OX=81935 GN=MT-CO2 PE=3 SV=1 |
| DNM3A_MOUSE | DNA (cytosine-5)-methyltransferase 3A OS=Mus musculus OX=10090 GN=Dnmt3a PE=1 SV=2 |
| F10A1_MOUSE | Hsc70-interacting protein OS=Mus musculus OX=10090 GN=St13 PE=1 SV=1 |

| | |
|---|---|
| FA49A_MOUSE | Protein FAM49A OS=Mus musculus OX=10090 GN=Fam49a PE=1 SV=1 |
| HINT1_MOUSE | Histidine triad nucleotide-binding protein 1 OS=Mus musculus OX=10090 GN=Hint1 PE=1 SV=3 |
| ITB1_MOUSE | Integrin beta-1 OS=Mus musculus OX=10090 GN=Itgb1 PE=1 SV=1 |
| JIP3_MOUSE | C-Jun-amino-terminal kinase-interacting protein 3 OS=Mus musculus OX=10090 GN=Mapk8ip3 PE=1 SV=1 |
| KCC2A_MOUSE | Calcium/calmodulin-dependent protein kinase type II subunit alpha OS=Mus musculus OX=10090 GN=Camk2a PE=1 SV=2 |
| MYL6_MOUSE | Myosin light polypeptide 6 OS=Mus musculus OX=10090 GN=Myl6 PE=1 SV=3 |
| NP1L4_MOUSE | Nucleosome assembly protein 1-like 4 OS=Mus musculus OX=10090 GN=Nap1l4 PE=1 SV=1 |
| ODO2_MOUSE | Dihydrolipoyllysine-residue succinyltransferase component of 2-oxoglutarate dehydrogenase complex, mitochondrial OS=Mus musculus OX=10090 GN=Dlst PE=1 SV=1 |
| RS5_MOUSE | 40S ribosomal protein S5 OS=Mus musculus OX=10090 GN=Rps5 PE=1 SV=3 |
| RS6_MOUSE | 40S ribosomal protein S6 OS=Mus musculus OX=10090 GN=Rps6 PE=1 SV=1 |
| STT3A_HUMAN | Dolichyl-diphosphooligosaccharide--protein glycosyltransferase subunit STT3A OS=Homo sapiens OX=9606 GN=STT3A PE=1 SV=2 |
| USO1_MOUSE | General vesicular transport factor p115 OS=Mus musculus OX=10090 GN=Uso1 PE=1 SV=2 |
| IDHG1_MOUSE | Isocitrate dehydrogenase [NAD] subunit gamma 1, mitochondrial OS=Mus musculus OX=10090 GN=Idh3g PE=1 SV=1 |
| H12_MOUSE | Histone H1.2 OS=Mus musculus OX=10090 GN=Hist1h1c PE=1 SV=2 |
| PPIB_MOUSE | Peptidyl-prolyl cis-trans isomerase B OS=Mus musculus OX=10090 GN=Ppib PE=1 SV=2 |
| PRDX5_MOUSE | Peroxiredoxin-5, mitochondrial OS=Mus musculus OX=10090 GN=Prdx5 PE=1 SV=2 |
| PSA5_MOUSE | Proteasome subunit alpha type-5 OS=Mus musculus OX=10090 GN=Psma5 PE=1 SV=1 |

| | |
|---|---|
| PUR9_MOUSE | Bifunctional purine biosynthesis protein PURH OS=Mus musculus OX=10090 GN=Atic PE=1 SV=2 |
| RU17_MOUSE | U1 small nuclear ribonucleoprotein 70 kDa OS=Mus musculus OX=10090 GN=Snrnp70 PE=1 SV=2 |
| STRAP_MOUSE | Serine-threonine kinase receptor-associated protein OS=Mus musculus OX=10090 GN=Strap PE=1 SV=2 |
| TEBP_MOUSE | Prostaglandin E synthase 3 OS=Mus musculus OX=10090 GN=Ptges3 PE=1 SV=1 |
| THIM_MOUSE | 3-ketoacyl-CoA thiolase, mitochondrial OS=Mus musculus OX=10090 GN=Acaa2 PE=1 SV=3 |
| PIPNA_MOUSE | Phosphatidylinositol transfer protein alpha isoform OS=Mus musculus OX=10090 GN=Pitpna PE=1 SV=2 |
| IMDH2_MOUSE | Inosine-5'-monophosphate dehydrogenase 2 OS=Mus musculus OX=10090 GN=Impdh2 PE=1 SV=2 |
| NACAM_MOUSE | Nascent polypeptide-associated complex subunit alpha, muscle-specific form OS=Mus musculus OX=10090 GN=Naca PE=1 SV=2 |
| GSLG1_MOUSE | Golgi apparatus protein 1 OS=Mus musculus OX=10090 GN=Glg1 PE=1 SV=1 |
| ACDSB_MOUSE | Short/branched chain specific acyl-CoA dehydrogenase, mitochondrial OS=Mus musculus OX=10090 GN=Acadsb PE=1 SV=1 |
| SYK_MOUSE | Lysine--tRNA ligase OS=Mus musculus OX=10090 GN=Kars1 PE=1 SV=1 |
| SC6A1_MOUSE | Sodium- and chloride-dependent GABA transporter 1 OS=Mus musculus OX=10090 GN=Slc6a1 PE=1 SV=2 |
| UBR4_MOUSE | E3 ubiquitin-protein ligase UBR4 OS=Mus musculus OX=10090 GN=Ubr4 PE=1 SV=1 |
| AINX_MOUSE | Alpha-internexin OS=Mus musculus OX=10090 GN=Ina PE=1 SV=3 |
| CANB1_MOUSE | Calcineurin subunit B type 1 OS=Mus musculus OX=10090 GN=Ppp3r1 PE=1 SV=3 |
| GNAI1_MOUSE | Guanine nucleotide-binding protein G(i) subunit alpha-1 OS=Mus musculus OX=10090 GN=Gnai1 PE=1 SV=1 |
| MBB1A_MOUSE | Myb-binding protein 1A OS=Mus musculus OX=10090 GN=Mybbp1a PE=1 SV=2 |

| | |
|---|---|
| PSD3_MOUSE | PH and SEC7 domain-containing protein 3 OS=Mus musculus OX=10090 GN=Psd3 PE=1 SV=2 |
| RBBP4_MOUSE | Histone-binding protein RBBP4 OS=Mus musculus OX=10090 GN=Rbbp4 PE=1 SV=5 |
| SUCA_MOUSE | Succinate--CoA ligase [ADP/GDP-forming] subunit alpha, mitochondrial OS=Mus musculus OX=10090 GN=Suclg1 PE=1 SV=4 |
| H2AV_MOUSE | Histone H2A.V OS=Mus musculus OX=10090 GN=H2afv PE=1 SV=3 |
| CKAP5_MOUSE | Cytoskeleton-associated protein 5 OS=Mus musculus OX=10090 GN=Ckap5 PE=1 SV=1 |
| FHL1_MOUSE | Four and a half LIM domains protein 1 OS=Mus musculus OX=10090 GN=Fhl1 PE=1 SV=3 |
| LASP1_MOUSE | LIM and SH3 domain protein 1 OS=Mus musculus OX=10090 GN=Lasp1 PE=1 SV=1 |
| PSA3_MOUSE | Proteasome subunit alpha type-3 OS=Mus musculus OX=10090 GN=Psma3 PE=1 SV=3 |
| RAC1_MOUSE | Ras-related C3 botulinum toxin substrate 1 OS=Mus musculus OX=10090 GN=Rac1 PE=1 SV=1 |
| RL23_MOUSE | 60S ribosomal protein L23 OS=Mus musculus OX=10090 GN=Rpl23 PE=1 SV=1 |
| RL8_MOUSE | 60S ribosomal protein L8 OS=Mus musculus OX=10090 GN=Rpl8 PE=1 SV=2 |
| SGTA_MOUSE | Small glutamine-rich tetratricopeptide repeat-containing protein alpha OS=Mus musculus OX=10090 GN=Sgta PE=1 SV=2 |
| SNAG_MOUSE | Gamma-soluble NSF attachment protein OS=Mus musculus OX=10090 GN=Napg PE=1 SV=1 |
| TRA2B_MOUSE | Transformer-2 protein homolog beta OS=Mus musculus OX=10090 GN=Tra2b PE=1 SV=1 |
| VATD_MOUSE | V-type proton ATPase subunit D OS=Mus musculus OX=10090 GN=Atp6v1d PE=1 SV=1 |
| SYWC_MOUSE | Tryptophan--tRNA ligase, cytoplasmic OS=Mus musculus OX=10090 GN=Wars PE=1 SV=2 |
| RLA2_MOUSE | 60S acidic ribosomal protein P2 OS=Mus musculus OX=10090 GN=Rplp2 PE=1 SV=3 |
| MIF_MOUSE | Macrophage migration inhibitory factor OS=Mus musculus OX=10090 GN=Mif PE=1 SV=2 |

| | |
|---|---|
| BZW1_MOUSE | Basic leucine zipper and W2 domain-containing protein 1 OS=Mus musculus OX=10090 GN=Bzw1 PE=1 SV=1 |
| SYFB_MOUSE | Phenylalanine--tRNA ligase beta subunit OS=Mus musculus OX=10090 GN=Farsb PE=1 SV=2 |
| ARPC3_MOUSE | Actin-related protein 2/3 complex subunit 3 OS=Mus musculus OX=10090 GN=Arpc3 PE=1 SV=3 |
| CAN2_MOUSE | Calpain-2 catalytic subunit OS=Mus musculus OX=10090 GN=Capn2 PE=1 SV=4 |
| DCE2_MOUSE | Glutamate decarboxylase 2 OS=Mus musculus OX=10090 GN=Gad2 PE=1 SV=1 |
| EIF3F_MOUSE | Eukaryotic translation initiation factor 3 subunit F OS=Mus musculus OX=10090 GN=Eif3f PE=1 SV=2 |
| G3BP2_MOUSE | Ras GTPase-activating protein-binding protein 2 OS=Mus musculus OX=10090 GN=G3bp2 PE=1 SV=2 |
| NSF1C_MOUSE | NSFL1 cofactor p47 OS=Mus musculus OX=10090 GN=Nsfl1c PE=1 SV=1 |
| PSA4_MOUSE | Proteasome subunit alpha type-4 OS=Mus musculus OX=10090 GN=Psma4 PE=1 SV=1 |
| PSB7_MOUSE | Proteasome subunit beta type-7 OS=Mus musculus OX=10090 GN=Psmb7 PE=1 SV=1 |
| PYRG1_MOUSE | CTP synthase 1 OS=Mus musculus OX=10090 GN=Ctps1 PE=1 SV=2 |
| TSN_MOUSE | Translin OS=Mus musculus OX=10090 GN=Tsn PE=1 SV=1 |
| SRC8_MOUSE | Src substrate cortactin OS=Mus musculus OX=10090 GN=Cttn PE=1 SV=2 |
| SYFA_MOUSE | Phenylalanine--tRNA ligase alpha subunit OS=Mus musculus OX=10090 GN=Farsa PE=1 SV=1 |
| SYIC_MOUSE | Isoleucine--tRNA ligase, cytoplasmic OS=Mus musculus OX=10090 GN=Iars PE=1 SV=2 |
| EHD1_MOUSE | EH domain-containing protein 1 OS=Mus musculus OX=10090 GN=Ehd1 PE=1 SV=1 |
| VIGLN_MOUSE | Vigilin OS=Mus musculus OX=10090 GN=Hdlbp PE=1 SV=1 |
| UBE2N_MOUSE | Ubiquitin-conjugating enzyme E2 N OS=Mus musculus OX=10090 GN=Ube2n PE=1 SV=1 |

| | |
|---|---|
| IF4A2_MOUSE | Eukaryotic initiation factor 4A-II OS=Mus musculus OX=10090 GN=Eif4a2 PE=1 SV=2 |
| CHD4_MOUSE | Chromodomain-helicase-DNA-binding protein 4 OS=Mus musculus OX=10090 GN=Chd4 PE=1 SV=1 |
| CALU_MOUSE | Calumenin OS=Mus musculus OX=10090 GN=Calu PE=1 SV=1 |
| CUL3_MOUSE | Cullin-3 OS=Mus musculus OX=10090 GN=Cul3 PE=1 SV=1 |
| EIF3L_MOUSE | Eukaryotic translation initiation factor 3 subunit L OS=Mus musculus OX=10090 GN=Eif3l PE=1 SV=1 |
| ELAV1_MOUSE | ELAV-like protein 1 OS=Mus musculus OX=10090 GN=Elavl1 PE=1 SV=2 |
| PAIRB_MOUSE | Plasminogen activator inhibitor 1 RNA-binding protein OS=Mus musculus OX=10090 GN=Serbp1 PE=1 SV=2 |
| RAP1B_MOUSE | Ras-related protein Rap-1b OS=Mus musculus OX=10090 GN=Rap1b PE=1 SV=2 |
| RL11_MOUSE | 60S ribosomal protein L11 OS=Mus musculus OX=10090 GN=Rpl11 PE=1 SV=4 |
| RS16_MOUSE | 40S ribosomal protein S16 OS=Mus musculus OX=10090 GN=Rps16 PE=1 SV=4 |
| SDHB_MOUSE | Succinate dehydrogenase [ubiquinone] iron-sulfur subunit, mitochondrial OS=Mus musculus OX=10090 GN=Sdhb PE=1 SV=1 |
| STX1A_MOUSE | Syntaxin-1A OS=Mus musculus OX=10090 GN=Stx1a PE=1 SV=3 |
| TALDO_MOUSE | Transaldolase OS=Mus musculus OX=10090 GN=Taldo1 PE=1 SV=2 |
| TXNL1_MOUSE | Thioredoxin-like protein 1 OS=Mus musculus OX=10090 GN=Txnl1 PE=1 SV=3 |
| PSPC1_MOUSE | Paraspeckle component 1 OS=Mus musculus OX=10090 GN=Pspc1 PE=1 SV=1 |
| FARP1_MOUSE | FERM, ARHGEF and pleckstrin domain-containing protein 1 OS=Mus musculus OX=10090 GN=Farp1 PE=1 SV=1 |
| S61A1_MOUSE | Protein transport protein Sec61 subunit alpha isoform 1 OS=Mus musculus OX=10090 GN=Sec61a1 PE=1 SV=2 |
| UGGG1_MOUSE | UDP-glucose:glycoprotein glucosyltransferase 1 OS=Mus musculus OX=10090 GN=Uggt1 PE=1 SV=4 |

| | |
|---|---|
| GLOD4_MOUSE | Glyoxalase domain-containing protein 4 OS=Mus musculus OX=10090 GN=Glod4 PE=1 SV=1 |
| DPP3_MOUSE | Dipeptidyl peptidase 3 OS=Mus musculus OX=10090 GN=Dpp3 PE=1 SV=2 |
| SKP1_HUMAN | S-phase kinase-associated protein 1 OS=Homo sapiens OX=9606 GN=SKP1 PE=1 SV=2 |
| KAD3_MOUSE | GTP:AMP phosphotransferase AK3, mitochondrial OS=Mus musculus OX=10090 GN=Ak3 PE=1 SV=3 |
| KPCA_MOUSE | Protein kinase C alpha type OS=Mus musculus OX=10090 GN=Prkca PE=1 SV=3 |
| ITAV_MOUSE | Integrin alpha-V OS=Mus musculus OX=10090 GN=Itgav PE=1 SV=2 |
| ADDB_MOUSE | Beta-adducin OS=Mus musculus OX=10090 GN=Add2 PE=1 SV=4 |
| CNDP2_MOUSE | Cytosolic non-specific dipeptidase OS=Mus musculus OX=10090 GN=Cndp2 PE=1 SV=1 |
| CON__P02533 | CON__P02533 |
| H10_MOUSE | Histone H1.0 OS=Mus musculus OX=10090 GN=H1f0 PE=2 SV=4 |
| KCRU_MOUSE | Creatine kinase U-type, mitochondrial OS=Mus musculus OX=10090 GN=Ckmt1 PE=1 SV=1 |
| LAT1_MOUSE | Large neutral amino acids transporter small subunit 1 OS=Mus musculus OX=10090 GN=Slc7a5 PE=1 SV=2 |
| EF1B_MOUSE | Elongation factor 1-beta OS=Mus musculus OX=10090 GN=Eef1b PE=1 SV=5 |
| TCTP_MOUSE | Translationally-controlled tumor protein OS=Mus musculus OX=10090 GN=Tpt1 PE=1 SV=1 |
| TNPO2_MOUSE | Transportin-2 OS=Mus musculus OX=10090 GN=Tnpo2 PE=1 SV=1 |
| OGA_MOUSE | Protein O-GlcNAcase OS=Mus musculus OX=10090 GN=Oga PE=1 SV=2 |
| AK1A1_MOUSE | Aldo-keto reductase family 1 member A1 OS=Mus musculus OX=10090 GN=Akr1a1 PE=1 SV=3 |
| APEX1_MOUSE | DNA-(apurinic or apyrimidinic site) lyase OS=Mus musculus OX=10090 GN=Apex1 PE=1 SV=2 |
| ASNA_MOUSE | ATPase Asna1 OS=Mus musculus OX=10090 GN=Asna1 PE=1 SV=2 |

| | |
|---|---|
| LONM_MOUSE | Lon protease homolog, mitochondrial OS=Mus musculus OX=10090 GN=Lonp1 PE=1 SV=2 |
| LRC59_MOUSE | Leucine-rich repeat-containing protein 59 OS=Mus musculus OX=10090 GN=Lrrc59 PE=1 SV=1 |
| PLPR4_MOUSE | Phospholipid phosphatase-related protein type 4 OS=Mus musculus OX=10090 GN=Plppr4 PE=1 SV=2 |
| PSA2_MOUSE | Proteasome subunit alpha type-2 OS=Mus musculus OX=10090 GN=Psma2 PE=1 SV=3 |
| PTN11_MOUSE | Tyrosine-protein phosphatase non-receptor type 11 OS=Mus musculus OX=10090 GN=Ptpn11 PE=1 SV=2 |
| RS19_MOUSE | 40S ribosomal protein S19 OS=Mus musculus OX=10090 GN=Rps19 PE=1 SV=3 |
| SNP47_MOUSE | Synaptosomal-associated protein 47 OS=Mus musculus OX=10090 GN=Snap47 PE=1 SV=1 |
| TR150_MOUSE | Thyroid hormone receptor-associated protein 3 OS=Mus musculus OX=10090 GN=Thrap3 PE=1 SV=1 |
| THIKA_MOUSE | 3-ketoacyl-CoA thiolase A, peroxisomal OS=Mus musculus OX=10090 GN=Acaa1a PE=1 SV=1 |
| VP26B_MOUSE | Vacuolar protein sorting-associated protein 26B OS=Mus musculus OX=10090 GN=Vps26b PE=1 SV=1 |
| AP1G1_MOUSE | AP-1 complex subunit gamma-1 OS=Mus musculus OX=10090 GN=Ap1g1 PE=1 SV=3 |
| FUBP1_MOUSE | Far upstream element-binding protein 1 OS=Mus musculus OX=10090 GN=Fubp1 PE=1 SV=1 |
| APLP2_HUMAN | Amyloid-like protein 2 OS=Homo sapiens OX=9606 GN=APLP2 PE=1 SV=2 |
| PSME1_MOUSE | Proteasome activator complex subunit 1 OS=Mus musculus OX=10090 GN=Psme1 PE=1 SV=2 |
| MY18A_MOUSE | Unconventional myosin-XVIIIa OS=Mus musculus OX=10090 GN=Myo18a PE=1 SV=2 |
| HUWE1_MOUSE | E3 ubiquitin-protein ligase HUWE1 OS=Mus musculus OX=10090 GN=Huwe1 PE=1 SV=5 |
| PLXA4_MOUSE | Plexin-A4 OS=Mus musculus OX=10090 GN=Plxna4 PE=1 SV=3 |
| CH10_MOUSE | 10 kDa heat shock protein, mitochondrial OS=Mus musculus OX=10090 GN=Hspe1 PE=1 SV=2 |

| | |
|---|---|
| HNRLL_MOUSE | Heterogeneous nuclear ribonucleoprotein L-like OS=Mus musculus OX=10090 GN=Hnrnpll PE=1 SV=3 |
| LEG1_MOUSE | Galectin-1 OS=Mus musculus OX=10090 GN=Lgals1 PE=1 SV=3 |
| MTPN_MOUSE | Myotrophin OS=Mus musculus OX=10090 GN=Mtpn PE=1 SV=2 |
| NPTN_MOUSE | Neuroplastin OS=Mus musculus OX=10090 GN=Nptn PE=1 SV=3 |
| QCR7_MOUSE | Cytochrome b-c1 complex subunit 7 OS=Mus musculus OX=10090 GN=Uqcrb PE=1 SV=3 |
| RANG_MOUSE | Ran-specific GTPase-activating protein OS=Mus musculus OX=10090 GN=Ranbp1 PE=1 SV=2 |
| RL18_MOUSE | 60S ribosomal protein L18 OS=Mus musculus OX=10090 GN=Rpl18 PE=1 SV=3 |
| STML2_MOUSE | Stomatin-like protein 2, mitochondrial OS=Mus musculus OX=10090 GN=Stoml2 PE=1 SV=1 |
| EPMIP_MOUSE | EPM2A-interacting protein 1 OS=Mus musculus OX=10090 GN=Epm2aip1 PE=1 SV=1 |
| CSN1_MOUSE | COP9 signalosome complex subunit 1 OS=Mus musculus OX=10090 GN=Gps1 PE=1 SV=1 |
| ELAV3_MOUSE | ELAV-like protein 3 OS=Mus musculus OX=10090 GN=Elavl3 PE=1 SV=1 |
| PARK7_MOUSE | Protein/nucleic acid deglycase DJ-1 OS=Mus musculus OX=10090 GN=Park7 PE=1 SV=1 |
| COR1B_MOUSE | Coronin-1B OS=Mus musculus OX=10090 GN=Coro1b PE=1 SV=1 |
| G6PD1_MOUSE | Glucose-6-phosphate 1-dehydrogenase X OS=Mus musculus OX=10090 GN=G6pdx PE=1 SV=3 |
| LETM1_MOUSE | Mitochondrial proton/calcium exchanger protein OS=Mus musculus OX=10090 GN=Letm1 PE=1 SV=1 |
| CA2D1_MOUSE | Voltage-dependent calcium channel subunit alpha-2/delta-1 OS=Mus musculus OX=10090 GN=Cacna2d1 PE=1 SV=1 |
| PLEC_MOUSE | Plectin OS=Mus musculus OX=10090 GN=Plec PE=1 SV=3 |
| ADHX_MOUSE | Alcohol dehydrogenase class-3 OS=Mus musculus OX=10090 GN=Adh5 PE=1 SV=3 |

| | |
|---|---|
| AN32E_MOUSE | Acidic leucine-rich nuclear phosphoprotein 32 family member E OS=Mus musculus OX=10090 GN=Anp32e PE=1 SV=2 |
| COPD_MOUSE | Coatomer subunit delta OS=Mus musculus OX=10090 GN=Arcn1 PE=1 SV=2 |
| IF2B_MOUSE | Eukaryotic translation initiation factor 2 subunit 2 OS=Mus musculus OX=10090 GN=Eif2s2 PE=1 SV=1 |
| MP2K1_MOUSE | Dual specificity mitogen-activated protein kinase kinase 1 OS=Mus musculus OX=10090 GN=Map2k1 PE=1 SV=2 |
| PLMN_MOUSE | Plasminogen OS=Mus musculus OX=10090 GN=Plg PE=1 SV=3 |
| PROF2_MOUSE | Profilin-2 OS=Mus musculus OX=10090 GN=Pfn2 PE=1 SV=3 |
| RL10_MOUSE | 60S ribosomal protein L10 OS=Mus musculus OX=10090 GN=Rpl10 PE=1 SV=3 |
| ROAA_MOUSE | Heterogeneous nuclear ribonucleoprotein A/B OS=Mus musculus OX=10090 GN=Hnrnpab PE=1 SV=1 |
| SNX3_MOUSE | Sorting nexin-3 OS=Mus musculus OX=10090 GN=Snx3 PE=1 SV=3 |
| TPM3_MOUSE | Tropomyosin alpha-3 chain OS=Mus musculus OX=10090 GN=Tpm3 PE=1 SV=3 |
| WDR7_MOUSE | WD repeat-containing protein 7 OS=Mus musculus OX=10090 GN=Wdr7 PE=1 SV=3 |
| IVD_MOUSE | Isovaleryl-CoA dehydrogenase, mitochondrial OS=Mus musculus OX=10090 GN=Ivd PE=1 SV=1 |
| RENT1_MOUSE | Regulator of nonsense transcripts 1 OS=Mus musculus OX=10090 GN=Upf1 PE=1 SV=2 |
| OLA1_MOUSE | Obg-like ATPase 1 OS=Mus musculus OX=10090 GN=Ola1 PE=1 SV=1 |
| SRSF3_MOUSE | Serine/arginine-rich splicing factor 3 OS=Mus musculus OX=10090 GN=Srsf3 PE=1 SV=1 |
| UBQL2_MOUSE | Ubiquilin-2 OS=Mus musculus OX=10090 GN=Ubqln2 PE=1 SV=2 |
| CSN4_MOUSE | COP9 signalosome complex subunit 4 OS=Mus musculus OX=10090 GN=Cops4 PE=1 SV=1 |
| RL21_MOUSE | 60S ribosomal protein L21 OS=Mus musculus OX=10090 GN=Rpl21 PE=1 SV=3 |

| | |
|---|---|
| MLEC_MOUSE | Malectin OS=Mus musculus OX=10090 GN=Mlec PE=1 SV=2 |
| CSDE1_MOUSE | Cold shock domain-containing protein E1 OS=Mus musculus OX=10090 GN=Csde1 PE=1 SV=1 |
| 2ABA_MOUSE | Serine/threonine-protein phosphatase 2A 55 kDa regulatory subunit B alpha isoform OS=Mus musculus OX=10090 GN=Ppp2r2a PE=1 SV=1 |
| HPRT_MOUSE | Hypoxanthine-guanine phosphoribosyltransferase OS=Mus musculus OX=10090 GN=Hprt1 PE=1 SV=3 |
| HVM51_MOUSE | Ig heavy chain V region AC38 205.12 OS=Mus musculus OX=10090 PE=1 SV=1 |
| NDUS3_MOUSE | NADH dehydrogenase [ubiquinone] iron-sulfur protein 3, mitochondrial OS=Mus musculus OX=10090 GN=Ndufs3 PE=1 SV=2 |
| NDUS7_MOUSE | NADH dehydrogenase [ubiquinone] iron-sulfur protein 7, mitochondrial OS=Mus musculus OX=10090 GN=Ndufs7 PE=1 SV=1 |
| RS13_MOUSE | 40S ribosomal protein S13 OS=Mus musculus OX=10090 GN=Rps13 PE=1 SV=2 |
| NEB2_MOUSE | Neurabin-2 OS=Mus musculus OX=10090 GN=Ppp1r9b PE=1 SV=1 |
| RLA1_MOUSE | 60S acidic ribosomal protein P1 OS=Mus musculus OX=10090 GN=Rplp1 PE=1 SV=1 |
| NEUL_MOUSE | Neurolysin, mitochondrial OS=Mus musculus OX=10090 GN=Nln PE=1 SV=1 |
| SEPT3_MOUSE | Neuronal-specific septin-3 OS=Mus musculus OX=10090 GN=Septin3 PE=1 SV=2 |
| SCAM1_MOUSE | Secretory carrier-associated membrane protein 1 OS=Mus musculus OX=10090 GN=Scamp1 PE=1 SV=1 |
| NCPR_MOUSE | NADPH--cytochrome P450 reductase OS=Mus musculus OX=10090 GN=Por PE=1 SV=2 |
| CO4B_MOUSE | Complement C4-B OS=Mus musculus OX=10090 GN=C4b PE=1 SV=3 |
| DNJA2_MOUSE | DnaJ homolog subfamily A member 2 OS=Mus musculus OX=10090 GN=Dnaja2 PE=1 SV=1 |
| ADK_MOUSE | Adenosine kinase OS=Mus musculus OX=10090 GN=Adk PE=1 SV=2 |
| TP53B_MOUSE | TP53-binding protein 1 OS=Mus musculus OX=10090 GN=Tp53bp1 PE=1 SV=3 |

| | |
|---|---|
| CLIP2_MOUSE | CAP-Gly domain-containing linker protein 2 OS=Mus musculus OX=10090 GN=Clip2 PE=1 SV=2 |
| AGK_MOUSE | Acylglycerol kinase, mitochondrial OS=Mus musculus OX=10090 GN=Agk PE=1 SV=1 |
| SYMC_MOUSE | Methionine--tRNA ligase, cytoplasmic OS=Mus musculus OX=10090 GN=Mars PE=1 SV=1 |
| SGPL1_MOUSE | Sphingosine-1-phosphate lyase 1 OS=Mus musculus OX=10090 GN=Sgpl1 PE=1 SV=1 |
| ANXA7_MOUSE | Annexin A7 OS=Mus musculus OX=10090 GN=Anxa7 PE=1 SV=2 |
| CNBP_MOUSE | Cellular nucleic acid-binding protein OS=Mus musculus OX=10090 GN=Cnbp PE=1 SV=2 |
| GSTM5_MOUSE | Glutathione S-transferase Mu 5 OS=Mus musculus OX=10090 GN=Gstm5 PE=1 SV=1 |
| RL27A_MOUSE | 60S ribosomal protein L27a OS=Mus musculus OX=10090 GN=Rpl27a PE=1 SV=5 |
| SH3G2_MOUSE | Endophilin-A1 OS=Mus musculus OX=10090 GN=Sh3gl2 PE=1 SV=2 |
| NEUA_MOUSE | N-acylneuraminate cytidylyltransferase OS=Mus musculus OX=10090 GN=Cmas PE=1 SV=2 |
| AAK1_MOUSE | AP2-associated protein kinase 1 OS=Mus musculus OX=10090 GN=Aak1 PE=1 SV=2 |
| MGST3_MOUSE | Microsomal glutathione S-transferase 3 OS=Mus musculus OX=10090 GN=Mgst3 PE=1 SV=1 |
| SC31A_MOUSE | Protein transport protein Sec31A OS=Mus musculus OX=10090 GN=Sec31a PE=1 SV=2 |
| NDUBA_MOUSE | NADH dehydrogenase [ubiquinone] 1 beta subcomplex subunit 10 OS=Mus musculus OX=10090 GN=Ndufb10 PE=1 SV=3 |
| NEDD4_MOUSE | E3 ubiquitin-protein ligase NEDD4 OS=Mus musculus OX=10090 GN=Nedd4 PE=1 SV=3 |
| ASGL1_MOUSE | Isoaspartyl peptidase/L-asparaginase OS=Mus musculus OX=10090 GN=Asrgl1 PE=1 SV=1 |
| RTN3_MOUSE | Reticulon-3 OS=Mus musculus OX=10090 GN=Rtn3 PE=1 SV=2 |
| DHRS1_MOUSE | Dehydrogenase/reductase SDR family member 1 OS=Mus musculus OX=10090 GN=Dhrs1 PE=1 SV=1 |
| NP1L1_MOUSE | Nucleosome assembly protein 1-like 1 OS=Mus musculus OX=10090 GN=Nap1l1 PE=1 SV=2 |

| | |
|---|---|
| ABD12_MOUSE | Lysophosphatidylserine lipase ABHD12 OS=Mus musculus OX=10090 GN=Abhd12 PE=1 SV=2 |
| SMRC2_MOUSE | SWI/SNF complex subunit SMARCC2 OS=Mus musculus OX=10090 GN=Smarcc2 PE=1 SV=2 |
| ETFD_MOUSE | Electron transfer flavoprotein-ubiquinone oxidoreductase, mitochondrial OS=Mus musculus OX=10090 GN=Etfdh PE=1 SV=1 |
| PALM_MOUSE | Paralemmin-1 OS=Mus musculus OX=10090 GN=Palm PE=1 SV=1 |
| LANC2_MOUSE | LanC-like protein 2 OS=Mus musculus OX=10090 GN=Lancl2 PE=1 SV=1 |
| CLCB_MOUSE | Clathrin light chain B OS=Mus musculus OX=10090 GN=Cltb PE=1 SV=1 |
| COX5A_MOUSE | Cytochrome c oxidase subunit 5A, mitochondrial OS=Mus musculus OX=10090 GN=Cox5a PE=1 SV=2 |
| GLU2B_MOUSE | Glucosidase 2 subunit beta OS=Mus musculus OX=10090 GN=Prkcsh PE=1 SV=1 |
| LAP2B_MOUSE | Lamina-associated polypeptide 2, isoforms beta/delta/epsilon/gamma OS=Mus musculus OX=10090 GN=Tmpo PE=1 SV=4 |
| NIPS1_MOUSE | Protein NipSnap homolog 1 OS=Mus musculus OX=10090 GN=Nipsnap1 PE=1 SV=1 |
| PLPP_MOUSE | Pyridoxal phosphate phosphatase OS=Mus musculus OX=10090 GN=Pdxp PE=1 SV=1 |
| PSB4_MOUSE | Proteasome subunit beta type-4 OS=Mus musculus OX=10090 GN=Psmb4 PE=1 SV=1 |
| RAB18_MOUSE | Ras-related protein Rab-18 OS=Mus musculus OX=10090 GN=Rab18 PE=1 SV=2 |
| RL34_MOUSE | 60S ribosomal protein L34 OS=Mus musculus OX=10090 GN=Rpl34 PE=1 SV=2 |
| RS9_MOUSE | 40S ribosomal protein S9 OS=Mus musculus OX=10090 GN=Rps9 PE=1 SV=3 |
| SP16H_MOUSE | FACT complex subunit SPT16 OS=Mus musculus OX=10090 GN=Supt16h PE=1 SV=2 |
| LY6H_MOUSE | Lymphocyte antigen 6H OS=Mus musculus OX=10090 GN=Ly6h PE=1 SV=2 |
| SC22B_MOUSE | Vesicle-trafficking protein SEC22b OS=Mus musculus OX=10090 GN=Sec22b PE=1 SV=3 |
| AP1B1_MOUSE | AP-1 complex subunit beta-1 OS=Mus musculus OX=10090 GN=Ap1b1 PE=1 SV=2 |

| | |
|---|---|
| 6PGL_MOUSE | 6-phosphogluconolactonase OS=Mus musculus OX=10090 GN=Pgls PE=1 SV=1 |
| CENPV_MOUSE | Centromere protein V OS=Mus musculus OX=10090 GN=Cenpv PE=1 SV=2 |
| SNAA_MOUSE | Alpha-soluble NSF attachment protein OS=Mus musculus OX=10090 GN=Napa PE=1 SV=1 |
| AHSA1_MOUSE | Activator of 90 kDa heat shock protein ATPase homolog 1 OS=Mus musculus OX=10090 GN=Ahsa1 PE=1 SV=2 |
| GLNA_MOUSE | Glutamine synthetase OS=Mus musculus OX=10090 GN=Glul PE=1 SV=6 |
| TMED2_MOUSE | Transmembrane emp24 domain-containing protein 2 OS=Mus musculus OX=10090 GN=Tmed2 PE=1 SV=1 |
| GLRX3_MOUSE | Glutaredoxin-3 OS=Mus musculus OX=10090 GN=Glrx3 PE=1 SV=1 |
| PA1B2_MOUSE | Platelet-activating factor acetylhydrolase IB subunit beta OS=Mus musculus OX=10090 GN=Pafah1b2 PE=1 SV=2 |
| CLPT1_MOUSE | Cleft lip and palate transmembrane protein 1 homolog OS=Mus musculus OX=10090 GN=Clptm1 PE=1 SV=1 |
| E41L2_MOUSE | Band 4.1-like protein 2 OS=Mus musculus OX=10090 GN=Epb41l2 PE=1 SV=2 |
| HSDL1_MOUSE | Inactive hydroxysteroid dehydrogenase-like protein 1 OS=Mus musculus OX=10090 GN=Hsdl1 PE=1 SV=1 |
| NCLN_MOUSE | Nicalin OS=Mus musculus OX=10090 GN=Ncln PE=1 SV=2 |
| NRX1A_MOUSE | Neurexin-1 OS=Mus musculus OX=10090 GN=Nrxn1 PE=1 SV=3 |
| ROA0_MOUSE | Heterogeneous nuclear ribonucleoprotein A0 OS=Mus musculus OX=10090 GN=Hnrnpa0 PE=1 SV=1 |
| MPP2_MOUSE | MAGUK p55 subfamily member 2 OS=Mus musculus OX=10090 GN=Mpp2 PE=1 SV=1 |
| SRSF2_MOUSE | Serine/arginine-rich splicing factor 2 OS=Mus musculus OX=10090 GN=Srsf2 PE=1 SV=4 |
| NT5D3_MOUSE | 5'-nucleotidase domain-containing protein 3 OS=Mus musculus OX=10090 GN=Nt5dc3 PE=1 SV=1 |
| PYR1_MOUSE | CAD protein OS=Mus musculus OX=10090 GN=Cad PE=1 SV=1 |

| | |
|---|---|
| A2ASS6-DECOY | A2ASS6 |
| BASI_MOUSE | Basigin OS=Mus musculus OX=10090 GN=Bsg PE=1 SV=2 |
| EIF3D_MOUSE | Eukaryotic translation initiation factor 3 subunit D OS=Mus musculus OX=10090 GN=Eif3d PE=1 SV=2 |
| FA49B_MOUSE | Protein FAM49B OS=Mus musculus OX=10090 GN=Fam49b PE=1 SV=1 |
| GCYB1_MOUSE | Guanylate cyclase soluble subunit beta-1 OS=Mus musculus OX=10090 GN=Gucy1b1 PE=1 SV=1 |
| PPR1B_MOUSE | Protein phosphatase 1 regulatory subunit 1B OS=Mus musculus OX=10090 GN=Ppp1r1b PE=1 SV=2 |
| PSMD7_MOUSE | 26S proteasome non-ATPase regulatory subunit 7 OS=Mus musculus OX=10090 GN=Psmd7 PE=1 SV=2 |
| RL14_MOUSE | 60S ribosomal protein L14 OS=Mus musculus OX=10090 GN=Rpl14 PE=1 SV=3 |
| SYUA_MOUSE | Alpha-synuclein OS=Mus musculus OX=10090 GN=Snca PE=1 SV=2 |
| THIO_MOUSE | Thioredoxin OS=Mus musculus OX=10090 GN=Txn PE=1 SV=3 |
| NUDC_MOUSE | Nuclear migration protein nudC OS=Mus musculus OX=10090 GN=Nudc PE=1 SV=1 |
| ABCD3_MOUSE | ATP-binding cassette sub-family D member 3 OS=Mus musculus OX=10090 GN=Abcd3 PE=1 SV=2 |
| PP1R7_MOUSE | Protein phosphatase 1 regulatory subunit 7 OS=Mus musculus OX=10090 GN=Ppp1r7 PE=1 SV=2 |
| SATT_MOUSE | Neutral amino acid transporter A OS=Mus musculus OX=10090 GN=Slc1a4 PE=1 SV=1 |
| UB2V1_MOUSE | Ubiquitin-conjugating enzyme E2 variant 1 OS=Mus musculus OX=10090 GN=Ube2v1 PE=1 SV=1 |
| PA1B3_MOUSE | Platelet-activating factor acetylhydrolase IB subunit gamma OS=Mus musculus OX=10090 GN=Pafah1b3 PE=1 SV=1 |
| SEPT5_MOUSE | Septin-5 OS=Mus musculus OX=10090 GN=Septin5 PE=1 SV=2 |
| ATAD1_MOUSE | ATPase family AAA domain-containing protein 1 OS=Mus musculus OX=10090 GN=Atad1 PE=1 SV=1 |

| | |
|---|---|
| NLTP_MOUSE | Non-specific lipid-transfer protein OS=Mus musculus OX=10090 GN=Scp2 PE=1 SV=3 |
| HS71A_MOUSE | Heat shock 70 kDa protein 1A OS=Mus musculus OX=10090 GN=Hspa1a PE=1 SV=2 |
| TRFE_MOUSE | Serotransferrin OS=Mus musculus OX=10090 GN=Tf PE=1 SV=1 |
| GEPH_MOUSE | Gephyrin OS=Mus musculus OX=10090 GN=Gphn PE=1 SV=2 |
| SAE2_MOUSE | SUMO-activating enzyme subunit 2 OS=Mus musculus OX=10090 GN=Uba2 PE=1 SV=1 |
| MECP2_MOUSE | Methyl-CpG-binding protein 2 OS=Mus musculus OX=10090 GN=Mecp2 PE=1 SV=1 |
| ACADL_MOUSE | Long-chain specific acyl-CoA dehydrogenase, mitochondrial OS=Mus musculus OX=10090 GN=Acadl PE=1 SV=2 |
| DHPR_MOUSE | Dihydropteridine reductase OS=Mus musculus OX=10090 GN=Qdpr PE=1 SV=2 |
| RL13_MOUSE | 60S ribosomal protein L13 OS=Mus musculus OX=10090 GN=Rpl13 PE=1 SV=3 |
| RL17_MOUSE | 60S ribosomal protein L17 OS=Mus musculus OX=10090 GN=Rpl17 PE=1 SV=3 |
| SRSF7_MOUSE | Serine/arginine-rich splicing factor 7 OS=Mus musculus OX=10090 GN=Srsf7 PE=1 SV=1 |
| TICN3_MOUSE | Testican-3 OS=Mus musculus OX=10090 GN=Spock3 PE=2 SV=2 |
| VAMP2_MOUSE | Vesicle-associated membrane protein 2 OS=Mus musculus OX=10090 GN=Vamp2 PE=1 SV=2 |
| CBX5_MOUSE | Chromobox protein homolog 5 OS=Mus musculus OX=10090 GN=Cbx5 PE=1 SV=1 |
| KV5AB_MOUSE | Ig kappa chain V-V region HP R16.7 OS=Mus musculus OX=10090 PE=1 SV=1 |
| SMCA5_MOUSE | SWI/SNF-related matrix-associated actin-dependent regulator of chromatin subfamily A member 5 OS=Mus musculus OX=10090 GN=Smarca5 PE=1 SV=1 |
| GSK3B_MOUSE | Glycogen synthase kinase-3 beta OS=Mus musculus OX=10090 GN=Gsk3b PE=1 SV=2 |
| RS17_MOUSE | 40S ribosomal protein S17 OS=Mus musculus OX=10090 GN=Rps17 PE=1 SV=2 |
| ATP5L_MOUSE | ATP synthase subunit g, mitochondrial OS=Mus musculus OX=10090 GN=Atp5mg PE=1 SV=1 |

| | |
|---|---|
| CRIP2_MOUSE | Cysteine-rich protein 2 OS=Mus musculus OX=10090 GN=Crip2 PE=1 SV=1 |
| TPM1_MOUSE | Tropomyosin alpha-1 chain OS=Mus musculus OX=10090 GN=Tpm1 PE=1 SV=1 |
| IPO4_MOUSE | Importin-4 OS=Mus musculus OX=10090 GN=Ipo4 PE=1 SV=1 |
| ERP29_MOUSE | Endoplasmic reticulum resident protein 29 OS=Mus musculus OX=10090 GN=Erp29 PE=1 SV=2 |
| GTF2I_MOUSE | General transcription factor II-I OS=Mus musculus OX=10090 GN=Gtf2i PE=1 SV=3 |
| STX7_MOUSE | Syntaxin-7 OS=Mus musculus OX=10090 GN=Stx7 PE=1 SV=3 |
| SRC_MOUSE | Neuronal proto-oncogene tyrosine-protein kinase Src OS=Mus musculus OX=10090 GN=Src PE=1 SV=4 |
| AMPL_MOUSE | Cytosol aminopeptidase OS=Mus musculus OX=10090 GN=Lap3 PE=1 SV=3 |
| XPO5_MOUSE | Exportin-5 OS=Mus musculus OX=10090 GN=Xpo5 PE=1 SV=1 |
| 2A5E_MOUSE | Serine/threonine-protein phosphatase 2A 56 kDa regulatory subunit epsilon isoform OS=Mus musculus OX=10090 GN=Ppp2r5e PE=1 SV=3 |
| NCAN_MOUSE | Neurocan core protein OS=Mus musculus OX=10090 GN=Ncan PE=1 SV=1 |
| TM1L2_MOUSE | TOM1-like protein 2 OS=Mus musculus OX=10090 GN=Tom1l2 PE=1 SV=1 |
| NAA15_MOUSE | N-alpha-acetyltransferase 15, NatA auxiliary subunit OS=Mus musculus OX=10090 GN=Naa15 PE=1 SV=1 |
| DMXL2_MOUSE | DmX-like protein 2 OS=Mus musculus OX=10090 GN=Dmxl2 PE=1 SV=3 |
| PLXB2_MOUSE | Plexin-B2 OS=Mus musculus OX=10090 GN=Plxnb2 PE=1 SV=1 |
| CUL2_MOUSE | Cullin-2 OS=Mus musculus OX=10090 GN=Cul2 PE=1 SV=2 |
| GBG2_MOUSE | Guanine nucleotide-binding protein G(I)/G(S)/G(O) subunit gamma-2 OS=Mus musculus OX=10090 GN=Gng2 PE=1 SV=2 |
| NDUA4_MOUSE | Cytochrome c oxidase subunit NDUFA4 OS=Mus musculus OX=10090 GN=Ndufa4 PE=1 SV=2 |

| | |
|---|---|
| RL13A_MOUSE | 60S ribosomal protein L13a OS=Mus musculus OX=10090 GN=Rpl13a PE=1 SV=4 |
| TMEDA_MOUSE | Transmembrane emp24 domain-containing protein 10 OS=Mus musculus OX=10090 GN=Tmed10 PE=1 SV=1 |
| UBC12_MOUSE | NEDD8-conjugating enzyme Ubc12 OS=Mus musculus OX=10090 GN=Ube2m PE=1 SV=1 |
| CDK5_MOUSE | Cyclin-dependent-like kinase 5 OS=Mus musculus OX=10090 GN=Cdk5 PE=1 SV=1 |
| DNPEP_MOUSE | Aspartyl aminopeptidase OS=Mus musculus OX=10090 GN=Dnpep PE=1 SV=2 |
| ML12B_MOUSE | Myosin regulatory light chain 12B OS=Mus musculus OX=10090 GN=Myl12b PE=1 SV=2 |
| SAFB1_MOUSE | Scaffold attachment factor B1 OS=Mus musculus OX=10090 GN=Safb PE=1 SV=2 |
| CEND_MOUSE | Cell cycle exit and neuronal differentiation protein 1 OS=Mus musculus OX=10090 GN=Cend1 PE=1 SV=1 |
| RL38_MOUSE | 60S ribosomal protein L38 OS=Mus musculus OX=10090 GN=Rpl38 PE=1 SV=3 |
| TAGL2_MOUSE | Transgelin-2 OS=Mus musculus OX=10090 GN=Tagln2 PE=1 SV=4 |
| RL18A_MOUSE | 60S ribosomal protein L18a OS=Mus musculus OX=10090 GN=Rpl18a PE=1 SV=1 |
| AT1B2_MOUSE | Sodium/potassium-transporting ATPase subunit beta-2 OS=Mus musculus OX=10090 GN=Atp1b2 PE=1 SV=2 |
| NDUB9_MOUSE | NADH dehydrogenase [ubiquinone] 1 beta subcomplex subunit 9 OS=Mus musculus OX=10090 GN=Ndufb9 PE=1 SV=3 |
| RL24_MOUSE | 60S ribosomal protein L24 OS=Mus musculus OX=10090 GN=Rpl24 PE=1 SV=2 |
| TMOD2_MOUSE | Tropomodulin-2 OS=Mus musculus OX=10090 GN=Tmod2 PE=1 SV=2 |
| ERGI1_MOUSE | Endoplasmic reticulum-Golgi intermediate compartment protein 1 OS=Mus musculus OX=10090 GN=Ergic1 PE=1 SV=1 |
| PSB3_MOUSE | Proteasome subunit beta type-3 OS=Mus musculus OX=10090 GN=Psmb3 PE=1 SV=1 |
| AACS_MOUSE | Acetoacetyl-CoA synthetase OS=Mus musculus OX=10090 GN=Aacs PE=1 SV=1 |

| | |
|---|---|
| PLAP_MOUSE | Phospholipase A-2-activating protein OS=Mus musculus OX=10090 GN=Plaa PE=1 SV=4 |
| NIPS2_MOUSE | Protein NipSnap homolog 2 OS=Mus musculus OX=10090 GN=Nipsnap2 PE=1 SV=1 |
| KIF1A_MOUSE | Kinesin-like protein KIF1A OS=Mus musculus OX=10090 GN=Kif1a PE=1 SV=2 |
| PHP14_MOUSE | 14 kDa phosphohistidine phosphatase OS=Mus musculus OX=10090 GN=Phpt1 PE=1 SV=1 |
| ITSN1_MOUSE | Intersectin-1 OS=Mus musculus OX=10090 GN=Itsn1 PE=1 SV=2 |
| P5CR2_MOUSE | Pyrroline-5-carboxylate reductase 2 OS=Mus musculus OX=10090 GN=Pycr2 PE=1 SV=1 |
| SRSF6_MOUSE | Serine/arginine-rich splicing factor 6 OS=Mus musculus OX=10090 GN=Srsf6 PE=1 SV=1 |
| ASSY_MOUSE | Argininosuccinate synthase OS=Mus musculus OX=10090 GN=Ass1 PE=1 SV=1 |
| DNJC5_MOUSE | DnaJ homolog subfamily C member 5 OS=Mus musculus OX=10090 GN=Dnajc5 PE=1 SV=1 |
| UGDH_MOUSE | UDP-glucose 6-dehydrogenase OS=Mus musculus OX=10090 GN=Ugdh PE=1 SV=1 |
| RL31_MOUSE | 60S ribosomal protein L31 OS=Mus musculus OX=10090 GN=Rpl31 PE=1 SV=1 |
| PTBP1_MOUSE | Polypyrimidine tract-binding protein 1 OS=Mus musculus OX=10090 GN=Ptbp1 PE=1 SV=2 |
| RRBP1_MOUSE | Ribosome-binding protein 1 OS=Mus musculus OX=10090 GN=Rrbp1 PE=1 SV=2 |
| A4_RAT | Amyloid-beta A4 protein OS=Rattus norvegicus OX=10116 GN=App PE=1 SV=2 |
| RS10_MOUSE | 40S ribosomal protein S10 OS=Mus musculus OX=10090 GN=Rps10 PE=1 SV=1 |
| RS27_MOUSE | 40S ribosomal protein S27 OS=Mus musculus OX=10090 GN=Rps27 PE=1 SV=3 |
| ELOB_MOUSE | Elongin-B OS=Mus musculus OX=10090 GN=Elob PE=1 SV=1 |
| HACD3_MOUSE | Very-long-chain (3R)-3-hydroxyacyl-CoA dehydratase 3 OS=Mus musculus OX=10090 GN=Hacd3 PE=1 SV=2 |
| TPM4_MOUSE | Tropomyosin alpha-4 chain OS=Mus musculus OX=10090 GN=Tpm4 PE=1 SV=3 |
| CATD_MOUSE | Cathepsin D OS=Mus musculus OX=10090 GN=Ctsd PE=1 SV=1 |

| | |
|---|---|
| NASP_MOUSE | Nuclear autoantigenic sperm protein OS=Mus musculus OX=10090 GN=Nasp PE=1 SV=2 |
| LIPA2_MOUSE | Liprin-alpha-2 OS=Mus musculus OX=10090 GN=Ppfia2 PE=1 SV=2 |
| LSAMP_RAT | Limbic system-associated membrane protein OS=Rattus norvegicus OX=10116 GN=Lsamp PE=1 SV=1 |
| CELF2_MOUSE | CUGBP Elav-like family member 2 OS=Mus musculus OX=10090 GN=Celf2 PE=1 SV=1 |
| HDAC2_MOUSE | Histone deacetylase 2 OS=Mus musculus OX=10090 GN=Hdac2 PE=1 SV=1 |
| RL19_MOUSE | 60S ribosomal protein L19 OS=Mus musculus OX=10090 GN=Rpl19 PE=1 SV=1 |
| CYB5B_MOUSE | Cytochrome b5 type B OS=Mus musculus OX=10090 GN=Cyb5b PE=1 SV=1 |
| ANM5_MOUSE | Protein arginine N-methyltransferase 5 OS=Mus musculus OX=10090 GN=Prmt5 PE=1 SV=3 |
| EIF3I_MOUSE | Eukaryotic translation initiation factor 3 subunit I OS=Mus musculus OX=10090 GN=Eif3i PE=1 SV=1 |
| SAE1_MOUSE | SUMO-activating enzyme subunit 1 OS=Mus musculus OX=10090 GN=Sae1 PE=1 SV=1 |
| GRB2_MOUSE | Growth factor receptor-bound protein 2 OS=Mus musculus OX=10090 GN=Grb2 PE=1 SV=1 |
| ELP1_MOUSE | Elongator complex protein 1 OS=Mus musculus OX=10090 GN=Elp1 PE=1 SV=2 |
| ATRX_MOUSE | Transcriptional regulator ATRX OS=Mus musculus OX=10090 GN=Atrx PE=1 SV=3 |
| PP2BB_MOUSE | Serine/threonine-protein phosphatase 2B catalytic subunit beta isoform OS=Mus musculus OX=10090 GN=Ppp3cb PE=1 SV=2 |
| IF4G2_MOUSE | Eukaryotic translation initiation factor 4 gamma 2 OS=Mus musculus OX=10090 GN=Eif4g2 PE=1 SV=2 |
| SCG2_MOUSE | Secretogranin-2 OS=Mus musculus OX=10090 GN=Scg2 PE=1 SV=1 |
| TM9S2_MOUSE | Transmembrane 9 superfamily member 2 OS=Mus musculus OX=10090 GN=Tm9sf2 PE=1 SV=1 |
| EAA2_MOUSE | Excitatory amino acid transporter 2 OS=Mus musculus OX=10090 GN=Slc1a2 PE=1 SV=1 |

| | |
|---|---|
| LAC2_MOUSE | Ig lambda-2 chain C region OS=Mus musculus OX=10090 GN=Iglc2 PE=1 SV=1 |
| MARE2_MOUSE | Microtubule-associated protein RP/EB family member 2 OS=Mus musculus OX=10090 GN=Mapre2 PE=1 SV=1 |
| ACTB_BOVIN | Actin, cytoplasmic 1 OS=Bos taurus OX=9913 GN=ACTB PE=1 SV=1 |
| LRC47_MOUSE | Leucine-rich repeat-containing protein 47 OS=Mus musculus OX=10090 GN=Lrrc47 PE=1 SV=1 |
| RL28_MOUSE | 60S ribosomal protein L28 OS=Mus musculus OX=10090 GN=Rpl28 PE=1 SV=2 |
| TWF1_MOUSE | Twinfilin-1 OS=Mus musculus OX=10090 GN=Twf1 PE=1 SV=2 |
| U2AF1_MOUSE | Splicing factor U2AF 35 kDa subunit OS=Mus musculus OX=10090 GN=U2af1 PE=1 SV=4 |
| NDUV2_MOUSE | NADH dehydrogenase [ubiquinone] flavoprotein 2, mitochondrial OS=Mus musculus OX=10090 GN=Ndufv2 PE=1 SV=2 |
| NCEH1_MOUSE | Neutral cholesterol ester hydrolase 1 OS=Mus musculus OX=10090 GN=Nceh1 PE=1 SV=1 |
| NDUS8_MOUSE | NADH dehydrogenase [ubiquinone] iron-sulfur protein 8, mitochondrial OS=Mus musculus OX=10090 GN=Ndufs8 PE=1 SV=1 |
| SGT1_MOUSE | Protein SGT1 homolog OS=Mus musculus OX=10090 GN=Sugt1 PE=1 SV=3 |
| AIMP2_MOUSE | Aminoacyl tRNA synthase complex-interacting multifunctional protein 2 OS=Mus musculus OX=10090 GN=Aimp2 PE=1 SV=2 |
| DECR_MOUSE | 2,4-dienoyl-CoA reductase, mitochondrial OS=Mus musculus OX=10090 GN=Decr1 PE=1 SV=1 |
| RS15A_MOUSE | 40S ribosomal protein S15a OS=Mus musculus OX=10090 GN=Rps15a PE=1 SV=2 |
| CPNS1_MOUSE | Calpain small subunit 1 OS=Mus musculus OX=10090 GN=Capns1 PE=1 SV=1 |
| SFXN1_MOUSE | Sideroflexin-1 OS=Mus musculus OX=10090 GN=Sfxn1 PE=1 SV=3 |
| CD47_MOUSE | Leukocyte surface antigen CD47 OS=Mus musculus OX=10090 GN=Cd47 PE=1 SV=2 |
| TAGL3_MOUSE | Transgelin-3 OS=Mus musculus OX=10090 GN=Tagln3 PE=1 SV=1 |

| | |
|---|---|
| PCCA_MOUSE | Propionyl-CoA carboxylase alpha chain, mitochondrial OS=Mus musculus OX=10090 GN=Pcca PE=1 SV=2 |
| AOFA_MOUSE | Amine oxidase [flavin-containing] A OS=Mus musculus OX=10090 GN=Maoa PE=1 SV=3 |
| CON__P00735 | CON__P00735 |
| CXAR_MOUSE | Coxsackievirus and adenovirus receptor homolog OS=Mus musculus OX=10090 GN=Cxadr PE=1 SV=1 |
| GDN_MOUSE | Glia-derived nexin OS=Mus musculus OX=10090 GN=Serpine2 PE=1 SV=2 |
| CPSF6_MOUSE | Cleavage and polyadenylation specificity factor subunit 6 OS=Mus musculus OX=10090 GN=Cpsf6 PE=1 SV=1 |
| TMM65_MOUSE | Transmembrane protein 65 OS=Mus musculus OX=10090 GN=Tmem65 PE=1 SV=1 |
| KAP2_MOUSE | cAMP-dependent protein kinase type II-alpha regulatory subunit OS=Mus musculus OX=10090 GN=Prkar2a PE=1 SV=2 |
| MAOX_MOUSE | NADP-dependent malic enzyme OS=Mus musculus OX=10090 GN=Me1 PE=1 SV=2 |
| NRX3A_MOUSE | Neurexin-3 OS=Mus musculus OX=10090 GN=Nrxn3 PE=1 SV=2 |
| LMAN2_MOUSE | Vesicular integral-membrane protein VIP36 OS=Mus musculus OX=10090 GN=Lman2 PE=1 SV=2 |
| SORC2_MOUSE | VPS10 domain-containing receptor SorCS2 OS=Mus musculus OX=10090 GN=Sorcs2 PE=1 SV=2 |
| XPP1_MOUSE | Xaa-Pro aminopeptidase 1 OS=Mus musculus OX=10090 GN=Xpnpep1 PE=1 SV=1 |
| BCAL2_ARATH | Branched-chain-amino-acid aminotransferase-like protein 2 OS=Arabidopsis thaliana OX=3702 GN=At5g27410 PE=2 SV=1 |
| FIS1_MOUSE | Mitochondrial fission 1 protein OS=Mus musculus OX=10090 GN=Fis1 PE=1 SV=1 |
| FKB1A_MOUSE | Peptidyl-prolyl cis-trans isomerase FKBP1A OS=Mus musculus OX=10090 GN=Fkbp1a PE=1 SV=2 |
| NDUA5_MOUSE | NADH dehydrogenase [ubiquinone] 1 alpha subcomplex subunit 5 OS=Mus musculus OX=10090 GN=Ndufa5 PE=1 SV=3 |
| CON__P13647 | CON__P13647 |

| | |
|---|---|
| THOC4_MOUSE | THO complex subunit 4 OS=Mus musculus OX=10090 GN=Alyref PE=1 SV=3 |
| SRP68_MOUSE | Signal recognition particle subunit SRP68 OS=Mus musculus OX=10090 GN=Srp68 PE=1 SV=2 |
| APT_MOUSE | Adenine phosphoribosyltransferase OS=Mus musculus OX=10090 GN=Aprt PE=1 SV=2 |
| CLIC4_MOUSE | Chloride intracellular channel protein 4 OS=Mus musculus OX=10090 GN=Clic4 PE=1 SV=3 |
| CYC_MOUSE | Cytochrome c, somatic OS=Mus musculus OX=10090 GN=Cycs PE=1 SV=2 |
| ACADM_MOUSE | Medium-chain specific acyl-CoA dehydrogenase, mitochondrial OS=Mus musculus OX=10090 GN=Acadm PE=1 SV=1 |
| sp\|Q6ZPJ3\|UBE2O_MOUSE | E2 ubiquitin-conjugating enzyme UBE2O OS=Mus musculus OX=10090 GN=Ube2o PE=1 SV=3 |
| BAF_MOUSE | Barrier-to-autointegration factor OS=Mus musculus OX=10090 GN=Banf1 PE=1 SV=1 |
| ERF1_HUMAN | Eukaryotic peptide chain release factor subunit 1 OS=Homo sapiens OX=9606 GN=ETF1 PE=1 SV=3 |
| RL10A_MOUSE | 60S ribosomal protein L10a OS=Mus musculus OX=10090 GN=Rpl10a PE=1 SV=3 |
| NOVA1_MOUSE | RNA-binding protein Nova-1 OS=Mus musculus OX=10090 GN=Nova1 PE=1 SV=2 |
| TRXR1_MOUSE | Thioredoxin reductase 1, cytoplasmic OS=Mus musculus OX=10090 GN=Txnrd1 PE=1 SV=3 |
| TSP1_MOUSE | Thrombospondin-1 OS=Mus musculus OX=10090 GN=Thbs1 PE=1 SV=1 |
| HIP1_MOUSE | Huntingtin-interacting protein 1 OS=Mus musculus OX=10090 GN=Hip1 PE=1 SV=2 |
| PSDE_MOUSE | 26S proteasome non-ATPase regulatory subunit 14 OS=Mus musculus OX=10090 GN=Psmd14 PE=1 SV=2 |
| PSMD4_MOUSE | 26S proteasome non-ATPase regulatory subunit 4 OS=Mus musculus OX=10090 GN=Psmd4 PE=1 SV=1 |
| DYN2_MOUSE | Dynamin-2 OS=Mus musculus OX=10090 GN=Dnm2 PE=1 SV=2 |
| TMED9_MOUSE | Transmembrane emp24 domain-containing protein 9 OS=Mus musculus OX=10090 GN=Tmed9 PE=1 SV=2 |

| | |
|---|---|
| CSK2B_HUMAN | Casein kinase II subunit beta OS=Homo sapiens OX=9606 GN=CSNK2B PE=1 SV=1 |
| DC1L2_MOUSE | Cytoplasmic dynein 1 light intermediate chain 2 OS=Mus musculus OX=10090 GN=Dync1li2 PE=1 SV=2 |
| GD1L1_MOUSE | Ganglioside-induced differentiation-associated protein 1-like 1 OS=Mus musculus OX=10090 GN=Gdap1l1 PE=1 SV=1 |
| SEPT2_MOUSE | Septin-2 OS=Mus musculus OX=10090 GN=Septin2 PE=1 SV=2 |
| MINK1_HUMAN | Misshapen-like kinase 1 OS=Homo sapiens OX=9606 GN=MINK1 PE=1 SV=2 |
| APMAP_MOUSE | Adipocyte plasma membrane-associated protein OS=Mus musculus OX=10090 GN=Apmap PE=1 SV=1 |
| SMC3_MOUSE | Structural maintenance of chromosomes protein 3 OS=Mus musculus OX=10090 GN=Smc3 PE=1 SV=2 |
| KV3AA_MOUSE | Ig kappa chain V-III region ABPC 22/PC 9245 OS=Mus musculus OX=10090 PE=1 SV=1 |
| AL4A1_MOUSE | Delta-1-pyrroline-5-carboxylate dehydrogenase, mitochondrial OS=Mus musculus OX=10090 GN=Aldh4a1 PE=1 SV=3 |
| DCAKD_MOUSE | Dephospho-CoA kinase domain-containing protein OS=Mus musculus OX=10090 GN=Dcakd PE=1 SV=1 |
| CALB2_MOUSE | Calretinin OS=Mus musculus OX=10090 GN=Calb2 PE=1 SV=3 |
| DYN3_MOUSE | Dynamin-3 OS=Mus musculus OX=10090 GN=Dnm3 PE=1 SV=1 |
| MIRO1_MOUSE | Mitochondrial Rho GTPase 1 OS=Mus musculus OX=10090 GN=Rhot1 PE=1 SV=1 |
| FKBP3_MOUSE | Peptidyl-prolyl cis-trans isomerase FKBP3 OS=Mus musculus OX=10090 GN=Fkbp3 PE=1 SV=2 |
| HPCL4_MOUSE | Hippocalcin-like protein 4 OS=Mus musculus OX=10090 GN=Hpcal4 PE=1 SV=3 |
| IF4G3_MOUSE | Eukaryotic translation initiation factor 4 gamma 3 OS=Mus musculus OX=10090 GN=Eif4g3 PE=1 SV=2 |
| SEPT9_MOUSE | Septin-9 OS=Mus musculus OX=10090 GN=Septin9 PE=1 SV=1 |

| | |
|---|---|
| API5_MOUSE | Apoptosis inhibitor 5 OS=Mus musculus OX=10090 GN=Api5 PE=1 SV=2 |
| TBB2B_MOUSE | Tubulin beta-2B chain OS=Mus musculus OX=10090 GN=Tubb2b PE=1 SV=1 |
| PSMD5_MOUSE | 26S proteasome non-ATPase regulatory subunit 5 OS=Mus musculus OX=10090 GN=Psmd5 PE=1 SV=4 |
| RS14_MOUSE | 40S ribosomal protein S14 OS=Mus musculus OX=10090 GN=Rps14 PE=1 SV=3 |
| EFHD2_MOUSE | EF-hand domain-containing protein D2 OS=Mus musculus OX=10090 GN=Efhd2 PE=1 SV=1 |
| HBB1_MOUSE | Hemoglobin subunit beta-1 OS=Mus musculus OX=10090 GN=Hbb-b1 PE=1 SV=2 |
| ARL1_MOUSE | ADP-ribosylation factor-like protein 1 OS=Mus musculus OX=10090 GN=Arl1 PE=1 SV=1 |
| ANXA4_MOUSE | Annexin A4 OS=Mus musculus OX=10090 GN=Anxa4 PE=1 SV=4 |
| ATPD_MOUSE | ATP synthase subunit delta, mitochondrial OS=Mus musculus OX=10090 GN=Atp5f1d PE=1 SV=1 |
| ITPA_MOUSE | Inosine triphosphate pyrophosphatase OS=Mus musculus OX=10090 GN=Itpa PE=1 SV=2 |
| ARF4_MOUSE | ADP-ribosylation factor 4 OS=Mus musculus OX=10090 GN=Arf4 PE=1 SV=2 |
| DAZP1_MOUSE | DAZ-associated protein 1 OS=Mus musculus OX=10090 GN=Dazap1 PE=1 SV=2 |
| RAB6B_MOUSE | Ras-related protein Rab-6B OS=Mus musculus OX=10090 GN=Rab6b PE=1 SV=1 |
| VATF_MOUSE | V-type proton ATPase subunit F OS=Mus musculus OX=10090 GN=Atp6v1f PE=1 SV=2 |
| PSB6_MOUSE | Proteasome subunit beta type-6 OS=Mus musculus OX=10090 GN=Psmb6 PE=1 SV=3 |
| MIC19_MOUSE | MICOS complex subunit Mic19 OS=Mus musculus OX=10090 GN=Chchd3 PE=1 SV=1 |
| TECR_MOUSE | Very-long-chain enoyl-CoA reductase OS=Mus musculus OX=10090 GN=Tecr PE=1 SV=1 |
| CPNE6_MOUSE | Copine-6 OS=Mus musculus OX=10090 GN=Cpne6 PE=1 SV=1 |
| PCCB_MOUSE | Propionyl-CoA carboxylase beta chain, mitochondrial OS=Mus musculus OX=10090 GN=Pccb PE=1 SV=2 |

| | |
|---|---|
| KI21A_MOUSE | Kinesin-like protein KIF21A OS=Mus musculus OX=10090 GN=Kif21a PE=1 SV=2 |
| RCN2_MOUSE | Reticulocalbin-2 OS=Mus musculus OX=10090 GN=Rcn2 PE=1 SV=1 |
| SNX1_MOUSE | Sorting nexin-1 OS=Mus musculus OX=10090 GN=Snx1 PE=1 SV=1 |
| CLIC1_MOUSE | Chloride intracellular channel protein 1 OS=Mus musculus OX=10090 GN=Clic1 PE=1 SV=3 |
| CLCA_MOUSE | Clathrin light chain A OS=Mus musculus OX=10090 GN=Clta PE=1 SV=2 |
| PDC6I_MOUSE | Programmed cell death 6-interacting protein OS=Mus musculus OX=10090 GN=Pdcd6ip PE=1 SV=3 |
| SYHC_MOUSE | Histidine--tRNA ligase, cytoplasmic OS=Mus musculus OX=10090 GN=Hars PE=1 SV=2 |
| DCPS_MOUSE | m7GpppX diphosphatase OS=Mus musculus OX=10090 GN=Dcps PE=1 SV=1 |
| SAC1_MOUSE | Phosphatidylinositide phosphatase SAC1 OS=Mus musculus OX=10090 GN=Sacm1l PE=1 SV=1 |
| RBGP1_MOUSE | Rab GTPase-activating protein 1 OS=Mus musculus OX=10090 GN=Rabgap1 PE=1 SV=1 |
| GRIA2_MOUSE | Glutamate receptor 2 OS=Mus musculus OX=10090 GN=Gria2 PE=1 SV=3 |
| CAB39_MOUSE | Calcium-binding protein 39 OS=Mus musculus OX=10090 GN=Cab39 PE=1 SV=2 |
| PSB1_MOUSE | Proteasome subunit beta type-1 OS=Mus musculus OX=10090 GN=Psmb1 PE=1 SV=1 |
| KV3AJ_MOUSE | Ig kappa chain V-III region PC 7175 OS=Mus musculus OX=10090 PE=1 SV=1 |
| NLGN2_MOUSE | Neuroligin-2 OS=Mus musculus OX=10090 GN=Nlgn2 PE=1 SV=2 |
| IF4E_MOUSE | Eukaryotic translation initiation factor 4E OS=Mus musculus OX=10090 GN=Eif4e PE=1 SV=1 |
| SRCN1_MOUSE | SRC kinase signaling inhibitor 1 OS=Mus musculus OX=10090 GN=Srcin1 PE=1 SV=2 |
| ATP5J_MOUSE | ATP synthase-coupling factor 6, mitochondrial OS=Mus musculus OX=10090 GN=Atp5pf PE=1 SV=1 |
| CAZA1_MOUSE | F-actin-capping protein subunit alpha-1 OS=Mus musculus OX=10090 GN=Capza1 PE=1 SV=4 |

| | |
|---|---|
| KV5A3_MOUSE | Ig kappa chain V-V region K2 (Fragment) OS=Mus musculus OX=10090 PE=1 SV=1 |
| FA98B_MOUSE | Protein FAM98B OS=Mus musculus OX=10090 GN=Fam98b PE=1 SV=1 |
| PSMD8_MOUSE | 26S proteasome non-ATPase regulatory subunit 8 OS=Mus musculus OX=10090 GN=Psmd8 PE=1 SV=2 |
| RAB1A_MOUSE | Ras-related protein Rab-1A OS=Mus musculus OX=10090 GN=Rab1A PE=1 SV=3 |
| CASP3_MOUSE | Caspase-3 OS=Mus musculus OX=10090 GN=Casp3 PE=1 SV=1 |
| RASN_MOUSE | GTPase NRas OS=Mus musculus OX=10090 GN=Nras PE=1 SV=1 |
| ARPC4_MOUSE | Actin-related protein 2/3 complex subunit 4 OS=Mus musculus OX=10090 GN=Arpc4 PE=1 SV=3 |
| STMN1_MOUSE | Stathmin OS=Mus musculus OX=10090 GN=Stmn1 PE=1 SV=2 |
| CDS2_MOUSE | Phosphatidate cytidylyltransferase 2 OS=Mus musculus OX=10090 GN=Cds2 PE=1 SV=1 |
| H11_MOUSE | Histone H1.1 OS=Mus musculus OX=10090 GN=H1-1 PE=1 SV=2 |
| PAK1_MOUSE | Serine/threonine-protein kinase PAK 1 OS=Mus musculus OX=10090 GN=Pak1 PE=1 SV=1 |
| DYL2_MOUSE | Dynein light chain 2, cytoplasmic OS=Mus musculus OX=10090 GN=Dynll2 PE=1 SV=1 |
| KTN1_MOUSE | Kinectin OS=Mus musculus OX=10090 GN=Ktn1 PE=1 SV=1 |
| SCRB2_MOUSE | Lysosome membrane protein 2 OS=Mus musculus OX=10090 GN=Scarb2 PE=1 SV=3 |
| CSN3_MOUSE | COP9 signalosome complex subunit 3 OS=Mus musculus OX=10090 GN=Cops3 PE=1 SV=3 |
| EWS_MOUSE | RNA-binding protein EWS OS=Mus musculus OX=10090 GN=Ewsr1 PE=1 SV=2 |
| TXND5_MOUSE | Thioredoxin domain-containing protein 5 OS=Mus musculus OX=10090 GN=Txndc5 PE=1 SV=2 |
| SAR1B_MOUSE | GTP-binding protein SAR1b OS=Mus musculus OX=10090 GN=Sar1b PE=1 SV=1 |
| EZRI_MOUSE | Ezrin OS=Mus musculus OX=10090 GN=Ezr PE=1 SV=3 |

| | |
|---|---|
| MARE3_MOUSE | Microtubule-associated protein RP/EB family member 3 OS=Mus musculus OX=10090 GN=Mapre3 PE=1 SV=1 |
| GBG4_MOUSE | Guanine nucleotide-binding protein G(I)/G(S)/G(O) subunit gamma-4 OS=Mus musculus OX=10090 GN=Gng4 PE=1 SV=1 |
| EP15R_MOUSE | Epidermal growth factor receptor substrate 15-like 1 OS=Mus musculus OX=10090 GN=Eps15l1 PE=1 SV=3 |
| IF4B_MOUSE | Eukaryotic translation initiation factor 4B OS=Mus musculus OX=10090 GN=Eif4b PE=1 SV=1 |
| PCYXL_MOUSE | Prenylcysteine oxidase-like OS=Mus musculus OX=10090 GN=Pcyox1l PE=1 SV=1 |
| CP51A_MOUSE | Lanosterol 14-alpha demethylase OS=Mus musculus OX=10090 GN=Cyp51a1 PE=1 SV=1 |
| MCCB_MOUSE | Methylcrotonoyl-CoA carboxylase beta chain, mitochondrial OS=Mus musculus OX=10090 GN=Mccc2 PE=1 SV=1 |
| NDKB_MOUSE | Nucleoside diphosphate kinase B OS=Mus musculus OX=10090 GN=Nme2 PE=1 SV=1 |
| ARLY_MOUSE | Argininosuccinate lyase OS=Mus musculus OX=10090 GN=Asl PE=1 SV=1 |
| RB6I2_MOUSE | ELKS/Rab6-interacting/CAST family member 1 OS=Mus musculus OX=10090 GN=Erc1 PE=1 SV=1 |
| RAB35_MOUSE | Ras-related protein Rab-35 OS=Mus musculus OX=10090 GN=Rab35 PE=1 SV=1 |
| ATAT_MOUSE | Alpha-tubulin N-acetyltransferase 1 OS=Mus musculus OX=10090 GN=Atat1 PE=1 SV=1 |
| STT3B_MOUSE | Dolichyl-diphosphooligosaccharide--protein glycosyltransferase subunit STT3B OS=Mus musculus OX=10090 GN=Stt3b PE=1 SV=2 |
| SYGP1_MOUSE | Ras/Rap GTPase-activating protein SynGAP OS=Mus musculus OX=10090 GN=Syngap1 PE=1 SV=2 |
| SMC1A_MOUSE | Structural maintenance of chromosomes protein 1A OS=Mus musculus OX=10090 GN=Smc1a PE=1 SV=4 |
| CLAP1_MOUSE | CLIP-associating protein 1 OS=Mus musculus OX=10090 GN=Clasp1 PE=1 SV=2 |

| | |
|---|---|
| SRGP2_MOUSE | SLIT-ROBO Rho GTPase-activating protein 2 OS=Mus musculus OX=10090 GN=Srgap2 PE=1 SV=2 |
| ZRAB2_MOUSE | Zinc finger Ran-binding domain-containing protein 2 OS=Mus musculus OX=10090 GN=Zranb2 PE=1 SV=2 |
| NDUA8_MOUSE | NADH dehydrogenase [ubiquinone] 1 alpha subcomplex subunit 8 OS=Mus musculus OX=10090 GN=Ndufa8 PE=1 SV=3 |
| GPD1L_MOUSE | Glycerol-3-phosphate dehydrogenase 1-like protein OS=Mus musculus OX=10090 GN=Gpd1l PE=1 SV=2 |
| CADM2_MOUSE | Cell adhesion molecule 2 OS=Mus musculus OX=10090 GN=Cadm2 PE=1 SV=2 |
| RS15_CHICK | 40S ribosomal protein S15 OS=Gallus gallus OX=9031 GN=RPS15 PE=2 SV=2 |
| F120A_MOUSE | Constitutive coactivator of PPAR-gamma-like protein 1 OS=Mus musculus OX=10090 GN=FAM120A PE=1 SV=2 |
| TRIM2_MOUSE | Tripartite motif-containing protein 2 OS=Mus musculus OX=10090 GN=Trim2 PE=1 SV=1 |
| SSRP1_MOUSE | FACT complex subunit SSRP1 OS=Mus musculus OX=10090 GN=Ssrp1 PE=1 SV=2 |
| SYQ_MOUSE | Glutamine--tRNA ligase OS=Mus musculus OX=10090 GN=Qars PE=1 SV=1 |
| NDRG4_MOUSE | Protein NDRG4 OS=Mus musculus OX=10090 GN=Ndrg4 PE=1 SV=1 |
| Q3E960-DECOY | Q3E960 |
| RL22_MOUSE | 60S ribosomal protein L22 OS=Mus musculus OX=10090 GN=Rpl22 PE=1 SV=2 |
| PP1A_MOUSE | Serine/threonine-protein phosphatase PP1-alpha catalytic subunit OS=Mus musculus OX=10090 GN=Ppp1ca PE=1 SV=1 |
| CISD1_MOUSE | CDGSH iron-sulfur domain-containing protein 1 OS=Mus musculus OX=10090 GN=Cisd1 PE=1 SV=1 |
| NRX2A_MOUSE | Neurexin-2 OS=Mus musculus OX=10090 GN=Nrxn2 PE=1 SV=1 |
| HVM32_MOUSE | Ig heavy chain V-III region J606 OS=Mus musculus OX=10090 PE=1 SV=1 |
| CX7A2_MOUSE | Cytochrome c oxidase subunit 7A2, mitochondrial OS=Mus musculus OX=10090 GN=Cox7a2 PE=1 SV=2 |

| | |
| --- | --- |
| GNA11_MOUSE | Guanine nucleotide-binding protein subunit alpha-11 OS=Mus musculus OX=10090 GN=Gna11 PE=1 SV=1 |
| CATB_MOUSE | Cathepsin B OS=Mus musculus OX=10090 GN=Ctsb PE=1 SV=2 |
| KCD12_MOUSE | BTB/POZ domain-containing protein KCTD12 OS=Mus musculus OX=10090 GN=Kctd12 PE=1 SV=1 |
| OX2G_MOUSE | OX-2 membrane glycoprotein OS=Mus musculus OX=10090 GN=Cd200 PE=1 SV=1 |
| BIN1_MOUSE | Myc box-dependent-interacting protein 1 OS=Mus musculus OX=10090 GN=Bin1 PE=1 SV=1 |
| PPME1_MOUSE | Protein phosphatase methylesterase 1 OS=Mus musculus OX=10090 GN=Ppme1 PE=1 SV=5 |
| PURA_MOUSE | Transcriptional activator protein Pur-alpha OS=Mus musculus OX=10090 GN=Pura PE=1 SV=1 |
| VPS29_MOUSE | Vacuolar protein sorting-associated protein 29 OS=Mus musculus OX=10090 GN=Vps29 PE=1 SV=1 |
| BUB3_MOUSE | Mitotic checkpoint protein BUB3 OS=Mus musculus OX=10090 GN=Bub3 PE=1 SV=2 |
| NECA2_MOUSE | N-terminal EF-hand calcium-binding protein 2 OS=Mus musculus OX=10090 GN=Necab2 PE=1 SV=1 |
| DC1I2_MOUSE | Cytoplasmic dynein 1 intermediate chain 2 OS=Mus musculus OX=10090 GN=Dync1i2 PE=1 SV=1 |
| HDGF_MOUSE | Hepatoma-derived growth factor OS=Mus musculus OX=10090 GN=Hdgf PE=1 SV=2 |
| CPSF5_MOUSE | Cleavage and polyadenylation specificity factor subunit 5 OS=Mus musculus OX=10090 GN=Nudt21 PE=1 SV=1 |
| CDC37_MOUSE | Hsp90 co-chaperone Cdc37 OS=Mus musculus OX=10090 GN=Cdc37 PE=1 SV=1 |
| DDAH2_MOUSE | N(G),N(G)-dimethylarginine dimethylaminohydrolase 2 OS=Mus musculus OX=10090 GN=Ddah2 PE=1 SV=1 |
| SRRT_MOUSE | Serrate RNA effector molecule homolog OS=Mus musculus OX=10090 GN=Srrt PE=1 SV=1 |
| CTND2_MOUSE | Catenin delta-2 OS=Mus musculus OX=10090 GN=Ctnnd2 PE=1 SV=1 |

| | |
|---|---|
| HVM36_MOUSE | Ig heavy chain V region 441 OS=Mus musculus OX=10090 PE=4 SV=1 |
| SYTC_MOUSE | Threonine--tRNA ligase 1, cytoplasmic OS=Mus musculus OX=10090 GN=Tars1 PE=1 SV=2 |
| SNG3_MOUSE | Synaptogyrin-3 OS=Mus musculus OX=10090 GN=Syngr3 PE=1 SV=1 |
| RL27_MOUSE | 60S ribosomal protein L27 OS=Mus musculus OX=10090 GN=Rpl27 PE=1 SV=2 |
| TCAL5_MOUSE | Transcription elongation factor A protein-like 5 OS=Mus musculus OX=10090 GN=Tceal5 PE=1 SV=1 |
| PTMA_MOUSE | Prothymosin alpha OS=Mus musculus OX=10090 GN=Ptma PE=1 SV=2 |
| RS20_MOUSE | 40S ribosomal protein S20 OS=Mus musculus OX=10090 GN=Rps20 PE=1 SV=1 |
| NOLC1_MOUSE | Nucleolar and coiled-body phosphoprotein 1 OS=Mus musculus OX=10090 GN=Nolc1 PE=1 SV=1 |
| GDPD1_MOUSE | Lysophospholipase D GDPD1 OS=Mus musculus OX=10090 GN=Gdpd1 PE=1 SV=1 |
| TBA4A_MOUSE | Tubulin alpha-4A chain OS=Mus musculus OX=10090 GN=Tuba4a PE=1 SV=1 |
| RHOB_MOUSE | Rho-related GTP-binding protein RhoB OS=Mus musculus OX=10090 GN=Rhob PE=1 SV=1 |
| SEPT6_MOUSE | Septin-6 OS=Mus musculus OX=10090 GN=Septin6 PE=1 SV=4 |
| NMT2_MOUSE | Glycylpeptide N-tetradecanoyltransferase 2 OS=Mus musculus OX=10090 GN=Nmt2 PE=1 SV=1 |
| SF3A1_MOUSE | Splicing factor 3A subunit 1 OS=Mus musculus OX=10090 GN=Sf3a1 PE=1 SV=1 |
| KIF3A_MOUSE | Kinesin-like protein KIF3A OS=Mus musculus OX=10090 GN=Kif3a PE=1 SV=2 |
| ATAD3_MOUSE | ATPase family AAA domain-containing protein 3 OS=Mus musculus OX=10090 GN=Atad3 PE=1 SV=1 |
| BT3L4_MOUSE | Transcription factor BTF3 homolog 4 OS=Mus musculus OX=10090 GN=Btf3l4 PE=1 SV=1 |
| NDUB6_MOUSE | NADH dehydrogenase [ubiquinone] 1 beta subcomplex subunit 6 OS=Mus musculus OX=10090 GN=Ndufb6 PE=1 SV=3 |

| | |
|---|---|
| TMM43_MOUSE | Transmembrane protein 43 OS=Mus musculus OX=10090 GN=Tmem43 PE=1 SV=1 |
| UBE3A_MOUSE | Ubiquitin-protein ligase E3A OS=Mus musculus OX=10090 GN=Ube3a PE=1 SV=2 |
| IF4H_MOUSE | Eukaryotic translation initiation factor 4H OS=Mus musculus OX=10090 GN=Eif4h PE=1 SV=3 |
| CSKP_MOUSE | Peripheral plasma membrane protein CASK OS=Mus musculus OX=10090 GN=Cask PE=1 SV=2 |
| RINI_MOUSE | Ribonuclease inhibitor OS=Mus musculus OX=10090 GN=Rnh1 PE=1 SV=1 |
| TOM40_MOUSE | Mitochondrial import receptor subunit TOM40 homolog OS=Mus musculus OX=10090 GN=Tomm40 PE=1 SV=3 |
| OGFR_MOUSE | Opioid growth factor receptor OS=Mus musculus OX=10090 GN=Ogfr PE=1 SV=1 |
| ANFY1_MOUSE | Rabankyrin-5 OS=Mus musculus OX=10090 GN=Ankfy1 PE=1 SV=2 |
| ABCE1_MOUSE | ATP-binding cassette sub-family E member 1 OS=Mus musculus OX=10090 GN=Abce1 PE=1 SV=1 |
| ATLA1_MOUSE | Atlastin-1 OS=Mus musculus OX=10090 GN=Atl1 PE=1 SV=1 |
| PPM1E_MOUSE | Protein phosphatase 1E OS=Mus musculus OX=10090 GN=Ppm1e PE=1 SV=2 |
| ERH_MOUSE | Enhancer of rudimentary homolog OS=Mus musculus OX=10090 GN=Erh PE=1 SV=1 |
| UBE2K_MOUSE | Ubiquitin-conjugating enzyme E2 K OS=Mus musculus OX=10090 GN=Ube2k PE=1 SV=3 |
| SYPH_MOUSE | Synaptophysin OS=Mus musculus OX=10090 GN=Syp PE=1 SV=2 |
| TCEA1_MOUSE | Transcription elongation factor A protein 1 OS=Mus musculus OX=10090 GN=Tcea1 PE=1 SV=2 |
| CC50A_MOUSE | Cell cycle control protein 50A OS=Mus musculus OX=10090 GN=Tmem30a PE=1 SV=1 |
| SMD1_MOUSE | Small nuclear ribonucleoprotein Sm D1 OS=Mus musculus OX=10090 GN=Snrpd1 PE=1 SV=1 |
| TMX2_MOUSE | Thioredoxin-related transmembrane protein 2 OS=Mus musculus OX=10090 GN=Tmx2 PE=1 SV=1 |

| | |
|---|---|
| RS23_MOUSE | 40S ribosomal protein S23 OS=Mus musculus OX=10090 GN=Rps23 PE=1 SV=3 |
| LGUL_MOUSE | Lactoylglutathione lyase OS=Mus musculus OX=10090 GN=Glo1 PE=1 SV=3 |
| ACBG1_MOUSE | Long-chain-fatty-acid--CoA ligase ACSBG1 OS=Mus musculus OX=10090 GN=Acsbg1 PE=1 SV=1 |
| ZN512_MOUSE | Zinc finger protein 512 OS=Mus musculus OX=10090 GN=Znf512 PE=2 SV=2 |
| CNRP1_MOUSE | CB1 cannabinoid receptor-interacting protein 1 OS=Mus musculus OX=10090 GN=Cnrip1 PE=1 SV=1 |
| NCAM2_MOUSE | Neural cell adhesion molecule 2 OS=Mus musculus OX=10090 GN=Ncam2 PE=1 SV=1 |
| LC7L2_MOUSE | Putative RNA-binding protein Luc7-like 2 OS=Mus musculus OX=10090 GN=Luc7l2 PE=1 SV=1 |
| CSN6_MOUSE | COP9 signalosome complex subunit 6 OS=Mus musculus OX=10090 GN=Cops6 PE=1 SV=1 |
| NECP1_MOUSE | Adaptin ear-binding coat-associated protein 1 OS=Mus musculus OX=10090 GN=Necap1 PE=1 SV=2 |
| DLG3_MOUSE | Disks large homolog 3 OS=Mus musculus OX=10090 GN=Dlg3 PE=1 SV=1 |
| TOP2A_MOUSE | DNA topoisomerase 2-alpha OS=Mus musculus OX=10090 GN=Top2a PE=1 SV=2 |
| RBM14_MOUSE | RNA-binding protein 14 OS=Mus musculus OX=10090 GN=Rbm14 PE=1 SV=1 |
| TPR_MOUSE | Nucleoprotein TPR OS=Mus musculus OX=10090 GN=Tpr PE=1 SV=1 |
| OGT1_MOUSE | UDP-N-acetylglucosamine--peptide N-acetylglucosaminyltransferase 110 kDa subunit OS=Mus musculus OX=10090 GN=Ogt PE=1 SV=2 |
| GCYA1_MOUSE | Guanylate cyclase soluble subunit alpha-1 OS=Mus musculus OX=10090 GN=Gucy1a1 PE=1 SV=2 |
| AL1L2_MOUSE | Mitochondrial 10-formyltetrahydrofolate dehydrogenase OS=Mus musculus OX=10090 GN=Aldh1l2 PE=1 SV=2 |
| LYRIC_MOUSE | Protein LYRIC OS=Mus musculus OX=10090 GN=Mtdh PE=1 SV=1 |

| | |
|---|---|
| 3HIDH_MOUSE | 3-hydroxyisobutyrate dehydrogenase, mitochondrial OS=Mus musculus OX=10090 GN=Hibadh PE=1 SV=1 |
| AL7A1_MOUSE | Alpha-aminoadipic semialdehyde dehydrogenase OS=Mus musculus OX=10090 GN=Aldh7a1 PE=1 SV=4 |
| ACOT1_MOUSE | Acyl-coenzyme A thioesterase 1 OS=Mus musculus OX=10090 GN=Acot1 PE=1 SV=1 |
| NUCB1_MOUSE | Nucleobindin-1 OS=Mus musculus OX=10090 GN=Nucb1 PE=1 SV=2 |
| NUP93_MOUSE | Nuclear pore complex protein Nup93 OS=Mus musculus OX=10090 GN=Nup93 PE=1 SV=1 |
| SYT5_MOUSE | Synaptotagmin-5 OS=Mus musculus OX=10090 GN=Syt5 PE=1 SV=1 |
| TIM44_MOUSE | Mitochondrial import inner membrane translocase subunit TIM44 OS=Mus musculus OX=10090 GN=Timm44 PE=1 SV=2 |
| BZW2_MOUSE | Basic leucine zipper and W2 domain-containing protein 2 OS=Mus musculus OX=10090 GN=Bzw2 PE=1 SV=1 |
| CPLX2_MOUSE | Complexin-2 OS=Mus musculus OX=10090 GN=Cplx2 PE=1 SV=1 |
| SARM1_MOUSE | Sterile alpha and TIR motif-containing protein 1 OS=Mus musculus OX=10090 GN=Sarm1 PE=1 SV=1 |
| THY1_MOUSE | Thy-1 membrane glycoprotein OS=Mus musculus OX=10090 GN=Thy1 PE=1 SV=1 |
| MTX2_MOUSE | Metaxin-2 OS=Mus musculus OX=10090 GN=Mtx2 PE=1 SV=1 |
| CON__P81644 | CON__P81644 |
| HVM57_MOUSE | Ig heavy chain V region 6.96 OS=Mus musculus OX=10090 PE=4 SV=1 |
| CSRP1_MOUSE | Cysteine and glycine-rich protein 1 OS=Mus musculus OX=10090 GN=Csrp1 PE=1 SV=3 |
| PDCD6_MOUSE | Programmed cell death protein 6 OS=Mus musculus OX=10090 GN=Pdcd6 PE=1 SV=2 |
| CSN2_MOUSE | COP9 signalosome complex subunit 2 OS=Mus musculus OX=10090 GN=Cops2 PE=1 SV=1 |
| WASF1_MOUSE | Wiskott-Aldrich syndrome protein family member 1 OS=Mus musculus OX=10090 GN=Wasf1 PE=1 SV=2 |

| | |
|---|---|
| PRAF3_MOUSE | PRA1 family protein 3 OS=Mus musculus OX=10090 GN=Arl6ip5 PE=1 SV=2 |
| RTRAF_MOUSE | RNA transcription, translation and transport factor protein OS=Mus musculus OX=10090 GN=RTRAF PE=1 SV=1 |
| CIRBP_MOUSE | Cold-inducible RNA-binding protein OS=Mus musculus OX=10090 GN=Cirbp PE=1 SV=1 |
| ACTY_MOUSE | Beta-centractin OS=Mus musculus OX=10090 GN=Actr1b PE=1 SV=1 |
| HCDH_MOUSE | Hydroxyacyl-coenzyme A dehydrogenase, mitochondrial OS=Mus musculus OX=10090 GN=Hadh PE=1 SV=2 |
| CHIP_MOUSE | STIP1 homology and U box-containing protein 1 OS=Mus musculus OX=10090 GN=Stub1 PE=1 SV=1 |
| LTOR1_MOUSE | Ragulator complex protein LAMTOR1 OS=Mus musculus OX=10090 GN=Lamtor1 PE=1 SV=1 |
| ARL3_MOUSE | ADP-ribosylation factor-like protein 3 OS=Mus musculus OX=10090 GN=Arl3 PE=1 SV=1 |
| MGN2_MOUSE | Protein mago nashi homolog 2 OS=Mus musculus OX=10090 GN=Magohb PE=2 SV=1 |
| SMD3_MOUSE | Small nuclear ribonucleoprotein Sm D3 OS=Mus musculus OX=10090 GN=Snrpd3 PE=1 SV=1 |
| DCLK2_MOUSE | Serine/threonine-protein kinase DCLK2 OS=Mus musculus OX=10090 GN=Dclk2 PE=1 SV=1 |
| LKHA4_MOUSE | Leukotriene A-4 hydrolase OS=Mus musculus OX=10090 GN=Lta4h PE=1 SV=4 |
| RALA_MOUSE | Ras-related protein Ral-A OS=Mus musculus OX=10090 GN=Rala PE=1 SV=1 |
| AIFM1_MOUSE | Apoptosis-inducing factor 1, mitochondrial OS=Mus musculus OX=10090 GN=Aifm1 PE=1 SV=1 |
| KV3AD_MOUSE | Ig kappa chain V-III region PC 7043 OS=Mus musculus OX=10090 PE=1 SV=1 |
| BAG6_MOUSE | Large proline-rich protein BAG6 OS=Mus musculus OX=10090 GN=Bag6 PE=1 SV=1 |
| SRRM2_MOUSE | Serine/arginine repetitive matrix protein 2 OS=Mus musculus OX=10090 GN=Srrm2 PE=1 SV=3 |
| VTNC_MOUSE | Vitronectin OS=Mus musculus OX=10090 GN=Vtn PE=1 SV=2 |

| | |
|---|---|
| PACS1_MOUSE | Phosphofurin acidic cluster sorting protein 1 OS=Mus musculus OX=10090 GN=Pacs1 PE=1 SV=2 |
| ACTBL_MOUSE | Beta-actin-like protein 2 OS=Mus musculus OX=10090 GN=Actbl2 PE=1 SV=1 |
| CYFP1_MOUSE | Cytoplasmic FMR1-interacting protein 1 OS=Mus musculus OX=10090 GN=Cyfip1 PE=1 SV=1 |
| H2A2A_HUMAN | Histone H2A type 2-A OS=Homo sapiens OX=9606 GN=HIST2H2AA3 PE=1 SV=3 |
| HNRL1_MOUSE | Heterogeneous nuclear ribonucleoprotein U-like protein 1 OS=Mus musculus OX=10090 GN=Hnrnpul1 PE=1 SV=1 |
| RHG01_MOUSE | Rho GTPase-activating protein 1 OS=Mus musculus OX=10090 GN=Arhgap1 PE=1 SV=1 |
| FKBP8_MOUSE | Peptidyl-prolyl cis-trans isomerase FKBP8 OS=Mus musculus OX=10090 GN=Fkbp8 PE=1 SV=2 |
| TRA2A_MOUSE | Transformer-2 protein homolog alpha OS=Mus musculus OX=10090 GN=Tra2a PE=1 SV=1 |
| CEMIP_MOUSE | Cell migration-inducing and hyaluronan-binding protein OS=Mus musculus OX=10090 GN=Cemip PE=1 SV=4 |
| PHOCN_MOUSE | MOB-like protein phocein OS=Mus musculus OX=10090 GN=Mob4 PE=1 SV=1 |
| ELOC_MOUSE | Elongin-C OS=Mus musculus OX=10090 GN=Eloc PE=1 SV=1 |
| PFKAL_MOUSE | ATP-dependent 6-phosphofructokinase, liver type OS=Mus musculus OX=10090 GN=Pfkl PE=1 SV=4 |
| RELCH_MOUSE | RAB11-binding protein RELCH OS=Mus musculus OX=10090 GN=Relch PE=1 SV=1 |
| ACOX1_MOUSE | Peroxisomal acyl-coenzyme A oxidase 1 OS=Mus musculus OX=10090 GN=Acox1 PE=1 SV=5 |
| HMGB3_MOUSE | High mobility group protein B3 OS=Mus musculus OX=10090 GN=Hmgb3 PE=1 SV=3 |
| LXN_MOUSE | Latexin OS=Mus musculus OX=10090 GN=Lxn PE=1 SV=2 |
| ARF5_MOUSE | ADP-ribosylation factor 5 OS=Mus musculus OX=10090 GN=Arf5 PE=1 SV=2 |
| GAL3A_MOUSE | Glutamine amidotransferase-like class 1 domain-containing protein 3A, mitochondrial OS=Mus musculus OX=10090 GN=Gatd3a PE=1 SV=1 |

| | |
|---|---|
| TICN1_MOUSE | Testican-1 OS=Mus musculus OX=10090 GN=Spock1 PE=2 SV=2 |
| PCBP3_MOUSE | Poly(rC)-binding protein 3 OS=Mus musculus OX=10090 GN=Pcbp3 PE=1 SV=3 |
| PGRC2_MOUSE | Membrane-associated progesterone receptor component 2 OS=Mus musculus OX=10090 GN=Pgrmc2 PE=1 SV=2 |
| AP2S1_MOUSE | AP-2 complex subunit sigma OS=Mus musculus OX=10090 GN=Ap2s1 PE=1 SV=1 |
| RAP2B_MOUSE | Ras-related protein Rap-2b OS=Mus musculus OX=10090 GN=Rap2b PE=1 SV=1 |
| MIC26_MOUSE | MICOS complex subunit Mic26 OS=Mus musculus OX=10090 GN=Apoo PE=1 SV=2 |
| BPNT1_MOUSE | 3'(2'),5'-bisphosphate nucleotidase 1 OS=Mus musculus OX=10090 GN=Bpnt1 PE=1 SV=2 |
| RU2A_MOUSE | U2 small nuclear ribonucleoprotein A' OS=Mus musculus OX=10090 GN=Snrpa1 PE=1 SV=2 |
| PI42B_MOUSE | Phosphatidylinositol 5-phosphate 4-kinase type-2 beta OS=Mus musculus OX=10090 GN=Pip4k2b PE=1 SV=1 |
| SMD2_MOUSE | Small nuclear ribonucleoprotein Sm D2 OS=Mus musculus OX=10090 GN=Snrpd2 PE=1 SV=1 |
| RBM39_MOUSE | RNA-binding protein 39 OS=Mus musculus OX=10090 GN=Rbm39 PE=1 SV=2 |
| SORCN_MOUSE | Sorcin OS=Mus musculus OX=10090 GN=Sri PE=1 SV=1 |
| SNX2_MOUSE | Sorting nexin-2 OS=Mus musculus OX=10090 GN=Snx2 PE=1 SV=2 |
| AUHM_MOUSE | Methylglutaconyl-CoA hydratase, mitochondrial OS=Mus musculus OX=10090 GN=Auh PE=1 SV=1 |
| SNX27_MOUSE | Sorting nexin-27 OS=Mus musculus OX=10090 GN=Snx27 PE=1 SV=2 |
| MCAT_MOUSE | Mitochondrial carnitine/acylcarnitine carrier protein OS=Mus musculus OX=10090 GN=Slc25a20 PE=1 SV=1 |
| AL1B1_MOUSE | Aldehyde dehydrogenase X, mitochondrial OS=Mus musculus OX=10090 GN=Aldh1b1 PE=1 SV=1 |
| ACSL1_MOUSE | Long-chain-fatty-acid--CoA ligase 1 OS=Mus musculus OX=10090 GN=Acsl1 PE=1 SV=2 |

| | |
|---|---|
| RAGP1_MOUSE | Ran GTPase-activating protein 1 OS=Mus musculus OX=10090 GN=Rangap1 PE=1 SV=2 |
| MK03_MOUSE | Mitogen-activated protein kinase 3 OS=Mus musculus OX=10090 GN=Mapk3 PE=1 SV=5 |
| PPP5_MOUSE | Serine/threonine-protein phosphatase 5 OS=Mus musculus OX=10090 GN=Ppp5c PE=1 SV=3 |
| TLN1_MOUSE | Talin-1 OS=Mus musculus OX=10090 GN=Tln1 PE=1 SV=2 |
| AMRP_MOUSE | Alpha-2-macroglobulin receptor-associated protein OS=Mus musculus OX=10090 GN=Lrpap1 PE=1 SV=1 |
| NFIX_MOUSE | Nuclear factor 1 X-type OS=Mus musculus OX=10090 GN=Nfix PE=1 SV=2 |
| H32_HUMAN | Histone H3.2 OS=Homo sapiens OX=9606 GN=HIST2H3A PE=1 SV=3 |
| UBP7_MOUSE | Ubiquitin carboxyl-terminal hydrolase 7 OS=Mus musculus OX=10090 GN=Usp7 PE=1 SV=1 |
| PTBP2_MOUSE | Polypyrimidine tract-binding protein 2 OS=Mus musculus OX=10090 GN=Ptbp2 PE=1 SV=2 |
| G3BP1_MOUSE | Ras GTPase-activating protein-binding protein 1 OS=Mus musculus OX=10090 GN=G3bp1 PE=1 SV=1 |
| TMX4_MOUSE | Thioredoxin-related transmembrane protein 4 OS=Mus musculus OX=10090 GN=Tmx4 PE=1 SV=2 |
| PRIO_MOUSE | Major prion protein OS=Mus musculus OX=10090 GN=Prnp PE=1 SV=2 |
| CAD13_MOUSE | Cadherin-13 OS=Mus musculus OX=10090 GN=Cdh13 PE=1 SV=2 |
| UCHL3_MOUSE | Ubiquitin carboxyl-terminal hydrolase isozyme L3 OS=Mus musculus OX=10090 GN=Uchl3 PE=1 SV=2 |
| RP3A_MOUSE | Rabphilin-3A OS=Mus musculus OX=10090 GN=Rph3a PE=1 SV=2 |
| AGAP3_MOUSE | Arf-GAP with GTPase, ANK repeat and PH domain-containing protein 3 OS=Mus musculus OX=10090 GN=Agap3 PE=1 SV=1 |
| KV5A1_MOUSE | Ig kappa chain V19-17 OS=Mus musculus OX=10090 GN=Igk-V19-17 PE=1 SV=1 |
| SSRA_MOUSE | Translocon-associated protein subunit alpha OS=Mus musculus OX=10090 GN=Ssr1 PE=1 SV=1 |

| | |
|---|---|
| RAB3B_MOUSE | Ras-related protein Rab-3B OS=Mus musculus OX=10090 GN=Rab3b PE=1 SV=1 |
| KV5A6_MOUSE | Ig kappa chain V-V region L6 (Fragment) OS=Mus musculus OX=10090 PE=4 SV=1 |
| PTPA_MOUSE | Serine/threonine-protein phosphatase 2A activator OS=Mus musculus OX=10090 GN=Ptpa PE=1 SV=1 |
| HDGR3_MOUSE | Hepatoma-derived growth factor-related protein 3 OS=Mus musculus OX=10090 GN=Hdgfl3 PE=1 SV=2 |
| KAPCA_MOUSE | cAMP-dependent protein kinase catalytic subunit alpha OS=Mus musculus OX=10090 GN=Prkaca PE=1 SV=3 |
| SUGP2_MOUSE | SURP and G-patch domain-containing protein 2 OS=Mus musculus OX=10090 GN=Sugp2 PE=1 SV=2 |
| SPCS2_MOUSE | Signal peptidase complex subunit 2 OS=Mus musculus OX=10090 GN=Spcs2 PE=1 SV=1 |
| EMC2_MOUSE | ER membrane protein complex subunit 2 OS=Mus musculus OX=10090 GN=Emc2 PE=1 SV=1 |
| STX12_MOUSE | Syntaxin-12 OS=Mus musculus OX=10090 GN=Stx12 PE=1 SV=1 |
| BAP31_MOUSE | B-cell receptor-associated protein 31 OS=Mus musculus OX=10090 GN=Bcap31 PE=1 SV=4 |
| FBLL1_MOUSE | rRNA/tRNA 2'-O-methyltransferase fibrillarin-like protein 1 OS=Mus musculus OX=10090 GN=Fbll1 PE=1 SV=1 |
| DYH8_HUMAN | Dynein heavy chain 8, axonemal OS=Homo sapiens OX=9606 GN=DNAH8 PE=1 SV=2 |
| S6A11_MOUSE | Sodium- and chloride-dependent GABA transporter 3 OS=Mus musculus OX=10090 GN=Slc6a11 PE=1 SV=2 |
| TOLIP_MOUSE | Toll-interacting protein OS=Mus musculus OX=10090 GN=Tollip PE=1 SV=1 |
| ATP9A_MOUSE | Probable phospholipid-transporting ATPase IIA OS=Mus musculus OX=10090 GN=Atp9a PE=1 SV=3 |
| GBRB3_MOUSE | Gamma-aminobutyric acid receptor subunit beta-3 OS=Mus musculus OX=10090 GN=Gabrb3 PE=1 SV=1 |
| GELS_MOUSE | Gelsolin OS=Mus musculus OX=10090 GN=Gsn PE=1 SV=3 |

| | |
|---|---|
| K0513_MOUSE | Uncharacterized protein KIAA0513 OS=Mus musculus OX=10090 GN=Kiaa0513 PE=1 SV=1 |
| SYCC_MOUSE | Cysteine--tRNA ligase, cytoplasmic OS=Mus musculus OX=10090 GN=Cars PE=1 SV=2 |
| IF1AX_MOUSE | Eukaryotic translation initiation factor 1A, X-chromosomal OS=Mus musculus OX=10090 GN=Eif1ax PE=2 SV=3 |
| KAD2_MOUSE | Adenylate kinase 2, mitochondrial OS=Mus musculus OX=10090 GN=Ak2 PE=1 SV=5 |
| TOM20_MOUSE | Mitochondrial import receptor subunit TOM20 homolog OS=Mus musculus OX=10090 GN=Tomm20 PE=1 SV=1 |
| DNER_MOUSE | Delta and Notch-like epidermal growth factor-related receptor OS=Mus musculus OX=10090 GN=Dner PE=1 SV=1 |
| NDRG2_MOUSE | Protein NDRG2 OS=Mus musculus OX=10090 GN=Ndrg2 PE=1 SV=1 |
| AL9A1_MOUSE | 4-trimethylaminobutyraldehyde dehydrogenase OS=Mus musculus OX=10090 GN=Aldh9a1 PE=1 SV=1 |
| GLO2_MOUSE | Hydroxyacylglutathione hydrolase, mitochondrial OS=Mus musculus OX=10090 GN=Hagh PE=1 SV=2 |
| WDR37_MOUSE | WD repeat-containing protein 37 OS=Mus musculus OX=10090 GN=Wdr37 PE=1 SV=1 |
| EIF3G_MOUSE | Eukaryotic translation initiation factor 3 subunit G OS=Mus musculus OX=10090 GN=Eif3g PE=1 SV=2 |
| GSTA4_MOUSE | Glutathione S-transferase A4 OS=Mus musculus OX=10090 GN=Gsta4 PE=1 SV=3 |
| NUMA1_MOUSE | Nuclear mitotic apparatus protein 1 OS=Mus musculus OX=10090 GN=Numa1 PE=1 SV=1 |
| CELF1_MOUSE | CUGBP Elav-like family member 1 OS=Mus musculus OX=10090 GN=Celf1 PE=1 SV=2 |
| NNRD_MOUSE | ATP-dependent (S)-NAD(P)H-hydrate dehydratase OS=Mus musculus OX=10090 GN=Naxd PE=1 SV=1 |
| HDAC6_MOUSE | Histone deacetylase 6 OS=Mus musculus OX=10090 GN=Hdac6 PE=1 SV=3 |
| SARNP_MOUSE | SAP domain-containing ribonucleoprotein OS=Mus musculus OX=10090 GN=Sarnp PE=1 SV=3 |

| | |
|---|---|
| GPM6A_MOUSE | Neuronal membrane glycoprotein M6-a OS=Mus musculus OX=10090 GN=Gpm6a PE=1 SV=1 |
| MPC2_MOUSE | Mitochondrial pyruvate carrier 2 OS=Mus musculus OX=10090 GN=Mpc2 PE=1 SV=1 |
| RADI_MOUSE | Radixin OS=Mus musculus OX=10090 GN=Rdx PE=1 SV=3 |
| QCR8_MOUSE | Cytochrome b-c1 complex subunit 8 OS=Mus musculus OX=10090 GN=Uqcrq PE=1 SV=3 |
| YKT6_MOUSE | Synaptobrevin homolog YKT6 OS=Mus musculus OX=10090 GN=Ykt6 PE=1 SV=1 |
| KI21B_MOUSE | Kinesin-like protein KIF21B OS=Mus musculus OX=10090 GN=Kif21b PE=1 SV=2 |
| HSP72_MOUSE | Heat shock-related 70 kDa protein 2 OS=Mus musculus OX=10090 GN=Hspa2 PE=1 SV=2 |
| PYGL_RAT | Glycogen phosphorylase, liver form OS=Rattus norvegicus OX=10116 GN=Pygl PE=1 SV=5 |
| SR140_MOUSE | U2 snRNP-associated SURP motif-containing protein OS=Mus musculus OX=10090 GN=U2surp PE=1 SV=3 |
| ERP44_MOUSE | Endoplasmic reticulum resident protein 44 OS=Mus musculus OX=10090 GN=Erp44 PE=1 SV=1 |
| SNX6_MOUSE | Sorting nexin-6 OS=Mus musculus OX=10090 GN=Snx6 PE=1 SV=2 |
| GGT7_MOUSE | Glutathione hydrolase 7 OS=Mus musculus OX=10090 GN=Ggt7 PE=1 SV=2 |
| SOGA3_MOUSE | Protein SOGA3 OS=Mus musculus OX=10090 GN=Soga3 PE=1 SV=2 |
| PIMT_MOUSE | Protein-L-isoaspartate(D-aspartate) O-methyltransferase OS=Mus musculus OX=10090 GN=Pcmt1 PE=1 SV=3 |
| TM35A_MOUSE | Transmembrane protein 35A OS=Mus musculus OX=10090 GN=Tmem35a PE=1 SV=1 |
| ATP5I_MOUSE | ATP synthase subunit e, mitochondrial OS=Mus musculus OX=10090 GN=Atp5me PE=1 SV=2 |
| RS21_MOUSE | 40S ribosomal protein S21 OS=Mus musculus OX=10090 GN=Rps21 PE=1 SV=1 |
| RL36A_MOUSE | 60S ribosomal protein L36a OS=Mus musculus OX=10090 GN=Rpl36a PE=3 SV=2 |
| RTCA_MOUSE | RNA 3'-terminal phosphate cyclase OS=Mus musculus OX=10090 GN=RtcA PE=1 SV=2 |

| | |
|---|---|
| DYLT1_MOUSE | Dynein light chain Tctex-type 1 OS=Mus musculus OX=10090 GN=Dynlt1 PE=1 SV=1 |
| PRKRA_MOUSE | Interferon-inducible double-stranded RNA-dependent protein kinase activator A OS=Mus musculus OX=10090 GN=Prkra PE=1 SV=1 |
| EIF3H_MOUSE | Eukaryotic translation initiation factor 3 subunit H OS=Mus musculus OX=10090 GN=Eif3h PE=1 SV=1 |
| DEK_MOUSE | Protein DEK OS=Mus musculus OX=10090 GN=Dek PE=1 SV=1 |
| RAB31_MOUSE | Ras-related protein Rab-31 OS=Mus musculus OX=10090 GN=Rab31 PE=1 SV=1 |
| ASPH_MOUSE | Aspartyl/asparaginyl beta-hydroxylase OS=Mus musculus OX=10090 GN=Asph PE=1 SV=1 |
| CYB5_MOUSE | Cytochrome b5 OS=Mus musculus OX=10090 GN=Cyb5a PE=1 SV=2 |
| GORS2_MOUSE | Golgi reassembly-stacking protein 2 OS=Mus musculus OX=10090 GN=Gorasp2 PE=1 SV=3 |
| S4A10_MOUSE | Sodium-driven chloride bicarbonate exchanger OS=Mus musculus OX=10090 GN=Slc4a10 PE=1 SV=2 |
| SYIM_MOUSE | Isoleucine--tRNA ligase, mitochondrial OS=Mus musculus OX=10090 GN=Iars2 PE=1 SV=1 |
| RL32_MOUSE | 60S ribosomal protein L32 OS=Mus musculus OX=10090 GN=Rpl32 PE=1 SV=2 |
| AMPB_MOUSE | Aminopeptidase B OS=Mus musculus OX=10090 GN=Rnpep PE=1 SV=2 |
| KVM5_MOUSE | Ig kappa chain V region Mem5 (Fragment) OS=Mus musculus OX=10090 PE=1 SV=1 |
| SC11A_MOUSE | Signal peptidase complex catalytic subunit SEC11A OS=Mus musculus OX=10090 GN=Sec11a PE=1 SV=1 |
| TBA1C_MOUSE | Tubulin alpha-1C chain OS=Mus musculus OX=10090 GN=Tuba1c PE=1 SV=1 |
| IMPCT_MOUSE | Protein IMPACT OS=Mus musculus OX=10090 GN=Impact PE=1 SV=2 |
| TOM34_MOUSE | Mitochondrial import receptor subunit TOM34 OS=Mus musculus OX=10090 GN=Tomm34 PE=1 SV=1 |
| SFXN5_MOUSE | Sideroflexin-5 OS=Mus musculus OX=10090 GN=Sfxn5 PE=1 SV=2 |

| | |
|---|---|
| CLCN6_MOUSE | Chloride transport protein 6 OS=Mus musculus OX=10090 GN=Clcn6 PE=1 SV=1 |
| P19137-DECOY | P19137 |
| GATM_MOUSE | Glycine amidinotransferase, mitochondrial OS=Mus musculus OX=10090 GN=Gatm PE=1 SV=1 |
| XPO7_MOUSE | Exportin-7 OS=Mus musculus OX=10090 GN=Xpo7 PE=1 SV=3 |
| LMNB2_MOUSE | Lamin-B2 OS=Mus musculus OX=10090 GN=Lmnb2 PE=1 SV=2 |
| PDS5B_MOUSE | Sister chromatid cohesion protein PDS5 homolog B OS=Mus musculus OX=10090 GN=Pds5b PE=1 SV=1 |
| SHLB2_MOUSE | Endophilin-B2 OS=Mus musculus OX=10090 GN=Sh3glb2 PE=1 SV=2 |
| IDI1_MOUSE | Isopentenyl-diphosphate Delta-isomerase 1 OS=Mus musculus OX=10090 GN=Idi1 PE=1 SV=1 |
| TMX3_MOUSE | Protein disulfide-isomerase TMX3 OS=Mus musculus OX=10090 GN=Tmx3 PE=1 SV=2 |
| TLN2_MOUSE | Talin-2 OS=Mus musculus OX=10090 GN=Tln2 PE=1 SV=3 |
| CUL5_MOUSE | Cullin-5 OS=Mus musculus OX=10090 GN=Cul5 PE=1 SV=3 |
| TPD54_MOUSE | Tumor protein D54 OS=Mus musculus OX=10090 GN=Tpd52l2 PE=1 SV=1 |
| PCLO_MOUSE | Protein piccolo OS=Mus musculus OX=10090 GN=Pclo PE=1 SV=4 |
| ABR_MOUSE | Active breakpoint cluster region-related protein OS=Mus musculus OX=10090 GN=Abr PE=1 SV=1 |
| PLRKT_MOUSE | Plasminogen receptor (KT) OS=Mus musculus OX=10090 GN=Plgrkt PE=1 SV=1 |
| GNAZ_MOUSE | Guanine nucleotide-binding protein G(z) subunit alpha OS=Mus musculus OX=10090 GN=Gnaz PE=1 SV=4 |
| GNAI3_MOUSE | Guanine nucleotide-binding protein G(i) subunit alpha OS=Mus musculus OX=10090 GN=Gnai3 PE=1 SV=3 |
| DHCR7_MOUSE | 7-dehydrocholesterol reductase OS=Mus musculus OX=10090 GN=Dhcr7 PE=1 SV=1 |

| | |
|---|---|
| NMT1_MOUSE | Glycylpeptide N-tetradecanoyltransferase 1 OS=Mus musculus OX=10090 GN=Nmt1 PE=1 SV=1 |
| COX5B_MOUSE | Cytochrome c oxidase subunit 5B, mitochondrial OS=Mus musculus OX=10090 GN=Cox5b PE=1 SV=1 |
| CON__P02672 | CON__P02672 |
| TCP4_MOUSE | Activated RNA polymerase II transcriptional coactivator p15 OS=Mus musculus OX=10090 GN=Sub1 PE=1 SV=3 |
| SRS10_MOUSE | Serine/arginine-rich splicing factor 10 OS=Mus musculus OX=10090 GN=Srsf10 PE=1 SV=2 |
| ACSF2_MOUSE | Medium-chain acyl-CoA ligase ACSF2, mitochondrial OS=Mus musculus OX=10090 GN=Acsf2 PE=1 SV=1 |
| MARK3_HUMAN | MAP/microtubule affinity-regulating kinase 3 OS=Homo sapiens OX=9606 GN=MARK3 PE=1 SV=5 |
| HPLN1_MOUSE | Hyaluronan and proteoglycan link protein 1 OS=Mus musculus OX=10090 GN=Hapln1 PE=1 SV=1 |
| SCPDL_MOUSE | Saccharopine dehydrogenase-like oxidoreductase OS=Mus musculus OX=10090 GN=Sccpdh PE=1 SV=1 |
| CD166_MOUSE | CD166 antigen OS=Mus musculus OX=10090 GN=Alcam PE=1 SV=3 |
| NDUAC_MOUSE | NADH dehydrogenase [ubiquinone] 1 alpha subcomplex subunit 12 OS=Mus musculus OX=10090 GN=Ndufa12 PE=1 SV=2 |
| RNPS1_MOUSE | RNA-binding protein with serine-rich domain 1 OS=Mus musculus OX=10090 GN=Rnps1 PE=1 SV=1 |
| NRDC_MOUSE | Nardilysin OS=Mus musculus OX=10090 GN=Nrdc PE=1 SV=1 |
| ARPC5_MOUSE | Actin-related protein 2/3 complex subunit 5 OS=Mus musculus OX=10090 GN=Arpc5 PE=1 SV=3 |
| SF3A3_MOUSE | Splicing factor 3A subunit 3 OS=Mus musculus OX=10090 GN=Sf3a3 PE=1 SV=2 |
| KV3A1_MOUSE | Ig kappa chain V-III region PC 2880/PC 1229 OS=Mus musculus OX=10090 PE=1 SV=1 |
| RAB21_MOUSE | Ras-related protein Rab-21 OS=Mus musculus OX=10090 GN=Rab21 PE=1 SV=4 |

| | |
|---|---|
| GDAP1_MOUSE | Ganglioside-induced differentiation-associated protein 1 OS=Mus musculus OX=10090 GN=Gdap1 PE=1 SV=1 |
| RDH11_MOUSE | Retinol dehydrogenase 11 OS=Mus musculus OX=10090 GN=Rdh11 PE=1 SV=2 |
| GPX4_MOUSE | Phospholipid hydroperoxide glutathione peroxidase OS=Mus musculus OX=10090 GN=Gpx4 PE=1 SV=4 |
| GFPT1_MOUSE | Glutamine--fructose-6-phosphate aminotransferase [isomerizing] 1 OS=Mus musculus OX=10090 GN=Gfpt1 PE=1 SV=3 |
| ARK72_MOUSE | Aflatoxin B1 aldehyde reductase member 2 OS=Mus musculus OX=10090 GN=Akr7a2 PE=1 SV=3 |
| NU4M_MOUSE | NADH-ubiquinone oxidoreductase chain 4 OS=Mus musculus OX=10090 GN=Mtnd4 PE=1 SV=1 |
| H13_MOUSE | Histone H1.3 OS=Mus musculus OX=10090 GN=Hist1h1d PE=1 SV=2 |
| CD81_MOUSE | CD81 antigen OS=Mus musculus OX=10090 GN=Cd81 PE=1 SV=2 |
| RER1_MOUSE | Protein RER1 OS=Mus musculus OX=10090 GN=Rer1 PE=1 SV=1 |
| ARAF_MOUSE | Serine/threonine-protein kinase A-Raf OS=Mus musculus OX=10090 GN=Araf PE=1 SV=2 |
| RNF14_MOUSE | E3 ubiquitin-protein ligase RNF14 OS=Mus musculus OX=10090 GN=Rnf14 PE=1 SV=2 |
| TXD17_MOUSE | Thioredoxin domain-containing protein 17 OS=Mus musculus OX=10090 GN=Txndc17 PE=1 SV=1 |
| RL35A_MOUSE | 60S ribosomal protein L35a OS=Mus musculus OX=10090 GN=Rpl35a PE=1 SV=2 |
| SPTN2_HUMAN | Spectrin beta chain, non-erythrocytic 2 OS=Homo sapiens OX=9606 GN=SPTBN2 PE=1 SV=3 |
| RL37A_MOUSE | 60S ribosomal protein L37a OS=Mus musculus OX=10090 GN=Rpl37a PE=1 SV=2 |
| EIF1_MOUSE | Eukaryotic translation initiation factor 1 OS=Mus musculus OX=10090 GN=Eif1 PE=1 SV=2 |
| FSD1_MOUSE | Fibronectin type III and SPRY domain-containing protein 1 OS=Mus musculus OX=10090 GN=Fsd1 PE=1 SV=1 |
| BIEA_MOUSE | Biliverdin reductase A OS=Mus musculus OX=10090 GN=Blvra PE=1 SV=1 |

| | |
|---|---|
| MTA1_MOUSE | Metastasis-associated protein MTA1 OS=Mus musculus OX=10090 GN=Mta1 PE=1 SV=1 |
| RAB12_MOUSE | Ras-related protein Rab-12 OS=Mus musculus OX=10090 GN=Rab12 PE=1 SV=3 |
| RRAGC_MOUSE | Ras-related GTP-binding protein C OS=Mus musculus OX=10090 GN=Rragc PE=1 SV=1 |
| F120C_MOUSE | Constitutive coactivator of PPAR-gamma-like protein 2 OS=Mus musculus OX=10090 GN=Fam120c PE=1 SV=3 |
| DRG2_MOUSE | Developmentally-regulated GTP-binding protein 2 OS=Mus musculus OX=10090 GN=Drg2 PE=1 SV=1 |
| PPM1G_MOUSE | Protein phosphatase 1G OS=Mus musculus OX=10090 GN=Ppm1g PE=1 SV=3 |
| SAP18_MOUSE | Histone deacetylase complex subunit SAP18 OS=Mus musculus OX=10090 GN=Sap18 PE=1 SV=1 |
| CADH2_MOUSE | Cadherin-2 OS=Mus musculus OX=10090 GN=Cdh2 PE=1 SV=2 |
| CSK22_MOUSE | Casein kinase II subunit alpha' OS=Mus musculus OX=10090 GN=Csnk2a2 PE=1 SV=1 |
| MTX1_MOUSE | Metaxin-1 OS=Mus musculus OX=10090 GN=Mtx1 PE=1 SV=1 |
| PUF60_MOUSE | Poly(U)-binding-splicing factor PUF60 OS=Mus musculus OX=10090 GN=Puf60 PE=1 SV=2 |
| SH3L1_MOUSE | SH3 domain-binding glutamic acid-rich-like protein OS=Mus musculus OX=10090 GN=Sh3bgrl PE=1 SV=1 |
| ATPK_MOUSE | ATP synthase subunit f, mitochondrial OS=Mus musculus OX=10090 GN=Atp5mf PE=1 SV=3 |
| VATG1_MOUSE | V-type proton ATPase subunit G 1 OS=Mus musculus OX=10090 GN=Atp6v1g1 PE=1 SV=3 |
| RS25_MOUSE | 40S ribosomal protein S25 OS=Mus musculus OX=10090 GN=Rps25 PE=1 SV=1 |
| SCAM5_MOUSE | Secretory carrier-associated membrane protein 5 OS=Mus musculus OX=10090 GN=Scamp5 PE=1 SV=1 |
| GPM6B_MOUSE | Neuronal membrane glycoprotein M6-b OS=Mus musculus OX=10090 GN=Gpm6b PE=1 SV=2 |
| ECI2_MOUSE | Enoyl-CoA delta isomerase 2, mitochondrial OS=Mus musculus OX=10090 GN=Eci2 PE=1 SV=2 |

| | |
|---|---|
| RAB5B_MOUSE | Ras-related protein Rab-5B OS=Mus musculus OX=10090 GN=Rab5b PE=1 SV=1 |
| METK2_MOUSE | S-adenosylmethionine synthase isoform type-2 OS=Mus musculus OX=10090 GN=Mat2a PE=1 SV=2 |
| SYNPR_MOUSE | Synaptoporin OS=Mus musculus OX=10090 GN=Synpr PE=1 SV=1 |
| NDUB5_MOUSE | NADH dehydrogenase [ubiquinone] 1 beta subcomplex subunit 5, mitochondrial OS=Mus musculus OX=10090 GN=Ndufb5 PE=1 SV=1 |
| MCU_MOUSE | Calcium uniporter protein, mitochondrial OS=Mus musculus OX=10090 GN=Mcu PE=1 SV=2 |
| RS24_MOUSE | 40S ribosomal protein S24 OS=Mus musculus OX=10090 GN=Rps24 PE=1 SV=1 |
| RALY_MOUSE | RNA-binding protein Raly OS=Mus musculus OX=10090 GN=Raly PE=1 SV=3 |
| ARXS1_MOUSE | Adipocyte-related X-chromosome expressed sequence 1 OS=Mus musculus OX=10090 GN=Arxes1 PE=1 SV=1 |
| MLP3A_MOUSE | Microtubule-associated proteins 1A/1B light chain 3A OS=Mus musculus OX=10090 GN=Map1lc3a PE=1 SV=1 |
| EMD_MOUSE | Emerin OS=Mus musculus OX=10090 GN=Emd PE=1 SV=1 |
| GRPE1_MOUSE | GrpE protein homolog 1, mitochondrial OS=Mus musculus OX=10090 GN=Grpel1 PE=1 SV=1 |
| GSTO1_MOUSE | Glutathione S-transferase omega-1 OS=Mus musculus OX=10090 GN=Gsto1 PE=1 SV=2 |
| SDCB1_MOUSE | Syntenin-1 OS=Mus musculus OX=10090 GN=Sdcbp PE=1 SV=1 |
| TM9S3_MOUSE | Transmembrane 9 superfamily member 3 OS=Mus musculus OX=10090 GN=Tm9sf3 PE=1 SV=1 |
| ODB2_MOUSE | Lipoamide acyltransferase component of branched-chain alpha-keto acid dehydrogenase complex, mitochondrial OS=Mus musculus OX=10090 GN=Dbt PE=1 SV=2 |
| DIRA2_MOUSE | GTP-binding protein Di-Ras2 OS=Mus musculus OX=10090 GN=Diras2 PE=1 SV=1 |
| GMFB_MOUSE | Glia maturation factor beta OS=Mus musculus OX=10090 GN=Gmfb PE=1 SV=3 |
| SEC13_MOUSE | Protein SEC13 homolog OS=Mus musculus OX=10090 GN=Sec13 PE=1 SV=3 |

| | |
|---|---|
| IDE_MOUSE | Insulin-degrading enzyme OS=Mus musculus OX=10090 GN=Ide PE=1 SV=1 |
| HSDL2_MOUSE | Hydroxysteroid dehydrogenase-like protein 2 OS=Mus musculus OX=10090 GN=Hsdl2 PE=1 SV=1 |
| LC7L3_MOUSE | Luc7-like protein 3 OS=Mus musculus OX=10090 GN=Luc7l3 PE=1 SV=1 |
| SNG1_MOUSE | Synaptogyrin-1 OS=Mus musculus OX=10090 GN=Syngr1 PE=1 SV=2 |
| SNX4_MOUSE | Sorting nexin-4 OS=Mus musculus OX=10090 GN=Snx4 PE=1 SV=1 |
| FETUA_MOUSE | Alpha-2-HS-glycoprotein OS=Mus musculus OX=10090 GN=Ahsg PE=1 SV=1 |
| CX6B1_MOUSE | Cytochrome c oxidase subunit 6B1 OS=Mus musculus OX=10090 GN=Cox6b1 PE=1 SV=2 |
| H2AX_MOUSE | Histone H2AX OS=Mus musculus OX=10090 GN=H2afx PE=1 SV=2 |
| ATPBM_ARATH | ATP synthase subunit beta-1, mitochondrial OS=Arabidopsis thaliana OX=3702 GN=At5g08670 PE=1 SV=1 |
| PEX5_MOUSE | Peroxisomal targeting signal 1 receptor OS=Mus musculus OX=10090 GN=Pex5 PE=1 SV=2 |
| ADPRH_MOUSE | [Protein ADP-ribosylarginine] hydrolase OS=Mus musculus OX=10090 GN=Adprh PE=1 SV=1 |
| SNRPA_MOUSE | U1 small nuclear ribonucleoprotein A OS=Mus musculus OX=10090 GN=Snrpa PE=1 SV=3 |
| PRP6_MOUSE | Pre-mRNA-processing factor 6 OS=Mus musculus OX=10090 GN=Prpf6 PE=1 SV=1 |
| DIP2B_MOUSE | Disco-interacting protein 2 homolog B OS=Mus musculus OX=10090 GN=Dip2b PE=1 SV=1 |
| PDE1B_MOUSE | Calcium/calmodulin-dependent 3',5'-cyclic nucleotide phosphodiesterase 1B OS=Mus musculus OX=10090 GN=Pde1b PE=1 SV=2 |
| COF2_MOUSE | Cofilin-2 OS=Mus musculus OX=10090 GN=Cfl2 PE=1 SV=1 |
| NAC1_MOUSE | Sodium/calcium exchanger 1 OS=Mus musculus OX=10090 GN=Slc8a1 PE=1 SV=1 |
| CSN7B_MOUSE | COP9 signalosome complex subunit 7b OS=Mus musculus OX=10090 GN=Cops7b PE=1 SV=1 |
| UBP2L_MOUSE | Ubiquitin-associated protein 2-like OS=Mus musculus OX=10090 GN=Ubap2l PE=1 SV=1 |

| | |
|---|---|
| RBP2_MOUSE | E3 SUMO-protein ligase RanBP2 OS=Mus musculus OX=10090 GN=Ranbp2 PE=1 SV=2 |
| PSME2_MOUSE | Proteasome activator complex subunit 2 OS=Mus musculus OX=10090 GN=Psme2 PE=1 SV=4 |
| NTRK2_MOUSE | BDNF/NT-3 growth factors receptor OS=Mus musculus OX=10090 GN=Ntrk2 PE=1 SV=1 |
| ATCAY_MOUSE | Caytaxin OS=Mus musculus OX=10090 GN=Atcay PE=1 SV=1 |
| CCD47_MOUSE | Coiled-coil domain-containing protein 47 OS=Mus musculus OX=10090 GN=Ccdc47 PE=1 SV=2 |
| NU155_MOUSE | Nuclear pore complex protein Nup155 OS=Mus musculus OX=10090 GN=Nup155 PE=1 SV=1 |
| ZO1_MOUSE | Tight junction protein ZO-1 OS=Mus musculus OX=10090 GN=Tjp1 PE=1 SV=2 |
| DYL1_MOUSE | Dynein light chain 1, cytoplasmic OS=Mus musculus OX=10090 GN=Dynll1 PE=1 SV=1 |
| NCBP1_MOUSE | Nuclear cap-binding protein subunit 1 OS=Mus musculus OX=10090 GN=Ncbp1 PE=1 SV=2 |
| GPSM1_MOUSE | G-protein-signaling modulator 1 OS=Mus musculus OX=10090 GN=Gpsm1 PE=1 SV=3 |
| L1CAM_MOUSE | Neural cell adhesion molecule L1 OS=Mus musculus OX=10090 GN=L1cam PE=1 SV=1 |
| ACACA_MOUSE | Acetyl-CoA carboxylase 1 OS=Mus musculus OX=10090 GN=Acaca PE=1 SV=1 |
| ERF3A_MOUSE | Eukaryotic peptide chain release factor GTP-binding subunit ERF3A OS=Mus musculus OX=10090 GN=Gspt1 PE=1 SV=2 |
| NDUB8_MOUSE | NADH dehydrogenase [ubiquinone] 1 beta subcomplex subunit 8, mitochondrial OS=Mus musculus OX=10090 GN=Ndufb8 PE=1 SV=1 |
| DTD1_MOUSE | D-aminoacyl-tRNA deacylase 1 OS=Mus musculus OX=10090 GN=Dtd1 PE=1 SV=2 |
| VATG2_MOUSE | V-type proton ATPase subunit G 2 OS=Mus musculus OX=10090 GN=Atp6v1g2 PE=1 SV=1 |
| ARL8A_MOUSE | ADP-ribosylation factor-like protein 8A OS=Mus musculus OX=10090 GN=Arl8a PE=1 SV=1 |
| SF01_MOUSE | Splicing factor 1 OS=Mus musculus OX=10090 GN=Sf1 PE=1 SV=6 |
| UGPA_MOUSE | UTP--glucose-1-phosphate uridylyltransferase OS=Mus musculus OX=10090 GN=Ugp2 PE=1 SV=3 |

| | |
|---|---|
| COX6C_MOUSE | Cytochrome c oxidase subunit 6C OS=Mus musculus OX=10090 GN=Cox6c PE=1 SV=3 |
| VTI1B_MOUSE | Vesicle transport through interaction with t-SNAREs homolog 1B OS=Mus musculus OX=10090 GN=Vti1b PE=1 SV=1 |
| FLOT1_MOUSE | Flotillin-1 OS=Mus musculus OX=10090 GN=Flot1 PE=1 SV=1 |
| SUCB2_MOUSE | Succinate--CoA ligase [GDP-forming] subunit beta, mitochondrial OS=Mus musculus OX=10090 GN=Suclg2 PE=1 SV=3 |
| DCTN3_MOUSE | Dynactin subunit 3 OS=Mus musculus OX=10090 GN=Dctn3 PE=1 SV=2 |
| TBB6_MOUSE | Tubulin beta-6 chain OS=Mus musculus OX=10090 GN=Tubb6 PE=1 SV=1 |
| UB2L3_MOUSE | Ubiquitin-conjugating enzyme E2 L3 OS=Mus musculus OX=10090 GN=Ube2l3 PE=1 SV=1 |
| KC1A_MOUSE | Casein kinase I isoform alpha OS=Mus musculus OX=10090 GN=Csnk1a1 PE=1 SV=2 |
| KCC2G_MOUSE | Calcium/calmodulin-dependent protein kinase type II subunit gamma OS=Mus musculus OX=10090 GN=Camk2g PE=1 SV=1 |
| SRSF5_HUMAN | Serine/arginine-rich splicing factor 5 OS=Homo sapiens OX=9606 GN=SRSF5 PE=1 SV=1 |
| ELAV4_MOUSE | ELAV-like protein 4 OS=Mus musculus OX=10090 GN=Elavl4 PE=1 SV=1 |
| BCAT1_MOUSE | Branched-chain-amino-acid aminotransferase, cytosolic OS=Mus musculus OX=10090 GN=Bcat1 PE=1 SV=2 |
| HVM44_MOUSE | Ig heavy chain V region PJ14 OS=Mus musculus OX=10090 PE=1 SV=1 |
| LMAN1_MOUSE | Protein ERGIC-53 OS=Mus musculus OX=10090 GN=Lman1 PE=1 SV=1 |
| TTL12_MOUSE | Tubulin--tyrosine ligase-like protein 12 OS=Mus musculus OX=10090 GN=Ttll12 PE=1 SV=1 |
| GLYR1_MOUSE | Putative oxidoreductase GLYR1 OS=Mus musculus OX=10090 GN=Glyr1 PE=1 SV=1 |
| MTCH1_MOUSE | Mitochondrial carrier homolog 1 OS=Mus musculus OX=10090 GN=Mtch1 PE=1 SV=1 |
| PFD3_MOUSE | Prefoldin subunit 3 OS=Mus musculus OX=10090 GN=Vbp1 PE=1 SV=2 |

| | |
|---|---|
| ULA1_MOUSE | NEDD8-activating enzyme E1 regulatory subunit OS=Mus musculus OX=10090 GN=Nae1 PE=1 SV=1 |
| PCYOX_MOUSE | Prenylcysteine oxidase OS=Mus musculus OX=10090 GN=Pcyox1 PE=1 SV=1 |
| FXR1_MOUSE | Fragile X mental retardation syndrome-related protein 1 OS=Mus musculus OX=10090 GN=Fxr1 PE=1 SV=2 |
| PI4KA_MOUSE | Phosphatidylinositol 4-kinase alpha OS=Mus musculus OX=10090 GN=Pi4ka PE=1 SV=2 |
| NDUB4_MOUSE | NADH dehydrogenase [ubiquinone] 1 beta subcomplex subunit 4 OS=Mus musculus OX=10090 GN=Ndufb4 PE=1 SV=3 |
| TXTP_MOUSE | Tricarboxylate transport protein, mitochondrial OS=Mus musculus OX=10090 GN=Slc25a1 PE=1 SV=1 |
| OXR1_MOUSE | Oxidation resistance protein 1 OS=Mus musculus OX=10090 GN=Oxr1 PE=1 SV=3 |
| VPS51_MOUSE | Vacuolar protein sorting-associated protein 51 homolog OS=Mus musculus OX=10090 GN=Vps51 PE=1 SV=2 |
| NRCAM_MOUSE | Neuronal cell adhesion molecule OS=Mus musculus OX=10090 GN=Nrcam PE=1 SV=2 |
| DJB11_MOUSE | DnaJ homolog subfamily B member 11 OS=Mus musculus OX=10090 GN=Dnajb11 PE=1 SV=1 |
| TRNT1_MOUSE | CCA tRNA nucleotidyltransferase 1, mitochondrial OS=Mus musculus OX=10090 GN=Trnt1 PE=1 SV=1 |
| HPCA_HUMAN | Neuron-specific calcium-binding protein hippocalcin OS=Homo sapiens OX=9606 GN=HPCA PE=1 SV=2 |
| STMN3_MOUSE | Stathmin-3 OS=Mus musculus OX=10090 GN=Stmn3 PE=1 SV=1 |
| S4A4_MOUSE | Electrogenic sodium bicarbonate cotransporter 1 OS=Mus musculus OX=10090 GN=Slc4a4 PE=1 SV=2 |
| RL26_MOUSE | 60S ribosomal protein L26 OS=Mus musculus OX=10090 GN=Rpl26 PE=1 SV=1 |
| AIMP1_MOUSE | Aminoacyl tRNA synthase complex-interacting multifunctional protein 1 OS=Mus musculus OX=10090 GN=Aimp1 PE=1 SV=2 |

| | |
|---|---|
| HIBCH_MOUSE | 3-hydroxyisobutyryl-CoA hydrolase, mitochondrial OS=Mus musculus OX=10090 GN=Hibch PE=1 SV=1 |
| AT1B3_MOUSE | Sodium/potassium-transporting ATPase subunit beta-3 OS=Mus musculus OX=10090 GN=Atp1b3 PE=1 SV=1 |
| AFG32_MOUSE | AFG3-like protein 2 OS=Mus musculus OX=10090 GN=Afg3l2 PE=1 SV=1 |
| SRPK2_MOUSE | SRSF protein kinase 2 OS=Mus musculus OX=10090 GN=Srpk2 PE=1 SV=2 |
| ACINU_MOUSE | Apoptotic chromatin condensation inducer in the nucleus OS=Mus musculus OX=10090 GN=Acin1 PE=1 SV=3 |
| MCCA_MOUSE | Methylcrotonoyl-CoA carboxylase subunit alpha, mitochondrial OS=Mus musculus OX=10090 GN=Mccc1 PE=1 SV=2 |
| RHG35_MOUSE | Rho GTPase-activating protein 35 OS=Mus musculus OX=10090 GN=Arhgap35 PE=1 SV=3 |
| 2A5G_MOUSE | Serine/threonine-protein phosphatase 2A 56 kDa regulatory subunit gamma isoform OS=Mus musculus OX=10090 GN=Ppp2r5c PE=1 SV=2 |
| SEM3C_MOUSE | Semaphorin-3C OS=Mus musculus OX=10090 GN=Sema3c PE=1 SV=2 |
| CNOT1_HUMAN | CCR4-NOT transcription complex subunit 1 OS=Homo sapiens OX=9606 GN=CNOT1 PE=1 SV=2 |
| NDUAD_MOUSE | NADH dehydrogenase [ubiquinone] 1 alpha subcomplex subunit 13 OS=Mus musculus OX=10090 GN=Ndufa13 PE=1 SV=3 |
| HA11_MOUSE | H-2 class I histocompatibility antigen, D-B alpha chain OS=Mus musculus OX=10090 GN=H2-D1 PE=1 SV=2 |
| LAMP1_MOUSE | Lysosome-associated membrane glycoprotein 1 OS=Mus musculus OX=10090 GN=Lamp1 PE=1 SV=2 |
| CON__P07477 | CON__P07477 |
| UBC9_MOUSE | SUMO-conjugating enzyme UBC9 OS=Mus musculus OX=10090 GN=Ube2i PE=1 SV=1 |
| NNRE_MOUSE | NAD(P)H-hydrate epimerase OS=Mus musculus OX=10090 GN=Naxe PE=1 SV=1 |
| RM12_MOUSE | 39S ribosomal protein L12, mitochondrial OS=Mus musculus OX=10090 GN=Mrpl12 PE=1 SV=2 |

| | |
|---|---|
| RU1C_MOUSE | U1 small nuclear ribonucleoprotein C OS=Mus musculus OX=10090 GN=Snrpc PE=1 SV=1 |
| HMGB2_MOUSE | High mobility group protein B2 OS=Mus musculus OX=10090 GN=Hmgb2 PE=1 SV=3 |
| PPR21_MOUSE | Protein phosphatase 1 regulatory subunit 21 OS=Mus musculus OX=10090 GN=Ppp1r21 PE=1 SV=2 |
| MSI2H_MOUSE | RNA-binding protein Musashi homolog 2 OS=Mus musculus OX=10090 GN=Msi2 PE=1 SV=1 |
| ACBP_MOUSE | Acyl-CoA-binding protein OS=Mus musculus OX=10090 GN=Dbi PE=1 SV=2 |
| UBXN1_MOUSE | UBX domain-containing protein 1 OS=Mus musculus OX=10090 GN=Ubxn1 PE=1 SV=1 |
| RSMB_MOUSE | Small nuclear ribonucleoprotein-associated protein B OS=Mus musculus OX=10090 GN=Snrpb PE=1 SV=1 |
| AT2B3_HUMAN | Plasma membrane calcium-transporting ATPase 3 OS=Homo sapiens OX=9606 GN=ATP2B3 PE=1 SV=3 |
| STMN2_MOUSE | Stathmin-2 OS=Mus musculus OX=10090 GN=Stmn2 PE=1 SV=1 |
| MYLK_MOUSE | Myosin light chain kinase, smooth muscle OS=Mus musculus OX=10090 GN=Mylk PE=1 SV=3 |
| BCLF1_MOUSE | Bcl-2-associated transcription factor 1 OS=Mus musculus OX=10090 GN=Bclaf1 PE=1 SV=2 |
| IMA4_MOUSE | Importin subunit alpha-4 OS=Mus musculus OX=10090 GN=Kpna3 PE=1 SV=1 |
| MT3_MOUSE | Metallothionein-3 OS=Mus musculus OX=10090 GN=Mt3 PE=1 SV=1 |
| LEGL_MOUSE | Galectin-related protein OS=Mus musculus OX=10090 GN=Lgalsl PE=1 SV=1 |
| RO60_MOUSE | 60 kDa SS-A/Ro ribonucleoprotein OS=Mus musculus OX=10090 GN=RO60 PE=1 SV=1 |
| COPZ1_MOUSE | Coatomer subunit zeta-1 OS=Mus musculus OX=10090 GN=Copz1 PE=1 SV=1 |
| THTR_MOUSE | Thiosulfate sulfurtransferase OS=Mus musculus OX=10090 GN=Tst PE=1 SV=3 |
| GBG7_MOUSE | Guanine nucleotide-binding protein G(I)/G(S)/G(O) subunit gamma-7 OS=Mus musculus OX=10090 GN=Gng7 PE=1 SV=2 |

| | |
|---|---|
| SRRM1_MOUSE | Serine/arginine repetitive matrix protein 1 OS=Mus musculus OX=10090 GN=Srrm1 PE=1 SV=2 |
| DOCK7_MOUSE | Dedicator of cytokinesis protein 7 OS=Mus musculus OX=10090 GN=Dock7 PE=1 SV=3 |
| PPM1A_MOUSE | Protein phosphatase 1A OS=Mus musculus OX=10090 GN=Ppm1a PE=1 SV=1 |
| NMDZ1_MOUSE | Glutamate receptor ionotropic, NMDA 1 OS=Mus musculus OX=10090 GN=Grin1 PE=1 SV=1 |
| ATLA2_MOUSE | Atlastin-2 OS=Mus musculus OX=10090 GN=Atl2 PE=1 SV=1 |
| TM9S4_MOUSE | Transmembrane 9 superfamily member 4 OS=Mus musculus OX=10090 GN=Tm9sf4 PE=1 SV=1 |
| IMPA1_MOUSE | Inositol monophosphatase 1 OS=Mus musculus OX=10090 GN=Impa1 PE=1 SV=1 |
| RT23_MOUSE | 28S ribosomal protein S23, mitochondrial OS=Mus musculus OX=10090 GN=Mrps23 PE=1 SV=1 |
| TTYH3_MOUSE | Protein tweety homolog 3 OS=Mus musculus OX=10090 GN=Ttyh3 PE=1 SV=1 |
| ZFR_MOUSE | Zinc finger RNA-binding protein OS=Mus musculus OX=10090 GN=Zfr PE=1 SV=2 |
| SRPRB_MOUSE | Signal recognition particle receptor subunit beta OS=Mus musculus OX=10090 GN=Srprb PE=1 SV=1 |
| MCES_MOUSE | mRNA cap guanine-N7 methyltransferase OS=Mus musculus OX=10090 GN=Rnmt PE=1 SV=1 |
| TIAR_MOUSE | Nucleolysin TIAR OS=Mus musculus OX=10090 GN=Tial1 PE=1 SV=1 |
| KAP0_MOUSE | cAMP-dependent protein kinase type I-alpha regulatory subunit OS=Mus musculus OX=10090 GN=Prkar1a PE=1 SV=3 |
| PIGS_MOUSE | GPI transamidase component PIG-S OS=Mus musculus OX=10090 GN=Pigs PE=1 SV=3 |
| ARP10_MOUSE | Actin-related protein 10 OS=Mus musculus OX=10090 GN=Actr10 PE=1 SV=2 |
| ANM8_MOUSE | Protein arginine N-methyltransferase 8 OS=Mus musculus OX=10090 GN=Prmt8 PE=1 SV=2 |
| BRK1_MOUSE | Protein BRICK1 OS=Mus musculus OX=10090 GN=Brk1 PE=1 SV=1 |

| | |
|---|---|
| RPGP1_MOUSE | Rap1 GTPase-activating protein 1 OS=Mus musculus OX=10090 GN=Rap1gap PE=1 SV=2 |
| CP131_MOUSE | Centrosomal protein of 131 kDa OS=Mus musculus OX=10090 GN=Cep131 PE=1 SV=2 |
| HCFC1_MOUSE | Host cell factor 1 OS=Mus musculus OX=10090 GN=Hcfc1 PE=1 SV=2 |
| ADRM1_MOUSE | Proteasomal ubiquitin receptor ADRM1 OS=Mus musculus OX=10090 GN=Adrm1 PE=1 SV=2 |
| STRN4_MOUSE | Striatin-4 OS=Mus musculus OX=10090 GN=Strn4 PE=1 SV=2 |
| CAP2_MOUSE | Adenylyl cyclase-associated protein 2 OS=Mus musculus OX=10090 GN=Cap2 PE=1 SV=1 |
| CADM4_MOUSE | Cell adhesion molecule 4 OS=Mus musculus OX=10090 GN=Cadm4 PE=1 SV=1 |
| RBX1_MOUSE | E3 ubiquitin-protein ligase RBX1 OS=Mus musculus OX=10090 GN=Rbx1 PE=1 SV=1 |
| DENR_MOUSE | Density-regulated protein OS=Mus musculus OX=10090 GN=Denr PE=1 SV=1 |
| NH2L1_MOUSE | NHP2-like protein 1 OS=Mus musculus OX=10090 GN=Snu13 PE=1 SV=4 |
| LYPA1_MOUSE | Acyl-protein thioesterase 1 OS=Mus musculus OX=10090 GN=Lypla1 PE=1 SV=1 |
| NDUA7_MOUSE | NADH dehydrogenase [ubiquinone] 1 alpha subcomplex subunit 7 OS=Mus musculus OX=10090 GN=Ndufa7 PE=1 SV=3 |
| TIM50_MOUSE | Mitochondrial import inner membrane translocase subunit TIM50 OS=Mus musculus OX=10090 GN=Timm50 PE=1 SV=1 |
| CDIPT_MOUSE | CDP-diacylglycerol--inositol 3-phosphatidyltransferase OS=Mus musculus OX=10090 GN=Cdipt PE=1 SV=1 |
| EVL_MOUSE | Ena/VASP-like protein OS=Mus musculus OX=10090 GN=Evl PE=1 SV=2 |
| NUDT3_MOUSE | Diphosphoinositol polyphosphate phosphohydrolase 1 OS=Mus musculus OX=10090 GN=Nudt3 PE=1 SV=1 |
| SODC_MOUSE | Superoxide dismutase [Cu-Zn] OS=Mus musculus OX=10090 GN=Sod1 PE=1 SV=2 |
| EFTS_MOUSE | Elongation factor Ts, mitochondrial OS=Mus musculus OX=10090 GN=Tsfm PE=1 SV=1 |
| SMU1_MOUSE | WD40 repeat-containing protein SMU1 OS=Mus musculus OX=10090 GN=Smu1 PE=2 SV=2 |

| | |
|---|---|
| RS26_MOUSE | 40S ribosomal protein S26 OS=Mus musculus OX=10090 GN=Rps26 PE=1 SV=3 |
| RS51_ARATH | 40S ribosomal protein S5-1 OS=Arabidopsis thaliana OX=3702 GN=RPS5A PE=1 SV=1 |
| IGHA_MOUSE | Ig alpha chain C region OS=Mus musculus OX=10090 PE=1 SV=1 |
| TYB4_RAT | Thymosin beta-4 OS=Rattus norvegicus OX=10116 GN=Tmsb4x PE=1 SV=2 |
| VPS36_MOUSE | Vacuolar protein-sorting-associated protein 36 OS=Mus musculus OX=10090 GN=Vps36 PE=1 SV=1 |
| BPHL_MOUSE | Valacyclovir hydrolase OS=Mus musculus OX=10090 GN=Bphl PE=1 SV=1 |
| P35969-DECOY | P35969 |
| ELP3_MOUSE | Elongator complex protein 3 OS=Mus musculus OX=10090 GN=Elp3 PE=1 SV=1 |
| CHRD1_MOUSE | Cysteine and histidine-rich domain-containing protein 1 OS=Mus musculus OX=10090 GN=Chordc1 PE=1 SV=1 |
| PUR8_MOUSE | Adenylosuccinate lyase OS=Mus musculus OX=10090 GN=Adsl PE=1 SV=2 |
| COPE_MOUSE | Coatomer subunit epsilon OS=Mus musculus OX=10090 GN=Cope PE=1 SV=3 |
| EI2BA_MOUSE | Translation initiation factor eIF-2B subunit alpha OS=Mus musculus OX=10090 GN=Eif2b1 PE=1 SV=1 |
| SAHH3_MOUSE | Putative adenosylhomocysteinase 3 OS=Mus musculus OX=10090 GN=Ahcyl2 PE=1 SV=1 |
| ABCF1_MOUSE | ATP-binding cassette sub-family F member 1 OS=Mus musculus OX=10090 GN=Abcf1 PE=1 SV=1 |
| MYH14_MOUSE | Myosin-14 OS=Mus musculus OX=10090 GN=Myh14 PE=1 SV=1 |
| Q61644-DECOY | Q61644 |
| RCC2_MOUSE | Protein RCC2 OS=Mus musculus OX=10090 GN=Rcc2 PE=1 SV=1 |
| GSHR_MOUSE | Glutathione reductase, mitochondrial OS=Mus musculus OX=10090 GN=Gsr PE=1 SV=3 |
| VPS28_MOUSE | Vacuolar protein sorting-associated protein 28 homolog OS=Mus musculus OX=10090 GN=Vps28 PE=1 SV=1 |

| | |
|---|---|
| MAOM_MOUSE | NAD-dependent malic enzyme, mitochondrial OS=Mus musculus OX=10090 GN=Me2 PE=1 SV=1 |
| SMCA4_MOUSE | Transcription activator BRG1 OS=Mus musculus OX=10090 GN=Smarca4 PE=1 SV=1 |
| VPS45_MOUSE | Vacuolar protein sorting-associated protein 45 OS=Mus musculus OX=10090 GN=Vps45 PE=1 SV=1 |
| FRIL1_MOUSE | Ferritin light chain 1 OS=Mus musculus OX=10090 GN=Ftl1 PE=1 SV=2 |
| SRGP3_MOUSE | SLIT-ROBO Rho GTPase-activating protein 3 OS=Mus musculus OX=10090 GN=Srgap3 PE=1 SV=1 |
| PGCA_MOUSE | Aggrecan core protein OS=Mus musculus OX=10090 GN=Acan PE=1 SV=2 |
| EI3JA_MOUSE | Eukaryotic translation initiation factor 3 subunit J-A OS=Mus musculus OX=10090 GN=Eif3j1 PE=2 SV=1 |
| PI51C_MOUSE | Phosphatidylinositol 4-phosphate 5-kinase type-1 gamma OS=Mus musculus OX=10090 GN=Pip5k1c PE=1 SV=2 |
| DAD1_MOUSE | Dolichyl-diphosphooligosaccharide--protein glycosyltransferase subunit DAD1 OS=Mus musculus OX=10090 GN=Dad1 PE=1 SV=3 |
| SRP54_MOUSE | Signal recognition particle 54 kDa protein OS=Mus musculus OX=10090 GN=Srp54 PE=1 SV=2 |
| DD19A_MOUSE | ATP-dependent RNA helicase DDX19A OS=Mus musculus OX=10090 GN=Ddx19a PE=1 SV=2 |
| TBCB_MOUSE | Tubulin-folding cofactor B OS=Mus musculus OX=10090 GN=Tbcb PE=1 SV=2 |
| PDLI5_MOUSE | PDZ and LIM domain protein 5 OS=Mus musculus OX=10090 GN=Pdlim5 PE=1 SV=4 |
| RBM25_MOUSE | RNA-binding protein 25 OS=Mus musculus OX=10090 GN=Rbm25 PE=1 SV=2 |
| IF5_MOUSE | Eukaryotic translation initiation factor 5 OS=Mus musculus OX=10090 GN=Eif5 PE=1 SV=1 |
| CON__P78385 | CON__P78385 |
| VAS1_MOUSE | V-type proton ATPase subunit S1 OS=Mus musculus OX=10090 GN=Atp6ap1 PE=1 SV=1 |
| REEP5_MOUSE | Receptor expression-enhancing protein 5 OS=Mus musculus OX=10090 GN=Reep5 PE=1 SV=1 |

| | |
|---|---|
| NDUA2_MOUSE | NADH dehydrogenase [ubiquinone] 1 alpha subcomplex subunit 2 OS=Mus musculus OX=10090 GN=Ndufa2 PE=1 SV=3 |
| PPAC_MOUSE | Low molecular weight phosphotyrosine protein phosphatase OS=Mus musculus OX=10090 GN=Acp1 PE=1 SV=3 |
| RRAS2_MOUSE | Ras-related protein R-Ras2 OS=Mus musculus OX=10090 GN=Rras2 PE=1 SV=1 |
| RBBP7_MOUSE | Histone-binding protein RBBP7 OS=Mus musculus OX=10090 GN=Rbbp7 PE=1 SV=1 |
| SSRD_MOUSE | Translocon-associated protein subunit delta OS=Mus musculus OX=10090 GN=Ssr4 PE=1 SV=1 |
| TBCD_MOUSE | Tubulin-specific chaperone D OS=Mus musculus OX=10090 GN=Tbcd PE=1 SV=1 |
| RBM8A_MOUSE | RNA-binding protein 8A OS=Mus musculus OX=10090 GN=Rbm8a PE=1 SV=4 |
| CN37_MOUSE | 2',3'-cyclic-nucleotide 3'-phosphodiesterase OS=Mus musculus OX=10090 GN=Cnp PE=1 SV=3 |
| PTMS_MOUSE | Parathymosin OS=Mus musculus OX=10090 GN=Ptms PE=1 SV=3 |
| ACPM_MOUSE | Acyl carrier protein, mitochondrial OS=Mus musculus OX=10090 GN=Ndufab1 PE=1 SV=1 |
| ITM2C_MOUSE | Integral membrane protein 2C OS=Mus musculus OX=10090 GN=Itm2c PE=1 SV=2 |
| MIC27_MOUSE | MICOS complex subunit Mic27 OS=Mus musculus OX=10090 GN=Apool PE=1 SV=1 |
| PLXA1_MOUSE | Plexin-A1 OS=Mus musculus OX=10090 GN=Plxna1 PE=1 SV=1 |
| H14_MOUSE | Histone H1.4 OS=Mus musculus OX=10090 GN=Hist1h1e PE=1 SV=2 |
| SC61B_MOUSE | Protein transport protein Sec61 subunit beta OS=Mus musculus OX=10090 GN=Sec61b PE=1 SV=3 |
| AP1M1_MOUSE | AP-1 complex subunit mu-1 OS=Mus musculus OX=10090 GN=Ap1m1 PE=1 SV=3 |
| PACN2_MOUSE | Protein kinase C and casein kinase substrate in neurons protein 2 OS=Mus musculus OX=10090 GN=Pacsin2 PE=1 SV=1 |
| XRN2_MOUSE | 5'-3' exoribonuclease 2 OS=Mus musculus OX=10090 GN=Xrn2 PE=1 SV=1 |

| | |
|---|---|
| RS28_MOUSE | 40S ribosomal protein S28 OS=Mus musculus OX=10090 GN=Rps28 PE=1 SV=1 |
| UB2D2_MOUSE | Ubiquitin-conjugating enzyme E2 D2 OS=Mus musculus OX=10090 GN=Ube2d2 PE=1 SV=1 |
| AN32B_MOUSE | Acidic leucine-rich nuclear phosphoprotein 32 family member B OS=Mus musculus OX=10090 GN=Anp32b PE=1 SV=1 |
| PABP2_MOUSE | Polyadenylate-binding protein 2 OS=Mus musculus OX=10090 GN=Pabpn1 PE=1 SV=3 |
| QCR9_MOUSE | Cytochrome b-c1 complex subunit 9 OS=Mus musculus OX=10090 GN=Uqcr10 PE=1 SV=1 |
| UBA6_MOUSE | Ubiquitin-like modifier-activating enzyme 6 OS=Mus musculus OX=10090 GN=Uba6 PE=1 SV=1 |
| WDR47_MOUSE | WD repeat-containing protein 47 OS=Mus musculus OX=10090 GN=Wdr47 PE=1 SV=2 |
| RM37_MOUSE | 39S ribosomal protein L37, mitochondrial OS=Mus musculus OX=10090 GN=Mrpl37 PE=1 SV=1 |
| KLC2_MOUSE | Kinesin light chain 2 OS=Mus musculus OX=10090 GN=Klc2 PE=1 SV=1 |
| MESD_MOUSE | LRP chaperone MESD OS=Mus musculus OX=10090 GN=Mesd PE=1 SV=1 |
| FRIH_MOUSE | Ferritin heavy chain OS=Mus musculus OX=10090 GN=Fth1 PE=1 SV=2 |
| MPP6_MOUSE | MAGUK p55 subfamily member 6 OS=Mus musculus OX=10090 GN=Mpp6 PE=1 SV=1 |
| EIF3M_MOUSE | Eukaryotic translation initiation factor 3 subunit M OS=Mus musculus OX=10090 GN=Eif3m PE=1 SV=1 |
| MYCT_MOUSE | Proton myo-inositol cotransporter OS=Mus musculus OX=10090 GN=Slc2a13 PE=1 SV=2 |
| NFASC_MOUSE | Neurofascin OS=Mus musculus OX=10090 GN=Nfasc PE=1 SV=1 |
| PTPRS_MOUSE | Receptor-type tyrosine-protein phosphatase S OS=Mus musculus OX=10090 GN=Ptprs PE=1 SV=1 |
| NAKD2_MOUSE | NAD kinase 2, mitochondrial OS=Mus musculus OX=10090 GN=Nadk2 PE=1 SV=2 |
| ACADS_MOUSE | Short-chain specific acyl-CoA dehydrogenase, mitochondrial OS=Mus musculus OX=10090 GN=Acads PE=1 SV=2 |

| | |
|---|---|
| CON__P02070 | CON__P02070 |
| HOOK3_MOUSE | Protein Hook homolog 3 OS=Mus musculus OX=10090 GN=Hook3 PE=1 SV=2 |
| F162A_MOUSE | Protein FAM162A OS=Mus musculus OX=10090 GN=Fam162a PE=1 SV=1 |
| NAV1_MOUSE | Neuron navigator 1 OS=Mus musculus OX=10090 GN=Nav1 PE=1 SV=2 |
| TIP_MOUSE | T-cell immunomodulatory protein OS=Mus musculus OX=10090 GN=Itfg1 PE=1 SV=2 |
| PP6R3_MOUSE | Serine/threonine-protein phosphatase 6 regulatory subunit 3 OS=Mus musculus OX=10090 GN=Ppp6r3 PE=1 SV=1 |
| GRM5_MOUSE | Metabotropic glutamate receptor 5 OS=Mus musculus OX=10090 GN=Grm5 PE=1 SV=2 |
| CON__ENSEMBL:ENSBTAP00000024146 | CON__ENSEMBL:ENSBTAP00000024146 |
| PLST_MOUSE | Plastin-3 OS=Mus musculus OX=10090 GN=Pls3 PE=1 SV=3 |
| TS101_MOUSE | Tumor susceptibility gene 101 protein OS=Mus musculus OX=10090 GN=Tsg101 PE=1 SV=2 |
| MADD_HUMAN | MAP kinase-activating death domain protein OS=Homo sapiens OX=9606 GN=MADD PE=1 SV=2 |
| CPNE2_MOUSE | Copine-2 OS=Mus musculus OX=10090 GN=Cpne2 PE=1 SV=1 |
| PSMD9_MOUSE | 26S proteasome non-ATPase regulatory subunit 9 OS=Mus musculus OX=10090 GN=Psmd9 PE=1 SV=1 |
| VAT1L_MOUSE | Synaptic vesicle membrane protein VAT-1 homolog-like OS=Mus musculus OX=10090 GN=Vat1l PE=1 SV=2 |
| EXOC4_MOUSE | Exocyst complex component 4 OS=Mus musculus OX=10090 GN=Exoc4 PE=1 SV=2 |
| PARP1_MOUSE | Poly [ADP-ribose] polymerase 1 OS=Mus musculus OX=10090 GN=Parp1 PE=1 SV=3 |
| TBC24_MOUSE | TBC1 domain family member 24 OS=Mus musculus OX=10090 GN=Tbc1d24 PE=1 SV=2 |
| CUL1_MOUSE | Cullin-1 OS=Mus musculus OX=10090 GN=Cul1 PE=1 SV=1 |
| AOXA_MOUSE | Aldehyde oxidase 1 OS=Mus musculus OX=10090 GN=Aox1 PE=1 SV=2 |
| KIF3B_MOUSE | Kinesin-like protein KIF3B OS=Mus musculus OX=10090 GN=Kif3b PE=1 SV=1 |

| | |
|---|---|
| PPCEL_MOUSE | Prolyl endopeptidase-like OS=Mus musculus OX=10090 GN=Prepl PE=1 SV=1 |
| DCTN4_MOUSE | Dynactin subunit 4 OS=Mus musculus OX=10090 GN=Dctn4 PE=1 SV=1 |
| APOB_MOUSE | Apolipoprotein B-100 OS=Mus musculus OX=10090 GN=Apob PE=1 SV=1 |
| DUS3_MOUSE | Dual specificity protein phosphatase 3 OS=Mus musculus OX=10090 GN=Dusp3 PE=1 SV=1 |
| RL35_MOUSE | 60S ribosomal protein L35 OS=Mus musculus OX=10090 GN=Rpl35 PE=1 SV=1 |
| ZN207_MOUSE | BUB3-interacting and GLEBS motif-containing protein ZNF207 OS=Mus musculus OX=10090 GN=Znf207 PE=1 SV=1 |
| ATPMD_MOUSE | ATP synthase membrane subunit DAPIT, mitochondrial OS=Mus musculus OX=10090 GN=Atp5md PE=1 SV=1 |
| MIC25_MOUSE | MICOS complex subunit Mic25 OS=Mus musculus OX=10090 GN=Chchd6 PE=1 SV=2 |
| MAGD1_MOUSE | Melanoma-associated antigen D1 OS=Mus musculus OX=10090 GN=Maged1 PE=1 SV=1 |
| PRAF2_MOUSE | PRA1 family protein 2 OS=Mus musculus OX=10090 GN=Praf2 PE=1 SV=1 |
| AP3S1_MOUSE | AP-3 complex subunit sigma-1 OS=Mus musculus OX=10090 GN=Ap3s1 PE=1 SV=2 |
| HVM18_MOUSE | Ig heavy chain V regions TEPC 15/S107/HPCM1/HPCM2/HPCM3 OS=Mus musculus OX=10090 PE=1 SV=1 |
| MBOA7_MOUSE | Lysophospholipid acyltransferase 7 OS=Mus musculus OX=10090 GN=Mboat7 PE=1 SV=1 |
| STAM1_MOUSE | Signal transducing adapter molecule 1 OS=Mus musculus OX=10090 GN=Stam PE=1 SV=3 |
| CON__Q86YZ3 | CON__Q86YZ3 |
| KPCB_MOUSE | Protein kinase C beta type OS=Mus musculus OX=10090 GN=Prkcb PE=1 SV=4 |
| KCAB2_MOUSE | Voltage-gated potassium channel subunit beta-2 OS=Mus musculus OX=10090 GN=Kcnab2 PE=1 SV=1 |
| SQSTM_MOUSE | Sequestosome-1 OS=Mus musculus OX=10090 GN=Sqstm1 PE=1 SV=1 |
| S38A3_MOUSE | Sodium-coupled neutral amino acid transporter 3 OS=Mus musculus OX=10090 GN=Slc38a3 PE=1 SV=1 |

| | |
|---|---|
| M4K4_MOUSE | Mitogen-activated protein kinase kinase kinase kinase 4 OS=Mus musculus OX=10090 GN=Map4k4 PE=1 SV=1 |
| ARF6_MOUSE | ADP-ribosylation factor 6 OS=Mus musculus OX=10090 GN=Arf6 PE=1 SV=2 |
| SLTM_MOUSE | SAFB-like transcription modulator OS=Mus musculus OX=10090 GN=Sltm PE=1 SV=1 |
| DPY30_MOUSE | Protein dpy-30 homolog OS=Mus musculus OX=10090 GN=Dpy30 PE=1 SV=1 |
| UMPS_MOUSE | Uridine 5'-monophosphate synthase OS=Mus musculus OX=10090 GN=Umps PE=1 SV=3 |
| MUTA_MOUSE | Methylmalonyl-CoA mutase, mitochondrial OS=Mus musculus OX=10090 GN=Mmut PE=1 SV=2 |
| RT29_MOUSE | 28S ribosomal protein S29, mitochondrial OS=Mus musculus OX=10090 GN=Dap3 PE=1 SV=1 |
| MFN2_MOUSE | Mitofusin-2 OS=Mus musculus OX=10090 GN=Mfn2 PE=1 SV=3 |
| ITM2B_MOUSE | Integral membrane protein 2B OS=Mus musculus OX=10090 GN=Itm2b PE=1 SV=1 |
| EPN1_MOUSE | Epsin-1 OS=Mus musculus OX=10090 GN=Epn1 PE=1 SV=3 |
| DDX21_MOUSE | Nucleolar RNA helicase 2 OS=Mus musculus OX=10090 GN=Ddx21 PE=1 SV=3 |
| CSN5_MOUSE | COP9 signalosome complex subunit 5 OS=Mus musculus OX=10090 GN=Cops5 PE=1 SV=3 |
| GPX1_MOUSE | Glutathione peroxidase 1 OS=Mus musculus OX=10090 GN=Gpx1 PE=1 SV=2 |
| ENSA_MOUSE | Alpha-endosulfine OS=Mus musculus OX=10090 GN=Ensa PE=1 SV=1 |
| ACSL6_MOUSE | Long-chain-fatty-acid--CoA ligase 6 OS=Mus musculus OX=10090 GN=Acsl6 PE=1 SV=1 |
| THTM_MOUSE | 3-mercaptopyruvate sulfurtransferase OS=Mus musculus OX=10090 GN=Mpst PE=1 SV=4 |
| RBM3_MOUSE | RNA-binding protein 3 OS=Mus musculus OX=10090 GN=Rbm3 PE=1 SV=1 |
| HMOX2_MOUSE | Heme oxygenase 2 OS=Mus musculus OX=10090 GN=Hmox2 PE=1 SV=1 |
| KGUA_MOUSE | Guanylate kinase OS=Mus musculus OX=10090 GN=Guk1 PE=1 SV=2 |

| | |
|---|---|
| MAGI1_MOUSE | Membrane-associated guanylate kinase, WW and PDZ domain-containing protein 1 OS=Mus musculus OX=10090 GN=Magi1 PE=1 SV=1 |
| S6A17_MOUSE | Sodium-dependent neutral amino acid transporter SLC6A17 OS=Mus musculus OX=10090 GN=Slc6a17 PE=1 SV=1 |
| ARL2_MOUSE | ADP-ribosylation factor-like protein 2 OS=Mus musculus OX=10090 GN=Arl2 PE=1 SV=1 |
| VAPB_MOUSE | Vesicle-associated membrane protein-associated protein B OS=Mus musculus OX=10090 GN=Vapb PE=1 SV=3 |
| SART3_MOUSE | Squamous cell carcinoma antigen recognized by T-cells 3 OS=Mus musculus OX=10090 GN=Sart3 PE=1 SV=1 |
| BHMT1_MOUSE | Betaine--homocysteine S-methyltransferase 1 OS=Mus musculus OX=10090 GN=Bhmt PE=1 SV=1 |
| KPRB_MOUSE | Phosphoribosyl pyrophosphate synthase-associated protein 2 OS=Mus musculus OX=10090 GN=Prpsap2 PE=1 SV=1 |
| PPM1H_MOUSE | Protein phosphatase 1H OS=Mus musculus OX=10090 GN=Ppm1h PE=1 SV=1 |
| BAIP2_MOUSE | Brain-specific angiogenesis inhibitor 1-associated protein 2 OS=Mus musculus OX=10090 GN=Baiap2 PE=1 SV=2 |
| ACOT9_MOUSE | Acyl-coenzyme A thioesterase 9, mitochondrial OS=Mus musculus OX=10090 GN=Acot9 PE=1 SV=1 |
| RAB5A_MOUSE | Ras-related protein Rab-5A OS=Mus musculus OX=10090 GN=Rab5a PE=1 SV=1 |
| MEST_MOUSE | Mesoderm-specific transcript protein OS=Mus musculus OX=10090 GN=Mest PE=2 SV=1 |
| RCC1_MOUSE | Regulator of chromosome condensation OS=Mus musculus OX=10090 GN=Rcc1 PE=1 SV=1 |
| GOGA2_MOUSE | Golgin subfamily A member 2 OS=Mus musculus OX=10090 GN=Golga2 PE=1 SV=3 |
| LAMA5_MOUSE | Laminin subunit alpha-5 OS=Mus musculus OX=10090 GN=Lama5 PE=1 SV=4 |
| FABPH_MOUSE | Fatty acid-binding protein, heart OS=Mus musculus OX=10090 GN=Fabp3 PE=1 SV=5 |
| PO210_MOUSE | Nuclear pore membrane glycoprotein 210 OS=Mus musculus OX=10090 GN=Nup210 PE=1 SV=2 |

| | |
|---|---|
| CLPP_MOUSE | ATP-dependent Clp protease proteolytic subunit, mitochondrial OS=Mus musculus OX=10090 GN=Clpp PE=1 SV=1 |
| BROX_MOUSE | BRO1 domain-containing protein BROX OS=Mus musculus OX=10090 GN=Brox PE=1 SV=1 |
| CHM4B_MOUSE | Charged multivesicular body protein 4b OS=Mus musculus OX=10090 GN=Chmp4b PE=1 SV=2 |
| P4HA1_MOUSE | Prolyl 4-hydroxylase subunit alpha-1 OS=Mus musculus OX=10090 GN=P4ha1 PE=1 SV=2 |
| MT2_MOUSE | Metallothionein-2 OS=Mus musculus OX=10090 GN=Mt2 PE=1 SV=2 |
| IQGA1_MOUSE | Ras GTPase-activating-like protein IQGAP1 OS=Mus musculus OX=10090 GN=Iqgap1 PE=1 SV=2 |
| WDR48_MOUSE | WD repeat-containing protein 48 OS=Mus musculus OX=10090 GN=Wdr48 PE=1 SV=1 |
| Q27IK6-DECOY | Q27IK6 |
| MCM6_MOUSE | DNA replication licensing factor MCM6 OS=Mus musculus OX=10090 GN=Mcm6 PE=1 SV=1 |
| SIAS_MOUSE | Sialic acid synthase OS=Mus musculus OX=10090 GN=Nans PE=1 SV=1 |
| CTND1_MOUSE | Catenin delta-1 OS=Mus musculus OX=10090 GN=Ctnnd1 PE=1 SV=2 |
| NPS3B_MOUSE | Protein NipSnap homolog 3B OS=Mus musculus OX=10090 GN=Nipsnap3b PE=1 SV=1 |
| CSK_MOUSE | Tyrosine-protein kinase CSK OS=Mus musculus OX=10090 GN=Csk PE=1 SV=2 |
| VMA5A_MOUSE | von Willebrand factor A domain-containing protein 5A OS=Mus musculus OX=10090 GN=Vwa5a PE=1 SV=2 |
| ISOC1_MOUSE | Isochorismatase domain-containing protein 1 OS=Mus musculus OX=10090 GN=Isoc1 PE=1 SV=1 |
| C1QT4_MOUSE | Complement C1q tumor necrosis factor-related protein 4 OS=Mus musculus OX=10090 GN=C1qtnf4 PE=1 SV=1 |
| RBM12_MOUSE | RNA-binding protein 12 OS=Mus musculus OX=10090 GN=Rbm12 PE=1 SV=3 |
| CDV3_MOUSE | Protein CDV3 OS=Mus musculus OX=10090 GN=Cdv3 PE=1 SV=2 |

| | |
|---|---|
| MPPA_MOUSE | Mitochondrial-processing peptidase subunit alpha OS=Mus musculus OX=10090 GN=Pmpca PE=1 SV=1 |
| CTTB2_MOUSE | Cortactin-binding protein 2 OS=Mus musculus OX=10090 GN=Cttnbp2 PE=1 SV=2 |
| PGP_MOUSE | Glycerol-3-phosphate phosphatase OS=Mus musculus OX=10090 GN=Pgp PE=1 SV=1 |
| GNL1_MOUSE | Guanine nucleotide-binding protein-like 1 OS=Mus musculus OX=10090 GN=Gnl1 PE=1 SV=4 |
| ACAP2_MOUSE | Arf-GAP with coiled-coil, ANK repeat and PH domain-containing protein 2 OS=Mus musculus OX=10090 GN=Acap2 PE=1 SV=2 |
| Q80VH0-DECOY | Q80VH0 |
| FLOT2_MOUSE | Flotillin-2 OS=Mus musculus OX=10090 GN=Flot2 PE=1 SV=2 |
| PAK2_MOUSE | Serine/threonine-protein kinase PAK 2 OS=Mus musculus OX=10090 GN=Pak2 PE=1 SV=1 |
| RN213_MOUSE | E3 ubiquitin-protein ligase RNF213 OS=Mus musculus OX=10090 GN=Rnf213 PE=1 SV=2 |
| F210B_MOUSE | Protein FAM210B, mitochondrial OS=Mus musculus OX=10090 GN=Fam210b PE=1 SV=3 |
| COQ9_MOUSE | Ubiquinone biosynthesis protein COQ9, mitochondrial OS=Mus musculus OX=10090 GN=Coq9 PE=1 SV=1 |
| RANB3_MOUSE | Ran-binding protein 3 OS=Mus musculus OX=10090 GN=Ranbp3 PE=1 SV=2 |
| NDUAB_MOUSE | NADH dehydrogenase [ubiquinone] 1 alpha subcomplex subunit 11 OS=Mus musculus OX=10090 GN=Ndufa11 PE=1 SV=2 |
| LIN7C_MOUSE | Protein lin-7 homolog C OS=Mus musculus OX=10090 GN=Lin7c PE=1 SV=2 |
| ACO13_MOUSE | Acyl-coenzyme A thioesterase 13 OS=Mus musculus OX=10090 GN=Acot13 PE=1 SV=1 |
| HAP28_MOUSE | 28 kDa heat- and acid-stable phosphoprotein OS=Mus musculus OX=10090 GN=Pdap1 PE=1 SV=1 |
| KV2A7_MOUSE | Ig kappa chain V-II region 26-10 OS=Mus musculus OX=10090 PE=1 SV=1 |
| IMA3_MOUSE | Importin subunit alpha-3 OS=Mus musculus OX=10090 GN=Kpna4 PE=1 SV=1 |

| | |
|---|---|
| ATP8_MOUSE | ATP synthase protein 8 OS=Mus musculus OX=10090 GN=Mtatp8 PE=1 SV=1 |
| SUMO2_MOUSE | Small ubiquitin-related modifier 2 OS=Mus musculus OX=10090 GN=Sumo2 PE=1 SV=1 |
| MOT1_MOUSE | Monocarboxylate transporter 1 OS=Mus musculus OX=10090 GN=Slc16a1 PE=1 SV=1 |
| NAMPT_MOUSE | Nicotinamide phosphoribosyltransferase OS=Mus musculus OX=10090 GN=Nampt PE=1 SV=1 |
| MXRA7_MOUSE | Matrix-remodeling-associated protein 7 OS=Mus musculus OX=10090 GN=Mxra7 PE=1 SV=2 |
| GMPPA_MOUSE | Mannose-1-phosphate guanyltransferase alpha OS=Mus musculus OX=10090 GN=Gmppa PE=1 SV=1 |
| PCDG4_MOUSE | Protocadherin gamma-A4 OS=Mus musculus OX=10090 GN=Pcdhga4 PE=1 SV=1 |
| OCAD1_MOUSE | OCIA domain-containing protein 1 OS=Mus musculus OX=10090 GN=Ociad1 PE=1 SV=1 |
| FMR1_MOUSE | Synaptic functional regulator FMR1 OS=Mus musculus OX=10090 GN=Fmr1 PE=1 SV=1 |
| NHRF1_MOUSE | Na(+)/H(+) exchange regulatory cofactor NHE-RF1 OS=Mus musculus OX=10090 GN=Slc9a3r1 PE=1 SV=3 |
| SRSF4_MOUSE | Serine/arginine-rich splicing factor 4 OS=Mus musculus OX=10090 GN=Srsf4 PE=2 SV=1 |
| HSBP1_MOUSE | Heat shock factor-binding protein 1 OS=Mus musculus OX=10090 GN=Hsbp1 PE=1 SV=1 |
| TPPC3_MOUSE | Trafficking protein particle complex subunit 3 OS=Mus musculus OX=10090 GN=Trappc3 PE=1 SV=1 |
| ICAM1_MOUSE | Intercellular adhesion molecule 1 OS=Mus musculus OX=10090 GN=Icam1 PE=1 SV=1 |
| NET1B_ARATH | Protein NETWORKED 1B OS=Arabidopsis thaliana OX=3702 GN=NET1B PE=2 SV=1 |
| CASC4_MOUSE | Protein CASC4 OS=Mus musculus OX=10090 GN=Casc4 PE=1 SV=1 |
| THUM1_MOUSE | THUMP domain-containing protein 1 OS=Mus musculus OX=10090 GN=Thumpd1 PE=1 SV=1 |
| GMPR1_MOUSE | GMP reductase 1 OS=Mus musculus OX=10090 GN=Gmpr PE=1 SV=1 |
| COX3_MOUSE | Cytochrome c oxidase subunit 3 OS=Mus musculus OX=10090 GN=mt-Co3 PE=1 SV=2 |

| | |
|---|---|
| NU1M_MOUSE | NADH-ubiquinone oxidoreductase chain 1 OS=Mus musculus OX=10090 GN=Mtnd1 PE=1 SV=3 |
| NUD10_MOUSE | Diphosphoinositol polyphosphate phosphohydrolase 3-alpha OS=Mus musculus OX=10090 GN=Nudt10 PE=1 SV=1 |
| RHEB_MOUSE | GTP-binding protein Rheb OS=Mus musculus OX=10090 GN=Rheb PE=1 SV=1 |
| MPU1_MOUSE | Mannose-P-dolichol utilization defect 1 protein OS=Mus musculus OX=10090 GN=Mpdu1 PE=1 SV=1 |
| NCKX4_MOUSE | Sodium/potassium/calcium exchanger 4 OS=Mus musculus OX=10090 GN=Slc24a4 PE=1 SV=2 |
| ENDD1_MOUSE | Endonuclease domain-containing 1 protein OS=Mus musculus OX=10090 GN=Endod1 PE=1 SV=2 |
| STRN3_MOUSE | Striatin-3 OS=Mus musculus OX=10090 GN=Strn3 PE=1 SV=1 |
| DHX30_MOUSE | ATP-dependent RNA helicase DHX30 OS=Mus musculus OX=10090 GN=Dhx30 PE=1 SV=1 |
| KISHA_MOUSE | Protein kish-A OS=Mus musculus OX=10090 GN=Tmem167a PE=1 SV=1 |
| PBX1_MOUSE | Pre-B-cell leukemia transcription factor 1 OS=Mus musculus OX=10090 GN=Pbx1 PE=1 SV=2 |
| HGS_MOUSE | Hepatocyte growth factor-regulated tyrosine kinase substrate OS=Mus musculus OX=10090 GN=Hgs PE=1 SV=2 |
| KV5A2_MOUSE | Ig kappa chain V-V region MOPC 21 OS=Mus musculus OX=10090 PE=1 SV=1 |
| CA2D3_MOUSE | Voltage-dependent calcium channel subunit alpha-2/delta-3 OS=Mus musculus OX=10090 GN=Cacna2d3 PE=1 SV=1 |
| Q8VHE6-DECOY | Q8VHE6 |
| CFAH_MOUSE | Complement factor H OS=Mus musculus OX=10090 GN=Cfh PE=1 SV=2 |
| APLP2_MOUSE | Amyloid-like protein 2 OS=Mus musculus OX=10090 GN=Aplp2 PE=1 SV=4 |
| MAT2B_MOUSE | Methionine adenosyltransferase 2 subunit beta OS=Mus musculus OX=10090 GN=Mat2b PE=1 SV=1 |
| CTBP2_MOUSE | C-terminal-binding protein 2 OS=Mus musculus OX=10090 GN=Ctbp2 PE=1 SV=2 |

| | |
|---|---|
| GBRA5_MOUSE | Gamma-aminobutyric acid receptor subunit alpha-5 OS=Mus musculus OX=10090 GN=Gabra5 PE=1 SV=1 |
| ICAM5_MOUSE | Intercellular adhesion molecule 5 OS=Mus musculus OX=10090 GN=Icam5 PE=1 SV=2 |
| NEGR1_MOUSE | Neuronal growth regulator 1 OS=Mus musculus OX=10090 GN=Negr1 PE=1 SV=1 |
| RT05_MOUSE | 28S ribosomal protein S5, mitochondrial OS=Mus musculus OX=10090 GN=Mrps5 PE=1 SV=1 |
| DDX46_MOUSE | Probable ATP-dependent RNA helicase DDX46 OS=Mus musculus OX=10090 GN=Ddx46 PE=1 SV=2 |
| RGS7_MOUSE | Regulator of G-protein signaling 7 OS=Mus musculus OX=10090 GN=Rgs7 PE=1 SV=2 |
| IMA7_MOUSE | Importin subunit alpha-7 OS=Mus musculus OX=10090 GN=Kpna6 PE=1 SV=2 |
| FXR2_MOUSE | Fragile X mental retardation syndrome-related protein 2 OS=Mus musculus OX=10090 GN=Fxr2 PE=1 SV=1 |
| MTA2_MOUSE | Metastasis-associated protein MTA2 OS=Mus musculus OX=10090 GN=Mta2 PE=1 SV=1 |
| CPNE8_MOUSE | Copine-8 OS=Mus musculus OX=10090 GN=Cpne8 PE=2 SV=3 |
| CRNL1_MOUSE | Crooked neck-like protein 1 OS=Mus musculus OX=10090 GN=Crnkl1 PE=1 SV=1 |
| EFGM_MOUSE | Elongation factor G, mitochondrial OS=Mus musculus OX=10090 GN=Gfm1 PE=1 SV=1 |
| LYAG_MOUSE | Lysosomal alpha-glucosidase OS=Mus musculus OX=10090 GN=Gaa PE=1 SV=2 |
| SAP_MOUSE | Prosaposin OS=Mus musculus OX=10090 GN=Psap PE=1 SV=2 |
| STK39_MOUSE | STE20/SPS1-related proline-alanine-rich protein kinase OS=Mus musculus OX=10090 GN=Stk39 PE=1 SV=1 |
| ARMC6_MOUSE | Armadillo repeat-containing protein 6 OS=Mus musculus OX=10090 GN=Armc6 PE=1 SV=1 |
| NCOA5_MOUSE | Nuclear receptor coactivator 5 OS=Mus musculus OX=10090 GN=Ncoa5 PE=1 SV=1 |
| ECM29_MOUSE | Proteasome adapter and scaffold protein ECM29 OS=Mus musculus OX=10090 GN=Ecpas PE=1 SV=3 |

| | |
|---|---|
| MCTS1_MOUSE | Malignant T-cell-amplified sequence 1 OS=Mus musculus OX=10090 GN=Mcts1 PE=1 SV=1 |
| ITA6_MOUSE | Integrin alpha-6 OS=Mus musculus OX=10090 GN=Itga6 PE=1 SV=3 |
| SEPT8_MOUSE | Septin-8 OS=Mus musculus OX=10090 GN=Septin8 PE=1 SV=4 |
| HIP1R_MOUSE | Huntingtin-interacting protein 1-related protein OS=Mus musculus OX=10090 GN=Hip1r PE=1 SV=2 |
| PUM2_MOUSE | Pumilio homolog 2 OS=Mus musculus OX=10090 GN=Pum2 PE=1 SV=2 |
| CE170_MOUSE | Centrosomal protein of 170 kDa OS=Mus musculus OX=10090 GN=Cep170 PE=1 SV=2 |
| AT2B4_MOUSE | Plasma membrane calcium-transporting ATPase 4 OS=Mus musculus OX=10090 GN=Atp2b4 PE=1 SV=1 |
| BSN_MOUSE | Protein bassoon OS=Mus musculus OX=10090 GN=Bsn PE=1 SV=4 |
| SYTC2_MOUSE | Threonine--tRNA ligase 2, cytoplasmic OS=Mus musculus OX=10090 GN=Tarsl2 PE=1 SV=1 |
| CUL4A_MOUSE | Cullin-4A OS=Mus musculus OX=10090 GN=Cul4a PE=1 SV=1 |
| UBP47_MOUSE | Ubiquitin carboxyl-terminal hydrolase 47 OS=Mus musculus OX=10090 GN=Usp47 PE=1 SV=2 |
| EHD3_MOUSE | EH domain-containing protein 3 OS=Mus musculus OX=10090 GN=Ehd3 PE=1 SV=2 |
| DCAF7_MOUSE | DDB1- and CUL4-associated factor 7 OS=Mus musculus OX=10090 GN=Dcaf7 PE=1 SV=1 |
| S27A4_MOUSE | Long-chain fatty acid transport protein 4 OS=Mus musculus OX=10090 GN=Slc27a4 PE=1 SV=1 |
| CLIP1_MOUSE | CAP-Gly domain-containing linker protein 1 OS=Mus musculus OX=10090 GN=Clip1 PE=1 SV=1 |
| RAB23_MOUSE | Ras-related protein Rab-23 OS=Mus musculus OX=10090 GN=Rab23 PE=1 SV=2 |
| MRCKB_MOUSE | Serine/threonine-protein kinase MRCK beta OS=Mus musculus OX=10090 GN=Cdc42bpb PE=1 SV=2 |
| SH3G1_MOUSE | Endophilin-A2 OS=Mus musculus OX=10090 GN=Sh3gl1 PE=1 SV=1 |

| | |
|---|---|
| NF1_MOUSE | Neurofibromin OS=Mus musculus OX=10090 GN=Nf1 PE=1 SV=1 |
| PGAM5_MOUSE | Serine/threonine-protein phosphatase PGAM5, mitochondrial OS=Mus musculus OX=10090 GN=Pgam5 PE=1 SV=1 |
| RUXF_MOUSE | Small nuclear ribonucleoprotein F OS=Mus musculus OX=10090 GN=Snrpf PE=1 SV=1 |
| PHF24_MOUSE | PHD finger protein 24 OS=Mus musculus OX=10090 GN=Phf24 PE=1 SV=2 |
| SRR_MOUSE | Serine racemase OS=Mus musculus OX=10090 GN=Srr PE=1 SV=1 |
| NDUS4_MOUSE | NADH dehydrogenase [ubiquinone] iron-sulfur protein 4, mitochondrial OS=Mus musculus OX=10090 GN=Ndufs4 PE=1 SV=3 |
| TMX1_MOUSE | Thioredoxin-related transmembrane protein 1 OS=Mus musculus OX=10090 GN=Tmx1 PE=1 SV=1 |
| GAS7_MOUSE | Growth arrest-specific protein 7 OS=Mus musculus OX=10090 GN=Gas7 PE=1 SV=1 |
| KV5A7_MOUSE | Ig kappa chain V-V region MOPC 41 OS=Mus musculus OX=10090 GN=Gm5571 PE=1 SV=1 |
| ECI1_MOUSE | Enoyl-CoA delta isomerase 1, mitochondrial OS=Mus musculus OX=10090 GN=Eci1 PE=1 SV=2 |
| NUDC2_MOUSE | NudC domain-containing protein 2 OS=Mus musculus OX=10090 GN=Nudcd2 PE=1 SV=1 |
| SPRE_MOUSE | Sepiapterin reductase OS=Mus musculus OX=10090 GN=Spr PE=1 SV=1 |
| AL1A7_MOUSE | Aldehyde dehydrogenase, cytosolic 1 OS=Mus musculus OX=10090 GN=Aldh1a7 PE=1 SV=1 |
| PLPR2_MOUSE | Phospholipid phosphatase-related protein type 2 OS=Mus musculus OX=10090 GN=Plppr2 PE=1 SV=1 |
| S39A6_MOUSE | Zinc transporter ZIP6 OS=Mus musculus OX=10090 GN=Slc39a6 PE=1 SV=1 |
| KV3AM_MOUSE | Ig kappa chain V-III region PC 2154 OS=Mus musculus OX=10090 PE=1 SV=1 |
| PC4L1_MOUSE | Purkinje cell protein 4-like protein 1 OS=Mus musculus OX=10090 GN=Pcp4l1 PE=1 SV=1 |
| GBG3_MOUSE | Guanine nucleotide-binding protein G(I)/G(S)/G(O) subunit gamma-3 OS=Mus musculus OX=10090 GN=Gng3 PE=1 SV=1 |

| | |
|---|---|
| TBA3_ARATH | Tubulin alpha-3 chain OS=Arabidopsis thaliana OX=3702 GN=TUBA3 PE=1 SV=2 |
| CON__P12763 | CON__P12763 |
| RUXE_MOUSE | Small nuclear ribonucleoprotein E OS=Mus musculus OX=10090 GN=Snrpe PE=1 SV=1 |
| O82190-DECOY | O82190 |
| PFD2_MOUSE | Prefoldin subunit 2 OS=Mus musculus OX=10090 GN=Pfdn2 PE=1 SV=2 |
| DHB8_MOUSE | Estradiol 17-beta-dehydrogenase 8 OS=Mus musculus OX=10090 GN=Hsd17b8 PE=1 SV=2 |
| GHC2_MOUSE | Mitochondrial glutamate carrier 2 OS=Mus musculus OX=10090 GN=Slc25a18 PE=1 SV=4 |
| ABRAL_MOUSE | Costars family protein ABRACL OS=Mus musculus OX=10090 GN=Abracl PE=1 SV=1 |
| PK1IP_MOUSE | p21-activated protein kinase-interacting protein 1 OS=Mus musculus OX=10090 GN=Pak1ip1 PE=1 SV=2 |
| RT07_MOUSE | 28S ribosomal protein S7, mitochondrial OS=Mus musculus OX=10090 GN=Mrps7 PE=1 SV=1 |
| NSMA2_MOUSE | Sphingomyelin phosphodiesterase 3 OS=Mus musculus OX=10090 GN=Smpd3 PE=1 SV=1 |
| HBA_MOUSE | Hemoglobin subunit alpha OS=Mus musculus OX=10090 GN=Hba PE=1 SV=2 |
| CTNA1_MOUSE | Catenin alpha-1 OS=Mus musculus OX=10090 GN=Ctnna1 PE=1 SV=1 |
| CCG8_MOUSE | Voltage-dependent calcium channel gamma-8 subunit OS=Mus musculus OX=10090 GN=Cacng8 PE=1 SV=1 |
| LANC1_MOUSE | Glutathione S-transferase LANCL1 OS=Mus musculus OX=10090 GN=Lancl1 PE=1 SV=1 |
| FADS2_MOUSE | Acyl-CoA 6-desaturase OS=Mus musculus OX=10090 GN=Fads2 PE=1 SV=1 |
| PCP4_HUMAN | Calmodulin regulator protein PCP4 OS=Homo sapiens OX=9606 GN=PCP4 PE=1 SV=3 |
| GSK3A_MOUSE | Glycogen synthase kinase-3 alpha OS=Mus musculus OX=10090 GN=Gsk3a PE=1 SV=2 |
| TNPO1_MOUSE | Transportin-1 OS=Mus musculus OX=10090 GN=Tnpo1 PE=1 SV=2 |
| PDLI1_MOUSE | PDZ and LIM domain protein 1 OS=Mus musculus OX=10090 GN=Pdlim1 PE=1 SV=4 |
| PR38B_MOUSE | Pre-mRNA-splicing factor 38B OS=Mus musculus OX=10090 GN=Prpf38b PE=1 SV=1 |

| | |
|---|---|
| SRP14_MOUSE | Signal recognition particle 14 kDa protein OS=Mus musculus OX=10090 GN=Srp14 PE=1 SV=1 |
| ACSL3_MOUSE | Long-chain-fatty-acid--CoA ligase 3 OS=Mus musculus OX=10090 GN=Acsl3 PE=1 SV=2 |
| KV6A6_MOUSE | Ig kappa chain V-VI region NQ2-17.4.1 OS=Mus musculus OX=10090 PE=2 SV=1 |
| PITM1_MOUSE | Membrane-associated phosphatidylinositol transfer protein 1 OS=Mus musculus OX=10090 GN=Pitpnm1 PE=1 SV=1 |
| HDGR2_MOUSE | Hepatoma-derived growth factor-related protein 2 OS=Mus musculus OX=10090 GN=Hdgfl2 PE=1 SV=1 |
| AP4A_MOUSE | Bis(5'-nucleosyl)-tetraphosphatase [asymmetrical] OS=Mus musculus OX=10090 GN=Nudt2 PE=1 SV=3 |
| LOXL2_HUMAN | Lysyl oxidase homolog 2 OS=Homo sapiens OX=9606 GN=LOXL2 PE=1 SV=1 |
| VPS18_MOUSE | Vacuolar protein sorting-associated protein 18 homolog OS=Mus musculus OX=10090 GN=Vps18 PE=1 SV=2 |
| TFAM_MOUSE | Transcription factor A, mitochondrial OS=Mus musculus OX=10090 GN=Tfam PE=1 SV=2 |
| S10A6_MOUSE | Protein S100-A6 OS=Mus musculus OX=10090 GN=S100a6 PE=1 SV=3 |
| PICAL_MOUSE | Phosphatidylinositol-binding clathrin assembly protein OS=Mus musculus OX=10090 GN=Picalm PE=1 SV=1 |
| SF3B4_MOUSE | Splicing factor 3B subunit 4 OS=Mus musculus OX=10090 GN=Sf3b4 PE=1 SV=1 |
| BAX_MOUSE | Apoptosis regulator BAX OS=Mus musculus OX=10090 GN=Bax PE=1 SV=1 |
| INP4A_MOUSE | Inositol polyphosphate-4-phosphatase type I A OS=Mus musculus OX=10090 GN=Inpp4a PE=1 SV=1 |
| CS1A_MOUSE | Complement C1s-A subcomponent OS=Mus musculus OX=10090 GN=C1sa PE=2 SV=2 |
| ISLR2_MOUSE | Immunoglobulin superfamily containing leucine-rich repeat protein 2 OS=Mus musculus OX=10090 GN=Islr2 PE=1 SV=1 |
| GPC2_MOUSE | Glypican-2 OS=Mus musculus OX=10090 GN=Gpc2 PE=2 SV=1 |

| | |
|---|---|
| HECAM_MOUSE | Hepatocyte cell adhesion molecule OS=Mus musculus OX=10090 GN=Hepacam PE=1 SV=2 |
| DNAK_MYCTU | Chaperone protein DnaK OS=Mycobacterium tuberculosis (strain ATCC 25618 / H37Rv) OX=83332 GN=dnaK PE=1 SV=1 |
| TTC3_MOUSE | E3 ubiquitin-protein ligase TTC3 OS=Mus musculus OX=10090 GN=Ttc3 PE=1 SV=2 |
| AGO2_MOUSE | Protein argonaute-2 OS=Mus musculus OX=10090 GN=Ago2 PE=1 SV=3 |
| KIFA3_MOUSE | Kinesin-associated protein 3 OS=Mus musculus OX=10090 GN=Kifap3 PE=1 SV=1 |
| CPLX1_MOUSE | Complexin-1 OS=Mus musculus OX=10090 GN=Cplx1 PE=1 SV=1 |
| GIT1_MOUSE | ARF GTPase-activating protein GIT1 OS=Mus musculus OX=10090 GN=Git1 PE=1 SV=1 |
| WDR6_MOUSE | WD repeat-containing protein 6 OS=Mus musculus OX=10090 GN=Wdr6 PE=1 SV=1 |
| UBFD1_MOUSE | Ubiquitin domain-containing protein UBFD1 OS=Mus musculus OX=10090 GN=Ubfd1 PE=1 SV=2 |
| IGSF8_MOUSE | Immunoglobulin superfamily member 8 OS=Mus musculus OX=10090 GN=Igsf8 PE=1 SV=2 |
| SMAP1_MOUSE | Stromal membrane-associated protein 1 OS=Mus musculus OX=10090 GN=Smap1 PE=1 SV=1 |
| PININ_MOUSE | Pinin OS=Mus musculus OX=10090 GN=Pnn PE=1 SV=4 |
| VTA1_MOUSE | Vacuolar protein sorting-associated protein VTA1 homolog OS=Mus musculus OX=10090 GN=Vta1 PE=1 SV=1 |
| MEP50_MOUSE | Methylosome protein 50 OS=Mus musculus OX=10090 GN=Wdr77 PE=1 SV=1 |
| LRRK1_MOUSE | Leucine-rich repeat serine/threonine-protein kinase 1 OS=Mus musculus OX=10090 GN=Lrrk1 PE=1 SV=1 |
| AL1A1_MOUSE | Retinal dehydrogenase 1 OS=Mus musculus OX=10090 GN=Aldh1a1 PE=1 SV=5 |
| ILK_MOUSE | Integrin-linked protein kinase OS=Mus musculus OX=10090 GN=Ilk PE=1 SV=2 |
| DLG2_MOUSE | Disks large homolog 2 OS=Mus musculus OX=10090 GN=Dlg2 PE=1 SV=2 |
| CYTB_MOUSE | Cystatin-B OS=Mus musculus OX=10090 GN=Cstb PE=1 SV=1 |

| | |
|---|---|
| BRCA2_MOUSE | Breast cancer type 2 susceptibility protein homolog OS=Mus musculus OX=10090 GN=Brca2 PE=1 SV=2 |
| IPYR2_MOUSE | Inorganic pyrophosphatase 2, mitochondrial OS=Mus musculus OX=10090 GN=Ppa2 PE=1 SV=1 |
| CPNE3_MOUSE | Copine-3 OS=Mus musculus OX=10090 GN=Cpne3 PE=1 SV=2 |
| HMGN2_MOUSE | Non-histone chromosomal protein HMG-17 OS=Mus musculus OX=10090 GN=Hmgn2 PE=1 SV=2 |
| TBAL3_MOUSE | Tubulin alpha chain-like 3 OS=Mus musculus OX=10090 GN=Tubal3 PE=2 SV=2 |
| S23IP_MOUSE | SEC23-interacting protein OS=Mus musculus OX=10090 GN=Sec23ip PE=1 SV=2 |
| KV2A6_MOUSE | Ig kappa chain V-II region 7S34.1 OS=Mus musculus OX=10090 PE=1 SV=1 |
| NDUB3_MOUSE | NADH dehydrogenase [ubiquinone] 1 beta subcomplex subunit 3 OS=Mus musculus OX=10090 GN=Ndufb3 PE=1 SV=1 |
| TCAL3_MOUSE | Transcription elongation factor A protein-like 3 OS=Mus musculus OX=10090 GN=Tceal3 PE=1 SV=2 |
| RAA1E_ARATH | Ras-related protein RABA1e OS=Arabidopsis thaliana OX=3702 GN=RABA1E PE=2 SV=1 |
| TIM14_MOUSE | Mitochondrial import inner membrane translocase subunit TIM14 OS=Mus musculus OX=10090 GN=Dnajc19 PE=1 SV=3 |
| VKORL_MOUSE | Vitamin K epoxide reductase complex subunit 1-like protein 1 OS=Mus musculus OX=10090 GN=Vkorc1l1 PE=1 SV=1 |
| CSN7A_MOUSE | COP9 signalosome complex subunit 7a OS=Mus musculus OX=10090 GN=Cops7a PE=1 SV=2 |
| VATB1_ARATH | V-type proton ATPase subunit B1 OS=Arabidopsis thaliana OX=3702 GN=VHA-B1 PE=2 SV=2 |
| B2L13_MOUSE | Bcl-2-like protein 13 OS=Mus musculus OX=10090 GN=Bcl2l13 PE=1 SV=2 |
| NCS1_MOUSE | Neuronal calcium sensor 1 OS=Mus musculus OX=10090 GN=Ncs1 PE=1 SV=3 |
| ICLN_MOUSE | Methylosome subunit pICln OS=Mus musculus OX=10090 GN=Clns1a PE=1 SV=1 |

| | |
|---|---|
| MK09_MOUSE | Mitogen-activated protein kinase 9 OS=Mus musculus OX=10090 GN=Mapk9 PE=1 SV=2 |
| CHMP5_MOUSE | Charged multivesicular body protein 5 OS=Mus musculus OX=10090 GN=Chmp5 PE=1 SV=1 |
| CON__Q3SX28 | CON__Q3SX28 |
| MT1_MOUSE | Metallothionein-1 OS=Mus musculus OX=10090 GN=Mt1 PE=1 SV=1 |
| TC132_ARATH | Translocase of chloroplast 132, chloroplastic OS=Arabidopsis thaliana OX=3702 GN=TOC132 PE=1 SV=1 |
| CACP_MOUSE | Carnitine O-acetyltransferase OS=Mus musculus OX=10090 GN=Crat PE=1 SV=3 |
| 2ABD_MOUSE | Serine/threonine-protein phosphatase 2A 55 kDa regulatory subunit B delta isoform OS=Mus musculus OX=10090 GN=Ppp2r2d PE=1 SV=1 |
| 1433S_MOUSE | 14-3-3 protein sigma OS=Mus musculus OX=10090 GN=Sfn PE=1 SV=2 |
| H31_HUMAN | Histone H3.1 OS=Homo sapiens OX=9606 GN=H3C1 PE=1 SV=2 |
| HPCL1_MOUSE | Hippocalcin-like protein 1 OS=Mus musculus OX=10090 GN=Hpcal1 PE=1 SV=2 |
| CPSF7_MOUSE | Cleavage and polyadenylation specificity factor subunit 7 OS=Mus musculus OX=10090 GN=Cpsf7 PE=1 SV=2 |
| GAK_MOUSE | Cyclin-G-associated kinase OS=Mus musculus OX=10090 GN=Gak PE=1 SV=2 |
| RBMX_MOUSE | RNA-binding motif protein, X chromosome OS=Mus musculus OX=10090 GN=Rbmx PE=1 SV=1 |
| GCDH_MOUSE | Glutaryl-CoA dehydrogenase, mitochondrial OS=Mus musculus OX=10090 GN=Gcdh PE=1 SV=2 |
| TCAF1_MOUSE | TRPM8 channel-associated factor 1 OS=Mus musculus OX=10090 GN=Tcaf1 PE=2 SV=1 |
| LEA29_ARATH | Late embryogenesis abundant protein 29 OS=Arabidopsis thaliana OX=3702 GN=LEA29 PE=2 SV=1 |
| MCA3_MOUSE | Eukaryotic translation elongation factor 1 epsilon-1 OS=Mus musculus OX=10090 GN=Eef1e1 PE=1 SV=1 |
| CROCC_MOUSE | Rootletin OS=Mus musculus OX=10090 GN=Crocc PE=1 SV=2 |

| | |
|---|---|
| PNPH_MOUSE | Purine nucleoside phosphorylase OS=Mus musculus OX=10090 GN=Pnp PE=1 SV=2 |
| RM04_MOUSE | 39S ribosomal protein L4, mitochondrial OS=Mus musculus OX=10090 GN=Mrpl4 PE=1 SV=1 |
| PK3CD_MOUSE | Phosphatidylinositol 4,5-bisphosphate 3-kinase catalytic subunit delta isoform OS=Mus musculus OX=10090 GN=Pik3cd PE=1 SV=2 |
| FKBP5_MOUSE | Peptidyl-prolyl cis-trans isomerase FKBP5 OS=Mus musculus OX=10090 GN=Fkbp5 PE=1 SV=1 |
| RAB8B_MOUSE | Ras-related protein Rab-8B OS=Mus musculus OX=10090 GN=Rab8b PE=1 SV=1 |
| GDE1_MOUSE | Glycerophosphodiester phosphodiesterase 1 OS=Mus musculus OX=10090 GN=Gde1 PE=1 SV=1 |
| ECSIT_MOUSE | Evolutionarily conserved signaling intermediate in Toll pathway, mitochondrial OS=Mus musculus OX=10090 GN=Ecsit PE=1 SV=2 |
| LBR_MOUSE | Delta(14)-sterol reductase LBR OS=Mus musculus OX=10090 GN=Lbr PE=1 SV=2 |
| CUL4B_MOUSE | Cullin-4B OS=Mus musculus OX=10090 GN=Cul4b PE=1 SV=1 |
| NIT2_MOUSE | Omega-amidase NIT2 OS=Mus musculus OX=10090 GN=Nit2 PE=1 SV=1 |
| TEFF2_MOUSE | Tomoregulin-2 OS=Mus musculus OX=10090 GN=Tmeff2 PE=2 SV=1 |
| GPI8_MOUSE | GPI-anchor transamidase OS=Mus musculus OX=10090 GN=Pigk PE=1 SV=2 |
| AFAD_MOUSE | Afadin OS=Mus musculus OX=10090 GN=Afdn PE=1 SV=3 |
| GPC1_MOUSE | Glypican-1 OS=Mus musculus OX=10090 GN=Gpc1 PE=1 SV=1 |
| NUDC3_MOUSE | NudC domain-containing protein 3 OS=Mus musculus OX=10090 GN=Nudcd3 PE=1 SV=3 |
| PCNA_MOUSE | Proliferating cell nuclear antigen OS=Mus musculus OX=10090 GN=Pcna PE=1 SV=2 |
| PFD6_MOUSE | Prefoldin subunit 6 OS=Mus musculus OX=10090 GN=Pfdn6 PE=1 SV=1 |
| MCM7_MOUSE | DNA replication licensing factor MCM7 OS=Mus musculus OX=10090 GN=Mcm7 PE=1 SV=1 |
| PFD5_MOUSE | Prefoldin subunit 5 OS=Mus musculus OX=10090 GN=Pfdn5 PE=1 SV=1 |

| | |
|---|---|
| RNZ2_MOUSE | Zinc phosphodiesterase ELAC protein 2 OS=Mus musculus OX=10090 GN=Elac2 PE=1 SV=1 |
| NAGK_MOUSE | N-acetyl-D-glucosamine kinase OS=Mus musculus OX=10090 GN=Nagk PE=1 SV=3 |
| CBARP_MOUSE | Voltage-dependent calcium channel beta subunit-associated regulatory protein OS=Mus musculus OX=10090 GN=Cbarp PE=1 SV=4 |
| TBCA_MOUSE | Tubulin-specific chaperone A OS=Mus musculus OX=10090 GN=Tbca PE=1 SV=3 |
| PRUN1_MOUSE | Exopolyphosphatase PRUNE1 OS=Mus musculus OX=10090 GN=Prune1 PE=1 SV=1 |
| BTF3_MOUSE | Transcription factor BTF3 OS=Mus musculus OX=10090 GN=Btf3 PE=1 SV=3 |
| KPCE_MOUSE | Protein kinase C epsilon type OS=Mus musculus OX=10090 GN=Prkce PE=1 SV=1 |
| IPO11_MOUSE | Importin-11 OS=Mus musculus OX=10090 GN=Ipo11 PE=1 SV=1 |
| PAPS1_MOUSE | Bifunctional 3'-phosphoadenosine 5'-phosphosulfate synthase 1 OS=Mus musculus OX=10090 GN=Papss1 PE=1 SV=1 |
| FAF2_MOUSE | FAS-associated factor 2 OS=Mus musculus OX=10090 GN=Faf2 PE=1 SV=2 |
| EMC8_MOUSE | ER membrane protein complex subunit 8 OS=Mus musculus OX=10090 GN=Emc8 PE=1 SV=1 |
| NT5C_MOUSE | 5'(3')-deoxyribonucleotidase, cytosolic type OS=Mus musculus OX=10090 GN=Nt5c PE=1 SV=1 |
| CTBL1_MOUSE | Beta-catenin-like protein 1 OS=Mus musculus OX=10090 GN=Ctnnbl1 PE=1 SV=1 |
| Q94FN2-DECOY | Q94FN2 |
| FLII_MOUSE | Protein flightless-1 homolog OS=Mus musculus OX=10090 GN=Flii PE=1 SV=1 |
| CON__Q32PI4 | CON__Q32PI4 |
| PSMG2_MOUSE | Proteasome assembly chaperone 2 OS=Mus musculus OX=10090 GN=Psmg2 PE=1 SV=1 |
| ABI2_MOUSE | Abl interactor 2 OS=Mus musculus OX=10090 GN=Abi2 PE=1 SV=1 |
| AGRL3_MOUSE | Adhesion G protein-coupled receptor L3 OS=Mus musculus OX=10090 GN=Adgrl3 PE=1 SV=3 |
| MAP1S_MOUSE | Microtubule-associated protein 1S OS=Mus musculus OX=10090 GN=Map1s PE=1 SV=2 |

| | |
|---|---|
| NMRL1_MOUSE | NmrA-like family domain-containing protein 1 OS=Mus musculus OX=10090 GN=Nmral1 PE=1 SV=1 |
| HM13_MOUSE | Minor histocompatibility antigen H13 OS=Mus musculus OX=10090 GN=Hm13 PE=1 SV=1 |
| DYH5_HUMAN | Dynein heavy chain 5, axonemal OS=Homo sapiens OX=9606 GN=DNAH5 PE=1 SV=3 |
| MICU1_MOUSE | Calcium uptake protein 1, mitochondrial OS=Mus musculus OX=10090 GN=Micu1 PE=1 SV=1 |
| P5CR3_MOUSE | Pyrroline-5-carboxylate reductase 3 OS=Mus musculus OX=10090 GN=Pycr3 PE=1 SV=2 |
| TDRKH_MOUSE | Tudor and KH domain-containing protein OS=Mus musculus OX=10090 GN=Tdrkh PE=1 SV=1 |
| RBGPR_MOUSE | Rab3 GTPase-activating protein non-catalytic subunit OS=Mus musculus OX=10090 GN=Rab3gap2 PE=1 SV=2 |
| DJC11_MOUSE | DnaJ homolog subfamily C member 11 OS=Mus musculus OX=10090 GN=Dnajc11 PE=1 SV=2 |
| OXSR1_MOUSE | Serine/threonine-protein kinase OSR1 OS=Mus musculus OX=10090 GN=Oxsr1 PE=1 SV=1 |
| S4A7_MOUSE | Sodium bicarbonate cotransporter 3 OS=Mus musculus OX=10090 GN=Slc4a7 PE=1 SV=2 |
| PP4P2_MOUSE | Type 2 phosphatidylinositol 4,5-bisphosphate 4-phosphatase OS=Mus musculus OX=10090 GN=Pip4p2 PE=1 SV=1 |
| MCM3_MOUSE | DNA replication licensing factor MCM3 OS=Mus musculus OX=10090 GN=Mcm3 PE=1 SV=2 |
| P85A_MOUSE | Phosphatidylinositol 3-kinase regulatory subunit alpha OS=Mus musculus OX=10090 GN=Pik3r1 PE=1 SV=2 |
| KIF15_MOUSE | Kinesin-like protein KIF15 OS=Mus musculus OX=10090 GN=Kif15 PE=1 SV=1 |
| BUD31_MOUSE | Protein BUD31 homolog OS=Mus musculus OX=10090 GN=Bud31 PE=1 SV=2 |
| C1QBP_MOUSE | Complement component 1 Q subcomponent-binding protein, mitochondrial OS=Mus musculus OX=10090 GN=C1qbp PE=1 SV=1 |
| HVM17_MOUSE | Ig heavy chain V region MOPC 47A OS=Mus musculus OX=10090 PE=1 SV=1 |
| HVM35_MOUSE | Ig heavy chain V-III region HPC76 (Fragment) OS=Mus musculus OX=10090 PE=4 SV=1 |

| | |
|---|---|
| HVM54_MOUSE | Ig heavy chain V region 5-84 OS=Mus musculus OX=10090 PE=1 SV=1 |
| HVM55_MOUSE | Ig heavy chain V region 345 OS=Mus musculus OX=10090 PE=1 SV=1 |
| KV4A1_MOUSE | Ig kappa chain V-IV region S107B OS=Mus musculus OX=10090 PE=4 SV=1 |
| KV5A4_MOUSE | Ig kappa chain V-V region MOPC 149 OS=Mus musculus OX=10090 PE=1 SV=1 |
| KV5A9_MOUSE | Ig kappa chain V-V region L7 (Fragment) OS=Mus musculus OX=10090 GN=Gm10881 PE=1 SV=1 |
| KV6AB_MOUSE | Ig kappa chain V-VI region NQ2-6.1 OS=Mus musculus OX=10090 PE=2 SV=1 |
| MGST1_MOUSE | Microsomal glutathione S-transferase 1 OS=Mus musculus OX=10090 GN=Mgst1 PE=1 SV=3 |
| NXF1_MOUSE | Nuclear RNA export factor 1 OS=Mus musculus OX=10090 GN=Nxf1 PE=1 SV=3 |
| RL36_MOUSE | 60S ribosomal protein L36 OS=Mus musculus OX=10090 GN=Rpl36 PE=1 SV=2 |
| SUMO1_MOUSE | Small ubiquitin-related modifier 1 OS=Mus musculus OX=10090 GN=Sumo1 PE=1 SV=1 |
| RS30_MOUSE | 40S ribosomal protein S30 OS=Mus musculus OX=10090 GN=Fau PE=1 SV=1 |
| TOM7_MOUSE | Mitochondrial import receptor subunit TOM7 homolog OS=Mus musculus OX=10090 GN=Tomm7 PE=3 SV=1 |
| KV2A5_MOUSE | Ig kappa chain V-II region 17S29.1 OS=Mus musculus OX=10090 PE=1 SV=1 |
| MYB73_ARATH | Transcription factor MYB73 OS=Arabidopsis thaliana OX=3702 GN=MYB73 PE=1 SV=1 |
| INV3_ARATH | Beta-fructofuranosidase, insoluble isoenzyme CWINV3 OS=Arabidopsis thaliana OX=3702 GN=CWINV3 PE=1 SV=2 |
| ARI1_HUMAN | E3 ubiquitin-protein ligase ARIH1 OS=Homo sapiens OX=9606 GN=ARIH1 PE=1 SV=2 |
| CON__Q7Z794 | CON__Q7Z794 |
| SP8_MOUSE | Transcription factor Sp8 OS=Mus musculus OX=10090 GN=Sp8 PE=2 SV=1 |
| INPP_MOUSE | Inositol polyphosphate 1-phosphatase OS=Mus musculus OX=10090 GN=Inpp1 PE=1 SV=2 |

| | |
|---|---|
| GALD1_MOUSE | Glutamine amidotransferase-like class 1 domain-containing protein 1 OS=Mus musculus OX=10090 GN=Gatd1 PE=1 SV=1 |
| ATE1_MOUSE | Arginyl-tRNA--protein transferase 1 OS=Mus musculus OX=10090 GN=Ate1 PE=1 SV=2 |
| SENP8_MOUSE | Sentrin-specific protease 8 OS=Mus musculus OX=10090 GN=Senp8 PE=1 SV=2 |
| ADDG_MOUSE | Gamma-adducin OS=Mus musculus OX=10090 GN=Add3 PE=1 SV=2 |
| Q9JMH9-DECOY | Q9JMH9 |
| SUN2_MOUSE | SUN domain-containing protein 2 OS=Mus musculus OX=10090 GN=Sun2 PE=1 SV=3 |
| ENAH_MOUSE | Protein enabled homolog OS=Mus musculus OX=10090 GN=Enah PE=1 SV=2 |
| NR4A2_MOUSE | Nuclear receptor subfamily 4 group A member 2 OS=Mus musculus OX=10090 GN=Nr4a2 PE=1 SV=1 |
| GNB5_MOUSE | Guanine nucleotide-binding protein subunit beta-5 OS=Mus musculus OX=10090 GN=Gnb5 PE=1 SV=1 |
| MK08_MOUSE | Mitogen-activated protein kinase 8 OS=Mus musculus OX=10090 GN=Mapk8 PE=1 SV=1 |
| CERS6_MOUSE | Ceramide synthase 6 OS=Mus musculus OX=10090 GN=Cers6 PE=1 SV=1 |
| IF6_MOUSE | Eukaryotic translation initiation factor 6 OS=Mus musculus OX=10090 GN=Eif6 PE=1 SV=2 |
| ARP5L_MOUSE | Actin-related protein 2/3 complex subunit 5-like protein OS=Mus musculus OX=10090 GN=Arpc5l PE=1 SV=1 |
| CBX1_MOUSE | Chromobox protein homolog 1 OS=Mus musculus OX=10090 GN=Cbx1 PE=1 SV=1 |
| WDR13_MOUSE | WD repeat-containing protein 13 OS=Mus musculus OX=10090 GN=Wdr13 PE=1 SV=1 |
| RUXG_MOUSE | Small nuclear ribonucleoprotein G OS=Mus musculus OX=10090 GN=Snrpg PE=1 SV=1 |
| TSYL4_MOUSE | Testis-specific Y-encoded-like protein 4 OS=Mus musculus OX=10090 GN=Tspyl4 PE=1 SV=1 |
| SNR40_MOUSE | U5 small nuclear ribonucleoprotein 40 kDa protein OS=Mus musculus OX=10090 GN=Snrnp40 PE=1 SV=1 |
| SMRD3_MOUSE | SWI/SNF-related matrix-associated actin-dependent regulator of chromatin subfamily D |

| | |
|---|---|
| | member 3 OS=Mus musculus OX=10090 GN=Smarcd3 PE=1 SV=2 |
| STAT3_MOUSE | Signal transducer and activator of transcription 3 OS=Mus musculus OX=10090 GN=Stat3 PE=1 SV=2 |
| ADAS_MOUSE | Alkyldihydroxyacetonephosphate synthase, peroxisomal OS=Mus musculus OX=10090 GN=Agps PE=1 SV=1 |
| P4HA3_MOUSE | Prolyl 4-hydroxylase subunit alpha-3 OS=Mus musculus OX=10090 GN=P4ha3 PE=2 SV=1 |
| X3CL1_MOUSE | Fractalkine OS=Mus musculus OX=10090 GN=Cx3cl1 PE=1 SV=3 |
| ATX2L_MOUSE | Ataxin-2-like protein OS=Mus musculus OX=10090 GN=Atxn2l PE=1 SV=1 |
| SHOT1_MOUSE | Shootin-1 OS=Mus musculus OX=10090 GN=Shtn1 PE=1 SV=1 |
| RM22_MOUSE | 39S ribosomal protein L22, mitochondrial OS=Mus musculus OX=10090 GN=Mrpl22 PE=1 SV=1 |
| CHERP_MOUSE | Calcium homeostasis endoplasmic reticulum protein OS=Mus musculus OX=10090 GN=Cherp PE=1 SV=1 |
| MEMO1_MOUSE | Protein MEMO1 OS=Mus musculus OX=10090 GN=Memo1 PE=1 SV=1 |
| MLF2_MOUSE | Myeloid leukemia factor 2 OS=Mus musculus OX=10090 GN=Mlf2 PE=1 SV=1 |
| I2BPL_MOUSE | Probable E3 ubiquitin-protein ligase IRF2BPL OS=Mus musculus OX=10090 GN=Irf2bpl PE=1 SV=1 |
| DKC1_MOUSE | H/ACA ribonucleoprotein complex subunit DKC1 OS=Mus musculus OX=10090 GN=Dkc1 PE=1 SV=4 |
| DNJC7_MOUSE | DnaJ homolog subfamily C member 7 OS=Mus musculus OX=10090 GN=Dnajc7 PE=1 SV=2 |
| ZN326_MOUSE | DBIRD complex subunit ZNF326 OS=Mus musculus OX=10090 GN=Znf326 PE=1 SV=1 |
| SHPS1_MOUSE | Tyrosine-protein phosphatase non-receptor type substrate 1 OS=Mus musculus OX=10090 GN=Sirpa PE=1 SV=2 |
| MBD3_MOUSE | Methyl-CpG-binding domain protein 3 OS=Mus musculus OX=10090 GN=Mbd3 PE=1 SV=1 |

| | |
|---|---|
| VPS52_MOUSE | Vacuolar protein sorting-associated protein 52 homolog OS=Mus musculus OX=10090 GN=Vps52 PE=1 SV=1 |
| SIR2_MOUSE | NAD-dependent protein deacetylase sirtuin-2 OS=Mus musculus OX=10090 GN=Sirt2 PE=1 SV=2 |
| NSDHL_MOUSE | Sterol-4-alpha-carboxylate 3-dehydrogenase, decarboxylating OS=Mus musculus OX=10090 GN=Nsdhl PE=1 SV=1 |
| T126A_MOUSE | Transmembrane protein 126A OS=Mus musculus OX=10090 GN=Tmem126a PE=1 SV=1 |
| NPL4_MOUSE | Nuclear protein localization protein 4 homolog OS=Mus musculus OX=10090 GN=Nploc4 PE=1 SV=3 |
| CPIN1_MOUSE | Anamorsin OS=Mus musculus OX=10090 GN=Ciapin1 PE=1 SV=1 |
| DC1I1_HUMAN | Cytoplasmic dynein 1 intermediate chain 1 OS=Homo sapiens OX=9606 GN=DYNC1I1 PE=1 SV=2 |
| BRSK1_MOUSE | Serine/threonine-protein kinase BRSK1 OS=Mus musculus OX=10090 GN=Brsk1 PE=1 SV=1 |
| DLG4_MOUSE | Disks large homolog 4 OS=Mus musculus OX=10090 GN=Dlg4 PE=1 SV=1 |
| UBP11_MOUSE | Ubiquitin carboxyl-terminal hydrolase 11 OS=Mus musculus OX=10090 GN=Usp11 PE=1 SV=4 |
| QRIC1_MOUSE | Glutamine-rich protein 1 OS=Mus musculus OX=10090 GN=Qrich1 PE=1 SV=1 |
| VPS4A_MOUSE | Vacuolar protein sorting-associated protein 4A OS=Mus musculus OX=10090 GN=Vps4a PE=1 SV=1 |
| FNTA_MOUSE | Protein farnesyltransferase/geranylgeranyltransferase type-1 subunit alpha OS=Mus musculus OX=10090 GN=Fnta PE=1 SV=1 |
| TUSC3_MOUSE | Tumor suppressor candidate 3 OS=Mus musculus OX=10090 GN=Tusc3 PE=1 SV=1 |
| ARRB1_MOUSE | Beta-arrestin-1 OS=Mus musculus OX=10090 GN=Arrb1 PE=1 SV=1 |
| HMGCL_MOUSE | Hydroxymethylglutaryl-CoA lyase, mitochondrial OS=Mus musculus OX=10090 GN=Hmgcl PE=1 SV=2 |

| | |
|---|---|
| CC181_MOUSE | Coiled-coil domain-containing protein 181 OS=Mus musculus OX=10090 GN=Ccdc181 PE=1 SV=1 |
| WDR82_MOUSE | WD repeat-containing protein 82 OS=Mus musculus OX=10090 GN=Wdr82 PE=1 SV=1 |
| STAU2_MOUSE | Double-stranded RNA-binding protein Staufen homolog 2 OS=Mus musculus OX=10090 GN=Stau2 PE=1 SV=1 |
| DNJB1_MOUSE | DnaJ homolog subfamily B member 1 OS=Mus musculus OX=10090 GN=Dnajb1 PE=1 SV=3 |
| RUN3A_MOUSE | RUN domain-containing protein 3A OS=Mus musculus OX=10090 GN=Rundc3a PE=1 SV=1 |
| ELMO2_MOUSE | Engulfment and cell motility protein 2 OS=Mus musculus OX=10090 GN=Elmo2 PE=1 SV=1 |
| BIG3_MOUSE | Brefeldin A-inhibited guanine nucleotide-exchange protein 3 OS=Mus musculus OX=10090 GN=Arfgef3 PE=1 SV=1 |
| PREP_MOUSE | Presequence protease, mitochondrial OS=Mus musculus OX=10090 GN=Pitrm1 PE=1 SV=1 |
| GCC2_MOUSE | GRIP and coiled-coil domain-containing protein 2 OS=Mus musculus OX=10090 GN=Gcc2 PE=1 SV=2 |
| CPEB3_MOUSE | Cytoplasmic polyadenylation element-binding protein 3 OS=Mus musculus OX=10090 GN=Cpeb3 PE=1 SV=1 |
| PDE2A_MOUSE | cGMP-dependent 3',5'-cyclic phosphodiesterase OS=Mus musculus OX=10090 GN=Pde2a PE=1 SV=4 |
| PIN1_MOUSE | Peptidyl-prolyl cis-trans isomerase NIMA-interacting 1 OS=Mus musculus OX=10090 GN=Pin1 PE=1 SV=1 |
| RRFM_MOUSE | Ribosome-recycling factor, mitochondrial OS=Mus musculus OX=10090 GN=Mrrf PE=1 SV=1 |
| VP37B_MOUSE | Vacuolar protein sorting-associated protein 37B OS=Mus musculus OX=10090 GN=Vps37b PE=1 SV=1 |
| TTYH1_MOUSE | Protein tweety homolog 1 OS=Mus musculus OX=10090 GN=Ttyh1 PE=1 SV=1 |
| RWDD1_MOUSE | RWD domain-containing protein 1 OS=Mus musculus OX=10090 GN=Rwdd1 PE=1 SV=1 |
| ACTN2_MOUSE | Alpha-actinin-2 OS=Mus musculus OX=10090 GN=Actn2 PE=1 SV=2 |

| | |
|---|---|
| TENA_MOUSE | Tenascin OS=Mus musculus OX=10090 GN=Tnc PE=1 SV=1 |
| FTO_MOUSE | Alpha-ketoglutarate-dependent dioxygenase FTO OS=Mus musculus OX=10090 GN=Fto PE=1 SV=1 |
| CK5P1_MOUSE | CDK5 regulatory subunit-associated protein 1 OS=Mus musculus OX=10090 GN=Cdk5rap1 PE=2 SV=2 |
| ACYP2_MOUSE | Acylphosphatase-2 OS=Mus musculus OX=10090 GN=Acyp2 PE=1 SV=2 |
| RPAB3_MOUSE | DNA-directed RNA polymerases I, II, and III subunit RPABC3 OS=Mus musculus OX=10090 GN=Polr2h PE=1 SV=3 |
| QCR10_MOUSE | Cytochrome b-c1 complex subunit 10 OS=Mus musculus OX=10090 GN=Uqcr11 PE=3 SV=1 |
| MYADM_MOUSE | Myeloid-associated differentiation marker OS=Mus musculus OX=10090 GN=Myadm PE=1 SV=2 |
| PP1G_MOUSE | Serine/threonine-protein phosphatase PP1-gamma catalytic subunit OS=Mus musculus OX=10090 GN=Ppp1cc PE=1 SV=1 |
| RRP1_MOUSE | Ribosomal RNA processing protein 1 homolog A OS=Mus musculus OX=10090 GN=Rrp1 PE=1 SV=2 |
| NEDD8_MOUSE | NEDD8 OS=Mus musculus OX=10090 GN=Nedd8 PE=1 SV=2 |
| HBB_MYOVE | Hemoglobin subunit beta OS=Myotis velifer OX=9435 GN=HBB PE=1 SV=1 |
| ZN428_MOUSE | Zinc finger protein 428 OS=Mus musculus OX=10090 GN=Znf428 PE=1 SV=1 |
| WASF3_MOUSE | Wiskott-Aldrich syndrome protein family member 3 OS=Mus musculus OX=10090 GN=Wasf3 PE=1 SV=1 |
| SESD1_MOUSE | SEC14 domain and spectrin repeat-containing protein 1 OS=Mus musculus OX=10090 GN=Sestd1 PE=1 SV=1 |
| Q9SJL2-DECOY | Q9SJL2 |
| HIG1A_MOUSE | HIG1 domain family member 1A, mitochondrial OS=Mus musculus OX=10090 GN=Higd1a PE=1 SV=1 |
| SC61G_MOUSE | Protein transport protein Sec61 subunit gamma OS=Mus musculus OX=10090 GN=Sec61g PE=3 SV=1 |

| | |
|---|---|
| 2AAB_MOUSE | Serine/threonine-protein phosphatase 2A 65 kDa regulatory subunit A beta isoform OS=Mus musculus OX=10090 GN=Ppp2r1b PE=1 SV=2 |
| NU3M_MOUSE | NADH-ubiquinone oxidoreductase chain 3 OS=Mus musculus OX=10090 GN=Mtnd3 PE=1 SV=3 |
| PTH2_MOUSE | Peptidyl-tRNA hydrolase 2, mitochondrial OS=Mus musculus OX=10090 GN=Ptrh2 PE=1 SV=1 |
| LCLT1_MOUSE | Lysocardiolipin acyltransferase 1 OS=Mus musculus OX=10090 GN=Lclat1 PE=1 SV=2 |
| CDN1B_MOUSE | Cyclin-dependent kinase inhibitor 1B OS=Mus musculus OX=10090 GN=Cdkn1b PE=1 SV=2 |
| 7B2_MOUSE | Neuroendocrine protein 7B2 OS=Mus musculus OX=10090 GN=Scg5 PE=1 SV=1 |
| MYPT2_MOUSE | Protein phosphatase 1 regulatory subunit 12B OS=Mus musculus OX=10090 GN=Ppp1r12b PE=1 SV=2 |
| FAHD2_MOUSE | Fumarylacetoacetate hydrolase domain-containing protein 2A OS=Mus musculus OX=10090 GN=Fahd2 PE=1 SV=1 |
| C1RA_MOUSE | Complement C1r-A subcomponent OS=Mus musculus OX=10090 GN=C1ra PE=1 SV=1 |
| GLRX5_MOUSE | Glutaredoxin-related protein 5, mitochondrial OS=Mus musculus OX=10090 GN=Glrx5 PE=1 SV=2 |
| EMC4_MOUSE | ER membrane protein complex subunit 4 OS=Mus musculus OX=10090 GN=Emc4 PE=1 SV=1 |
| ARMC1_MOUSE | Armadillo repeat-containing protein 1 OS=Mus musculus OX=10090 GN=Armc1 PE=1 SV=1 |
| LIPL_MOUSE | Lipoprotein lipase OS=Mus musculus OX=10090 GN=Lpl PE=1 SV=3 |
| RL29_MOUSE | 60S ribosomal protein L29 OS=Mus musculus OX=10090 GN=Rpl29 PE=1 SV=2 |
| SDC4_MOUSE | Syndecan-4 OS=Mus musculus OX=10090 GN=Sdc4 PE=1 SV=1 |
| NET1_MOUSE | Netrin-1 OS=Mus musculus OX=10090 GN=Ntn1 PE=1 SV=3 |
| RM14_MOUSE | 39S ribosomal protein L14, mitochondrial OS=Mus musculus OX=10090 GN=Mrpl14 PE=1 SV=1 |

| | |
|---|---|
| KCD19_MOUSE | BTB/POZ domain-containing protein KCTD19 OS=Mus musculus OX=10090 GN=Kctd19 PE=1 SV=2 |
| SDC3_MOUSE | Syndecan-3 OS=Mus musculus OX=10090 GN=Sdc3 PE=1 SV=2 |
| DJB12_MOUSE | DnaJ homolog subfamily B member 12 OS=Mus musculus OX=10090 GN=Dnajb12 PE=1 SV=2 |
| CAND2_MOUSE | Cullin-associated NEDD8-dissociated protein 2 OS=Mus musculus OX=10090 GN=Cand2 PE=1 SV=2 |
| CPSF3_MOUSE | Cleavage and polyadenylation specificity factor subunit 3 OS=Mus musculus OX=10090 GN=Cpsf3 PE=1 SV=2 |
| PXL2A_MOUSE | Peroxiredoxin-like 2A OS=Mus musculus OX=10090 GN=Prxl2a PE=1 SV=2 |
| DHX29_MOUSE | ATP-dependent RNA helicase DHX29 OS=Mus musculus OX=10090 GN=Dhx29 PE=1 SV=1 |
| MAST4_MOUSE | Microtubule-associated serine/threonine-protein kinase 4 OS=Mus musculus OX=10090 GN=Mast4 PE=1 SV=3 |
| P4K2A_MOUSE | Phosphatidylinositol 4-kinase type 2-alpha OS=Mus musculus OX=10090 GN=Pi4k2a PE=1 SV=1 |
| BGLR_MOUSE | Beta-glucuronidase OS=Mus musculus OX=10090 GN=Gusb PE=1 SV=2 |
| ARFP2_MOUSE | Arfaptin-2 OS=Mus musculus OX=10090 GN=Arfip2 PE=1 SV=2 |
| RT02_MOUSE | 28S ribosomal protein S2, mitochondrial OS=Mus musculus OX=10090 GN=Mrps2 PE=1 SV=1 |
| HOME1_MOUSE | Homer protein homolog 1 OS=Mus musculus OX=10090 GN=Homer1 PE=1 SV=2 |
| MOGS_MOUSE | Mannosyl-oligosaccharide glucosidase OS=Mus musculus OX=10090 GN=Mogs PE=1 SV=1 |
| ERGI3_MOUSE | Endoplasmic reticulum-Golgi intermediate compartment protein 3 OS=Mus musculus OX=10090 GN=Ergic3 PE=1 SV=1 |
| MTAP_MOUSE | S-methyl-5'-thioadenosine phosphorylase OS=Mus musculus OX=10090 GN=Mtap PE=1 SV=1 |
| SMCE1_MOUSE | SWI/SNF-related matrix-associated actin-dependent regulator of chromatin subfamily E member 1 OS=Mus musculus OX=10090 GN=Smarce1 PE=1 SV=1 |

| | |
|---|---|
| WFS1_MOUSE | Wolframin OS=Mus musculus OX=10090 GN=Wfs1 PE=1 SV=1 |
| PLCA_MOUSE | 1-acyl-sn-glycerol-3-phosphate acyltransferase alpha OS=Mus musculus OX=10090 GN=Agpat1 PE=1 SV=1 |
| NDEL1_MOUSE | Nuclear distribution protein nudE-like 1 OS=Mus musculus OX=10090 GN=Ndel1 PE=1 SV=2 |
| VAC14_MOUSE | Protein VAC14 homolog OS=Mus musculus OX=10090 GN=Vac14 PE=1 SV=1 |
| MYO5A_HUMAN | Unconventional myosin-Va OS=Homo sapiens OX=9606 GN=MYO5A PE=1 SV=2 |
| SACS_MOUSE | Sacsin OS=Mus musculus OX=10090 GN=Sacs PE=1 SV=2 |
| 5NTC_MOUSE | Cytosolic purine 5'-nucleotidase OS=Mus musculus OX=10090 GN=Nt5c2 PE=1 SV=2 |
| CCHL_MOUSE | Cytochrome c-type heme lyase OS=Mus musculus OX=10090 GN=Hccs PE=1 SV=2 |
| FPRP_MOUSE | Prostaglandin F2 receptor negative regulator OS=Mus musculus OX=10090 GN=Ptgfrn PE=1 SV=2 |
| MS5L2_ARATH | Protein POLLENLESS 3-LIKE 2 OS=Arabidopsis thaliana OX=3702 GN=At3g51280 PE=2 SV=1 |
| CLVS2_MOUSE | Clavesin-2 OS=Mus musculus OX=10090 GN=Clvs2 PE=1 SV=1 |
| DOCK3_MOUSE | Dedicator of cytokinesis protein 3 OS=Mus musculus OX=10090 GN=Dock3 PE=1 SV=1 |
| CATZ_MOUSE | Cathepsin Z OS=Mus musculus OX=10090 GN=Ctsz PE=1 SV=1 |
| T132A_MOUSE | Transmembrane protein 132A OS=Mus musculus OX=10090 GN=Tmem132a PE=1 SV=2 |
| NSG2_MOUSE | Neuronal vesicle trafficking-associated protein 2 OS=Mus musculus OX=10090 GN=Nsg2 PE=1 SV=1 |
| AL3A2_MOUSE | Aldehyde dehydrogenase family 3 member A2 OS=Mus musculus OX=10090 GN=Aldh3a2 PE=1 SV=2 |
| A8R7K9-DECOY | A8R7K9 |
| TCRG1_MOUSE | Transcription elongation regulator 1 OS=Mus musculus OX=10090 GN=Tcerg1 PE=1 SV=2 |
| ABCF2_MOUSE | ATP-binding cassette sub-family F member 2 OS=Mus musculus OX=10090 GN=Abcf2 PE=1 SV=1 |

| | |
|---|---|
| TENR_MOUSE | Tenascin-R OS=Mus musculus OX=10090 GN=Tnr PE=1 SV=2 |
| PEX19_MOUSE | Peroxisomal biogenesis factor 19 OS=Mus musculus OX=10090 GN=Pex19 PE=1 SV=1 |
| ERI3_MOUSE | ERI1 exoribonuclease 3 OS=Mus musculus OX=10090 GN=Eri3 PE=1 SV=1 |
| TMCO1_MOUSE | Calcium load-activated calcium channel OS=Mus musculus OX=10090 GN=Tmco1 PE=1 SV=1 |
| UBA3_MOUSE | NEDD8-activating enzyme E1 catalytic subunit OS=Mus musculus OX=10090 GN=Uba3 PE=1 SV=2 |
| IF2P_MOUSE | Eukaryotic translation initiation factor 5B OS=Mus musculus OX=10090 GN=Eif5b PE=1 SV=2 |
| GTR3_MOUSE | Solute carrier family 2, facilitated glucose transporter member 3 OS=Mus musculus OX=10090 GN=Slc2a3 PE=1 SV=1 |
| ACL6B_MOUSE | Actin-like protein 6B OS=Mus musculus OX=10090 GN=Actl6b PE=1 SV=1 |
| ARM10_MOUSE | Armadillo repeat-containing protein 10 OS=Mus musculus OX=10090 GN=Armc10 PE=1 SV=1 |
| ACAD8_MOUSE | Isobutyryl-CoA dehydrogenase, mitochondrial OS=Mus musculus OX=10090 GN=Acad8 PE=1 SV=2 |
| PGTB2_MOUSE | Geranylgeranyl transferase type-2 subunit beta OS=Mus musculus OX=10090 GN=Rabggtb PE=1 SV=2 |
| ATIF1_MOUSE | ATPase inhibitor, mitochondrial OS=Mus musculus OX=10090 GN=ATP5IF1 PE=1 SV=2 |
| LARP7_MOUSE | La-related protein 7 OS=Mus musculus OX=10090 GN=Larp7 PE=1 SV=2 |
| AFAP1_MOUSE | Actin filament-associated protein 1 OS=Mus musculus OX=10090 GN=Afap1 PE=1 SV=1 |
| KALRN_MOUSE | Kalirin OS=Mus musculus OX=10090 GN=Kalrn PE=1 SV=1 |
| CON__P08779 | CON__P08779 |
| UBP24_MOUSE | Ubiquitin carboxyl-terminal hydrolase 24 OS=Mus musculus OX=10090 GN=Usp24 PE=1 SV=1 |
| MYO1C_MOUSE | Unconventional myosin-Ic OS=Mus musculus OX=10090 GN=Myo1c PE=1 SV=2 |

| | |
|---|---|
| COA3_MOUSE | Cytochrome c oxidase assembly factor 3 homolog, mitochondrial OS=Mus musculus OX=10090 GN=Coa3 PE=1 SV=1 |
| SYUB_MOUSE | Beta-synuclein OS=Mus musculus OX=10090 GN=Sncb PE=1 SV=1 |
| LUC7L_MOUSE | Putative RNA-binding protein Luc7-like 1 OS=Mus musculus OX=10090 GN=Luc7l PE=1 SV=2 |
| PCSK1_MOUSE | ProSAAS OS=Mus musculus OX=10090 GN=Pcsk1n PE=1 SV=2 |
| CP46A_MOUSE | Cholesterol 24-hydroxylase OS=Mus musculus OX=10090 GN=Cyp46a1 PE=1 SV=1 |
| GBRA3_MOUSE | Gamma-aminobutyric acid receptor subunit alpha-3 OS=Mus musculus OX=10090 GN=Gabra3 PE=1 SV=1 |
| TIM8B_MOUSE | Mitochondrial import inner membrane translocase subunit Tim8 B OS=Mus musculus OX=10090 GN=Timm8b PE=1 SV=1 |
| MPPB_MOUSE | Mitochondrial-processing peptidase subunit beta OS=Mus musculus OX=10090 GN=Pmpcb PE=1 SV=1 |
| PP4R2_MOUSE | Serine/threonine-protein phosphatase 4 regulatory subunit 2 OS=Mus musculus OX=10090 GN=Ppp4r2 PE=1 SV=1 |
| GBG12_MOUSE | Guanine nucleotide-binding protein G(I)/G(S)/G(O) subunit gamma-12 OS=Mus musculus OX=10090 GN=Gng12 PE=1 SV=3 |
| COASY_MOUSE | Bifunctional coenzyme A synthase OS=Mus musculus OX=10090 GN=Coasy PE=1 SV=2 |
| NIT1_MOUSE | Deaminated glutathione amidase OS=Mus musculus OX=10090 GN=Nit1 PE=1 SV=2 |
| TMM33_MOUSE | Transmembrane protein 33 OS=Mus musculus OX=10090 GN=Tmem33 PE=1 SV=1 |
| ARF2_MOUSE | ADP-ribosylation factor 2 OS=Mus musculus OX=10090 GN=Arf2 PE=1 SV=2 |
| TF3C5_MOUSE | General transcription factor 3C polypeptide 5 OS=Mus musculus OX=10090 GN=Gtf3c5 PE=2 SV=2 |
| PEX14_MOUSE | Peroxisomal membrane protein PEX14 OS=Mus musculus OX=10090 GN=Pex14 PE=1 SV=1 |
| TM163_MOUSE | Transmembrane protein 163 OS=Mus musculus OX=10090 GN=Tmem163 PE=1 SV=1 |

| | |
|---|---|
| S10AB_MOUSE | Protein S100-A11 OS=Mus musculus OX=10090 GN=S100a11 PE=1 SV=1 |
| QOR_MOUSE | Quinone oxidoreductase OS=Mus musculus OX=10090 GN=Cryz PE=1 SV=1 |
| ACOX1_ARATH | Peroxisomal acyl-coenzyme A oxidase 1 OS=Arabidopsis thaliana OX=3702 GN=ACX1 PE=1 SV=1 |
| OTC_MOUSE | Ornithine carbamoyltransferase, mitochondrial OS=Mus musculus OX=10090 GN=Otc PE=1 SV=1 |
| LACTB_MOUSE | Serine beta-lactamase-like protein LACTB, mitochondrial OS=Mus musculus OX=10090 GN=Lactb PE=1 SV=1 |
| ZYX_MOUSE | Zyxin OS=Mus musculus OX=10090 GN=Zyx PE=1 SV=2 |
| NACAD_MOUSE | NAC-alpha domain-containing protein 1 OS=Mus musculus OX=10090 GN=Nacad PE=1 SV=1 |
| HAT1_MOUSE | Histone acetyltransferase type B catalytic subunit OS=Mus musculus OX=10090 GN=Hat1 PE=1 SV=1 |
| TPC6B_MOUSE | Trafficking protein particle complex subunit 6B OS=Mus musculus OX=10090 GN=Trappc6b PE=1 SV=1 |
| F169A_MOUSE | Soluble lamin-associated protein of 75 kDa OS=Mus musculus OX=10090 GN=Fam169a PE=1 SV=3 |
| CPSF1_MOUSE | Cleavage and polyadenylation specificity factor subunit 1 OS=Mus musculus OX=10090 GN=Cpsf1 PE=1 SV=1 |
| RN214_MOUSE | RING finger protein 214 OS=Mus musculus OX=10090 GN=Rnf214 PE=1 SV=1 |
| EVI5_MOUSE | Ecotropic viral integration site 5 protein OS=Mus musculus OX=10090 GN=Evi5 PE=1 SV=2 |
| MMGT1_MOUSE | Membrane magnesium transporter 1 OS=Mus musculus OX=10090 GN=Mmgt1 PE=1 SV=1 |
| TIAM1_MOUSE | T-lymphoma invasion and metastasis-inducing protein 1 OS=Mus musculus OX=10090 GN=Tiam1 PE=1 SV=1 |
| BABA2_CHLAE | BRISC and BRCA1-A complex member 2 OS=Chlorocebus aethiops OX=9534 GN=BABAM2 PE=2 SV=1 |
| Q9WUH2-DECOY | Q9WUH2 |

| | |
|---|---|
| FACE1_MOUSE | CAAX prenyl protease 1 homolog OS=Mus musculus OX=10090 GN=Zmpste24 PE=1 SV=2 |
| UBE3C_MOUSE | Ubiquitin-protein ligase E3C OS=Mus musculus OX=10090 GN=Ube3c PE=1 SV=2 |
| VAMP4_MOUSE | Vesicle-associated membrane protein 4 OS=Mus musculus OX=10090 GN=Vamp4 PE=1 SV=1 |
| MPI_MOUSE | Mannose-6-phosphate isomerase OS=Mus musculus OX=10090 GN=Mpi PE=1 SV=1 |
| TSN7_MOUSE | Tetraspanin-7 OS=Mus musculus OX=10090 GN=Tspan7 PE=1 SV=2 |
| SCG1_MOUSE | Secretogranin-1 OS=Mus musculus OX=10090 GN=Chgb PE=1 SV=2 |
| P66B_MOUSE | Transcriptional repressor p66-beta OS=Mus musculus OX=10090 GN=Gatad2b PE=1 SV=1 |
| Q1PEY6-DECOY | Q1PEY6 |
| SMRD1_MOUSE | SWI/SNF-related matrix-associated actin-dependent regulator of chromatin subfamily D member 1 OS=Mus musculus OX=10090 GN=Smarcd1 PE=1 SV=3 |
| PELP1_MOUSE | Proline-, glutamic acid- and leucine-rich protein 1 OS=Mus musculus OX=10090 GN=Pelp1 PE=1 SV=2 |
| HSP13_MOUSE | Heat shock 70 kDa protein 13 OS=Mus musculus OX=10090 GN=Hspa13 PE=1 SV=1 |
| C560_MOUSE | Succinate dehydrogenase cytochrome b560 subunit, mitochondrial OS=Mus musculus OX=10090 GN=Sdhc PE=1 SV=1 |
| HDAC1_MOUSE | Histone deacetylase 1 OS=Mus musculus OX=10090 GN=Hdac1 PE=1 SV=1 |
| CCAR1_MOUSE | Cell division cycle and apoptosis regulator protein 1 OS=Mus musculus OX=10090 GN=Ccar1 PE=1 SV=1 |
| CPNE1_MOUSE | Copine-1 OS=Mus musculus OX=10090 GN=Cpne1 PE=1 SV=1 |
| TMM11_MOUSE | Transmembrane protein 11, mitochondrial OS=Mus musculus OX=10090 GN=Tmem11 PE=1 SV=1 |
| SFRP1_MOUSE | Secreted frizzled-related protein 1 OS=Mus musculus OX=10090 GN=Sfrp1 PE=1 SV=3 |
| MYPT1_MOUSE | Protein phosphatase 1 regulatory subunit 12A OS=Mus musculus OX=10090 GN=Ppp1r12a PE=1 SV=2 |

| | |
|---|---|
| GBB4_MOUSE | Guanine nucleotide-binding protein subunit beta-4 OS=Mus musculus OX=10090 GN=Gnb4 PE=1 SV=4 |
| CAN5_MOUSE | Calpain-5 OS=Mus musculus OX=10090 GN=Capn5 PE=1 SV=1 |
| SPRY4_MOUSE | SPRY domain-containing protein 4 OS=Mus musculus OX=10090 GN=Spryd4 PE=1 SV=1 |
| APEH_MOUSE | Acylamino-acid-releasing enzyme OS=Mus musculus OX=10090 GN=Apeh PE=1 SV=3 |
| GSH0_MOUSE | Glutamate--cysteine ligase regulatory subunit OS=Mus musculus OX=10090 GN=Gclm PE=1 SV=1 |
| SAM50_MOUSE | Sorting and assembly machinery component 50 homolog OS=Mus musculus OX=10090 GN=Samm50 PE=1 SV=1 |
| DEN10_MOUSE | DENN domain-containing protein 10 OS=Mus musculus OX=10090 GN=Dennd10 PE=1 SV=2 |
| Y1285_ARATH | Putative UPF0725 protein At1g28500 OS=Arabidopsis thaliana OX=3702 GN=At1g28500 PE=3 SV=1 |
| PMVK_MOUSE | Phosphomevalonate kinase OS=Mus musculus OX=10090 GN=Pmvk PE=1 SV=3 |
| GAPD1_MOUSE | GTPase-activating protein and VPS9 domain-containing protein 1 OS=Mus musculus OX=10090 GN=Gapvd1 PE=1 SV=2 |
| A2AGT5-DECOY | A2AGT5 |
| IBP3_MOUSE | Insulin-like growth factor-binding protein 3 OS=Mus musculus OX=10090 GN=Igfbp3 PE=2 SV=2 |
| TGRM1_MOUSE | TOG array regulator of axonemal microtubules protein 1 OS=Mus musculus OX=10090 GN=Togaram1 PE=1 SV=3 |
| TACO1_MOUSE | Translational activator of cytochrome c oxidase 1 OS=Mus musculus OX=10090 GN=Taco1 PE=1 SV=1 |
| ACBD6_MOUSE | Acyl-CoA-binding domain-containing protein 6 OS=Mus musculus OX=10090 GN=Acbd6 PE=1 SV=2 |
| TNPO3_MOUSE | Transportin-3 OS=Mus musculus OX=10090 GN=Tnpo3 PE=1 SV=1 |
| PCD10_HUMAN | Protocadherin-10 OS=Homo sapiens OX=9606 GN=PCDH10 PE=2 SV=2 |

| | |
|---|---|
| CHTOP_MOUSE | Chromatin target of PRMT1 protein OS=Mus musculus OX=10090 GN=Chtop PE=1 SV=2 |
| ARHL2_MOUSE | ADP-ribose glycohydrolase ARH3 OS=Mus musculus OX=10090 GN=Adprhl2 PE=1 SV=1 |
| QCR6_MOUSE | Cytochrome b-c1 complex subunit 6, mitochondrial OS=Mus musculus OX=10090 GN=Uqcrh PE=1 SV=2 |
| FXYD6_MOUSE | FXYD domain-containing ion transport regulator 6 OS=Mus musculus OX=10090 GN=Fxyd6 PE=1 SV=2 |
| RFOX1_MOUSE | RNA binding protein fox-1 homolog 1 OS=Mus musculus OX=10090 GN=Rbfox1 PE=1 SV=3 |
| DPYS_MOUSE | Dihydropyrimidinase OS=Mus musculus OX=10090 GN=Dpys PE=1 SV=2 |
| S2539_MOUSE | Solute carrier family 25 member 39 OS=Mus musculus OX=10090 GN=Slc25a39 PE=2 SV=1 |
| YES_MOUSE | Tyrosine-protein kinase Yes OS=Mus musculus OX=10090 GN=Yes1 PE=1 SV=3 |
| UBQL1_MOUSE | Ubiquilin-1 OS=Mus musculus OX=10090 GN=Ubqln1 PE=1 SV=1 |
| MYL9_MOUSE | Myosin regulatory light polypeptide 9 OS=Mus musculus OX=10090 GN=Myl9 PE=1 SV=3 |
| CF97D_MOUSE | Uncharacterized protein CFAP97D1 OS=Mus musculus OX=10090 GN=Cfap97d1 PE=2 SV=1 |
| CC115_MOUSE | Coiled-coil domain-containing protein 115 OS=Mus musculus OX=10090 GN=Ccdc115 PE=1 SV=1 |
| WRB_MOUSE | Tail-anchored protein insertion receptor WRB OS=Mus musculus OX=10090 GN=Wrb PE=1 SV=1 |
| SCAI_MOUSE | Protein SCAI OS=Mus musculus OX=10090 GN=Scai PE=1 SV=2 |
| IHH_MOUSE | Indian hedgehog protein OS=Mus musculus OX=10090 GN=Ihh PE=1 SV=2 |
| RAB2B_MOUSE | Ras-related protein Rab-2B OS=Mus musculus OX=10090 GN=Rab2b PE=1 SV=1 |
| PP4C_MOUSE | Serine/threonine-protein phosphatase 4 catalytic subunit OS=Mus musculus OX=10090 GN=Ppp4c PE=1 SV=2 |
| SPZ1_ARATH | Serpin-Z1 OS=Arabidopsis thaliana OX=3702 GN=At1g64030 PE=2 SV=2 |

| | |
|---|---|
| KAD5_MOUSE | Adenylate kinase isoenzyme 5 OS=Mus musculus OX=10090 GN=Ak5 PE=1 SV=2 |
| RM46_MOUSE | 39S ribosomal protein L46, mitochondrial OS=Mus musculus OX=10090 GN=Mrpl46 PE=1 SV=1 |
| RT10_MOUSE | 28S ribosomal protein S10, mitochondrial OS=Mus musculus OX=10090 GN=Mrps10 PE=1 SV=1 |
| LCAP_MOUSE | Leucyl-cystinyl aminopeptidase OS=Mus musculus OX=10090 GN=Lnpep PE=1 SV=1 |
| TSNAX_MOUSE | Translin-associated protein X OS=Mus musculus OX=10090 GN=Tsnax PE=1 SV=1 |
| ABCB7_MOUSE | ATP-binding cassette sub-family B member 7, mitochondrial OS=Mus musculus OX=10090 GN=Abcb7 PE=1 SV=3 |
| PARVA_MOUSE | Alpha-parvin OS=Mus musculus OX=10090 GN=Parva PE=1 SV=1 |
| ACM1_MOUSE | Muscarinic acetylcholine receptor M1 OS=Mus musculus OX=10090 GN=Chrm1 PE=1 SV=2 |
| RCAN2_MOUSE | Calcipressin-2 OS=Mus musculus OX=10090 GN=Rcan2 PE=1 SV=1 |
| SPF27_MOUSE | Pre-mRNA-splicing factor SPF27 OS=Mus musculus OX=10090 GN=Bcas2 PE=1 SV=1 |
| CBX6_MOUSE | Chromobox protein homolog 6 OS=Mus musculus OX=10090 GN=Cbx6 PE=1 SV=2 |
| RCN3_MOUSE | Reticulocalbin-3 OS=Mus musculus OX=10090 GN=Rcn3 PE=1 SV=1 |
| KV3A8_MOUSE | Ig kappa chain V-III region PC 3741/TEPC 111 OS=Mus musculus OX=10090 PE=1 SV=1 |
| HMGN1_MOUSE | Non-histone chromosomal protein HMG-14 OS=Mus musculus OX=10090 GN=Hmgn1 PE=1 SV=2 |
| CNOT9_MOUSE | CCR4-NOT transcription complex subunit 9 OS=Mus musculus OX=10090 GN=Cnot9 PE=1 SV=1 |
| SBP1_RAT | Methanethiol oxidase OS=Rattus norvegicus OX=10116 GN=Selenbp1 PE=1 SV=1 |
| MAON_MOUSE | NADP-dependent malic enzyme, mitochondrial OS=Mus musculus OX=10090 GN=Me3 PE=1 SV=2 |
| TIM10_MOUSE | Mitochondrial import inner membrane translocase subunit Tim10 OS=Mus musculus OX=10090 GN=Timm10 PE=1 SV=1 |

| | |
|---|---|
| ARHG7_MOUSE | Rho guanine nucleotide exchange factor 7 OS=Mus musculus OX=10090 GN=Arhgef7 PE=1 SV=2 |
| GCP60_MOUSE | Golgi resident protein GCP60 OS=Mus musculus OX=10090 GN=Acbd3 PE=1 SV=3 |
| MK_MOUSE | Midkine OS=Mus musculus OX=10090 GN=Mdk PE=1 SV=2 |
| CSN8_MOUSE | COP9 signalosome complex subunit 8 OS=Mus musculus OX=10090 GN=Cops8 PE=1 SV=1 |
| CON__Q15323 | CON__Q15323 |
| CCD22_MOUSE | Coiled-coil domain-containing protein 22 OS=Mus musculus OX=10090 GN=Ccdc22 PE=1 SV=1 |
| KT3K_MOUSE | Ketosamine-3-kinase OS=Mus musculus OX=10090 GN=Fn3krp PE=1 SV=2 |
| UBR3_MOUSE | E3 ubiquitin-protein ligase UBR3 OS=Mus musculus OX=10090 GN=Ubr3 PE=1 SV=3 |
| MIRO2_MOUSE | Mitochondrial Rho GTPase 2 OS=Mus musculus OX=10090 GN=Rhot2 PE=1 SV=1 |
| MTSS2_MOUSE | Protein MTSS 2 OS=Mus musculus OX=10090 GN=Mtss2 PE=1 SV=1 |
| COCA1_HUMAN | Collagen alpha-1(XII) chain OS=Homo sapiens OX=9606 GN=COL12A1 PE=1 SV=2 |
| T22D1_MOUSE | TSC22 domain family protein 1 OS=Mus musculus OX=10090 GN=Tsc22d1 PE=1 SV=2 |
| ZC3HF_HUMAN | Zinc finger CCCH domain-containing protein 15 OS=Homo sapiens OX=9606 GN=ZC3H15 PE=1 SV=1 |
| EXOC8_MOUSE | Exocyst complex component 8 OS=Mus musculus OX=10090 GN=Exoc8 PE=1 SV=1 |
| SNX30_MOUSE | Sorting nexin-30 OS=Mus musculus OX=10090 GN=Snx30 PE=1 SV=1 |
| MON2_MOUSE | Protein MON2 homolog OS=Mus musculus OX=10090 GN=Mon2 PE=1 SV=2 |
| AKP8L_MOUSE | A-kinase anchor protein 8-like OS=Mus musculus OX=10090 GN=Akap8l PE=1 SV=1 |
| KI26B_MOUSE | Kinesin-like protein KIF26B OS=Mus musculus OX=10090 GN=Kif26b PE=1 SV=3 |
| TBCE_MOUSE | Tubulin-specific chaperone E OS=Mus musculus OX=10090 GN=Tbce PE=1 SV=1 |

| | |
|---|---|
| NU5M_MOUSE | NADH-ubiquinone oxidoreductase chain 5 OS=Mus musculus OX=10090 GN=Mtnd5 PE=1 SV=3 |
| PPM1L_MOUSE | Protein phosphatase 1L OS=Mus musculus OX=10090 GN=Ppm1l PE=1 SV=1 |
| CON__Q3ZBS7 | CON__Q3ZBS7 |
| PESC_MOUSE | Pescadillo homolog OS=Mus musculus OX=10090 GN=Pes1 PE=1 SV=1 |
| NRP1_MOUSE | Neuropilin-1 OS=Mus musculus OX=10090 GN=Nrp1 PE=1 SV=2 |
| LYPA2_MOUSE | Acyl-protein thioesterase 2 OS=Mus musculus OX=10090 GN=Lypla2 PE=1 SV=1 |
| DCMC_MOUSE | Malonyl-CoA decarboxylase, mitochondrial OS=Mus musculus OX=10090 GN=Mlycd PE=1 SV=1 |
| KN7L_ARATH | Kinesin-like protein KIN-7L, chloroplastic OS=Arabidopsis thaliana OX=3702 GN=KIN7L PE=3 SV=2 |
| MA7D2_MOUSE | MAP7 domain-containing protein 2 OS=Mus musculus OX=10090 GN=Map7d2 PE=1 SV=1 |
| A16L1_HUMAN | Autophagy-related protein 16-1 OS=Homo sapiens OX=9606 GN=ATG16L1 PE=1 SV=2 |
| CDC23_ARATH | Anaphase-promoting complex subunit 8 OS=Arabidopsis thaliana OX=3702 GN=APC8 PE=1 SV=1 |
| IMA1_MOUSE | Importin subunit alpha-1 OS=Mus musculus OX=10090 GN=Kpna2 PE=1 SV=2 |
| O48671-DECOY | O48671 |
| ODPAT_MOUSE | Pyruvate dehydrogenase E1 component subunit alpha, testis-specific form, mitochondrial OS=Mus musculus OX=10090 GN=Pdha2 PE=1 SV=1 |
| PGPI_MOUSE | Pyroglutamyl-peptidase 1 OS=Mus musculus OX=10090 GN=Pgpep1 PE=1 SV=1 |
| PRPS2_MOUSE | Ribose-phosphate pyrophosphokinase 2 OS=Mus musculus OX=10090 GN=Prps2 PE=1 SV=4 |
| SFR1_MOUSE | Swi5-dependent recombination DNA repair protein 1 homolog OS=Mus musculus OX=10090 GN=Sfr1 PE=1 SV=2 |
| TXD12_MOUSE | Thioredoxin domain-containing protein 12 OS=Mus musculus OX=10090 GN=Txndc12 PE=1 SV=1 |

| | |
|---|---|
| UQCC1_MOUSE | Ubiquinol-cytochrome-c reductase complex assembly factor 1 OS=Mus musculus OX=10090 GN=Uqcc1 PE=1 SV=1 |
| ZN235_HUMAN | Zinc finger protein 235 OS=Homo sapiens OX=9606 GN=ZNF235 PE=2 SV=3 |
