## Supplementary Data 1 for "The low complexity motif of cytoplasmic polyadenylation element binding protein 3 (CPEB3) is critical for the trafficking of its targets in neurons"

Supplementary Data 1. Proteomics results, HEK

| | |
|---|---|
| Q3TKA2_MOUSE | Uncharacterized protein OS=Mus musculus OX=10090 GN=Hsp90aa1 PE=2 SV=1 |
| NP_001277756.1 cytoplasmic polyadenylation element-binding protein 3 isoform a [Mus... | cytoplasmic polyadenylation element-binding protein 3 isoform a [Mus musculus] |
| Q5SXR6_MOUSE | Clathrin heavy chain OS=Mus musculus OX=10090 GN=Cltc PE=1 SV=1 |
| Q5FWJ3_MOUSE | Vimentin OS=Mus musculus OX=10090 GN=Vim PE=1 SV=1 |
| EF2_MOUSE | Elongation factor 2 OS=Mus musculus OX=10090 GN=Eef2 PE=1 SV=2 |
| Q3TQ13_MOUSE | Uncharacterized protein OS=Mus musculus OX=10090 GN=Hspa8 PE=2 SV=1 |
| Q52L87_MOUSE | Tubulin alpha chain OS=Mus musculus OX=10090 GN=Tuba1c PE=1 SV=1 |
| sp\|P52480\|KPYM_MOUSE | Pyruvate kinase PKM OS=Mus musculus OX=10090 GN=Pkm PE=1 SV=4 |
| Q3TWN5_MOUSE | Uncharacterized protein OS=Mus musculus OX=10090 GN=Hnrnpu PE=2 SV=1 |
| Q3UH59_MOUSE | Myosin-10 OS=Mus musculus OX=10090 GN=Myh10 PE=1 SV=1 |
| sp\|Q8BTM8\|FLNA_MOUSE | Filamin-A OS=Mus musculus OX=10090 GN=Flna PE=1 SV=5 |
| TLN1_MOUSE | Talin-1 OS=Mus musculus OX=10090 GN=Tln1 PE=1 SV=2 |
| Q3TII3_MOUSE | Elongation factor 1-alpha OS=Mus musculus OX=10090 GN=Eef1a1 PE=2 SV=1 |
| ACLY_MOUSE | ATP-citrate synthase OS=Mus musculus OX=10090 GN=Acly PE=1 SV=1 |
| G5E866_MOUSE | Splicing factor 3B subunit 1 OS=Mus musculus OX=10090 GN=Sf3b1 PE=1 SV=1 |
| Q6A0B9_MOUSE | Eukaryotic translation initiation factor 3 subunit A (Fragment) OS=Mus musculus OX=10090 GN=Eif3a PE=2 SV=1 |
| Q3TFG3_MOUSE | Uncharacterized protein OS=Mus musculus OX=10090 GN=Eif4a1 PE=2 SV=1 |
| Q570Z0_MOUSE | MKIAA4193 protein (Fragment) OS=Mus musculus OX=10090 GN=Hnrnpm PE=2 SV=1 |

| | |
|---|---|
| Q8BNF8_MOUSE | Uncharacterized protein OS=Mus musculus OX=10090 GN=Vcp PE=2 SV=1 |
| XPO2_MOUSE | Exportin-2 OS=Mus musculus OX=10090 GN=Cse1l PE=1 SV=1 |
| Q63ZW9_MOUSE | Copa protein (Fragment) OS=Mus musculus OX=10090 GN=Copa PE=2 SV=1 |
| E9Q8K5_MOUSE | Titin OS=Mus musculus OX=10090 GN=Ttn PE=1 SV=1 |
| Q3TIV3_MOUSE | Poly [ADP-ribose] polymerase OS=Mus musculus OX=10090 GN=Parp1 PE=2 SV=1 |
| Q3U4U6_MOUSE | T-complex protein 1 subunit gamma OS=Mus musculus OX=10090 GN=Cct3 PE=1 SV=1 |
| tr\|A1E2B8\|A1E2B8_MOUSE Inducible heat shock protein 70 OS=Mus musculus OX=10090 PE=3... | Inducible heat shock protein 70 OS=Mus musculus OX=10090 PE=3 SV=1 |
| G3X956_MOUSE | Suppressor of Ty 16 OS=Mus musculus OX=10090 GN=Supt16 PE=1 SV=1 |
| CHD4_MOUSE | Chromodomain-helicase-DNA-binding protein 4 OS=Mus musculus OX=10090 GN=Chd4 PE=1 SV=1 |
| MYH9_MOUSE | Myosin-9 OS=Mus musculus OX=10090 GN=Myh9 PE=1 SV=4 |
| sp\|Q7TMY8-3\|HUWE1_MOUSE | Isoform 3 of E3 ubiquitin-protein ligase HUWE1 OS=Mus musculus OX=10090 GN=Huwe1 |
| Q5BL18_MOUSE | Matrin 3 OS=Mus musculus OX=10090 GN=Matr3 PE=1 SV=1 |
| RL4_MOUSE | 60S ribosomal protein L4 OS=Mus musculus OX=10090 GN=Rpl4 PE=1 SV=3 |
| sp\|P11983\|TCPA_MOUSE | T-complex protein 1 subunit alpha OS=Mus musculus OX=10090 GN=Tcp1 PE=1 SV=3 |
| sp\|P38647\|GRP75_MOUSEStress-70 protein, mitochondrial OS=Mus musculus OX=10090 GN=Hspa9 PE=1 SV=3 | sp\|P38647\|GRP75_MOUSE Stress-70 protein, mitochondrial OS=Mus musculus OX=10090 GN=Hspa9 PE=1 SV=3 |
| tr\|Q80U36\|Q80U36_MOUSE MKIAA0325 protein (Fragment) OS=Mus musculus OX=10090 GN=Dync1h1... | MKIAA0325 protein (Fragment) OS=Mus musculus OX=10090 GN=Dync1h1 PE=2 SV=1 |
| Q6ZQK2_MOUSE | MKIAA0051 protein (Fragment) OS=Mus musculus OX=10090 GN=Iqgap1 PE=2 SV=1 |

| | |
|---|---|
| tr\|Q3TWF2\|Q3TWF2_MOUSE Uncharacterized protein OS=Mus musculus OX=10090 GN=Hspa5... | Uncharacterized protein OS=Mus musculus OX=10090 GN=Hspa5 PE=2 SV=1 |
| H3BL49_MOUSE | T-complex protein 1 subunit theta OS=Mus musculus OX=10090 GN=Cct8 PE=1 SV=1 |
| sp\|Q61656\|DDX5_MOUSE Probable ATP-dependent RNA helicase DDX5 OS=Mus musculus OX=10090... | Probable ATP-dependent RNA helicase DDX5 OS=Mus musculus OX=10090 GN=Ddx5 PE=1 SV=2 |
| sp\|Q9Z1X4-3\|ILF3_MOUSE | Isoform 3 of Interleukin enhancer-binding factor 3 OS=Mus musculus OX=10090 GN=Ilf3 |
| PLST_MOUSE | Plastin-3 OS=Mus musculus OX=10090 GN=Pls3 PE=1 SV=3 |
| Q3TMY8_MOUSE | Uncharacterized protein OS=Mus musculus OX=10090 GN=Eftud2 PE=2 SV=1 |
| Q5DTS8_MOUSE | DNA helicase (Fragment) OS=Mus musculus OX=10090 GN=Mcm4 PE=2 SV=1 |
| tr\|E9Q8K5\|E9Q8K5_MOUSE-DECOY Titin OS=Mus musculus OX=10090 GN=Ttn PE=1 SV=1 | tr\|E9Q8K5\|E9Q8K5_MOUSE-DECOY Titin OS=Mus musculus OX=10090 GN=Ttn PE=1 SV=1 |
| Q3THH8_MOUSE | Uncharacterized protein OS=Mus musculus OX=10090 GN=Cct7 PE=2 SV=1 |
| tr\|Q3TFE5\|Q3TFE5_MOUSE Uncharacterized protein OS=Mus musculus OX=10090 GN=Dhx15... | Uncharacterized protein OS=Mus musculus OX=10090 GN=Dhx15 PE=2 SV=1 |
| tr\|Q3UAD6\|Q3UAD6_MOUSE Heat shock protein 90kDa beta (Grp94), member 1 OS=Mus musculus... | Heat shock protein 90kDa beta (Grp94), member 1 OS=Mus musculus OX=10090 GN=Hsp90b1 PE=1 SV=1 |
| Q3UJN1_MOUSE | DNA helicase OS=Mus musculus OX=10090 GN=Mcm2 PE=2 SV=1 |
| Q3ULD6_MOUSE | DNA helicase OS=Mus musculus OX=10090 GN=Mcm3 PE=2 SV=1 |
| sp\|Q03265\|ATPA_MOUSE ATP synthase subunit alpha, mitochondrial OS=Mus musculus OX=10090... | ATP synthase subunit alpha, mitochondrial OS=Mus musculus OX=10090 GN=Atp5f1a PE=1 SV=1 |
| RENT1_MOUSE | Isoform 2 of Regulator of nonsense transcripts 1 OS=Mus musculus OX=10090 GN=Upf1 |
| G3X9G4_MOUSE | Dynamin-2 OS=Mus musculus OX=10090 GN=Dnm2 PE=1 SV=1 |
| Q3TIX8_MOUSE | Uncharacterized protein OS=Mus musculus OX=10090 GN=Cct6a PE=2 SV=1 |

| | |
|---|---|
| VINC_MOUSE | Vinculin OS=Mus musculus OX=10090 GN=Vcl PE=1 SV=4 |
| sp\|E9PVA8\|GCN1_MOUSE eIF-2-alpha kinase activator GCN1 OS=Mus musculus OX=10090 GN=Gcn1... | eIF-2-alpha kinase activator GCN1 OS=Mus musculus OX=10090 GN=Gcn1 PE=1 SV=1 |
| sp\|P09411\|PGK1_MOUSE Phosphoglycerate kinase 1 OS=Mus musculus OX=10090 GN=Pgk1 PE=1... | sp\|P09411\|PGK1_MOUSE Phosphoglycerate kinase 1 OS=Mus musculus OX=10090 GN=Pgk1 PE=1... |
| Q3UB15_MOUSE | Uncharacterized protein OS=Mus musculus OX=10090 GN=Rpl3 PE=2 SV=1 |
| XPO1_MOUSE | Exportin-1 OS=Mus musculus OX=10090 GN=Xpo1 PE=1 SV=1 |
| sp\|Q91WQ3\|SYYC_MOUSE Tyrosine--tRNA ligase, cytoplasmic OS=Mus musculus OX=10090... | sp\|Q91WQ3\|SYYC_MOUSE Tyrosine--tRNA ligase, cytoplasmic OS=Mus musculus OX=10090... |
| E9Q039_MOUSE | Aminopeptidase OS=Mus musculus OX=10090 GN=Npepps PE=1 SV=1 |
| Q3UGW8_MOUSE | Uncharacterized protein (Fragment) OS=Mus musculus OX=10090 GN=Aldh18a1 PE=2 SV=1 |
| tr\|Q7TSZ6\|Q7TSZ6_MOUSE Karyopherin (Importin) beta 1 OS=Mus musculus OX=10090 GN=Kpnb1... | tr\|Q7TSZ6\|Q7TSZ6_MOUSE Karyopherin (Importin) beta 1 OS=Mus musculus OX=10090 GN=Kpnb1... |
| FLNB_MOUSE | Filamin-B OS=Mus musculus OX=10090 GN=Flnb PE=1 SV=3 |
| sp\|Q6ZQ38\|CAND1_MOUSECullin-associated NEDD8-dissociated protein 1 OS=Mus musculus OX=10090 GN=Cand1 PE=1 SV=2 | sp\|Q6ZQ38\|CAND1_MOUSE Cullin-associated NEDD8-dissociated protein 1 OS=Mus musculus OX=10090 GN=Cand1 PE=1 SV=2 |
| Q3TFC2_MOUSE | Uncharacterized protein OS=Mus musculus OX=10090 GN=Nono PE=2 SV=1 |
| Q80Y09_MOUSE | Pdcd6ip protein OS=Mus musculus OX=10090 GN=Pdcd6ip PE=2 SV=1 |
| tr\|Q3TJ38\|Q3TJ38_MOUSE Uncharacterized protein OS=Mus musculus OX=10090 GN=Hnrnpk... | tr\|Q3TJ38\|Q3TJ38_MOUSE Uncharacterized protein OS=Mus musculus OX=10090 GN=Hnrnpk... |
| tr\|Q811H6\|Q811H6_MOUSE Hsp90ab1 protein (Fragment) OS=Mus musculus OX=10090 GN=Hsp90ab1... | tr\|Q811H6\|Q811H6_MOUSE Hsp90ab1 protein (Fragment) OS=Mus musculus OX=10090 GN=Hsp90ab1... |
| Q3TW51_MOUSE | Uncharacterized protein OS=Mus musculus OX=10090 GN=Snd1 PE=2 SV=1 |

| | |
|---|---|
| Q3TWL6_MOUSE | 26S proteasome non-ATPase regulatory subunit 2 OS=Mus musculus OX=10090 GN=Psmd2 PE=2 SV=1 |
| tr\|Q3V1M8\|Q3V1M8_MOUSE Uncharacterized protein OS=Mus musculus OX=10090 GN=Hdlbp... | Uncharacterized protein OS=Mus musculus OX=10090 GN=Hdlbp PE=2 SV=1 |
| tr\|Q542X7\|Q542X7_MOUSE Chaperonin subunit 2 (Beta), isoform CRA_a OS=Mus musculus... | Chaperonin subunit 2 (Beta), isoform CRA_a OS=Mus musculus OX=10090 GN=Cct2 PE=1 SV=1 |
| A0A1S6GWH2_MOUSE | Uncharacterized protein OS=Mus musculus OX=10090 GN=Ddx39b PE=2 SV=1 |
| B7ZWF1_MOUSE | Ddx3x protein OS=Mus musculus OX=10090 GN=Ddx3x PE=2 SV=1 |
| Q3UC10_MOUSE | DNA helicase OS=Mus musculus OX=10090 GN=Mcm6 PE=2 SV=1 |
| Q52KC3_MOUSE | DNA helicase OS=Mus musculus OX=10090 GN=Mcm5 PE=1 SV=1 |
| sp\|Q62261-2\|SPTB2_MOUSE Isoform 2 of Spectrin beta chain, non-erythrocytic 1 OS=Mus... | Isoform 2 of Spectrin beta chain, non-erythrocytic 1 OS=Mus musculus OX=10090 GN=Sptbn1 |
| tr\|Q3UIG0\|Q3UIG0_MOUSE Eukaryotic translation initiation factor 3 subunit E OS=Mus... | tr\|Q3UIG0\|Q3UIG0_MOUSE Eukaryotic translation initiation factor 3 subunit E OS=Mus... |
| tr\|Q8C2Q7\|Q8C2Q7_MOUSE Heterogeneous nuclear ribonucleoprotein H OS=Mus musculus... | Heterogeneous nuclear ribonucleoprotein H OS=Mus musculus OX=10090 GN=Hnrnph1 PE=1 SV=1 |
| E9QAI5_MOUSE | CAD protein OS=Mus musculus OX=10090 GN=Cad PE=1 SV=1 |
| NSF_MOUSE | Vesicle-fusing ATPase OS=Mus musculus OX=10090 GN=Nsf PE=1 SV=2 |
| tr\|Q3UJN2\|Q3UJN2_MOUSE RuvB-like helicase OS=Mus musculus OX=10090 GN=Ruvbl1 PE=2... | RuvB-like helicase OS=Mus musculus OX=10090 GN=Ruvbl1 PE=2 SV=1 |
| B9EIU1_MOUSE | Glutamyl-prolyl-tRNA synthetase OS=Mus musculus OX=10090 GN=Eprs PE=2 SV=1 |
| Q05DR5_MOUSE | FACT complex subunit SSRP1 (Fragment) OS=Mus musculus OX=10090 GN=Ssrp1 PE=2 SV=1 |
| SFPQ_MOUSE | Splicing factor, proline- and glutamine-rich OS=Mus musculus OX=10090 GN=Sfpq PE=1 SV=1 |

| | |
|---|---|
| sp\|P16125\|LDHB_MOUSE L-lactate dehydrogenase B chain OS=Mus musculus OX=10090 GN=Ldhb... | L-lactate dehydrogenase B chain OS=Mus musculus OX=10090 GN=Ldhb PE=1 SV=2 |
| Q32P04_MOUSE | Keratin 5 OS=Mus musculus OX=10090 GN=Krt5 PE=1 SV=2 |
| tr\|Q3TXF9\|Q3TXF9_MOUSE Sodium/potassium-transporting ATPase subunit alpha OS=Mus... | Sodium/potassium-transporting ATPase subunit alpha OS=Mus musculus OX=10090 GN=Atp1a1 PE=2 SV=1 |
| tr\|Q3UZH3\|Q3UZH3_MOUSE Uncharacterized protein (Fragment) OS=Mus musculus OX=10090... | Uncharacterized protein (Fragment) OS=Mus musculus OX=10090 GN=Mthfd1 PE=2 SV=1 |
| sp\|Q8BKC5\|IPO5_MOUSE | Importin-5 OS=Mus musculus OX=10090 GN=Ipo5 PE=1 SV=3 |
| Q545A2_MOUSE | MCG11560 OS=Mus musculus OX=10090 GN=Slc25a5 PE=1 SV=1 |
| sp\|Q61753\|SERA_MOUSE D-3-phosphoglycerate dehydrogenase OS=Mus musculus OX=10090... | sp\|Q61753\|SERA_MOUSE D-3-phosphoglycerate dehydrogenase OS=Mus musculus OX=10090... |
| Q3U4D1_MOUSE | Adenosylhomocysteinase OS=Mus musculus OX=10090 PE=2 SV=1 |
| sp\|Q8QZY1\|EIF3L_MOUSEEukaryotic translation initiation factor 3 subunit L OS=Mus musculus OX=10090 GN=Eif3l PE=1 SV=1 | sp\|Q8QZY1\|EIF3L_MOUSE Eukaryotic translation initiation factor 3 subunit L OS=Mus musculus OX=10090 GN=Eif3l PE=1 SV=1 |
| sp\|Q8BFR5\|EFTU_MOUSE Elongation factor Tu, mitochondrial OS=Mus musculus OX=10090... | sp\|Q8BFR5\|EFTU_MOUSE Elongation factor Tu, mitochondrial OS=Mus musculus OX=10090... |
| FAS_MOUSE | Fatty acid synthase OS=Mus musculus OX=10090 GN=Fasn PE=1 SV=2 |
| Q5EBP8_MOUSE | Heterogeneous nuclear ribonucleoprotein A1 OS=Mus musculus OX=10090 GN=Hnrnpa1 PE=1 SV=1 |
| tr\|B2RSV4\|B2RSV4_MOUSE Splicing factor 3b, subunit 3 OS=Mus musculus OX=10090 GN=Sf3b3... | tr\|B2RSV4\|B2RSV4_MOUSE Splicing factor 3b, subunit 3 OS=Mus musculus OX=10090 GN=Sf3b3... |
| CUL1_MOUSE | Cullin-1 OS=Mus musculus OX=10090 GN=Cul1 PE=1 SV=1 |
| Q5SWR1_MOUSE | AP complex subunit beta OS=Mus musculus OX=10090 GN=Ap2b1 PE=2 SV=1 |
| Q8BVQ9_MOUSE | 26S proteasome regulatory subunit 7 OS=Mus musculus OX=10090 GN=Psmc2 PE=1 SV=1 |

| | |
|---|---|
| sp\|Q8VHG2-2\|AMOT_MOUSE | Isoform 2 of Angiomotin OS=Mus musculus OX=10090 GN=Amot |
| tr\|A0JLM6\|A0JLM6_MOUSE Smc1a protein (Fragment) OS=Mus musculus OX=10090 GN=Smc1a... | tr\|A0JLM6\|A0JLM6_MOUSE Smc1a protein (Fragment) OS=Mus musculus OX=10090 GN=Smc1a... |
| tr\|Q3U8U8\|Q3U8U8_MOUSE Polyadenylate-binding protein OS=Mus musculus OX=10090 GN=Pabpc1... | tr\|Q3U8U8\|Q3U8U8_MOUSE Polyadenylate-binding protein OS=Mus musculus OX=10090 GN=Pabpc1... |
| tr\|Q6PCP0\|Q6PCP0_MOUSE DEAD (Asp-Glu-Ala-Asp) box polypeptide 21 OS=Mus musculus... | tr\|Q6PCP0\|Q6PCP0_MOUSE DEAD (Asp-Glu-Ala-Asp) box polypeptide 21 OS=Mus musculus... |
| E9QNN1_MOUSE | ATP-dependent RNA helicase A OS=Mus musculus OX=10090 GN=Dhx9 PE=1 SV=1 |
| tr\|Q3UXQ6\|Q3UXQ6_MOUSE 40S ribosomal protein S4 OS=Mus musculus OX=10090 GN=Rps4x... | 40S ribosomal protein S4 OS=Mus musculus OX=10090 GN=Rps4x PE=2 SV=1 |
| ILF2_MOUSE | Interleukin enhancer-binding factor 2 OS=Mus musculus OX=10090 GN=Ilf2 PE=1 SV=1 |
| Q58ET1_MOUSE | MCG11348 OS=Mus musculus OX=10090 GN=Rpl7a PE=1 SV=1 |
| B2RRX1_MOUSE | Actin, beta OS=Mus musculus OX=10090 GN=Actb PE=2 SV=1 |
| Q5BLK1_MOUSE | 40S ribosomal protein S6 OS=Mus musculus OX=10090 GN=Rps6 PE=1 SV=1 |
| Q6NXX7_MOUSE | Abce1 protein (Fragment) OS=Mus musculus OX=10090 GN=Abce1 PE=2 SV=1 |
| sp\|P14733\|LMNB1_MOUSELamin-B1 OS=Mus musculus OX=10090 GN=Lmnb1 PE=1 SV=3 | sp\|P14733\|LMNB1_MOUSE Lamin-B1 OS=Mus musculus OX=10090 GN=Lmnb1 PE=1 SV=3 |
| sp\|Q61699-2\|HS105_MOUSE Isoform HSP105-beta of Heat shock protein 105 kDa OS=Mus... | sp\|Q61699-2\|HS105_MOUSE Isoform HSP105-beta of Heat shock protein 105 kDa OS=Mus... |
| sp\|Q8BJ71\|NUP93_MOUSENuclear pore complex protein Nup93 OS=Mus musculus OX=10090 GN=Nup93 PE=1 SV=1 | sp\|Q8BJ71\|NUP93_MOUSE Nuclear pore complex protein Nup93 OS=Mus musculus OX=10090 GN=Nup93 PE=1 SV=1 |
| sp\|Q8CAQ8-5\|MIC60_MOUSE Isoform 5 of MICOS complex subunit Mic60 OS=Mus musculus... | sp\|Q8CAQ8-5\|MIC60_MOUSE Isoform 5 of MICOS complex subunit Mic60 OS=Mus musculus... |

| | |
|---|---|
| sp\|Q9ET01\|PYGL_MOUSE Glycogen phosphorylase, liver form OS=Mus musculus OX=10090... | Glycogen phosphorylase, liver form OS=Mus musculus OX=10090 GN=Pygl PE=1 SV=4 |
| tr\|A0A0H3XWX3\|A0A0H3XWX3_MOUSE Insulin-like growth factor 2 mRNA binding protein... | Insulin-like growth factor 2 mRNA binding protein 1 dN CRDBP-2 OS=Mus musculus OX=10090 GN=Igf2bp1 PE=2 SV=1 |
| tr\|Q3UAC2\|Q3UAC2_MOUSE 40S ribosomal protein S3a OS=Mus musculus OX=10090 GN=Rps3a1... | 40S ribosomal protein S3a OS=Mus musculus OX=10090 GN=Rps3a1 PE=2 SV=1 |
| Q3U4P3_MOUSE | Uncharacterized protein (Fragment) OS=Mus musculus OX=10090 GN=Dnm1l PE=2 SV=1 |
| sp\|P62196\|PRS8_MOUSE 26S proteasome regulatory subunit 8 OS=Mus musculus OX=10090... | 26S proteasome regulatory subunit 8 OS=Mus musculus OX=10090 GN=Psmc5 PE=1 SV=1 |
| tr\|B2RRH9\|B2RRH9_MOUSE Guanine monphosphate synthetase OS=Mus musculus OX=10090 GN=Gmps... | Guanine monphosphate synthetase OS=Mus musculus OX=10090 GN=Gmps PE=1 SV=1 |
| Q9Z1R9_MOUSE | MCG124046 OS=Mus musculus OX=10090 GN=Prss1 PE=1 SV=1 |
| A0A0A6YW80_MOUSE | N-alpha-acetyltransferase 15, NatA auxiliary subunit OS=Mus musculus OX=10090 GN=Naa15 PE=1 SV=1 |
| sp\|Q3U0V1\|FUBP2_MOUSEFar upstream element-binding protein 2 OS=Mus musculus OX=10090 GN=Khsrp PE=1 SV=2 | sp\|Q3U0V1\|FUBP2_MOUSE Far upstream element-binding protein 2 OS=Mus musculus OX=10090 GN=Khsrp PE=1 SV=2 |
| sp\|Q8K224\|NAT10_MOUSERNA cytidine acetyltransferase OS=Mus musculus OX=10090 GN=Nat10 PE=1 SV=1 | sp\|Q8K224\|NAT10_MOUSE RNA cytidine acetyltransferase OS=Mus musculus OX=10090 GN=Nat10 PE=1 SV=1 |
| sp\|Q8R1B4\|EIF3C_MOUSEEukaryotic translation initiation factor 3 subunit C OS=Mus musculus OX=10090 GN=Eif3c PE=1 SV=1 | sp\|Q8R1B4\|EIF3C_MOUSE Eukaryotic translation initiation factor 3 subunit C OS=Mus musculus OX=10090 GN=Eif3c PE=1 SV=1 |
| sp\|Q9D0R2\|SYTC_MOUSE Threonine--tRNA ligase, cytoplasmic OS=Mus musculus OX=10090... | Threonine--tRNA ligase, cytoplasmic OS=Mus musculus OX=10090 GN=Tars PE=1 SV=2 |
| sp\|Q9DBR1\|XRN2_MOUSE | 5'-3' exoribonuclease 2 OS=Mus musculus OX=10090 GN=Xrn2 PE=1 SV=1 |
| sp\|Q9JIF7\|COPB_MOUSE Coatomer subunit beta OS=Mus musculus OX=10090 GN=Copb1 PE=1... | sp\|Q9JIF7\|COPB_MOUSE Coatomer subunit beta OS=Mus musculus OX=10090 GN=Copb1 PE=1... |

| | |
|---|---|
| tr\|E9Q3M3\|E9Q3M3_MOUSE Dynactin subunit 1 OS=Mus musculus OX=10090 GN=Dctn1 PE=1... | Dynactin subunit 1 OS=Mus musculus OX=10090 GN=Dctn1 PE=1 SV=1 |
| E9QKZ2_MOUSE | Importin-9 OS=Mus musculus OX=10090 GN=Ipo9 PE=1 SV=1 |
| Q3TK29_MOUSE | Uncharacterized protein OS=Mus musculus OX=10090 GN=Trap1 PE=2 SV=1 |
| Q3U2G2_MOUSE | Heat shock 70 kDa protein 4 OS=Mus musculus OX=10090 GN=Hspa4 PE=1 SV=1 |
| Q3ULS2_MOUSE | Structural maintenance of chromosomes protein OS=Mus musculus OX=10090 GN=Smc2 PE=2 SV=1 |
| sp\|P68040\|RACK1_MOUSEReceptor of activated protein C kinase 1 OS=Mus musculus OX=10090 GN=Rack1 PE=1 SV=3 | sp\|P68040\|RACK1_MOUSE Receptor of activated protein C kinase 1 OS=Mus musculus OX=10090 GN=Rack1 PE=1 SV=3 |
| sp\|Q02053\|UBA1_MOUSE Ubiquitin-like modifier-activating enzyme 1 OS=Mus musculus... | sp\|Q02053\|UBA1_MOUSE Ubiquitin-like modifier-activating enzyme 1 OS=Mus musculus... |
| tr\|B2M1R7\|B2M1R7_MOUSE Poly(RC) binding protein 2 OS=Mus musculus OX=10090 GN=Pcbp2... | tr\|B2M1R7\|B2M1R7_MOUSE Poly(RC) binding protein 2 OS=Mus musculus OX=10090 GN=Pcbp2... |
| tr\|E9Q1S3\|E9Q1S3_MOUSE Protein transport protein Sec23A OS=Mus musculus OX=10090... | Protein transport protein Sec23A OS=Mus musculus OX=10090 GN=Sec23a PE=1 SV=1 |
| tr\|Q5YLW3\|Q5YLW3_MOUSE Ribosomal protein S3 OS=Mus musculus OX=10090 GN=Rps3 PE=1... | tr\|Q5YLW3\|Q5YLW3_MOUSE Ribosomal protein S3 OS=Mus musculus OX=10090 GN=Rps3 PE=1... |
| MTA2_MOUSE | Metastasis-associated protein MTA2 OS=Mus musculus OX=10090 GN=Mta2 PE=1 SV=1 |
| Q3U1S6_MOUSE | Uncharacterized protein OS=Mus musculus OX=10090 GN=Csde1 PE=1 SV=1 |
| Q8CIJ3_MOUSE | Eukaryotic translation initiation factor 3 subunit B OS=Mus musculus OX=10090 GN=Eif3b PE=2 SV=1 |
| TOP1_MOUSE | DNA topoisomerase 1 OS=Mus musculus OX=10090 GN=Top1 PE=1 SV=2 |
| tr\|A1A4T2\|A1A4T2_MOUSE Alpha glucosidase 2 alpha neutral subunit OS=Mus musculus... | tr\|A1A4T2\|A1A4T2_MOUSE Alpha glucosidase 2 alpha neutral subunit OS=Mus musculus... |
| E9PV04_MOUSE | Predicted gene 8994 OS=Mus musculus OX=10090 GN=Gm8994 PE=3 SV=2 |

| | |
|---|---|
| MOES_MOUSE | Moesin OS=Mus musculus OX=10090 GN=Msn PE=1 SV=3 |
| Q3TH47_MOUSE | Uncharacterized protein OS=Mus musculus OX=10090 GN=Ubc PE=2 SV=1 |
| Q3TKG4_MOUSE | Uncharacterized protein (Fragment) OS=Mus musculus OX=10090 GN=Psmc3 PE=2 SV=1 |
| Q546G4_MOUSE | Albumin 1 OS=Mus musculus OX=10090 GN=Alb PE=1 SV=1 |
| Q58EU6_MOUSE | MCG13589 OS=Mus musculus OX=10090 GN=Rpl5 PE=1 SV=1 |
| SYRC_MOUSE | Arginine--tRNA ligase, cytoplasmic OS=Mus musculus OX=10090 GN=Rars PE=1 SV=2 |
| sp\|Q76MZ3\|2AAA_MOUSE Serine/threonine-protein phosphatase 2A 65 kDa regulatory subunit... | sp\|Q76MZ3\|2AAA_MOUSE Serine/threonine-protein phosphatase 2A 65 kDa regulatory subunit... |
| tr\|E9PVC5\|E9PVC5_MOUSE Eukaryotic translation initiation factor 4 gamma 1 OS=Mus... | Eukaryotic translation initiation factor 4 gamma 1 OS=Mus musculus OX=10090 GN=Eif4g1 PE=1 SV=1 |
| E9PXY1_MOUSE | Cullin-4B OS=Mus musculus OX=10090 GN=Cul4b PE=1 SV=2 |
| EF1G_MOUSE | Elongation factor 1-gamma OS=Mus musculus OX=10090 GN=Eef1g PE=1 SV=3 |
| KINH_MOUSE | Kinesin-1 heavy chain OS=Mus musculus OX=10090 GN=Kif5b PE=1 SV=3 |
| Q5XJF6_MOUSE | Ribosomal protein OS=Mus musculus OX=10090 GN=Rpl10a PE=1 SV=1 |
| Q71LX8_MOUSE | Heat shock protein 84b OS=Mus musculus OX=10090 GN=Hsp90ab1 PE=1 SV=1 |
| sp\|Q91YR7\|PRP6_MOUSE | Pre-mRNA-processing factor 6 OS=Mus musculus OX=10090 GN=Prpf6 PE=1 SV=1 |
| sp\|Q9CZN7-2\|GLYM_MOUSE Isoform 2 of Serine hydroxymethyltransferase, mitochondrial... | sp\|Q9CZN7-2\|GLYM_MOUSE Isoform 2 of Serine hydroxymethyltransferase, mitochondrial... |
| F8WHR6_MOUSE | Probable ATP-dependent RNA helicase DDX46 OS=Mus musculus OX=10090 GN=Ddx46 PE=1 SV=1 |
| G3UXL2_MOUSE | Phosphoribosyl pyrophosphate synthetase 1-like 3 OS=Mus musculus OX=10090 GN=Prps1l3 PE=3 SV=1 |

| | |
|---|---|
| IPO7_MOUSE | Importin-7 OS=Mus musculus OX=10090 GN=Ipo7 PE=1 SV=2 |
| Q3TIY6_MOUSE | Rab GDP dissociation inhibitor OS=Mus musculus OX=10090 GN=Gdi2 PE=2 SV=1 |
| Q9CZI7_MOUSE | Annexin OS=Mus musculus OX=10090 GN=Anxa2 PE=2 SV=1 |
| Q3UXP2_MOUSE | RuvB-like helicase OS=Mus musculus OX=10090 GN=Ruvbl2 PE=2 SV=1 |
| GFP | GFP |
| PRP8_MOUSE | Pre-mRNA-processing-splicing factor 8 OS=Mus musculus OX=10090 GN=Prpf8 PE=1 SV=2 |
| ECM29_MOUSE | Isoform 2 of Proteasome adapter and scaffold protein ECM29 OS=Mus musculus OX=10090 GN=Ecpas |
| Q99JX6_MOUSE | Annexin OS=Mus musculus OX=10090 GN=Anxa6 PE=2 SV=1 |
| sp\|Q99KP6-2\|PRP19_MOUSE Isoform 2 of Pre-mRNA-processing factor 19 OS=Mus musculus... | sp\|Q99KP6-2\|PRP19_MOUSE Isoform 2 of Pre-mRNA-processing factor 19 OS=Mus musculus... |
| tr\|Q8QZS9\|Q8QZS9_MOUSE Psmc6 protein (Fragment) OS=Mus musculus OX=10090 GN=Psmc6... | Psmc6 protein (Fragment) OS=Mus musculus OX=10090 GN=Psmc6 PE=2 SV=1 |
| G5E924_MOUSE | Heterogeneous nuclear ribonucleoprotein L (Fragment) OS=Mus musculus OX=10090 GN=Hnrnpl PE=1 SV=1 |
| Q3TJG1_MOUSE | Platelet-activating factor acetylhydrolase IB subunit alpha OS=Mus musculus OX=10090 GN=Pafah1b1 PE=2 SV=1 |
| Q3UJS2_MOUSE | Uncharacterized protein OS=Mus musculus OX=10090 GN=Fbl PE=2 SV=1 |
| Q8BNS0_MOUSE | Uncharacterized protein OS=Mus musculus OX=10090 GN=Drg1 PE=2 SV=1 |
| sp\|P70698\|PYRG1_MOUSECTP synthase 1 OS=Mus musculus OX=10090 GN=Ctps1 PE=1 SV=2 | sp\|P70698\|PYRG1_MOUSE CTP synthase 1 OS=Mus musculus OX=10090 GN=Ctps1 PE=1 SV=2 |
| DESP_MOUSE | Desmoplakin OS=Mus musculus OX=10090 GN=Dsp PE=1 SV=1 |
| GARS_MOUSE | Glycine--tRNA ligase OS=Mus musculus OX=10090 GN=Gars PE=1 SV=1 |
| KCRB_MOUSE | Creatine kinase B-type OS=Mus musculus OX=10090 GN=Ckb PE=1 SV=1 |

| | |
|---|---|
| Q569X4_MOUSE | Proteasome (Prosome, macropain) 26S subunit, ATPase, 4 OS=Mus musculus OX=10090 GN=Psmc4 PE=2 SV=1 |
| Q5CZY9_MOUSE | Rps16 protein OS=Mus musculus OX=10090 GN=Rps16 PE=2 SV=1 |
| sp\|O70194\|EIF3D_MOUSEEukaryotic translation initiation factor 3 subunit D OS=Mus musculus OX=10090 GN=Eif3d PE=1 SV=2 | sp\|O70194\|EIF3D_MOUSE Eukaryotic translation initiation factor 3 subunit D OS=Mus musculus OX=10090 GN=Eif3d PE=1 SV=2 |
| sp\|Q925I1-2\|ATAD3_MOUSE Isoform 2 of ATPase family AAA domain-containing protein... | Isoform 2 of ATPase family AAA domain-containing protein 3 OS=Mus musculus OX=10090 GN=Atad3 |
| sp\|Q9ESZ8-2\|GTF2I_MOUSE Isoform 2 of General transcription factor II-I OS=Mus musculus... | Isoform 2 of General transcription factor II-I OS=Mus musculus OX=10090 GN=Gtf2i |
| tr\|D3YZ35\|D3YZ35_MOUSE Neurofilament medium polypeptide OS=Mus musculus OX=10090... | Neurofilament medium polypeptide OS=Mus musculus OX=10090 GN=Nefm PE=1 SV=1 |
| tr\|Q69ZW4\|Q69ZW4_MOUSE MKIAA0899 protein (Fragment) OS=Mus musculus OX=10090 GN=Ap2a2... | MKIAA0899 protein (Fragment) OS=Mus musculus OX=10090 GN=Ap2a2 PE=2 SV=1 |
| TCPE_MOUSE | T-complex protein 1 subunit epsilon OS=Mus musculus OX=10090 GN=Cct5 PE=1 SV=1 |
| B2RXR3_MOUSE | Cnot1 protein OS=Mus musculus OX=10090 GN=Cnot1 PE=2 SV=1 |
| Q80WV6_MOUSE | Prmt1 protein OS=Mus musculus OX=10090 GN=Prmt1 PE=2 SV=1 |
| sp\|Q64737\|PUR2_MOUSE Trifunctional purine biosynthetic protein adenosine-3 OS=Mus... | sp\|Q64737\|PUR2_MOUSE Trifunctional purine biosynthetic protein adenosine-3 OS=Mus... |
| sp\|Q6PGB8-2\|SMCA1_MOUSE Isoform 2 of Probable global transcription activator SNF2L1... | Isoform 2 of Probable global transcription activator SNF2L1 OS=Mus musculus OX=10090 GN=Smarca1 |
| sp\|Q9JKR6\|HYOU1_MOUSEHypoxia up-regulated protein 1 OS=Mus musculus OX=10090 GN=Hyou1 PE=1 SV=1 | sp\|Q9JKR6\|HYOU1_MOUSE Hypoxia up-regulated protein 1 OS=Mus musculus OX=10090 GN=Hyou1 PE=1 SV=1 |
| sp\|Q9QXS1-3\|PLEC_MOUSE | Isoform PLEC-1A of Plectin OS=Mus musculus OX=10090 GN=Plec |
| tr\|B1AUX2\|B1AUX2_MOUSE Host cell factor 1 OS=Mus musculus OX=10090 GN=Hcfc1 PE=1... | Host cell factor 1 OS=Mus musculus OX=10090 GN=Hcfc1 PE=1 SV=1 |

| | |
|---|---|
| tr\|Q3UDI8\|Q3UDI8_MOUSE DNA replication licensing factor MCM7 OS=Mus musculus OX=10090... | DNA replication licensing factor MCM7 OS=Mus musculus OX=10090 GN=Mcm7 PE=2 SV=1 |
| tr\|Q3UYQ4\|Q3UYQ4_MOUSE Uncharacterized protein OS=Mus musculus OX=10090 GN=Api5 PE=2... | Uncharacterized protein OS=Mus musculus OX=10090 GN=Api5 PE=2 SV=1 |
| A2A513_MOUSE | Keratin, type I cytoskeletal 10 OS=Mus musculus OX=10090 GN=Krt10 PE=1 SV=1 |
| B2RXM2_MOUSE | EG627828 protein OS=Mus musculus OX=10090 GN=Gm6793 PE=1 SV=1 |
| G3UVV4_MOUSE | Hexokinase 1, isoform CRA_f OS=Mus musculus OX=10090 GN=Hk1 PE=1 SV=1 |
| PGM1_MOUSE | Phosphoglucomutase-1 OS=Mus musculus OX=10090 GN=Pgm1 PE=1 SV=4 |
| PUR9_MOUSE | Bifunctional purine biosynthesis protein PURH OS=Mus musculus OX=10090 GN=Atic PE=1 SV=2 |
| Q3TLE5_MOUSE | Uncharacterized protein OS=Mus musculus OX=10090 GN=Rps2 PE=2 SV=1 |
| Q3UBI6_MOUSE | Uncharacterized protein OS=Mus musculus OX=10090 GN=Rpl7 PE=2 SV=1 |
| Q91VZ1_MOUSE | Snx2 protein OS=Mus musculus OX=10090 GN=Snx2 PE=2 SV=1 |
| sp\|Q810A7-2\|DDX42_MOUSE Isoform 2 of ATP-dependent RNA helicase DDX42 OS=Mus musculus... | sp\|Q810A7-2\|DDX42_MOUSE Isoform 2 of ATP-dependent RNA helicase DDX42 OS=Mus musculus... |
| sp\|Q91ZW3\|SMCA5_MOUSESWI/SNF-related matrix-associated actin-dependent regulator of chromatin subfamily A member 5 OS=Mus musculus OX=10090 GN=Smarca5 PE=1 SV=1 | sp\|Q91ZW3\|SMCA5_MOUSE SWI/SNF-related matrix-associated actin-dependent regulator of chromatin subfamily A member 5 OS=Mus musculus OX=10090 GN=Smarca5 PE=1 SV=1 |
| sp\|Q9Z1Z0\|USO1_MOUSE | General vesicular transport factor p115 OS=Mus musculus OX=10090 GN=Uso1 PE=1 SV=2 |
| sp\|O55029\|COPB2_MOUSECoatomer subunit beta' OS=Mus musculus OX=10090 GN=Copb2 PE=1 SV=2 | sp\|O55029\|COPB2_MOUSE Coatomer subunit beta' OS=Mus musculus OX=10090 GN=Copb2 PE=1 SV=2 |
| SYIC_MOUSE | Isoleucine--tRNA ligase, cytoplasmic OS=Mus musculus OX=10090 GN=Iars PE=1 SV=2 |

| | |
|---|---|
| E9Q2X6_MOUSE | Structural maintenance of chromosomes protein OS=Mus musculus OX=10090 GN=Smc4 PE=1 SV=1 |
| Q3THJ6_MOUSE | Uncharacterized protein OS=Mus musculus OX=10090 PE=2 SV=1 |
| Q3TIA9_MOUSE | Uncharacterized protein OS=Mus musculus OX=10090 GN=Asns PE=2 SV=1 |
| Q3TJ43_MOUSE | Vacuolar protein sorting-associated protein 35 OS=Mus musculus OX=10090 GN=Vps35 PE=2 SV=1 |
| sp\|P62830\|RL23_MOUSE 60S ribosomal protein L23 OS=Mus musculus OX=10090 GN=Rpl23... | sp\|P62830\|RL23_MOUSE 60S ribosomal protein L23 OS=Mus musculus OX=10090 GN=Rpl23... |
| tr\|E9Q3D6\|E9Q3D6_MOUSE Heat shock protein HSP 90-beta (Fragment) OS=Mus musculus... | tr\|E9Q3D6\|E9Q3D6_MOUSE Heat shock protein HSP 90-beta (Fragment) OS=Mus musculus... |
| tr\|Q8BU34\|Q8BU34_MOUSE Uncharacterized protein (Fragment) OS=Mus musculus OX=10090... | Uncharacterized protein (Fragment) OS=Mus musculus OX=10090 GN=Psmd1 PE=2 SV=2 |
| Q8BZY3_MOUSE | DEAD (Asp-Glu-Ala-Asp) box polypeptide 19b OS=Mus musculus OX=10090 GN=Ddx19b PE=1 SV=1 |
| PHB_MOUSE | Prohibitin OS=Mus musculus OX=10090 GN=Phb PE=1 SV=1 |
| Q3ULT2_MOUSE | Actinin alpha 4 OS=Mus musculus OX=10090 GN=Actn4 PE=1 SV=1 |
| Q5M9L9_MOUSE | 40S ribosomal protein S8 OS=Mus musculus OX=10090 GN=Rps8 PE=2 SV=1 |
| sp\|P06745\|G6PI_MOUSE Glucose-6-phosphate isomerase OS=Mus musculus OX=10090 GN=Gpi... | sp\|P06745\|G6PI_MOUSE Glucose-6-phosphate isomerase OS=Mus musculus OX=10090 GN=Gpi... |
| sp\|P70388-2\|RAD50_MOUSE | Isoform 2 of DNA repair protein RAD50 OS=Mus musculus OX=10090 GN=Rad50 |
| tr\|Q3UIQ2\|Q3UIQ2_MOUSE Uncharacterized protein OS=Mus musculus OX=10090 GN=Ndufs1... | Uncharacterized protein OS=Mus musculus OX=10090 GN=Ndufs1 PE=2 SV=1 |
| tr\|Q3UJS0\|Q3UJS0_MOUSE Uncharacterized protein OS=Mus musculus OX=10090 GN=Rpl8 PE=2... | tr\|Q3UJS0\|Q3UJS0_MOUSE Uncharacterized protein OS=Mus musculus OX=10090 GN=Rpl8 PE=2... |
| tr\|Q5M8R8\|Q5M8R8_MOUSE 60S acidic ribosomal protein P0 OS=Mus musculus OX=10090 GN=Rplp0... | 60S acidic ribosomal protein P0 OS=Mus musculus OX=10090 GN=Rplp0 PE=2 SV=1 |

| | |
|---|---|
| tr\|Q91ZH2\|Q91ZH2_MOUSE Proliferating cell nuclear antigen OS=Mus musculus OX=10090... | tr\|Q91ZH2\|Q91ZH2_MOUSE Proliferating cell nuclear antigen OS=Mus musculus OX=10090... |
| Q08EK4_MOUSE | Keratin 77 OS=Mus musculus OX=10090 GN=Krt77 PE=2 SV=1 |
| Q3TWJ4_MOUSE | Uncharacterized protein OS=Mus musculus OX=10090 GN=Actr3 PE=2 SV=1 |
| Q3UJZ7_MOUSE | Uncharacterized protein OS=Mus musculus OX=10090 GN=Nop56 PE=2 SV=1 |
| Q8BK73_MOUSE | Uncharacterized protein OS=Mus musculus OX=10090 GN=Psmd11 PE=2 SV=1 |
| sp\|P54071\|IDHP_MOUSE Isocitrate dehydrogenase [NADP], mitochondrial OS=Mus musculus... | Isocitrate dehydrogenase [NADP], mitochondrial OS=Mus musculus OX=10090 GN=Idh2 PE=1 SV=3 |
| sp\|P60335\|PCBP1_MOUSEPoly(rC)-binding protein 1 OS=Mus musculus OX=10090 GN=Pcbp1 PE=1 SV=1 | sp\|P60335\|PCBP1_MOUSE Poly(rC)-binding protein 1 OS=Mus musculus OX=10090 GN=Pcbp1 PE=1 SV=1 |
| sp\|Q8C7X2-2\|EMC1_MOUSE Isoform 2 of ER membrane protein complex subunit 1 OS=Mus... | Isoform 2 of ER membrane protein complex subunit 1 OS=Mus musculus OX=10090 GN=Emc1 |
| sp\|Q99MR6\|SRRT_MOUSE | Serrate RNA effector molecule homolog OS=Mus musculus OX=10090 GN=Srrt PE=1 SV=1 |
| sp\|Q99PU8-2\|DHX30_MOUSE Isoform 2 of Putative ATP-dependent RNA helicase DHX30 OS=Mus... | Isoform 2 of Putative ATP-dependent RNA helicase DHX30 OS=Mus musculus OX=10090 GN=Dhx30 |
| sp\|Q9QXX4\|CMC2_MOUSE Calcium-binding mitochondrial carrier protein Aralar2 OS=Mus... | sp\|Q9QXX4\|CMC2_MOUSE Calcium-binding mitochondrial carrier protein Aralar2 OS=Mus... |
| tr\|A0A0R4J008\|A0A0R4J008_MOUSE Histone deacetylase OS=Mus musculus OX=10090 GN=Hdac2... | tr\|A0A0R4J008\|A0A0R4J008_MOUSE Histone deacetylase OS=Mus musculus OX=10090 GN=Hdac2... |
| tr\|A0A0R4J0K2\|A0A0R4J0K2_MOUSE Cytoskeleton-associated protein 5 OS=Mus musculus... | tr\|A0A0R4J0K2\|A0A0R4J0K2_MOUSE Cytoskeleton-associated protein 5 OS=Mus musculus... |
| tr\|A0JNY7\|A0JNY7_MOUSE Eukaryotic translation initiation factor 4, gamma 2 OS=Mus... | Eukaryotic translation initiation factor 4, gamma 2 OS=Mus musculus OX=10090 GN=Eif4g2 PE=2 SV=1 |
| tr\|Q3UKX8\|Q3UKX8_MOUSE Uncharacterized protein OS=Mus musculus OX=10090 GN=Ap1g1... | Uncharacterized protein OS=Mus musculus OX=10090 GN=Ap1g1 PE=2 SV=1 |

| | |
|---|---|
| tr\|Q8BTX6\|Q8BTX6_MOUSE Uncharacterized protein (Fragment) OS=Mus musculus OX=10090... | tr\|Q8BTX6\|Q8BTX6_MOUSE Uncharacterized protein (Fragment) OS=Mus musculus OX=10090... |
| Q3UVI9_MOUSE | Uncharacterized protein OS=Mus musculus OX=10090 GN=Abcf2 PE=2 SV=1 |
| sp\|Q9CU62\|SMC1A_MOUSEStructural maintenance of chromosomes protein 1A OS=Mus musculus OX=10090 GN=Smc1a PE=1 SV=4 | sp\|Q9CU62\|SMC1A_MOUSE Structural maintenance of chromosomes protein 1A OS=Mus musculus OX=10090 GN=Smc1a PE=1 SV=4 |
| Q3U5M8_MOUSE | Uncharacterized protein OS=Mus musculus OX=10090 GN=Psmd3 PE=2 SV=1 |
| Q5SW83_MOUSE | ARP2 actin-related protein 2 homolog (Yeast) OS=Mus musculus OX=10090 GN=Actr2 PE=1 SV=1 |
| Q80V79_MOUSE | DNA mismatch repair protein OS=Mus musculus OX=10090 GN=Msh2 PE=2 SV=1 |
| RPN1_MOUSE | Dolichyl-diphosphooligosaccharide--protein glycosyltransferase subunit 1 OS=Mus musculus OX=10090 GN=Rpn1 PE=1 SV=1 |
| sp\|P47857-3\|PFKAM_MOUSE | Isoform 3 of ATP-dependent 6-phosphofructokinase, muscle type OS=Mus musculus OX=10090 GN=Pfkm |
| sp\|Q5XJY5\|COPD_MOUSE Coatomer subunit delta OS=Mus musculus OX=10090 GN=Arcn1 PE=1... | sp\|Q5XJY5\|COPD_MOUSE Coatomer subunit delta OS=Mus musculus OX=10090 GN=Arcn1 PE=1... |
| tr\|A2A547\|A2A547_MOUSE Ribosomal protein L19 OS=Mus musculus OX=10090 GN=Rpl19 PE=1... | Ribosomal protein L19 OS=Mus musculus OX=10090 GN=Rpl19 PE=1 SV=1 |
| A0A0R4J119_MOUSE | Cytoplasmic FMR1-interacting protein 1 OS=Mus musculus OX=10090 GN=Cyfip1 PE=1 SV=1 |
| IF2H_MOUSE | Eukaryotic translation initiation factor 2 subunit 3, Y-linked OS=Mus musculus OX=10090 GN=Eif2s3y PE=1 SV=2 |
| Q3TGQ3_MOUSE | Uncharacterized protein OS=Mus musculus OX=10090 GN=Ctnna1 PE=2 SV=1 |
| Q80X48_MOUSE | Nucleoporin 155 OS=Mus musculus OX=10090 GN=Nup155 PE=2 SV=1 |
| RS11_MOUSE | 40S ribosomal protein S11 OS=Mus musculus OX=10090 GN=Rps11 PE=1 SV=3 |

| | |
|---|---|
| sp\|Q8BMJ2\|SYLC_MOUSE Leucine--tRNA ligase, cytoplasmic OS=Mus musculus OX=10090 GN=Lars... | sp\|Q8BMJ2\|SYLC_MOUSE Leucine--tRNA ligase, cytoplasmic OS=Mus musculus OX=10090 GN=Lars... |
| tr\|A0A0A0MQ76\|A0A0A0MQ76_MOUSE Nucleolar protein 58 OS=Mus musculus OX=10090 GN=Nop58... | tr\|A0A0A0MQ76\|A0A0A0MQ76_MOUSE Nucleolar protein 58 OS=Mus musculus OX=10090 GN=Nop58... |
| tr\|A2AJ72\|A2AJ72_MOUSE Far upstream element (FUSE)-binding protein 3 OS=Mus musculus... | Far upstream element (FUSE)-binding protein 3 OS=Mus musculus OX=10090 GN=Fubp3 PE=1 SV=1 |
| tr\|Q3UAG2\|Q3UAG2_MOUSE 6-phosphogluconate dehydrogenase, decarboxylating OS=Mus musculus... | 6-phosphogluconate dehydrogenase, decarboxylating OS=Mus musculus OX=10090 GN=Pgd PE=2 SV=1 |
| tr\|Q3ULW0\|Q3ULW0_MOUSE GTP-binding nuclear protein Ran OS=Mus musculus OX=10090 GN=Ran... | GTP-binding nuclear protein Ran OS=Mus musculus OX=10090 GN=Ran PE=2 SV=1 |
| tr\|Q3UZT7\|Q3UZT7_MOUSE Uncharacterized protein OS=Mus musculus OX=10090 GN=Ctnnb1... | tr\|Q3UZT7\|Q3UZT7_MOUSE Uncharacterized protein OS=Mus musculus OX=10090 GN=Ctnnb1... |
| tr\|Q5FW97\|Q5FW97_MOUSE Enolase 1, alpha non-neuron OS=Mus musculus OX=10090 GN=EG433182... | tr\|Q5FW97\|Q5FW97_MOUSE Enolase 1, alpha non-neuron OS=Mus musculus OX=10090 GN=EG433182... |
| A3KML3_MOUSE | MCG126220 OS=Mus musculus OX=10090 GN=Ywhaq PE=2 SV=1 |
| Q3TG12_MOUSE | Uncharacterized protein (Fragment) OS=Mus musculus OX=10090 GN=Farsb PE=2 SV=1 |
| Q3TMJ8_MOUSE | Mitogen activated protein kinase kinase 1 OS=Mus musculus OX=10090 GN=Map2k1 PE=1 SV=1 |
| Q3U292_MOUSE | Uncharacterized protein OS=Mus musculus OX=10090 GN=Hist1h1d PE=2 SV=1 |
| Q3U8F5_MOUSE | Uncharacterized protein OS=Mus musculus OX=10090 GN=Psmd12 PE=2 SV=1 |
| Q497D7_MOUSE | Rpl30 protein OS=Mus musculus OX=10090 GN=Rpl30 PE=2 SV=1 |
| sp\|Q3UKJ7\|SMU1_MOUSE WD40 repeat-containing protein SMU1 OS=Mus musculus OX=10090... | sp\|Q3UKJ7\|SMU1_MOUSE WD40 repeat-containing protein SMU1 OS=Mus musculus OX=10090... |
| sp\|Q8CGK3\|LONM_MOUSE Lon protease homolog, mitochondrial OS=Mus musculus OX=10090... | sp\|Q8CGK3\|LONM_MOUSE Lon protease homolog, mitochondrial OS=Mus musculus OX=10090... |

| | |
|---|---|
| sp\|Q9ESE1\|LRBA_MOUSE | Lipopolysaccharide-responsive and beige-like anchor protein OS=Mus musculus OX=10090 GN=Lrba PE=1 SV=1 |
| tr\|A0A1D5RLM8\|A0A1D5RLM8_MOUSE Predicted gene 11639 OS=Mus musculus OX=10090 GN=Gm11639... | Predicted gene 11639 OS=Mus musculus OX=10090 GN=Gm11639 PE=1 SV=1 |
| tr\|G3UX26\|G3UX26_MOUSE Voltage-dependent anion-selective channel protein 2 OS=Mus... | tr\|G3UX26\|G3UX26_MOUSE Voltage-dependent anion-selective channel protein 2 OS=Mus... |
| tr\|Q3TWH2\|Q3TWH2_MOUSE Uncharacterized protein OS=Mus musculus OX=10090 GN=Nnt PE=2... | Uncharacterized protein OS=Mus musculus OX=10090 GN=Nnt PE=2 SV=1 |
| tr\|Q3UAT9\|Q3UAT9_MOUSE Inosine-5'-monophosphate dehydrogenase OS=Mus musculus OX=10090... | tr\|Q3UAT9\|Q3UAT9_MOUSE Inosine-5'-monophosphate dehydrogenase OS=Mus musculus OX=10090... |
| tr\|Q8BK18\|Q8BK18_MOUSE Uncharacterized protein OS=Mus musculus OX=10090 GN=Dars PE=2... | tr\|Q8BK18\|Q8BK18_MOUSE Uncharacterized protein OS=Mus musculus OX=10090 GN=Dars PE=2... |
| tr\|Q91V55\|Q91V55_MOUSE 40S ribosomal protein S5 OS=Mus musculus OX=10090 GN=Rps5... | tr\|Q91V55\|Q91V55_MOUSE 40S ribosomal protein S5 OS=Mus musculus OX=10090 GN=Rps5... |
| A0A1B0GRU8_MOUSE | Pre-mRNA-processing factor 40 homolog A OS=Mus musculus OX=10090 GN=Prpf40a PE=1 SV=1 |
| H2B1P_MOUSE | Isoform 2 of Histone H2B type 1-P OS=Mus musculus OX=10090 GN=Hist1h2bp |
| H31_MOUSE | Histone H3.1 OS=Mus musculus OX=10090 GN=Hist1h3a PE=1 SV=2 |
| A0A286YDW8_MOUSE | 26S proteasome non-ATPase regulatory subunit 6 (Fragment) OS=Mus musculus OX=10090 GN=Psmd6 PE=1 SV=1 |
| B1AT82_MOUSE | MCG6846, isoform CRA_c OS=Mus musculus OX=10090 GN=Prpsap1 PE=1 SV=1 |
| B2RUC7_MOUSE | Serine/threonine kinase receptor associated protein OS=Mus musculus OX=10090 GN=Strap PE=1 SV=1 |
| D3Z2H2_MOUSE | Catenin delta-1 OS=Mus musculus OX=10090 GN=Ctnnd1 PE=1 SV=1 |
| Q3TJ01_MOUSE | tRNA-splicing ligase RtcB homolog OS=Mus musculus OX=10090 GN=Rtcb PE=2 SV=1 |

| | |
|---|---|
| Q3ULI4_MOUSE | Signal transducer and activator of transcription OS=Mus musculus OX=10090 GN=Stat3 PE=2 SV=1 |
| Q99KA2_MOUSE | Tubulin alpha chain (Fragment) OS=Mus musculus OX=10090 GN=Tuba1b PE=2 SV=1 |
| Q99PB4_MOUSE | Mage-d2 OS=Mus musculus OX=10090 GN=Maged2 PE=2 SV=1 |
| sp\|O55201-2\|SPT5H_MOUSE Isoform 2 of Transcription elongation factor SPT5 OS=Mus... | sp\|O55201-2\|SPT5H_MOUSE Isoform 2 of Transcription elongation factor SPT5 OS=Mus... |
| tr\|D3YWT1\|D3YWT1_MOUSE Heterogeneous nuclear ribonucleoprotein H3 OS=Mus musculus... | tr\|D3YWT1\|D3YWT1_MOUSE Heterogeneous nuclear ribonucleoprotein H3 OS=Mus musculus... |
| tr\|D3Z0M9\|D3Z0M9_MOUSE DEAD (Asp-Glu-Ala-Asp) box polypeptide 23 OS=Mus musculus... | DEAD (Asp-Glu-Ala-Asp) box polypeptide 23 OS=Mus musculus OX=10090 GN=Ddx23 PE=1 SV=1 |
| tr\|Q3TII4\|Q3TII4_MOUSE Uncharacterized protein (Fragment) OS=Mus musculus OX=10090... | Uncharacterized protein (Fragment) OS=Mus musculus OX=10090 GN=Vars PE=2 SV=1 |
| tr\|Q3TR62\|Q3TR62_MOUSE DNA topoisomerase 2 (Fragment) OS=Mus musculus OX=10090 GN=Top2b... | tr\|Q3TR62\|Q3TR62_MOUSE DNA topoisomerase 2 (Fragment) OS=Mus musculus OX=10090 GN=Top2b... |
| tr\|Q5M8Q0\|Q5M8Q0_MOUSE Ribosomal protein L15 OS=Mus musculus OX=10090 GN=Rpl15 PE=2... | Ribosomal protein L15 OS=Mus musculus OX=10090 GN=Rpl15 PE=2 SV=1 |
| A0A1W2P7Z8_MOUSE | Predicted gene 9045 OS=Mus musculus OX=10090 GN=Gm9045 PE=4 SV=1 |
| A0A0H2UH17_MOUSE | Ubiquitin associated protein 2-like, isoform CRA_b OS=Mus musculus OX=10090 GN=Ubap2l PE=1 SV=1 |
| A0A1D5RM85_MOUSE | 60S ribosomal protein L18a (Fragment) OS=Mus musculus OX=10090 GN=Rpl18a PE=1 SV=1 |
| A2AAN2_MOUSE | Signal recognition particle subunit SRP68 OS=Mus musculus OX=10090 GN=Srp68 PE=1 SV=1 |
| A2AVJ7_MOUSE | Ribosome-binding protein 1 OS=Mus musculus OX=10090 GN=Rrbp1 PE=1 SV=1 |
| B1AXN9_MOUSE | Non-specific serine/threonine protein kinase OS=Mus musculus OX=10090 GN=Rps6ka3 PE=1 SV=1 |

| | |
|---|---|
| B2RX05_MOUSE | Slit2 protein OS=Mus musculus OX=10090 GN=Slit2 PE=2 SV=1 |
| F8WGL3_MOUSE | Cofilin-1 OS=Mus musculus OX=10090 GN=Cfl1 PE=1 SV=1 |
| O70569_MOUSE | Ribosomal protein S14 OS=Mus musculus OX=10090 GN=rps14 PE=3 SV=1 |
| Q3TQW3_MOUSE | Uncharacterized protein OS=Mus musculus OX=10090 GN=Ptbp1 PE=2 SV=1 |
| Q3U2V0_MOUSE | Uncharacterized protein OS=Mus musculus OX=10090 GN=Sympk PE=2 SV=1 |
| Q3UAJ1_MOUSE | Peptidyl-prolyl cis-trans isomerase OS=Mus musculus OX=10090 GN=Ppia PE=2 SV=1 |
| Q3UXT6_MOUSE | Uncharacterized protein (Fragment) OS=Mus musculus OX=10090 GN=Rbm39 PE=2 SV=1 |
| sp\|P00493\|HPRT_MOUSE Hypoxanthine-guanine phosphoribosyltransferase OS=Mus musculus... | sp\|P00493\|HPRT_MOUSE Hypoxanthine-guanine phosphoribosyltransferase OS=Mus musculus... |
| sp\|P48722-2\|HS74L_MOUSE Isoform 2 of Heat shock 70 kDa protein 4L OS=Mus musculus... | Isoform 2 of Heat shock 70 kDa protein 4L OS=Mus musculus OX=10090 GN=Hspa4l |
| sp\|P97807\|FUMH_MOUSE | Fumarate hydratase, mitochondrial OS=Mus musculus OX=10090 GN=Fh PE=1 SV=3 |
| sp\|Q4VA53-3\|PDS5B_MOUSE | Isoform 3 of Sister chromatid cohesion protein PDS5 homolog B OS=Mus musculus OX=10090 GN=Pds5b |
| sp\|Q6ZWR6-4\|SYNE1_MOUSE | Isoform 4 of Nesprin-1 OS=Mus musculus OX=10090 GN=Syne1 |
| sp\|Q8BTI8-2\|SRRM2_MOUSE Isoform 2 of Serine/arginine repetitive matrix protein 2... | sp\|Q8BTI8-2\|SRRM2_MOUSE Isoform 2 of Serine/arginine repetitive matrix protein 2... |
| sp\|Q99KI0\|ACON_MOUSE Aconitate hydratase, mitochondrial OS=Mus musculus OX=10090... | sp\|Q99KI0\|ACON_MOUSE Aconitate hydratase, mitochondrial OS=Mus musculus OX=10090... |
| sp\|Q9WVA3\|BUB3_MOUSE Mitotic checkpoint protein BUB3 OS=Mus musculus OX=10090 GN=Bub3... | Mitotic checkpoint protein BUB3 OS=Mus musculus OX=10090 GN=Bub3 PE=1 SV=2 |
| tr\|B2CY77\|B2CY77_MOUSE Laminin receptor (Fragment) OS=Mus musculus OX=10090 GN=Rpsa... | tr\|B2CY77\|B2CY77_MOUSE Laminin receptor (Fragment) OS=Mus musculus OX=10090 GN=Rpsa... |
| tr\|Q3U6A2\|Q3U6A2_MOUSE Uncharacterized protein OS=Mus musculus OX=10090 GN=Rbm14... | Uncharacterized protein OS=Mus musculus OX=10090 GN=Rbm14 PE=2 SV=1 |

| | |
|---|---|
| tr\|Q8C605\|Q8C605_MOUSE ATP-dependent 6-phosphofructokinase OS=Mus musculus OX=10090... | tr\|Q8C605\|Q8C605_MOUSE ATP-dependent 6-phosphofructokinase OS=Mus musculus OX=10090... |
| Q6XMP4_MOUSE | Alanyl-tRNA synthase OS=Mus musculus OX=10090 GN=Aars PE=2 SV=1 |
| A0A1S6GWG6_MOUSE | Uncharacterized protein OS=Mus musculus OX=10090 GN=Atp6v1b2 PE=2 SV=1 |
| ACTZ_MOUSE | Alpha-centractin OS=Mus musculus OX=10090 GN=Actr1a PE=1 SV=1 |
| AMPB_MOUSE | Aminopeptidase B OS=Mus musculus OX=10090 GN=Rnpep PE=1 SV=2 |
| B1AXW6_MOUSE | Peroxiredoxin-1 (Fragment) OS=Mus musculus OX=10090 GN=Prdx1 PE=1 SV=1 |
| B2RRI2_MOUSE | Protein strawberry notch homolog 1 OS=Mus musculus OX=10090 GN=Sbno1 PE=1 SV=1 |
| CSK_MOUSE | Tyrosine-protein kinase CSK OS=Mus musculus OX=10090 GN=Csk PE=1 SV=2 |
| E9PUW7_MOUSE | Exportin-7 OS=Mus musculus OX=10090 GN=Xpo7 PE=1 SV=1 |
| Q3TIQ2_MOUSE | Uncharacterized protein OS=Mus musculus OX=10090 GN=Rpl12 PE=2 SV=1 |
| Q3TVM2_MOUSE | Uncharacterized protein OS=Mus musculus OX=10090 GN=Aldh2 PE=2 SV=1 |
| Q3TVN4_MOUSE | AP complex subunit beta OS=Mus musculus OX=10090 GN=Ap1b1 PE=2 SV=1 |
| Q3UDQ7_MOUSE | Uncharacterized protein OS=Mus musculus OX=10090 GN=Afg3l2 PE=2 SV=1 |
| Q3UIX8_MOUSE | Uncharacterized protein OS=Mus musculus OX=10090 GN=Ctbp2 PE=2 SV=1 |
| Q3V141_MOUSE | Uncharacterized protein (Fragment) OS=Mus musculus OX=10090 GN=Papola PE=2 SV=1 |
| Q5BKP5_MOUSE | DEAD (Asp-Glu-Ala-Asp) box polypeptide 50 OS=Mus musculus OX=10090 GN=Ddx50 PE=2 SV=1 |
| Q5SWZ0_MOUSE | Prpsap2 protein OS=Mus musculus OX=10090 GN=Prpsap2 PE=1 SV=1 |
| Q6IRT4_MOUSE | Eukaryotic translation initiation factor 3 subunit F OS=Mus musculus OX=10090 GN=Eif3f PE=2 SV=1 |

| | |
|---|---|
| Q80YG4_MOUSE | SEC63 OS=Mus musculus OX=10090 GN=Sec63 PE=2 SV=1 |
| Q8C2N0_MOUSE | DNA polymerase OS=Mus musculus OX=10090 GN=Pold1 PE=2 SV=1 |
| Q8CCE5_MOUSE | Uncharacterized protein (Fragment) OS=Mus musculus OX=10090 GN=Top2a PE=2 SV=1 |
| RS20_MOUSE | 40S ribosomal protein S20 OS=Mus musculus OX=10090 GN=Rps20 PE=1 SV=1 |
| Z4YLI8_MOUSE | Clustered mitochondria protein homolog OS=Mus musculus OX=10090 GN=Cluh PE=1 SV=1 |
| sp\|O35593\|PSDE_MOUSE 26S proteasome non-ATPase regulatory subunit 14 OS=Mus musculus... | sp\|O35593\|PSDE_MOUSE 26S proteasome non-ATPase regulatory subunit 14 OS=Mus musculus... |
| sp\|P47757-2\|CAPZB_MOUSE | Isoform 2 of F-actin-capping protein subunit beta OS=Mus musculus OX=10090 GN=Capzb |
| sp\|Q6NZN0-4\|RBM26_MOUSE | Isoform 4 of RNA-binding protein 26 OS=Mus musculus OX=10090 GN=Rbm26 |
| sp\|Q8CIG8\|ANM5_MOUSE Protein arginine N-methyltransferase 5 OS=Mus musculus OX=10090... | sp\|Q8CIG8\|ANM5_MOUSE Protein arginine N-methyltransferase 5 OS=Mus musculus OX=10090... |
| sp\|Q9Z204-4\|HNRPC_MOUSE | Isoform 4 of Heterogeneous nuclear ribonucleoproteins C1/C2 OS=Mus musculus OX=10090 GN=Hnrnpc |
| tr\|E9Q1W3\|E9Q1W3_MOUSE-DECOY Nebulin OS=Mus musculus OX=10090 GN=Neb PE=1 SV=1 | tr\|E9Q1W3\|E9Q1W3_MOUSE-DECOY Nebulin OS=Mus musculus OX=10090 GN=Neb PE=1 SV=1 |
| Q4FZG5_MOUSE | Arp2/3 complex 34 kDa subunit (Fragment) OS=Mus musculus OX=10090 GN=Arpc2 PE=2 SV=1 |
| sp\|Q9QZE5\|COPG1_MOUSECoatomer subunit gamma-1 OS=Mus musculus OX=10090 GN=Copg1 PE=1 SV=1 | sp\|Q9QZE5\|COPG1_MOUSE Coatomer subunit gamma-1 OS=Mus musculus OX=10090 GN=Copg1 PE=1 SV=1 |
| Q91YC8_MOUSE | Damaged-DNA recognition protein 1 OS=Mus musculus OX=10090 GN=Ddb1 PE=2 SV=1 |
| A0A087WR97_MOUSE | TAR DNA-binding protein 43 (Fragment) OS=Mus musculus OX=10090 GN=Tardbp PE=1 SV=1 |

| | |
|---|---|
| A0A2I3BRL0_MOUSE | 5'-nucleotidase domain containing 2 OS=Mus musculus OX=10090 GN=Nt5dc2 PE=1 SV=1 |
| A2ANY6_MOUSE | Midasin OS=Mus musculus OX=10090 GN=Mdn1 PE=1 SV=1 |
| B2RTP7_MOUSE | Krt2 protein OS=Mus musculus OX=10090 GN=Krt2 PE=2 SV=1 |
| E9QP46_MOUSE | Nesprin-2 OS=Mus musculus OX=10090 GN=Syne2 PE=1 SV=1 |
| Q3THH1_MOUSE | Uncharacterized protein OS=Mus musculus OX=10090 GN=Pdia6 PE=2 SV=1 |
| Q3TIJ1_MOUSE | Uncharacterized protein OS=Mus musculus OX=10090 GN=Cops4 PE=2 SV=1 |
| Q3UKP5_MOUSE | Uncharacterized protein OS=Mus musculus OX=10090 GN=Xab2 PE=2 SV=1 |
| Q9QWU1_MOUSE | Hsc70t (Fragment) OS=Mus musculus OX=10090 PE=3 SV=1 |
| RS9_MOUSE | 40S ribosomal protein S9 OS=Mus musculus OX=10090 GN=Rps9 PE=1 SV=3 |
| sp\|P28740-2\|KIF2A_MOUSE Isoform 2 of Kinesin-like protein KIF2A OS=Mus musculus OX=10090... | Isoform 2 of Kinesin-like protein KIF2A OS=Mus musculus OX=10090 GN=Kif2a |
| tr\|E9Q616\|E9Q616_MOUSE AHNAK nucleoprotein (desmoyokin) OS=Mus musculus OX=10090... | tr\|E9Q616\|E9Q616_MOUSE AHNAK nucleoprotein (desmoyokin) OS=Mus musculus OX=10090... |
| tr\|Q3TLE2\|Q3TLE2_MOUSE Pleckstrin homology domain containing, family C (With FERM... | tr\|Q3TLE2\|Q3TLE2_MOUSE Pleckstrin homology domain containing, family C (With FERM... |
| tr\|Q99J57\|Q99J57_MOUSE S-adenosylmethionine synthase OS=Mus musculus OX=10090 GN=Mat2a... | S-adenosylmethionine synthase OS=Mus musculus OX=10090 GN=Mat2a PE=1 SV=1 |
| B1ARU4_MOUSE | Microtubule-actin cross-linking factor 1 OS=Mus musculus OX=10090 GN=Macf1 PE=1 SV=1 |
| sp\|Q62318-2\|TIF1B_MOUSE Isoform 2 of Transcription intermediary factor 1-beta OS=Mus... | Isoform 2 of Transcription intermediary factor 1-beta OS=Mus musculus OX=10090 GN=Trim28 |
| Q3TEU8_MOUSE | Coronin OS=Mus musculus OX=10090 GN=Coro1c PE=2 SV=1 |
| A2A841_MOUSE | Protein 4.1 OS=Mus musculus OX=10090 GN=Epb41 PE=1 SV=1 |

| | |
|---|---|
| sp\|P97313\|PRKDC_MOUSEDNA-dependent protein kinase catalytic subunit OS=Mus musculus OX=10090 GN=Prkdc PE=1 SV=3 | sp\|P97313\|PRKDC_MOUSE DNA-dependent protein kinase catalytic subunit OS=Mus musculus OX=10090 GN=Prkdc PE=1 SV=3 |
| sp\|Q1HFZ0-2\|NSUN2_MOUSE Isoform 2 of tRNA (cytosine(34)-C(5))-methyltransferase OS=Mus... | sp\|Q1HFZ0-2\|NSUN2_MOUSE Isoform 2 of tRNA (cytosine(34)-C(5))-methyltransferase OS=Mus... |
| B2RUG7_MOUSE | Zinc finger RNA binding protein OS=Mus musculus OX=10090 GN=Zfr PE=2 SV=1 |
| E9Q1W0_MOUSE | Calcium/calmodulin-dependent protein kinase type II subunit delta OS=Mus musculus OX=10090 GN=Camk2d PE=1 SV=2 |
| ESYT1_MOUSE | Isoform 2 of Extended synaptotagmin-1 OS=Mus musculus OX=10090 GN=Esyt1 |
| H4_MOUSE | Histone H4 OS=Mus musculus OX=10090 GN=Hist1h4a PE=1 SV=2 |
| Q3TIP8_MOUSE | Chloride intracellular channel protein OS=Mus musculus OX=10090 GN=Clic1 PE=2 SV=1 |
| Q6PHZ1_MOUSE | Rpl17 protein OS=Mus musculus OX=10090 GN=Rpl17 PE=2 SV=1 |
| Q8BP16_MOUSE | Core histone macro-H2A OS=Mus musculus OX=10090 GN=H2afy2 PE=2 SV=1 |
| V9GX76_MOUSE | Unconventional myosin-VI OS=Mus musculus OX=10090 GN=Myo6 PE=1 SV=1 |
| sp\|P62192\|PRS4_MOUSE 26S proteasome regulatory subunit 4 OS=Mus musculus OX=10090... | sp\|P62192\|PRS4_MOUSE 26S proteasome regulatory subunit 4 OS=Mus musculus OX=10090... |
| sp\|Q8VHX6-2\|FLNC_MOUSE | Isoform 2 of Filamin-C OS=Mus musculus OX=10090 GN=Flnc |
| sp\|Q91V12\|BACH_MOUSE Cytosolic acyl coenzyme A thioester hydrolase OS=Mus musculus... | Cytosolic acyl coenzyme A thioester hydrolase OS=Mus musculus OX=10090 GN=Acot7 PE=1 SV=2 |
| sp\|Q9QYB1\|CLIC4_MOUSEChloride intracellular channel protein 4 OS=Mus musculus OX=10090 GN=Clic4 PE=1 SV=3 | sp\|Q9QYB1\|CLIC4_MOUSE Chloride intracellular channel protein 4 OS=Mus musculus OX=10090 GN=Clic4 PE=1 SV=3 |
| tr\|D3YTQ3\|D3YTQ3_MOUSE Heterogeneous nuclear ribonucleoprotein D-like OS=Mus musculus... | tr\|D3YTQ3\|D3YTQ3_MOUSE Heterogeneous nuclear ribonucleoprotein D-like OS=Mus musculus... |

| | |
|---|---|
| tr\|Q20BD0\|Q20BD0_MOUSE Heterogeneous nuclear ribonucleoprotein A/B OS=Mus musculus... | tr\|Q20BD0\|Q20BD0_MOUSE Heterogeneous nuclear ribonucleoprotein A/B OS=Mus musculus... |
| tr\|Q9CTC7\|Q9CTC7_MOUSE Uncharacterized protein (Fragment) OS=Mus musculus OX=10090... | tr\|Q9CTC7\|Q9CTC7_MOUSE Uncharacterized protein (Fragment) OS=Mus musculus OX=10090... |
| QCR1_MOUSE | Cytochrome b-c1 complex subunit 1, mitochondrial OS=Mus musculus OX=10090 GN=Uqcrc1 PE=1 SV=2 |
| B9EKK3_MOUSE | IQ motif containing GTPase activating protein 2 OS=Mus musculus OX=10090 GN=Iqgap2 PE=2 SV=1 |
| sp\|P50516\|VATA_MOUSE V-type proton ATPase catalytic subunit A OS=Mus musculus OX=10090... | sp\|P50516\|VATA_MOUSE V-type proton ATPase catalytic subunit A OS=Mus musculus OX=10090... |
| sp\|P24527\|LKHA4_MOUSELeukotriene A-4 hydrolase OS=Mus musculus OX=10090 GN=Lta4h PE=1 SV=4 | sp\|P24527\|LKHA4_MOUSE Leukotriene A-4 hydrolase OS=Mus musculus OX=10090 GN=Lta4h PE=1 SV=4 |
| Q4VA28_MOUSE | Ribosomal protein L21 OS=Mus musculus OX=10090 GN=Rpl21 PE=2 SV=1 |
| tr\|Q3U4W8\|Q3U4W8_MOUSE Ubiquitinyl hydrolase 1 OS=Mus musculus OX=10090 GN=Usp5 PE=1... | Ubiquitinyl hydrolase 1 OS=Mus musculus OX=10090 GN=Usp5 PE=1 SV=1 |
| Q5FWB7_MOUSE | Fructose-bisphosphate aldolase OS=Mus musculus OX=10090 GN=Aldoa PE=1 SV=1 |
| sp\|Q9JLN9\|MTOR_MOUSE | Serine/threonine-protein kinase mTOR OS=Mus musculus OX=10090 GN=Mtor PE=1 SV=2 |
| sp\|B2RY56\|RBM25_MOUSERNA-binding protein 25 OS=Mus musculus OX=10090 GN=Rbm25 PE=1 SV=2 | sp\|B2RY56\|RBM25_MOUSE RNA-binding protein 25 OS=Mus musculus OX=10090 GN=Rbm25 PE=1 SV=2 |
| sp\|O88477\|IF2B1_MOUSEInsulin-like growth factor 2 mRNA-binding protein 1 OS=Mus musculus OX=10090 GN=Igf2bp1 PE=1 SV=1 | sp\|O88477\|IF2B1_MOUSE Insulin-like growth factor 2 mRNA-binding protein 1 OS=Mus musculus OX=10090 GN=Igf2bp1 PE=1 SV=1 |
| A0A0R4J0I9_MOUSE | Low density lipoprotein receptor-related protein 1 OS=Mus musculus OX=10090 GN=Lrp1 PE=1 SV=1 |
| A1L3S7_MOUSE | Gatad2b protein OS=Mus musculus OX=10090 GN=Gatad2b PE=1 SV=1 |

| | |
|---|---|
| D6RE33_MOUSE | Enhancer of mRNA-decapping protein 4 OS=Mus musculus OX=10090 GN=Edc4 PE=1 SV=1 |
| E9Q6Q8_MOUSE | TBC1 domain family member 4 OS=Mus musculus OX=10090 GN=Tbc1d4 PE=1 SV=1 |
| E9QB02_MOUSE | Methionine--tRNA ligase, cytoplasmic OS=Mus musculus OX=10090 GN=Mars PE=1 SV=1 |
| Q05DU8_MOUSE | Ribonucleoside-diphosphate reductase (Fragment) OS=Mus musculus OX=10090 GN=Rrm1 PE=2 SV=1 |
| Q3TE06_MOUSE | Uncharacterized protein (Fragment) OS=Mus musculus OX=10090 GN=Wdr1 PE=2 SV=1 |
| Q3TKD0_MOUSE | Transportin-1 (Fragment) OS=Mus musculus OX=10090 GN=Tnpo1 PE=1 SV=1 |
| Q3TLX1_MOUSE | Uncharacterized protein OS=Mus musculus OX=10090 GN=Nampt PE=2 SV=1 |
| Q3UMM1_MOUSE | Tubulin beta chain OS=Mus musculus OX=10090 GN=Tubb6 PE=1 SV=1 |
| Q3UNG1_MOUSE | Uncharacterized protein (Fragment) OS=Mus musculus OX=10090 GN=Prpf8 PE=2 SV=1 |
| sp\|P62267\|RS23_MOUSE 40S ribosomal protein S23 OS=Mus musculus OX=10090 GN=Rps23... | sp\|P62267\|RS23_MOUSE 40S ribosomal protein S23 OS=Mus musculus OX=10090 GN=Rps23... |
| sp\|Q91VR5\|DDX1_MOUSE ATP-dependent RNA helicase DDX1 OS=Mus musculus OX=10090 GN=Ddx1... | sp\|Q91VR5\|DDX1_MOUSE ATP-dependent RNA helicase DDX1 OS=Mus musculus OX=10090 GN=Ddx1... |
| sp\|Q9DBE9\|SPB1_MOUSE pre-rRNA processing protein FTSJ3 OS=Mus musculus OX=10090 GN=Ftsj3... | sp\|Q9DBE9\|SPB1_MOUSE pre-rRNA processing protein FTSJ3 OS=Mus musculus OX=10090 GN=Ftsj3... |
| tr\|Q4FJZ2\|Q4FJZ2_MOUSE Importin subunit alpha OS=Mus musculus OX=10090 GN=Kpna6 PE=1... | tr\|Q4FJZ2\|Q4FJZ2_MOUSE Importin subunit alpha OS=Mus musculus OX=10090 GN=Kpna6 PE=1... |
| E9Q828_MOUSE | Calcium-transporting ATPase OS=Mus musculus OX=10090 GN=Atp2b4 PE=1 SV=1 |
| Q3TTV6_MOUSE | Tuftelin-interacting protein 11 OS=Mus musculus OX=10090 GN=Tfip11 PE=1 SV=1 |

| | |
|---|---|
| sp\|O88569\|ROA2_MOUSE | Heterogeneous nuclear ribonucleoproteins A2/B1 OS=Mus musculus OX=10090 GN=Hnrnpa2b1 PE=1 SV=2 |
| sp\|P11103\|PARP1_MOUSEPoly [ADP-ribose] polymerase 1 OS=Mus musculus OX=10090 GN=Parp1 PE=1 SV=3 | sp\|P11103\|PARP1_MOUSE Poly [ADP-ribose] polymerase 1 OS=Mus musculus OX=10090 GN=Parp1 PE=1 SV=3 |
| E9Q0F0_MOUSE | Keratin 78 OS=Mus musculus OX=10090 GN=Krt78 PE=1 SV=1 |
| F8WJ41_MOUSE | 40S ribosomal protein S15a (Fragment) OS=Mus musculus OX=10090 GN=Rps15a PE=1 SV=1 |
| Q3UKV5_MOUSE | Uncharacterized protein (Fragment) OS=Mus musculus OX=10090 GN=Hnrnpr PE=2 SV=1 |
| E9Q715_MOUSE | Putative RNA-binding protein Luc7-like 2 OS=Mus musculus OX=10090 GN=Luc7l2 PE=1 SV=1 |
| Q3TJF2_MOUSE | Obg-like ATPase 1 OS=Mus musculus OX=10090 GN=Ola1 PE=2 SV=1 |
| CUL3_MOUSE | Cullin-3 OS=Mus musculus OX=10090 GN=Cul3 PE=1 SV=1 |
| Z4YL78_MOUSE | Cytoskeleton-associated protein 5 OS=Mus musculus OX=10090 GN=Ckap5 PE=1 SV=1 |
| CISY_MOUSE | Citrate synthase, mitochondrial OS=Mus musculus OX=10090 GN=Cs PE=1 SV=1 |
| F6ZV59_MOUSE | Heterogeneous nuclear ribonucleoprotein D0 (Fragment) OS=Mus musculus OX=10090 GN=Hnrnpd PE=1 SV=1 |
| GDIA_MOUSE | Rab GDP dissociation inhibitor alpha OS=Mus musculus OX=10090 GN=Gdi1 PE=1 SV=3 |
| H3BJW3_MOUSE | Cleavage and polyadenylation-specificity factor subunit 6 OS=Mus musculus OX=10090 GN=Cpsf6 PE=1 SV=1 |
| Q3THL5_MOUSE | Protein pelota homolog OS=Mus musculus OX=10090 GN=Pelo PE=2 SV=1 |
| Q3TNL1_MOUSE | Glucose-6-phosphate 1-dehydrogenase OS=Mus musculus OX=10090 GN=G6pdx PE=2 SV=1 |
| Q6NWW1_MOUSE | Ubiquitination factor E4B, UFD2 homolog (S. cerevisiae) OS=Mus musculus OX=10090 GN=Ube4b PE=2 SV=1 |

| | |
|---|---|
| Q7TMW1_MOUSE | Rangap1 protein OS=Mus musculus OX=10090 GN=Rangap1 PE=2 SV=1 |
| Q80X98_MOUSE | DEAH (Asp-Glu-Ala-His) box polypeptide 38 OS=Mus musculus OX=10090 GN=Dhx38 PE=1 SV=1 |
| Q9CVR0_MOUSE | Tubulin beta chain (Fragment) OS=Mus musculus OX=10090 GN=Tubb4b PE=2 SV=2 |
| RS17_MOUSE | 40S ribosomal protein S17 OS=Mus musculus OX=10090 GN=Rps17 PE=1 SV=2 |
| RS27_MOUSE | 40S ribosomal protein S27 OS=Mus musculus OX=10090 GN=Rps27 PE=1 SV=3 |
| sp\|P04104\|K2C1_MOUSE Keratin, type II cytoskeletal 1 OS=Mus musculus OX=10090 GN=Krt1... | Keratin, type II cytoskeletal 1 OS=Mus musculus OX=10090 GN=Krt1 PE=1 SV=4 |
| sp\|Q60902-4\|EP15R_MOUSE | Isoform 4 of Epidermal growth factor receptor substrate 15-like 1 OS=Mus musculus OX=10090 GN=Eps15l1 |
| sp\|Q6ZWX6\|IF2A_MOUSE Eukaryotic translation initiation factor 2 subunit 1 OS=Mus... | Eukaryotic translation initiation factor 2 subunit 1 OS=Mus musculus OX=10090 GN=Eif2s1 PE=1 SV=3 |
| sp\|Q7M6Y3-2\|PICAL_MOUSE | Isoform 2 of Phosphatidylinositol-binding clathrin assembly protein OS=Mus musculus OX=10090 GN=Picalm |
| sp\|Q8CGY8\|OGT1_MOUSE | UDP-N-acetylglucosamine--peptide N-acetylglucosaminyltransferase 110 kDa subunit OS=Mus musculus OX=10090 GN=Ogt PE=1 SV=2 |
| sp\|Q9Z1F9\|SAE2_MOUSE SUMO-activating enzyme subunit 2 OS=Mus musculus OX=10090 GN=Uba2... | sp\|Q9Z1F9\|SAE2_MOUSE SUMO-activating enzyme subunit 2 OS=Mus musculus OX=10090 GN=Uba2... |
| tr\|A0A0N4SVT1\|A0A0N4SVT1_MOUSE Probable ATP-dependent RNA helicase DDX47 (Fragment)... | tr\|A0A0N4SVT1\|A0A0N4SVT1_MOUSE Probable ATP-dependent RNA helicase DDX47 (Fragment)... |
| tr\|E9QN31\|E9QN31_MOUSE Probable 28S rRNA (cytosine-C(5))-methyltransferase OS=Mus... | tr\|E9QN31\|E9QN31_MOUSE Probable 28S rRNA (cytosine-C(5))-methyltransferase OS=Mus... |
| tr\|Q3TJK3\|Q3TJK3_MOUSE Uncharacterized protein OS=Mus musculus OX=10090 GN=Serpinh1... | tr\|Q3TJK3\|Q3TJK3_MOUSE Uncharacterized protein OS=Mus musculus OX=10090 GN=Serpinh1... |

| | |
|---|---|
| tr\|Q3UJK5\|Q3UJK5_MOUSE Uncharacterized protein OS=Mus musculus OX=10090 GN=Oat PE=2... | Uncharacterized protein OS=Mus musculus OX=10090 GN=Oat PE=2 SV=1 |
| S4R1I3_MOUSE | Xaa-Pro aminopeptidase 1 OS=Mus musculus OX=10090 GN=Xpnpep1 PE=1 SV=1 |
| G3X8R5_MOUSE | Glutamine-rich protein 1 OS=Mus musculus OX=10090 GN=Qrich1 PE=1 SV=1 |
| sp\|A2AN08-5\|UBR4_MOUSE | Isoform 5 of E3 ubiquitin-protein ligase UBR4 OS=Mus musculus OX=10090 GN=Ubr4 |
| ERF1_MOUSE | Eukaryotic peptide chain release factor subunit 1 OS=Mus musculus OX=10090 GN=Etf1 PE=1 SV=4 |
| HNRPF_MOUSE | Isoform 2 of Heterogeneous nuclear ribonucleoprotein F OS=Mus musculus OX=10090 GN=Hnrnpf |
| sp\|Q9QXK3-3\|COPG2_MOUSE Isoform 3 of Coatomer subunit gamma-2 OS=Mus musculus OX=10090... | Isoform 3 of Coatomer subunit gamma-2 OS=Mus musculus OX=10090 GN=Copg2 |
| H7BX05_MOUSE | Obscurin OS=Mus musculus OX=10090 GN=Obscn PE=1 SV=2 |
| tr\|Q3UIF3\|Q3UIF3_MOUSE Uncharacterized protein OS=Mus musculus OX=10090 GN=Hsp90aa1... | tr\|Q3UIF3\|Q3UIF3_MOUSE Uncharacterized protein OS=Mus musculus OX=10090 GN=Hsp90aa1... |
| sp\|Q6P5E4\|UGGG1_MOUSEUDP-glucose:glycoprotein glucosyltransferase 1 OS=Mus musculus OX=10090 GN=Uggt1 PE=1 SV=4 | sp\|Q6P5E4\|UGGG1_MOUSE UDP-glucose:glycoprotein glucosyltransferase 1 OS=Mus musculus OX=10090 GN=Uggt1 PE=1 SV=4 |
| sp\|Q8K363\|DDX18_MOUSEATP-dependent RNA helicase DDX18 OS=Mus musculus OX=10090 GN=Ddx18 PE=1 SV=1 | sp\|Q8K363\|DDX18_MOUSE ATP-dependent RNA helicase DDX18 OS=Mus musculus OX=10090 GN=Ddx18 PE=1 SV=1 |
| sp\|Q8K2Z4\|CND1_MOUSE | Condensin complex subunit 1 OS=Mus musculus OX=10090 GN=Ncapd2 PE=1 SV=2 |
| sp\|Q00PI9\|HNRL2_MOUSEHeterogeneous nuclear ribonucleoprotein U-like protein 2 OS=Mus musculus OX=10090 GN=Hnrnpul2 PE=1 SV=2 | sp\|Q00PI9\|HNRL2_MOUSE Heterogeneous nuclear ribonucleoprotein U-like protein 2 OS=Mus musculus OX=10090 GN=Hnrnpul2 PE=1 SV=2 |
| sp\|Q64511\|TOP2B_MOUSEDNA topoisomerase 2-beta OS=Mus musculus OX=10090 GN=Top2b PE=1 SV=2 | sp\|Q64511\|TOP2B_MOUSE DNA topoisomerase 2-beta OS=Mus musculus OX=10090 GN=Top2b PE=1 SV=2 |

| | |
|---|---|
| C1TM_MOUSE | Monofunctional C1-tetrahydrofolate synthase, mitochondrial OS=Mus musculus OX=10090 GN=Mthfd1l PE=1 SV=2 |
| CUL5_MOUSE | Cullin-5 OS=Mus musculus OX=10090 GN=Cul5 PE=1 SV=3 |
| E9Q5B5_MOUSE | Hexokinase-2 OS=Mus musculus OX=10090 GN=Hk2 PE=1 SV=1 |
| G3X928_MOUSE | SEC23-interacting protein OS=Mus musculus OX=10090 GN=Sec23ip PE=1 SV=1 |
| M1VHG6_MOUSE | Metastasis-associated protein MTA1 isoform 11 OS=Mus musculus OX=10090 GN=Mta1 PE=2 SV=1 |
| Q3TQ70_MOUSE | Beta1 subnuit of GTP-binding protein OS=Mus musculus OX=10090 GN=Gnb1 PE=1 SV=1 |
| Q80W12_MOUSE | TROVE domain family, member 2 OS=Mus musculus OX=10090 GN=Trove2 PE=2 SV=1 |
| Q80ZX0_MOUSE | Sec24-related gene family, member B (S. cerevisiae) OS=Mus musculus OX=10090 GN=Sec24b PE=1 SV=1 |
| Q8VDC3_MOUSE | Cytoplasmic aconitase OS=Mus musculus OX=10090 GN=aco1 PE=3 SV=1 |
| sp\|P28481-5\|CO2A1_MOUSE | Isoform 5 of Collagen alpha-1(II) chain OS=Mus musculus OX=10090 GN=Col2a1 |
| sp\|P61205\|ARF3_MOUSE ADP-ribosylation factor 3 OS=Mus musculus OX=10090 GN=Arf3 PE=2... | ADP-ribosylation factor 3 OS=Mus musculus OX=10090 GN=Arf3 PE=2 SV=2 |
| sp\|P62141\|PP1B_MOUSE Serine/threonine-protein phosphatase PP1-beta catalytic subunit... | sp\|P62141\|PP1B_MOUSE Serine/threonine-protein phosphatase PP1-beta catalytic subunit... |
| sp\|Q3UA06\|PCH2_MOUSE Pachytene checkpoint protein 2 homolog OS=Mus musculus OX=10090... | sp\|Q3UA06\|PCH2_MOUSE Pachytene checkpoint protein 2 homolog OS=Mus musculus OX=10090... |
| sp\|Q61781\|K1C14_MOUSEKeratin, type I cytoskeletal 14 OS=Mus musculus OX=10090 GN=Krt14 PE=1 SV=2 | sp\|Q61781\|K1C14_MOUSE Keratin, type I cytoskeletal 14 OS=Mus musculus OX=10090 GN=Krt14 PE=1 SV=2 |
| sp\|Q6NS46\|RRP5_MOUSE Protein RRP5 homolog OS=Mus musculus OX=10090 GN=Pdcd11 PE=1... | sp\|Q6NS46\|RRP5_MOUSE Protein RRP5 homolog OS=Mus musculus OX=10090 GN=Pdcd11 PE=1... |

| | |
|---|---|
| sp\|Q8BRF7-3\|SCFD1_MOUSE Isoform 3 of Sec1 family domain-containing protein 1 OS=Mus... | sp\|Q8BRF7-3\|SCFD1_MOUSE Isoform 3 of Sec1 family domain-containing protein 1 OS=Mus... |
| tr\|A0A0R4J187\|A0A0R4J187_MOUSE X-ray repair cross-complementing protein 6 OS=Mus... | X-ray repair cross-complementing protein 6 OS=Mus musculus OX=10090 GN=Xrcc6 PE=1 SV=1 |
| tr\|A3KG93\|A3KG93_MOUSE Lysine-specific histone demethylase 1A OS=Mus musculus OX=10090... | tr\|A3KG93\|A3KG93_MOUSE Lysine-specific histone demethylase 1A OS=Mus musculus OX=10090... |
| tr\|E9QP46\|E9QP46_MOUSE-DECOY Nesprin-2 OS=Mus musculus OX=10090 GN=Syne2 PE=1 SV=1 | tr\|E9QP46\|E9QP46_MOUSE-DECOY Nesprin-2 OS=Mus musculus OX=10090 GN=Syne2 PE=1 SV=1 |
| tr\|Q3TC91\|Q3TC91_MOUSE Uncharacterized protein OS=Mus musculus OX=10090 GN=Cpsf3... | tr\|Q3TC91\|Q3TC91_MOUSE Uncharacterized protein OS=Mus musculus OX=10090 GN=Cpsf3... |
| tr\|Q8CD14\|Q8CD14_MOUSE Uncharacterized protein OS=Mus musculus OX=10090 GN=Ddx54... | Uncharacterized protein OS=Mus musculus OX=10090 GN=Ddx54 PE=2 SV=1 |
| sp\|P54276\|MSH6_MOUSE DNA mismatch repair protein Msh6 OS=Mus musculus OX=10090 GN=Msh6... | sp\|P54276\|MSH6_MOUSE DNA mismatch repair protein Msh6 OS=Mus musculus OX=10090 GN=Msh6... |
| tr\|Q3UB63\|Q3UB63_MOUSE Uncharacterized protein OS=Mus musculus OX=10090 GN=Slc25a3... | Uncharacterized protein OS=Mus musculus OX=10090 GN=Slc25a3 PE=2 SV=1 |
| A0A0G2JEP0_MOUSE | Fragile X mental retardation syndrome-related protein 1 OS=Mus musculus OX=10090 GN=Fxr1 PE=1 SV=1 |
| A0A0R4J0W6_MOUSE | Leucine rich repeat containing 40, isoform CRA_a OS=Mus musculus OX=10090 GN=Lrrc40 PE=1 SV=1 |
| IF2B2_MOUSE | Isoform 2 of Insulin-like growth factor 2 mRNA-binding protein 2 OS=Mus musculus OX=10090 GN=Igf2bp2 |
| tr\|Q3UR54\|Q3UR54_MOUSE Uncharacterized protein (Fragment) OS=Mus musculus OX=10090... | tr\|Q3UR54\|Q3UR54_MOUSE Uncharacterized protein (Fragment) OS=Mus musculus OX=10090... |
| A0A087WP14_MOUSE | Kinectin OS=Mus musculus OX=10090 GN=Ktn1 PE=1 SV=1 |
| A0A0R4J098_MOUSE | Zinc finger protein 326 OS=Mus musculus OX=10090 GN=Zfp326 PE=1 SV=1 |
| sp\|Q8K3H0\|DP13A_MOUSEDCC-interacting protein 13-alpha OS=Mus musculus OX=10090 GN=Appl1 PE=1 SV=1 | sp\|Q8K3H0\|DP13A_MOUSE DCC-interacting protein 13-alpha OS=Mus |

| | |
|---|---|
| | musculus OX=10090 GN=Appl1 PE=1 SV=1 |
| sp\|Q8VDS4\|RPR1A_MOUSERegulation of nuclear pre-mRNA domain-containing protein 1A OS=Mus musculus OX=10090 GN=Rprd1a PE=1 SV=1 | sp\|Q8VDS4\|RPR1A_MOUSE Regulation of nuclear pre-mRNA domain-containing protein 1A OS=Mus musculus OX=10090 GN=Rprd1a PE=1 SV=1 |
| sp\|P14115\|RL27A_MOUSE60S ribosomal protein L27a OS=Mus musculus OX=10090 GN=Rpl27a PE=1 SV=5 | sp\|P14115\|RL27A_MOUSE 60S ribosomal protein L27a OS=Mus musculus OX=10090 GN=Rpl27a PE=1 SV=5 |
| CHD5_MOUSE | Chromodomain-helicase-DNA-binding protein 5 OS=Mus musculus OX=10090 GN=Chd5 PE=1 SV=1 |
| sp\|P48036\|ANXA5_MOUSEAnnexin A5 OS=Mus musculus OX=10090 GN=Anxa5 PE=1 SV=1 | sp\|P48036\|ANXA5_MOUSE Annexin A5 OS=Mus musculus OX=10090 GN=Anxa5 PE=1 SV=1 |
| A2RRR3_MOUSE | Clasp2 protein OS=Mus musculus OX=10090 GN=Clasp2 PE=2 SV=1 |
| RCC2_MOUSE | Protein RCC2 OS=Mus musculus OX=10090 GN=Rcc2 PE=1 SV=1 |
| sp\|Q9CPN8\|IF2B3_MOUSEInsulin-like growth factor 2 mRNA-binding protein 3 OS=Mus musculus OX=10090 GN=Igf2bp3 PE=1 SV=1 | sp\|Q9CPN8\|IF2B3_MOUSE Insulin-like growth factor 2 mRNA-binding protein 3 OS=Mus musculus OX=10090 GN=Igf2bp3 PE=1 SV=1 |
| Q3U547_MOUSE | Uncharacterized protein OS=Mus musculus OX=10090 GN=Stt3a PE=2 SV=1 |
| E9Q481_MOUSE | Serine/threonine-protein phosphatase 4 regulatory subunit 3A OS=Mus musculus OX=10090 GN=Ppp4r3a PE=1 SV=1 |
| E9Q804_MOUSE | Ankyrin repeat domain-containing protein 17 OS=Mus musculus OX=10090 GN=Ankrd17 PE=1 SV=3 |
| K3W4Q5_MOUSE | Family with sequence similarity 186, member A OS=Mus musculus OX=10090 GN=Fam186a PE=4 SV=1 |
| Q8R1V9_MOUSE | Qars protein OS=Mus musculus OX=10090 GN=Qars PE=1 SV=1 |
| sp\|P47963\|RL13_MOUSE 60S ribosomal protein L13 OS=Mus musculus OX=10090 GN=Rpl13... | sp\|P47963\|RL13_MOUSE 60S ribosomal protein L13 OS=Mus musculus OX=10090 GN=Rpl13... |
| sp\|P55200-2\|KMT2A_MOUSE-DECOY Isoform 2 of Histone-lysine N-methyltransferase 2A... | sp\|P55200-2\|KMT2A_MOUSE-DECOY Isoform 2 of Histone-lysine N-methyltransferase 2A... |

| | |
|---|---|
| sp\|Q70FJ1-2\|AKAP9_MOUSE Isoform 2 of A-kinase anchor protein 9 OS=Mus musculus OX=10090... | Isoform 2 of A-kinase anchor protein 9 OS=Mus musculus OX=10090 GN=Akap9 |
| sp\|Q8BL97-3\|SRSF7_MOUSE Isoform 3 of Serine/arginine-rich splicing factor 7 OS=Mus... | sp\|Q8BL97-3\|SRSF7_MOUSE Isoform 3 of Serine/arginine-rich splicing factor 7 OS=Mus... |
| sp\|Q8K0D5\|EFGM_MOUSE Elongation factor G, mitochondrial OS=Mus musculus OX=10090... | sp\|Q8K0D5\|EFGM_MOUSE Elongation factor G, mitochondrial OS=Mus musculus OX=10090... |
| sp\|Q99JX4\|EIF3M_MOUSEEukaryotic translation initiation factor 3 subunit M OS=Mus musculus OX=10090 GN=Eif3m PE=1 SV=1 | sp\|Q99JX4\|EIF3M_MOUSE Eukaryotic translation initiation factor 3 subunit M OS=Mus musculus OX=10090 GN=Eif3m PE=1 SV=1 |
| tr\|A0A1W2P6U8\|A0A1W2P6U8_MOUSE-DECOY RIKEN cDNA 4932415D10 gene OS=Mus musculus OX=10090... | tr\|A0A1W2P6U8\|A0A1W2P6U8_MOUSE-DECOY RIKEN cDNA 4932415D10 gene OS=Mus musculus OX=10090... |
| tr\|A3KFU8\|A3KFU8_MOUSE Polyadenylate-binding protein OS=Mus musculus OX=10090 GN=Pabpc4... | Polyadenylate-binding protein OS=Mus musculus OX=10090 GN=Pabpc4 PE=1 SV=1 |
| tr\|Q3UFJ3\|Q3UFJ3_MOUSE Pyruvate dehydrogenase E1 component subunit alpha OS=Mus musculus... | tr\|Q3UFJ3\|Q3UFJ3_MOUSE Pyruvate dehydrogenase E1 component subunit alpha OS=Mus musculus... |
| A0A0R4J1E3_MOUSE | Drebrin OS=Mus musculus OX=10090 GN=Dbn1 PE=1 SV=1 |
| D3YYI8_MOUSE | Histone deacetylase OS=Mus musculus OX=10090 GN=Gm10093 PE=3 SV=1 |
| H9KV00_MOUSE | Protein SON OS=Mus musculus OX=10090 GN=Son PE=1 SV=1 |
| sp\|O88738-2\|BIRC6_MOUSE | Isoform 2 of Baculoviral IAP repeat-containing protein 6 OS=Mus musculus OX=10090 GN=Birc6 |
| sp\|Q99ME9\|NOG1_MOUSE Nucleolar GTP-binding protein 1 OS=Mus musculus OX=10090 GN=Gtpbp4... | sp\|Q99ME9\|NOG1_MOUSE Nucleolar GTP-binding protein 1 OS=Mus musculus OX=10090 GN=Gtpbp4... |
| Q4VAG4_MOUSE | MCG12304 OS=Mus musculus OX=10090 GN=Rpl22 PE=1 SV=1 |
| tr\|Q3U7B3\|Q3U7B3_MOUSE Uncharacterized protein OS=Mus musculus OX=10090 GN=Tmpo PE=2... | Uncharacterized protein OS=Mus musculus OX=10090 GN=Tmpo PE=2 SV=1 |
| A0A2I3BQS6_MOUSE | E3 ubiquitin-protein ligase UBR5 OS=Mus musculus OX=10090 GN=Ubr5 PE=1 SV=1 |

| | |
|---|---|
| sp\|Q60598\|SRC8_MOUSE Src substrate cortactin OS=Mus musculus OX=10090 GN=Cttn PE=1... | sp\|Q60598\|SRC8_MOUSE Src substrate cortactin OS=Mus musculus OX=10090 GN=Cttn PE=1... |
| Q6PFF0_MOUSE | SR-related CTD-associated factor 4 OS=Mus musculus OX=10090 GN=Scaf4 PE=1 SV=1 |
| B1AX98_MOUSE | Leucine rich repeat containing 47 OS=Mus musculus OX=10090 GN=Lrrc47 PE=1 SV=1 |
| sp\|Q8CCF0\|PRP31_MOUSEU4/U6 small nuclear ribonucleoprotein Prp31 OS=Mus musculus OX=10090 GN=Prpf31 PE=1 SV=3 | sp\|Q8CCF0\|PRP31_MOUSE U4/U6 small nuclear ribonucleoprotein Prp31 OS=Mus musculus OX=10090 GN=Prpf31 PE=1 SV=3 |
| A0A1B0GSX7_MOUSE | Nuclear pore complex protein Nup98-Nup96 OS=Mus musculus OX=10090 GN=Nup98 PE=1 SV=1 |
| Q99JZ4_MOUSE | GTP-binding protein SAR1a OS=Mus musculus OX=10090 GN=Sar1a PE=1 SV=1 |
| B2RQS6_MOUSE | Dhx36 protein OS=Mus musculus OX=10090 GN=Dhx36 PE=2 SV=1 |
| E9PZF0_MOUSE | Nucleoside diphosphate kinase OS=Mus musculus OX=10090 GN=Gm20390 PE=3 SV=1 |
| sp\|P47911\|RL6_MOUSE 60S ribosomal protein L6 OS=Mus musculus OX=10090 GN=Rpl6 PE=1... | sp\|P47911\|RL6_MOUSE 60S ribosomal protein L6 OS=Mus musculus OX=10090 GN=Rpl6 PE=1... |
| sp\|Q8BGH2\|SAM50_MOUSESorting and assembly machinery component 50 homolog OS=Mus musculus OX=10090 GN=Samm50 PE=1 SV=1 | sp\|Q8BGH2\|SAM50_MOUSE Sorting and assembly machinery component 50 homolog OS=Mus musculus OX=10090 GN=Samm50 PE=1 SV=1 |
| Q3UJI2_MOUSE | Uncharacterized protein OS=Mus musculus OX=10090 GN=Rbbp7 PE=2 SV=1 |
| sp\|Q8BWW9\|PKN2_MOUSE | Serine/threonine-protein kinase N2 OS=Mus musculus OX=10090 GN=Pkn2 PE=1 SV=3 |
| A0A1L1STE6_MOUSE | Isocitrate dehydrogenase [NAD] subunit, mitochondrial OS=Mus musculus OX=10090 GN=Idh3a PE=1 SV=1 |
| tr\|Q059T9\|Q059T9_MOUSE PRP4 pre-mRNA processing factor 4 homolog (Yeast) OS=Mus musculus... | tr\|Q059T9\|Q059T9_MOUSE PRP4 pre-mRNA processing factor 4 homolog (Yeast) OS=Mus musculus... |
| ATRX_MOUSE | Transcriptional regulator ATRX OS=Mus musculus OX=10090 GN=Atrx PE=1 SV=3 |

| | |
|---|---|
| ERF3A_MOUSE | Isoform 2 of Eukaryotic peptide chain release factor GTP-binding subunit ERF3A OS=Mus musculus OX=10090 GN=Gspt1 |
| A0A0R4J036_MOUSE | Neurofilament 3, medium OS=Mus musculus OX=10090 GN=Nefm PE=1 SV=1 |
| A0A0G2JDV8_MOUSE | Calponin OS=Mus musculus OX=10090 GN=Cnn3 PE=1 SV=1 |
| A1L366_MOUSE | RAD21 homolog (S. pombe) OS=Mus musculus OX=10090 GN=Rad21 PE=2 SV=1 |
| A2RRJ4_MOUSE | Exportin 5 OS=Mus musculus OX=10090 GN=Xpo5 PE=2 SV=1 |
| A8DUL0_MOUSE | Beta-globin OS=Mus musculus OX=10090 GN=Hbbt1 PE=3 SV=1 |
| BAG6_MOUSE | Large proline-rich protein BAG6 OS=Mus musculus OX=10090 GN=Bag6 PE=1 SV=1 |
| Q3U548_MOUSE | UTP--glucose-1-phosphate uridylyltransferase OS=Mus musculus OX=10090 GN=Ugp2 PE=2 SV=1 |
| Q3UDK3_MOUSE | Uncharacterized protein OS=Mus musculus OX=10090 GN=Rfc5 PE=2 SV=1 |
| Q3UIZ1_MOUSE | UDP-glucose 6-dehydrogenase OS=Mus musculus OX=10090 GN=Ugdh PE=2 SV=1 |
| Q5M9J8_MOUSE | MCG13936 OS=Mus musculus OX=10090 GN=Rpl28 PE=1 SV=1 |
| sp\|Q6ZWR6-4\|SYNE1_MOUSE-DECOY Isoform 4 of Nesprin-1 OS=Mus musculus OX=10090 GN=Syne1 | sp\|Q6ZWR6-4\|SYNE1_MOUSE-DECOY Isoform 4 of Nesprin-1 OS=Mus musculus OX=10090 GN=Syne1 |
| sp\|Q8BTI8-2\|SRRM2_MOUSE-DECOY Isoform 2 of Serine/arginine repetitive matrix protein... | sp\|Q8BTI8-2\|SRRM2_MOUSE-DECOY Isoform 2 of Serine/arginine repetitive matrix protein... |
| sp\|Q9D051\|ODPB_MOUSE Pyruvate dehydrogenase E1 component subunit beta, mitochondrial... | sp\|Q9D051\|ODPB_MOUSE Pyruvate dehydrogenase E1 component subunit beta, mitochondrial... |
| sp\|Q9EP69\|SAC1_MOUSE Phosphatidylinositide phosphatase SAC1 OS=Mus musculus OX=10090... | sp\|Q9EP69\|SAC1_MOUSE Phosphatidylinositide phosphatase SAC1 OS=Mus musculus OX=10090... |
| sp\|Q9JJ28\|FLII_MOUSE Protein flightless-1 homolog OS=Mus musculus OX=10090 GN=Flii... | Protein flightless-1 homolog OS=Mus musculus OX=10090 GN=Flii PE=1 SV=1 |

| | |
| --- | --- |
| sp\|Q9Z0E0\|NCDN_MOUSE | Neurochondrin OS=Mus musculus OX=10090 GN=Ncdn PE=1 SV=1 |
| tr\|D3YWF6\|D3YWF6_MOUSE Ubiquitin thioesterase OTUB1 OS=Mus musculus OX=10090 GN=Otub1... | tr\|D3YWF6\|D3YWF6_MOUSE Ubiquitin thioesterase OTUB1 OS=Mus musculus OX=10090 GN=Otub1... |
| tr\|E9PUC2\|E9PUC2_MOUSE Long-chain-fatty-acid--CoA ligase 3 OS=Mus musculus OX=10090... | tr\|E9PUC2\|E9PUC2_MOUSE Long-chain-fatty-acid--CoA ligase 3 OS=Mus musculus OX=10090... |
| tr\|E9Q5I9\|E9Q5I9_MOUSE 26S proteasome non-ATPase regulatory subunit 13 OS=Mus musculus... | tr\|E9Q5I9\|E9Q5I9_MOUSE 26S proteasome non-ATPase regulatory subunit 13 OS=Mus musculus... |
| tr\|E9Q616\|E9Q616_MOUSE-DECOY AHNAK nucleoprotein (desmoyokin) OS=Mus musculus OX=10090... | tr\|E9Q616\|E9Q616_MOUSE-DECOY AHNAK nucleoprotein (desmoyokin) OS=Mus musculus OX=10090... |
| tr\|Q497N1\|Q497N1_MOUSE 40S ribosomal protein S26 OS=Mus musculus OX=10090 GN=Rps26... | 40S ribosomal protein S26 OS=Mus musculus OX=10090 GN=Rps26 PE=1 SV=1 |
| tr\|Q4FJL0\|Q4FJL0_MOUSE RAB10, member RAS oncogene family OS=Mus musculus OX=10090... | tr\|Q4FJL0\|Q4FJL0_MOUSE RAB10, member RAS oncogene family OS=Mus musculus OX=10090... |
| tr\|Q80U83\|Q80U83_MOUSE MKIAA0079 protein (Fragment) OS=Mus musculus OX=10090 GN=Sec24c... | tr\|Q80U83\|Q80U83_MOUSE MKIAA0079 protein (Fragment) OS=Mus musculus OX=10090 GN=Sec24c... |
| tr\|Q3TL52\|Q3TL52_MOUSE Uncharacterized protein OS=Mus musculus OX=10090 GN=Ncl PE=2... | tr\|Q3TL52\|Q3TL52_MOUSE Uncharacterized protein OS=Mus musculus OX=10090 GN=Ncl PE=2... |
| Q545V8_MOUSE | Uncharacterized protein OS=Mus musculus OX=10090 GN=Csnk2a2 PE=1 SV=1 |
| sp\|P27773\|PDIA3_MOUSEProtein disulfide-isomerase A3 OS=Mus musculus OX=10090 GN=Pdia3 PE=1 SV=2 | sp\|P27773\|PDIA3_MOUSE Protein disulfide-isomerase A3 OS=Mus musculus OX=10090 GN=Pdia3 PE=1 SV=2 |
| tr\|E9Q1Y3\|E9Q1Y3_MOUSE-DECOY Apolipoprotein B-100 (Fragment) OS=Mus musculus OX=10090... | tr\|E9Q1Y3\|E9Q1Y3_MOUSE-DECOY Apolipoprotein B-100 (Fragment) OS=Mus musculus OX=10090... |
| Q3TSX8_MOUSE | Uncharacterized protein OS=Mus musculus OX=10090 GN=Tomm70a PE=2 SV=1 |
| Q3TVT4_MOUSE | Uncharacterized protein (Fragment) OS=Mus musculus OX=10090 GN=Dync1li1 PE=2 SV=1 |
| A0A0U1RPW2_MOUSE | Tight junction protein ZO-1 OS=Mus musculus OX=10090 GN=Tjp1 PE=1 SV=1 |

| | |
|---|---|
| sp\|P27546\|MAP4_MOUSE | Microtubule-associated protein 4 OS=Mus musculus OX=10090 GN=Map4 PE=1 SV=3 |
| sp\|Q9JLI8-2\|SART3_MOUSE Isoform 2 of Squamous cell carcinoma antigen recognized by... | sp\|Q9JLI8-2\|SART3_MOUSE Isoform 2 of Squamous cell carcinoma antigen recognized by... |
| A0A0R4J1G5_MOUSE | Erlin-1 OS=Mus musculus OX=10090 GN=Erlin1 PE=1 SV=1 |
| B1AT03_MOUSE | DNA ligase OS=Mus musculus OX=10090 GN=Lig3 PE=1 SV=1 |
| tr\|Q9D0K3\|Q9D0K3_MOUSE Uncharacterized protein OS=Mus musculus OX=10090 GN=Rfc2 PE=2... | Uncharacterized protein OS=Mus musculus OX=10090 GN=Rfc2 PE=2 SV=1 |
| APOB_MOUSE | Apolipoprotein B-100 OS=Mus musculus OX=10090 GN=Apob PE=1 SV=1 |
| sp\|Q9DBG7\|SRPRA_MOUSESignal recognition particle receptor subunit alpha OS=Mus musculus OX=10090 GN=Srpra PE=1 SV=1 | sp\|Q9DBG7\|SRPRA_MOUSE Signal recognition particle receptor subunit alpha OS=Mus musculus OX=10090 GN=Srpra PE=1 SV=1 |
| A2A4N9_MOUSE | ATP-dependent RNA helicase DHX8 (Fragment) OS=Mus musculus OX=10090 GN=Dhx8 PE=1 SV=8 |
| E9Q1G8_MOUSE | Septin-7 OS=Mus musculus OX=10090 GN=Sept7 PE=1 SV=2 |
| sp\|Q6A4J8-2\|UBP7_MOUSE | Isoform 2 of Ubiquitin carboxyl-terminal hydrolase 7 OS=Mus musculus OX=10090 GN=Usp7 |
| sp\|Q6NV83-2\|SR140_MOUSE Isoform 2 of U2 snRNP-associated SURP motif-containing protein... | sp\|Q6NV83-2\|SR140_MOUSE Isoform 2 of U2 snRNP-associated SURP motif-containing protein... |
| sp\|E9PVX6-2\|KI67_MOUSE-DECOY Isoform 2 of Proliferation marker protein Ki-67 OS=Mus... | sp\|E9PVX6-2\|KI67_MOUSE-DECOY Isoform 2 of Proliferation marker protein Ki-67 OS=Mus... |
| Q8BTU6_MOUSE | Eukaryotic initiation factor 4A-II OS=Mus musculus OX=10090 GN=Eif4a2 PE=1 SV=1 |
| G3UXW9_MOUSE | COP9 signalosome complex subunit 1 OS=Mus musculus OX=10090 GN=Gps1 PE=1 SV=1 |
| sp\|Q6P9L6\|KIF15_MOUSEKinesin-like protein KIF15 OS=Mus musculus OX=10090 GN=Kif15 PE=1 SV=1 | sp\|Q6P9L6\|KIF15_MOUSE Kinesin-like protein KIF15 OS=Mus musculus OX=10090 GN=Kif15 PE=1 SV=1 |

| | |
|---|---|
| PRAX_MOUSE | Periaxin OS=Mus musculus OX=10090 GN=Prx PE=1 SV=1 |
| sp\|Q6PB66\|LPPRC_MOUSELeucine-rich PPR motif-containing protein, mitochondrial OS=Mus musculus OX=10090 GN=Lrpprc PE=1 SV=2 | sp\|Q6PB66\|LPPRC_MOUSE Leucine-rich PPR motif-containing protein, mitochondrial OS=Mus musculus OX=10090 GN=Lrpprc PE=1 SV=2 |
| Q5SUH6_MOUSE | Clathrin interactor 1 OS=Mus musculus OX=10090 GN=Clint1 PE=1 SV=1 |
| Q8BPH1_MOUSE | Uncharacterized protein OS=Mus musculus OX=10090 GN=Ywhae PE=2 SV=1 |
| Q8BTQ1_MOUSE | ELAV-like protein (Fragment) OS=Mus musculus OX=10090 GN=Elavl1 PE=2 SV=1 |
| Q0VDR7_MOUSE | Krt6b protein OS=Mus musculus OX=10090 GN=Krt6b PE=2 SV=1 |
| A0A0R4J195_MOUSE | Regulation of nuclear pre-mRNA domain-containing protein 1B OS=Mus musculus OX=10090 GN=Rprd1b PE=1 SV=1 |
| A0A2C9F2D2_MOUSE | Annexin OS=Mus musculus OX=10090 GN=Anxa7 PE=1 SV=1 |
| A2BFF8_MOUSE | Cytoplasmic dynein 1 intermediate chain 2 OS=Mus musculus OX=10090 GN=Dync1i2 PE=1 SV=1 |
| B1AUY7_MOUSE | N-alpha-acetyltransferase 10 OS=Mus musculus OX=10090 GN=Naa10 PE=1 SV=1 |
| CDK2_MOUSE | Cyclin-dependent kinase 2 OS=Mus musculus OX=10090 GN=Cdk2 PE=1 SV=2 |
| DYH5_MOUSE | Dynein heavy chain 5, axonemal OS=Mus musculus OX=10090 GN=Dnah5 PE=1 SV=2 |
| Q8BSZ8_MOUSE | Uncharacterized protein OS=Mus musculus OX=10090 GN=Srm PE=2 SV=1 |
| S4R2F3_MOUSE | Ankyrin-2 OS=Mus musculus OX=10090 GN=Ank2 PE=1 SV=3 |
| sp\|A2AAE1-6\|K1109_MOUSE-DECOY Isoform 6 of Uncharacterized protein KIAA1109 OS=Mus... | sp\|A2AAE1-6\|K1109_MOUSE-DECOY Isoform 6 of Uncharacterized protein KIAA1109 OS=Mus... |
| sp\|P26443\|DHE3_MOUSE Glutamate dehydrogenase 1, mitochondrial OS=Mus musculus OX=10090... | sp\|P26443\|DHE3_MOUSE Glutamate dehydrogenase 1, mitochondrial OS=Mus musculus OX=10090... |

| | |
|---|---|
| sp\|P55200-2\|KMT2A_MOUSE Isoform 2 of Histone-lysine N-methyltransferase 2A OS=Mus... | sp\|P55200-2\|KMT2A_MOUSE Isoform 2 of Histone-lysine N-methyltransferase 2A OS=Mus... |
| sp\|Q6R0H7-4\|GNAS1_MOUSE Isoform XLas-4 of Guanine nucleotide-binding protein G(s)... | sp\|Q6R0H7-4\|GNAS1_MOUSE Isoform XLas-4 of Guanine nucleotide-binding protein G(s)... |
| sp\|Q80U72-2\|SCRIB_MOUSE Isoform 2 of Protein scribble homolog OS=Mus musculus OX=10090... | sp\|Q80U72-2\|SCRIB_MOUSE Isoform 2 of Protein scribble homolog OS=Mus musculus OX=10090... |
| sp\|Q922J9-3\|FACR1_MOUSE | Isoform 3 of Fatty acyl-CoA reductase 1 OS=Mus musculus OX=10090 GN=Far1 |
| sp\|Q9JHS4\|CLPX_MOUSE ATP-dependent Clp protease ATP-binding subunit clpX-like, mitochondrial... | sp\|Q9JHS4\|CLPX_MOUSE ATP-dependent Clp protease ATP-binding subunit clpX-like, mitochondrial... |
| sp\|Q9QZQ8-2\|H2AY_MOUSE Isoform 1 of Core histone macro-H2A.1 OS=Mus musculus OX=10090... | sp\|Q9QZQ8-2\|H2AY_MOUSE Isoform 1 of Core histone macro-H2A.1 OS=Mus musculus OX=10090... |
| tr\|H7BX05\|H7BX05_MOUSE-DECOY Obscurin OS=Mus musculus OX=10090 GN=Obscn PE=1 SV=2 | tr\|H7BX05\|H7BX05_MOUSE-DECOY Obscurin OS=Mus musculus OX=10090 GN=Obscn PE=1 SV=2 |
| tr\|Q3TQ33\|Q3TQ33_MOUSE Uncharacterized protein (Fragment) OS=Mus musculus OX=10090... | tr\|Q3TQ33\|Q3TQ33_MOUSE Uncharacterized protein (Fragment) OS=Mus musculus OX=10090... |
| Q3UE11_MOUSE | Uncharacterized protein OS=Mus musculus OX=10090 PE=2 SV=1 |
| sp\|Q8BTW9\|PAK4_MOUSE Serine/threonine-protein kinase PAK 4 OS=Mus musculus OX=10090... | Serine/threonine-protein kinase PAK 4 OS=Mus musculus OX=10090 GN=Pak4 PE=1 SV=1 |
| tr\|A2AD84\|A2AD84_MOUSE Serine/threonine-protein kinase 26 OS=Mus musculus OX=10090... | Serine/threonine-protein kinase 26 OS=Mus musculus OX=10090 GN=Stk26 PE=1 SV=1 |
| tr\|F8VQ05\|F8VQ05_MOUSE-DECOY FRY-like transcription coactivator OS=Mus musculus OX=10090... | tr\|F8VQ05\|F8VQ05_MOUSE-DECOY FRY-like transcription coactivator OS=Mus musculus OX=10090... |
| tr\|Q99K20\|Q99K20_MOUSE L-lactate dehydrogenase (Fragment) OS=Mus musculus OX=10090... | L-lactate dehydrogenase (Fragment) OS=Mus musculus OX=10090 GN=Ldha PE=2 SV=1 |
| A0A286YDJ6_MOUSE | Lysine-specific demethylase 3B OS=Mus musculus OX=10090 GN=Kdm3b PE=1 SV=1 |
| sp\|A2AWL7\|MGAP_MOUSE | MAX gene-associated protein OS=Mus musculus OX=10090 GN=Mga PE=2 SV=1 |

| | |
|---|---|
| E9PWV3_MOUSE | Ribosomal protein S6 kinase OS=Mus musculus OX=10090 GN=Rps6ka1 PE=1 SV=1 |
| sp\|P97412\|LYST_MOUSE | Lysosomal-trafficking regulator OS=Mus musculus OX=10090 GN=Lyst PE=1 SV=1 |
| Q6PFB2_MOUSE | Rcc1 protein OS=Mus musculus OX=10090 GN=Rcc1 PE=1 SV=1 |
| Q4FJZ4_MOUSE | Wars protein OS=Mus musculus OX=10090 GN=Wars PE=2 SV=1 |
| tr\|D3Z7E5\|D3Z7E5_MOUSE Glycogen synthase kinase-3 alpha OS=Mus musculus OX=10090... | tr\|D3Z7E5\|D3Z7E5_MOUSE Glycogen synthase kinase-3 alpha OS=Mus musculus OX=10090... |
| DLDH_MOUSE | Dihydrolipoyl dehydrogenase, mitochondrial OS=Mus musculus OX=10090 GN=Dld PE=1 SV=2 |
| Q5XJZ3_MOUSE | Grwd1 protein (Fragment) OS=Mus musculus OX=10090 GN=Grwd1 PE=2 SV=1 |
| B2RXQ8_MOUSE | MCG19514, isoform CRA_a OS=Mus musculus OX=10090 GN=Wdr82 PE=1 SV=1 |
| tr\|E9QPD7\|E9QPD7_MOUSE Pyruvate carboxylase OS=Mus musculus OX=10090 GN=Pcx PE=1... | tr\|E9QPD7\|E9QPD7_MOUSE Pyruvate carboxylase OS=Mus musculus OX=10090 GN=Pcx PE=1... |
| VP26A_MOUSE | Isoform 2 of Vacuolar protein sorting-associated protein 26A OS=Mus musculus OX=10090 GN=Vps26a |
| E9Q9M0_MOUSE | Centrosomal protein of 290 kDa OS=Mus musculus OX=10090 GN=Cep290 PE=1 SV=1 |
| tr\|O08821\|O08821_MOUSE Cytoplasmic dynein heavy chain (Fragment) OS=Mus musculus... | tr\|O08821\|O08821_MOUSE Cytoplasmic dynein heavy chain (Fragment) OS=Mus musculus... |
| tr\|Q3U6K8\|Q3U6K8_MOUSE Uncharacterized protein OS=Mus musculus OX=10090 GN=Vdac1... | tr\|Q3U6K8\|Q3U6K8_MOUSE Uncharacterized protein OS=Mus musculus OX=10090 GN=Vdac1... |
| A0A0U1RPL0_MOUSE | Ataxin-2-like protein OS=Mus musculus OX=10090 GN=Atxn2l PE=1 SV=1 |
| sp\|Q61553\|FSCN1_MOUSEFascin OS=Mus musculus OX=10090 GN=Fscn1 PE=1 SV=4 | sp\|Q61553\|FSCN1_MOUSE Fascin OS=Mus musculus OX=10090 GN=Fscn1 PE=1 SV=4 |

| | |
|---|---|
| tr\|Q9DCR1\|Q9DCR1_MOUSE Uncharacterized protein (Fragment) OS=Mus musculus OX=10090... | Uncharacterized protein (Fragment) OS=Mus musculus OX=10090 GN=Tubb4b PE=2 SV=2 |
| sp\|P05132-2\|KAPCA_MOUSE Isoform 2 of cAMP-dependent protein kinase catalytic subunit... | sp\|P05132-2\|KAPCA_MOUSE Isoform 2 of cAMP-dependent protein kinase catalytic subunit... |
| FAT3_MOUSE | Protocadherin Fat 3 OS=Mus musculus OX=10090 GN=Fat3 PE=1 SV=2 |
| SYAM_MOUSE | Alanine--tRNA ligase, mitochondrial OS=Mus musculus OX=10090 GN=Aars2 PE=1 SV=1 |
| G3X9B1_MOUSE | HEAT repeat-containing 1 OS=Mus musculus OX=10090 GN=Heatr1 PE=1 SV=1 |
| PLAK_MOUSE | Junction plakoglobin OS=Mus musculus OX=10090 GN=Jup PE=1 SV=3 |
| Q3TCI8_MOUSE | Glucosamine OS=Mus musculus OX=10090 GN=Gne PE=1 SV=1 |
| tr\|Q3UL32\|Q3UL32_MOUSE Uncharacterized protein OS=Mus musculus OX=10090 GN=Dnajc7... | Uncharacterized protein OS=Mus musculus OX=10090 GN=Dnajc7 PE=2 SV=1 |
| sp\|Q9D024-2\|CCD47_MOUSE | Isoform 2 of Coiled-coil domain-containing protein 47 OS=Mus musculus OX=10090 GN=Ccdc47 |
| TBG1_MOUSE | Tubulin gamma-1 chain OS=Mus musculus OX=10090 GN=Tubg1 PE=1 SV=1 |
| sp\|Q921N6\|DDX27_MOUSEProbable ATP-dependent RNA helicase DDX27 OS=Mus musculus OX=10090 GN=Ddx27 PE=1 SV=3 | sp\|Q921N6\|DDX27_MOUSE Probable ATP-dependent RNA helicase DDX27 OS=Mus musculus OX=10090 GN=Ddx27 PE=1 SV=3 |
| HERC2_MOUSE | Isoform 2 of E3 ubiquitin-protein ligase HERC2 OS=Mus musculus OX=10090 GN=Herc2 |
| D3YUP1_MOUSE | Histone-arginine methyltransferase CARM1 OS=Mus musculus OX=10090 GN=Carm1 PE=1 SV=1 |
| E9Q6R3_MOUSE | Vesicle-trafficking protein SEC22b OS=Mus musculus OX=10090 GN=Sec22b PE=1 SV=1 |
| Q3ZAW8_MOUSE | Keratin 16 OS=Mus musculus OX=10090 GN=Krt16 PE=2 SV=1 |

| | |
|---|---|
| Q8BZZ0_MOUSE | Gamma-tubulin complex component OS=Mus musculus OX=10090 GN=Tubgcp2 PE=2 SV=1 |
| sp\|E9Q8T7\|DYH1_MOUSE Dynein heavy chain 1, axonemal OS=Mus musculus OX=10090 GN=Dnah1... | Dynein heavy chain 1, axonemal OS=Mus musculus OX=10090 GN=Dnah1 PE=1 SV=1 |
| sp\|Q62009-4\|POSTN_MOUSE | Isoform 4 of Periostin OS=Mus musculus OX=10090 GN=Postn |
| tr\|A2AUY4\|A2AUY4_MOUSE-DECOY Bromodomain adjacent to zinc finger domain, 2B OS=Mus... | tr\|A2AUY4\|A2AUY4_MOUSE-DECOY Bromodomain adjacent to zinc finger domain, 2B OS=Mus... |
| tr\|B1ARU1\|B1ARU1_MOUSE-DECOY Microtubule-actin cross-linking factor 1 OS=Mus musculus... | tr\|B1ARU1\|B1ARU1_MOUSE-DECOY Microtubule-actin cross-linking factor 1 OS=Mus musculus... |
| tr\|B9EKJ3\|B9EKJ3_MOUSE-DECOY Mon2 protein OS=Mus musculus OX=10090 GN=Mon2 PE=1 SV=1 | tr\|B9EKJ3\|B9EKJ3_MOUSE-DECOY Mon2 protein OS=Mus musculus OX=10090 GN=Mon2 PE=1 SV=1 |
| tr\|E9Q740\|E9Q740_MOUSE Signal recognition particle subunit SRP72 OS=Mus musculus... | Signal recognition particle subunit SRP72 OS=Mus musculus OX=10090 GN=Srp72 PE=1 SV=1 |
| tr\|Q99J49\|Q99J49_MOUSE Tubb2a protein (Fragment) OS=Mus musculus OX=10090 GN=Tubb2a... | Tubb2a protein (Fragment) OS=Mus musculus OX=10090 GN=Tubb2a PE=2 SV=1 |
| tr\|W6PPR4\|W6PPR4_MOUSE-DECOY 480-kDa ankyrinG OS=Mus musculus OX=10090 GN=Ank3 PE=4... | tr\|W6PPR4\|W6PPR4_MOUSE-DECOY 480-kDa ankyrinG OS=Mus musculus OX=10090 GN=Ank3 PE=4... |
| A0A0R4J0S4_MOUSE | Lethal(2) giant larvae protein homolog 1 OS=Mus musculus OX=10090 GN=Llgl1 PE=1 SV=1 |
| sp\|P49615\|CDK5_MOUSE Cyclin-dependent-like kinase 5 OS=Mus musculus OX=10090 GN=Cdk5... | Cyclin-dependent-like kinase 5 OS=Mus musculus OX=10090 GN=Cdk5 PE=1 SV=1 |
| tr\|A0A087WNX7\|A0A087WNX7_MOUSE Snf2-related CREBBP activator protein OS=Mus musculus... | tr\|A0A087WNX7\|A0A087WNX7_MOUSE Snf2-related CREBBP activator protein OS=Mus musculus... |
| tr\|A0A0A6YWA9\|A0A0A6YWA9_MOUSE Guanine nucleotide-binding protein G(i) subunit alpha-2... | tr\|A0A0A6YWA9\|A0A0A6YWA9_MOUSE Guanine nucleotide-binding protein G(i) subunit alpha-2... |
| tr\|Q3TUQ7\|Q3TUQ7_MOUSE Non-specific serine/threonine protein kinase OS=Mus musculus... | Non-specific serine/threonine protein kinase OS=Mus musculus OX=10090 GN=Prkaa1 PE=1 SV=1 |

| | |
|---|---|
| E9Q1T3_MOUSE | Mitogen-activated protein kinase kinase kinase kinase OS=Mus musculus OX=10090 GN=Map4k5 PE=1 SV=1 |
| RL11_MOUSE | 60S ribosomal protein L11 OS=Mus musculus OX=10090 GN=Rpl11 PE=1 SV=4 |
| tr\|A2ANY6\|A2ANY6_MOUSE-DECOY Midasin OS=Mus musculus OX=10090 GN=Mdn1 PE=1 SV=1 | tr\|A2ANY6\|A2ANY6_MOUSE-DECOY Midasin OS=Mus musculus OX=10090 GN=Mdn1 PE=1 SV=1 |
| B2RXY7_MOUSE | Carbonyl reductase 1 OS=Mus musculus OX=10090 GN=Cbr1 PE=1 SV=1 |
| sp\|D3Z7P3-2\|GLSK_MOUSE | Isoform 2 of Glutaminase kidney isoform, mitochondrial OS=Mus musculus OX=10090 GN=Gls |
| tr\|A0A0G2JDI9\|A0A0G2JDI9_MOUSE ATP-binding cassette sub-family D member 3 OS=Mus... | tr\|A0A0G2JDI9\|A0A0G2JDI9_MOUSE ATP-binding cassette sub-family D member 3 OS=Mus... |
| sp\|Q9JLB0-2\|MPP6_MOUSE Isoform Alpha of MAGUK p55 subfamily member 6 OS=Mus musculus... | Isoform Alpha of MAGUK p55 subfamily member 6 OS=Mus musculus OX=10090 GN=Mpp6 |
| sp\|Q8VHE6\|DYH5_MOUSE-DECOY Dynein heavy chain 5, axonemal OS=Mus musculus OX=10090... | sp\|Q8VHE6\|DYH5_MOUSE-DECOY Dynein heavy chain 5, axonemal OS=Mus musculus OX=10090... |
| tr\|A0A087WNZ7\|A0A087WNZ7_MOUSE E3 ubiquitin-protein ligase TRIP12 OS=Mus musculus... | E3 ubiquitin-protein ligase TRIP12 OS=Mus musculus OX=10090 GN=Trip12 PE=1 SV=1 |
| A0A0N4SV80_MOUSE | Zinc finger protein 638 OS=Mus musculus OX=10090 GN=Zfp638 PE=1 SV=1 |
| AINX_MOUSE | Alpha-internexin OS=Mus musculus OX=10090 GN=Ina PE=1 SV=3 |
| E9QNH6_MOUSE | Unconventional myosin-Ib OS=Mus musculus OX=10090 GN=Myo1b PE=1 SV=1 |
| sp\|Q9JHR7\|IDE_MOUSE Insulin-degrading enzyme OS=Mus musculus OX=10090 GN=Ide PE=1... | Insulin-degrading enzyme OS=Mus musculus OX=10090 GN=Ide PE=1 SV=1 |
| HAT1_MOUSE | Histone acetyltransferase type B catalytic subunit OS=Mus musculus OX=10090 GN=Hat1 PE=1 SV=1 |
| sp\|Q62407\|SPEG_MOUSE-DECOY Striated muscle-specific serine/threonine-protein kinase... | sp\|Q62407\|SPEG_MOUSE-DECOY Striated muscle-specific serine/threonine-protein kinase... |

| | |
|---|---|
| Q3TW11_MOUSE | Signal transducer and activator of transcription OS=Mus musculus OX=10090 GN=Stat1 PE=2 SV=1 |
| sp\|Q05512-2\|MARK2_MOUSE Isoform 2 of Serine/threonine-protein kinase MARK2 OS=Mus... | sp\|Q05512-2\|MARK2_MOUSE Isoform 2 of Serine/threonine-protein kinase MARK2 OS=Mus... |
| B2RY05_MOUSE | Pik3r4 protein OS=Mus musculus OX=10090 GN=Pik3r4 PE=2 SV=1 |
| PUM3_MOUSE | Pumilio homolog 3 OS=Mus musculus OX=10090 GN=Pum3 PE=1 SV=2 |
| Q3UXN3_MOUSE | Uncharacterized protein OS=Mus musculus OX=10090 GN=Cars PE=2 SV=1 |
| tr\|J3QMX2\|J3QMX2_MOUSE Transportin-1 (Fragment) OS=Mus musculus OX=10090 GN=Tnpo1... | Transportin-1 (Fragment) OS=Mus musculus OX=10090 GN=Tnpo1 PE=1 SV=1 |
| F7CDT0_MOUSE | NEDD8-conjugating enzyme Ubc12 (Fragment) OS=Mus musculus OX=10090 GN=Ube2m PE=1 SV=1 |
| A2A9P6_MOUSE | Cyclin-dependent kinase 11B OS=Mus musculus OX=10090 GN=Cdk11b PE=1 SV=1 |
| tr\|Q3UHW2\|Q3UHW2_MOUSE Uncharacterized protein OS=Mus musculus OX=10090 GN=Hsd17b4... | Uncharacterized protein OS=Mus musculus OX=10090 GN=Hsd17b4 PE=2 SV=1 |
| A0A140T8W1_MOUSE | Collagen alpha-5(VI) chain OS=Mus musculus OX=10090 GN=Col6a5 PE=1 SV=1 |
| sp\|Q7TMY8-3\|HUWE1_MOUSE-DECOY Isoform 3 of E3 ubiquitin-protein ligase HUWE1 OS=Mus... | sp\|Q7TMY8-3\|HUWE1_MOUSE-DECOY Isoform 3 of E3 ubiquitin-protein ligase HUWE1 OS=Mus... |
| sp\|Q7TPH6-2\|MYCB2_MOUSE Isoform 2 of E3 ubiquitin-protein ligase MYCBP2 OS=Mus musculus... | Isoform 2 of E3 ubiquitin-protein ligase MYCBP2 OS=Mus musculus OX=10090 GN=Mycbp2 |
| sp\|Q5SUR0\|PUR4_MOUSE Phosphoribosylformylglycinamidine synthase OS=Mus musculus OX=10090... | Phosphoribosylformylglycinamidine synthase OS=Mus musculus OX=10090 GN=Pfas PE=1 SV=1 |
| Q561N4_MOUSE | MCG1032217 OS=Mus musculus OX=10090 GN=Ube2l3 PE=1 SV=1 |
| sp\|Q4U2R1-2\|HERC2_MOUSE-DECOY Isoform 2 of E3 ubiquitin-protein ligase HERC2 OS=Mus... | sp\|Q4U2R1-2\|HERC2_MOUSE-DECOY Isoform 2 of E3 ubiquitin-protein ligase HERC2 OS=Mus... |
| BYST_MOUSE | Bystin OS=Mus musculus OX=10090 GN=Bysl PE=1 SV=3 |

| | |
|---|---|
| sp\|Q61245-2\|COBA1_MOUSE-DECOY Isoform Short of Collagen alpha-1(XI) chain OS=Mus... | sp\|Q61245-2\|COBA1_MOUSE-DECOY Isoform Short of Collagen alpha-1(XI) chain OS=Mus... |
| sp\|Q80U78-2\|PUM1_MOUSE | Isoform 2 of Pumilio homolog 1 OS=Mus musculus OX=10090 GN=Pum1 |
| Q8C952_MOUSE | Flap endonuclease 1 OS=Mus musculus OX=10090 GN=Fen1 PE=2 SV=1 |
| FXR2_MOUSE | Fragile X mental retardation syndrome-related protein 2 OS=Mus musculus OX=10090 GN=Fxr2 PE=1 SV=1 |
| Q3TA68_MOUSE | WD repeat domain 36 OS=Mus musculus OX=10090 GN=Wdr36 PE=1 SV=1 |
| A1BN54_MOUSE | Alpha actinin 1a OS=Mus musculus OX=10090 GN=Actn1 PE=1 SV=1 |
| Q91XW2_MOUSE | Putative ubiquitin-specific protease OS=Mus musculus OX=10090 GN=Usp9y PE=2 SV=1 |
| PESC_MOUSE | Pescadillo homolog OS=Mus musculus OX=10090 GN=Pes1 PE=1 SV=1 |
| sp\|P70303-2\|PYRG2_MOUSE | Isoform 2 of CTP synthase 2 OS=Mus musculus OX=10090 GN=Ctps2 |
| sp\|Q09XV5\|CHD8_MOUSE Chromodomain-helicase-DNA-binding protein 8 OS=Mus musculus... | sp\|Q09XV5\|CHD8_MOUSE Chromodomain-helicase-DNA-binding protein 8 OS=Mus musculus... |
| sp\|O35218\|CPSF2_MOUSECleavage and polyadenylation specificity factor subunit 2 OS=Mus musculus OX=10090 GN=Cpsf2 PE=1 SV=1 | sp\|O35218\|CPSF2_MOUSE Cleavage and polyadenylation specificity factor subunit 2 OS=Mus musculus OX=10090 GN=Cpsf2 PE=1 SV=1 |
| tr\|Q3TL79\|Q3TL79_MOUSE Uncharacterized protein OS=Mus musculus OX=10090 GN=Ahsa1... | tr\|Q3TL79\|Q3TL79_MOUSE Uncharacterized protein OS=Mus musculus OX=10090 GN=Ahsa1... |
| SNX27_MOUSE | Isoform 2 of Sorting nexin-27 OS=Mus musculus OX=10090 GN=Snx27 |
| A0A1C9IC75_MOUSE | ABCC4-N1 OS=Mus musculus OX=10090 GN=Abcc4 PE=2 SV=1 |
| tr\|Q69ZD1\|Q69ZD1_MOUSE MKIAA1699 protein (Fragment) OS=Mus musculus OX=10090 GN=Exoc4... | MKIAA1699 protein (Fragment) OS=Mus musculus OX=10090 GN=Exoc4 PE=2 SV=1 |
| sp\|O54990-3\|PROM1_MOUSE | Isoform 3 of Prominin-1 OS=Mus musculus OX=10090 GN=Prom1 |
| Q3TJS0_MOUSE | Uncharacterized protein OS=Mus musculus OX=10090 GN=Flot1 PE=2 SV=1 |

| | |
|---|---|
| B2RQ80_MOUSE | Tnik protein OS=Mus musculus OX=10090 GN=Tnik PE=1 SV=1 |
| sp\|O08582\|GTPB1_MOUSEGTP-binding protein 1 OS=Mus musculus OX=10090 GN=Gtpbp1 PE=1 SV=2 | sp\|O08582\|GTPB1_MOUSE GTP-binding protein 1 OS=Mus musculus OX=10090 GN=Gtpbp1 PE=1 SV=2 |
| A2AQA9_MOUSE | Nebulin OS=Mus musculus OX=10090 GN=Neb PE=1 SV=1 |
| NAV3_MOUSE | Neuron navigator 3 OS=Mus musculus OX=10090 GN=Nav3 PE=1 SV=2 |
| B2RQA0_MOUSE | EG331392 protein OS=Mus musculus OX=10090 GN=Gm5124 PE=2 SV=1 |
| F7BGR7_MOUSE | Predicted gene 21992 OS=Mus musculus OX=10090 GN=Gm21992 PE=1 SV=1 |
| tr\|A0A1B0GR11\|A0A1B0GR11_MOUSE Transaldolase OS=Mus musculus OX=10090 GN=Taldo1 PE=1... | Transaldolase OS=Mus musculus OX=10090 GN=Taldo1 PE=1 SV=1 |
| tr\|Q8CD98\|Q8CD98_MOUSE Uncharacterized protein OS=Mus musculus OX=10090 GN=Pfkl PE=2... | tr\|Q8CD98\|Q8CD98_MOUSE Uncharacterized protein OS=Mus musculus OX=10090 GN=Pfkl PE=2... |
| sp\|Q8VDP4\|CCAR2_MOUSECell cycle and apoptosis regulator protein 2 OS=Mus musculus OX=10090 GN=Ccar2 PE=1 SV=2 | sp\|Q8VDP4\|CCAR2_MOUSE Cell cycle and apoptosis regulator protein 2 OS=Mus musculus OX=10090 GN=Ccar2 PE=1 SV=2 |
| sp\|P01786\|HVM17_MOUSEIg heavy chain V region MOPC 47A OS=Mus musculus OX=10090 PE=1 SV=1 | sp\|P01786\|HVM17_MOUSE Ig heavy chain V region MOPC 47A OS=Mus musculus OX=10090 PE=1 SV=1 |
| sp\|B2RR83\|YTDC2_MOUSEProbable ATP-dependent RNA helicase YTHDC2 OS=Mus musculus OX=10090 GN=Ythdc2 PE=1 SV=1 | sp\|B2RR83\|YTDC2_MOUSE Probable ATP-dependent RNA helicase YTHDC2 OS=Mus musculus OX=10090 GN=Ythdc2 PE=1 SV=1 |
| A0A1L1SS82_MOUSE | Coiled-coil domain-containing 7A OS=Mus musculus OX=10090 GN=Ccdc7a PE=4 SV=1 |
| sp\|Q9Z1T6-1\|FYV1_MOUSE | Isoform 2 of 1-phosphatidylinositol 3-phosphate 5-kinase OS=Mus musculus OX=10090 GN=Pikfyve |
| B2RRH9_MOUSE | Guanine monphosphate synthetase OS=Mus musculus OX=10090 GN=Gmps PE=1 SV=1 |
| Q155P7_MOUSE | LEK1 OS=Mus musculus OX=10090 GN=Cenpf PE=1 SV=1 |

| | |
|---|---|
| B2RSI3_MOUSE | Poly [ADP-ribose] polymerase OS=Mus musculus OX=10090 GN=Parp14 PE=2 SV=1 |
| Q3TRH1_MOUSE | Uncharacterized protein (Fragment) OS=Mus musculus OX=10090 GN=Eif3a PE=2 SV=1 |
| A2AFQ2_MOUSE | 3-hydroxyacyl-CoA dehydrogenase type-2 OS=Mus musculus OX=10090 GN=Hsd17b10 PE=1 SV=1 |
| sp\|O35344\|IMA4_MOUSE Importin subunit alpha-4 OS=Mus musculus OX=10090 GN=Kpna3 PE=1... | sp\|O35344\|IMA4_MOUSE Importin subunit alpha-4 OS=Mus musculus OX=10090 GN=Kpna3 PE=1... |
| sp\|O70546-2\|KDM6A_MOUSE-DECOY Isoform 2 of Lysine-specific demethylase 6A OS=Mus... | sp\|O70546-2\|KDM6A_MOUSE-DECOY Isoform 2 of Lysine-specific demethylase 6A OS=Mus... |
| sp\|Q9QYX7-2\|PCLO_MOUSE | Isoform 2 of Protein piccolo OS=Mus musculus OX=10090 GN=Pclo |
| tr\|Q3TXG1\|Q3TXG1_MOUSE Mitogen-activated protein kinase kinase kinase 7 OS=Mus musculus... | tr\|Q3TXG1\|Q3TXG1_MOUSE Mitogen-activated protein kinase kinase kinase 7 OS=Mus musculus... |
| sp\|Q9Z1K7\|APCL_MOUSE-DECOY Adenomatous polyposis coli protein 2 OS=Mus musculus OX=10090... | sp\|Q9Z1K7\|APCL_MOUSE-DECOY Adenomatous polyposis coli protein 2 OS=Mus musculus OX=10090... |
| F8WI37_MOUSE | Histone-lysine N-methyltransferase 2C OS=Mus musculus OX=10090 GN=Kmt2c PE=1 SV=1 |
| Q8CD76_MOUSE | Kinesin light chain 1 OS=Mus musculus OX=10090 GN=Klc1 PE=1 SV=1 |
| tr\|D3YYQ8\|D3YYQ8_MOUSE Dynein, axonemal, heavy chain 10 OS=Mus musculus OX=10090... | tr\|D3YYQ8\|D3YYQ8_MOUSE Dynein, axonemal, heavy chain 10 OS=Mus musculus OX=10090... |
| tr\|Z4YJT3\|Z4YJT3_MOUSE La-related protein 1 OS=Mus musculus OX=10090 GN=Larp1 PE=1... | tr\|Z4YJT3\|Z4YJT3_MOUSE La-related protein 1 OS=Mus musculus OX=10090 GN=Larp1 PE=1... |
| sp\|Q8BW94-2\|DYH3_MOUSE-DECOY Isoform 2 of Dynein heavy chain 3, axonemal OS=Mus musculus... | sp\|Q8BW94-2\|DYH3_MOUSE-DECOY Isoform 2 of Dynein heavy chain 3, axonemal OS=Mus musculus... |
| tr\|B9EK93\|B9EK93_MOUSE Insulin receptor substrate 4 OS=Mus musculus OX=10090 GN=Irs4... | tr\|B9EK93\|B9EK93_MOUSE Insulin receptor substrate 4 OS=Mus musculus OX=10090 GN=Irs4... |
| tr\|E9QAS5\|E9QAS5_MOUSE-DECOY Chromodomain-helicase-DNA-binding protein 4 OS=Mus musculus... | tr\|E9QAS5\|E9QAS5_MOUSE-DECOY Chromodomain-helicase-DNA-binding protein 4 OS=Mus musculus... |

| | |
|---|---|
| tr\|Q8BK66\|Q8BK66_MOUSE Uncharacterized protein OS=Mus musculus OX=10090 GN=Ddx56... | tr\|Q8BK66\|Q8BK66_MOUSE Uncharacterized protein OS=Mus musculus OX=10090 GN=Ddx56... |
| RYR2_MOUSE | Ryanodine receptor 2 OS=Mus musculus OX=10090 GN=Ryr2 PE=1 SV=1 |
| PUR8_MOUSE | Adenylosuccinate lyase OS=Mus musculus OX=10090 GN=Adsl PE=1 SV=2 |
| E9PX48_MOUSE | Dedicator of cytokinesis protein 7 OS=Mus musculus OX=10090 GN=Dock7 PE=1 SV=2 |
| SC31A_MOUSE | Isoform 2 of Protein transport protein Sec31A OS=Mus musculus OX=10090 GN=Sec31a |
| Q3UGZ4_MOUSE | Spectrin beta chain OS=Mus musculus OX=10090 GN=Sptbn2 PE=2 SV=1 |
| IPO11_MOUSE | Isoform 2 of Importin-11 OS=Mus musculus OX=10090 GN=Ipo11 |
| sp\|Q7TPH6-2\|MYCB2_MOUSE-DECOY Isoform 2 of E3 ubiquitin-protein ligase MYCBP2 OS=Mus... | sp\|Q7TPH6-2\|MYCB2_MOUSE-DECOY Isoform 2 of E3 ubiquitin-protein ligase MYCBP2 OS=Mus... |
| A0A1L1SQ51_MOUSE | Talin-2 OS=Mus musculus OX=10090 GN=Tln2 PE=1 SV=1 |
| tr\|Q3TQW0\|Q3TQW0_MOUSE Uncharacterized protein (Fragment) OS=Mus musculus OX=10090... | tr\|Q3TQW0\|Q3TQW0_MOUSE Uncharacterized protein (Fragment) OS=Mus musculus OX=10090... |
| E0CZE0_MOUSE | NEDD8-activating enzyme E1 regulatory subunit OS=Mus musculus OX=10090 GN=Nae1 PE=1 SV=1 |
| O08847_MOUSE | DNA-directed RNA polymerase subunit OS=Mus musculus OX=10090 GN=Polr2a PE=2 SV=1 |
| Q5DTH1_MOUSE | MKIAA4216 protein (Fragment) OS=Mus musculus OX=10090 GN=Ube3a PE=2 SV=1 |
| Q7TN20_MOUSE | Ythdf3 protein (Fragment) OS=Mus musculus OX=10090 GN=Ythdf3 PE=2 SV=1 |
| Q80UE3_MOUSE | Protein 4.1G OS=Mus musculus OX=10090 GN=Epb41l2 PE=2 SV=1 |
| sp\|P11440\|CDK1_MOUSE Cyclin-dependent kinase 1 OS=Mus musculus OX=10090 GN=Cdk1 PE=1... | sp\|P11440\|CDK1_MOUSE Cyclin-dependent kinase 1 OS=Mus musculus OX=10090 GN=Cdk1 PE=1... |

| | |
|---|---|
| sp\|Q8C8R3-2\|ANK2_MOUSE-DECOY Isoform 2 of Ankyrin-2 OS=Mus musculus OX=10090 GN=Ank2 | sp\|Q8C8R3-2\|ANK2_MOUSE-DECOY Isoform 2 of Ankyrin-2 OS=Mus musculus OX=10090 GN=Ank2 |
| tr\|A0A0G2JF52\|A0A0G2JF52_MOUSE Actin related protein 2/3 complex, subunit 1A, isoform... | tr\|A0A0G2JF52\|A0A0G2JF52_MOUSE Actin related protein 2/3 complex, subunit 1A, isoform... |
| tr\|A0A0G2JGX4\|A0A0G2JGX4_MOUSE Sodium/potassium-transporting ATPase subunit alpha... | tr\|A0A0G2JGX4\|A0A0G2JGX4_MOUSE Sodium/potassium-transporting ATPase subunit alpha... |
| tr\|D3YU58\|D3YU58_MOUSE Anaphase-promoting complex subunit 7 OS=Mus musculus OX=10090... | Anaphase-promoting complex subunit 7 OS=Mus musculus OX=10090 GN=Anapc7 PE=1 SV=1 |
| tr\|E9Q3P4\|E9Q3P4_MOUSE-DECOY Centromere protein F OS=Mus musculus OX=10090 GN=Cenpf... | tr\|E9Q3P4\|E9Q3P4_MOUSE-DECOY Centromere protein F OS=Mus musculus OX=10090 GN=Cenpf... |
| tr\|G3UWR2\|G3UWR2_MOUSE GRB10-interacting GYF protein 2 OS=Mus musculus OX=10090 GN=Gigyf2... | tr\|G3UWR2\|G3UWR2_MOUSE GRB10-interacting GYF protein 2 OS=Mus musculus OX=10090 GN=Gigyf2... |
| tr\|Q6NZK2\|Q6NZK2_MOUSE-DECOY NCK-associated protein 5 OS=Mus musculus OX=10090 GN=Nckap5... | tr\|Q6NZK2\|Q6NZK2_MOUSE-DECOY NCK-associated protein 5 OS=Mus musculus OX=10090 GN=Nckap5... |
| A2AMU9_MOUSE | Ribosome assembly factor mrt4 OS=Mus musculus OX=10090 GN=Mrto4 PE=1 SV=1 |
| E9PV41_MOUSE | Protein diaphanous homolog 1 OS=Mus musculus OX=10090 GN=Diaph1 PE=1 SV=1 |
| E9Q043_MOUSE | Fibronectin type III domain-containing 1 OS=Mus musculus OX=10090 GN=Fndc1 PE=1 SV=1 |
| tr\|Q571A1\|Q571A1_MOUSE MKIAA0765 splice variant 1 (Fragment) OS=Mus musculus OX=10090... | tr\|Q571A1\|Q571A1_MOUSE MKIAA0765 splice variant 1 (Fragment) OS=Mus musculus OX=10090... |
| A0A1W2P7Q6_MOUSE | Exportin-T OS=Mus musculus OX=10090 GN=Xpot PE=1 SV=1 |
| A7TU71_MOUSE | Protein Shroom2 OS=Mus musculus OX=10090 GN=Shroom2 PE=1 SV=1 |
| tr\|A2A520\|A2A520_MOUSE-DECOY Dynein heavy chain 17, axonemal OS=Mus musculus OX=10090... | tr\|A2A520\|A2A520_MOUSE-DECOY Dynein heavy chain 17, axonemal OS=Mus musculus OX=10090... |
| tr\|E9Q1M6\|E9Q1M6_MOUSE Ankyrin repeat and KH domain-containing 1 OS=Mus musculus... | tr\|E9Q1M6\|E9Q1M6_MOUSE Ankyrin repeat and KH domain-containing 1 OS=Mus musculus... |

| | |
|---|---|
| E9Q6A7_MOUSE | Bromodomain PHD finger transcription factor OS=Mus musculus OX=10090 GN=Bptf PE=1 SV=1 |
| Q9CWR5_MOUSE | Serine hydroxymethyltransferase OS=Mus musculus OX=10090 GN=Shmt1 PE=2 SV=1 |
| sp\|Q78DX7\|ROS1_MOUSE Proto-oncogene tyrosine-protein kinase ROS OS=Mus musculus OX=10090... | sp\|Q78DX7\|ROS1_MOUSE Proto-oncogene tyrosine-protein kinase ROS OS=Mus musculus OX=10090... |
| D3Z667_MOUSE | Dynein heavy chain 2, axonemal OS=Mus musculus OX=10090 GN=Dnah2 PE=1 SV=1 |
| tr\|Q5SUC3\|Q5SUC3_MOUSE Calnexin, isoform CRA_a OS=Mus musculus OX=10090 GN=Canx PE=1... | tr\|Q5SUC3\|Q5SUC3_MOUSE Calnexin, isoform CRA_a OS=Mus musculus OX=10090 GN=Canx PE=1... |
| sp\|Q8C0C7\|SYFA_MOUSE Phenylalanine--tRNA ligase alpha subunit OS=Mus musculus OX=10090... | sp\|Q8C0C7\|SYFA_MOUSE Phenylalanine--tRNA ligase alpha subunit OS=Mus musculus OX=10090... |
| sp\|Q8BP67\|RL24_MOUSE 60S ribosomal protein L24 OS=Mus musculus OX=10090 GN=Rpl24... | sp\|Q8BP67\|RL24_MOUSE 60S ribosomal protein L24 OS=Mus musculus OX=10090 GN=Rpl24... |
| A0A087WSE3_MOUSE | Striated muscle-specific serine/threonine-protein kinase (Fragment) OS=Mus musculus OX=10090 GN=Speg PE=1 SV=1 |
| sp\|E9Q401\|RYR2_MOUSE-DECOY Ryanodine receptor 2 OS=Mus musculus OX=10090 GN=Ryr2... | sp\|E9Q401\|RYR2_MOUSE-DECOY Ryanodine receptor 2 OS=Mus musculus OX=10090 GN=Ryr2... |
| tr\|D0VYV7\|D0VYV7_MOUSE Erythrocyte protein band 4.1-like 3 isoform C OS=Mus musculus... | Erythrocyte protein band 4.1-like 3 isoform C OS=Mus musculus OX=10090 GN=Epb41l3 PE=2 SV=1 |
| tr\|Q8BTJ1\|Q8BTJ1_MOUSE Phosphoserine aminotransferase OS=Mus musculus OX=10090 GN=Psat1... | Phosphoserine aminotransferase OS=Mus musculus OX=10090 GN=Psat1 PE=2 SV=1 |
| Q3UDN8_MOUSE | Uncharacterized protein (Fragment) OS=Mus musculus OX=10090 GN=Trim28 PE=2 SV=1 |
| ATRN_MOUSE | Attractin OS=Mus musculus OX=10090 GN=Atrn PE=1 SV=3 |
| sp\|Q9WVA4\|TAGL2_MOUSETransgelin-2 OS=Mus musculus OX=10090 GN=Tagln2 PE=1 SV=4 | sp\|Q9WVA4\|TAGL2_MOUSE Transgelin-2 OS=Mus musculus OX=10090 GN=Tagln2 PE=1 SV=4 |
| Q9D6F6_MOUSE | Dynein light chain OS=Mus musculus OX=10090 GN=Dynll1 PE=2 SV=1 |

| | |
|---|---|
| sp\|Q60974-2\|NCOR1_MOUSE-DECOY Isoform 2 of Nuclear receptor corepressor 1 OS=Mus... | sp\|Q60974-2\|NCOR1_MOUSE-DECOY Isoform 2 of Nuclear receptor corepressor 1 OS=Mus... |
| NCPR_MOUSE | NADPH--cytochrome P450 reductase OS=Mus musculus OX=10090 GN=Por PE=1 SV=2 |
| DPP3_MOUSE | Dipeptidyl peptidase 3 OS=Mus musculus OX=10090 GN=Dpp3 PE=1 SV=2 |
| A0A0A6YX18_MOUSE | V-type proton ATPase subunit H OS=Mus musculus OX=10090 GN=Atp6v1h PE=1 SV=1 |
| tr\|G5E8B1\|G5E8B1_MOUSE-DECOY Protein tyrosine phosphatase, non-receptor type 13 OS=Mus... | tr\|G5E8B1\|G5E8B1_MOUSE-DECOY Protein tyrosine phosphatase, non-receptor type 13 OS=Mus... |
| E9QK83_MOUSE | Small subunit processome component 20 homolog OS=Mus musculus OX=10090 GN=Utp20 PE=1 SV=1 |
| tr\|B2RX08\|B2RX08_MOUSE-DECOY Spectrin beta chain OS=Mus musculus OX=10090 GN=Sptb... | tr\|B2RX08\|B2RX08_MOUSE-DECOY Spectrin beta chain OS=Mus musculus OX=10090 GN=Sptb... |
| A0A076N9U7_MOUSE | Celsr3 OS=Mus musculus OX=10090 GN=Celsr3 PE=2 SV=1 |
| A2A6U3_MOUSE | Septin-9 OS=Mus musculus OX=10090 GN=Sept9 PE=1 SV=1 |
| sp\|E9PVX6-2\|KI67_MOUSE Isoform 2 of Proliferation marker protein Ki-67 OS=Mus musculus... | Isoform 2 of Proliferation marker protein Ki-67 OS=Mus musculus OX=10090 GN=Mki67 |
| APCL_MOUSE | Adenomatous polyposis coli protein 2 OS=Mus musculus OX=10090 GN=Apc2 PE=1 SV=1 |
| sp\|P46664\|PURA2_MOUSEAdenylosuccinate synthetase isozyme 2 OS=Mus musculus OX=10090 GN=Adss PE=1 SV=2 | sp\|P46664\|PURA2_MOUSE Adenylosuccinate synthetase isozyme 2 OS=Mus musculus OX=10090 GN=Adss PE=1 SV=2 |
| sp\|Q9Z2U1\|PSA5_MOUSE Proteasome subunit alpha type-5 OS=Mus musculus OX=10090 GN=Psma5... | sp\|Q9Z2U1\|PSA5_MOUSE Proteasome subunit alpha type-5 OS=Mus musculus OX=10090 GN=Psma5... |
| VP13A_MOUSE | Isoform 2 of Vacuolar protein sorting-associated protein 13A OS=Mus musculus OX=10090 GN=Vps13a |
| E9QM38_MOUSE | Solute carrier family 12 member 2 OS=Mus musculus OX=10090 GN=Slc12a2 PE=1 SV=1 |

| | |
|---|---|
| sp\|Q9CSH3\|RRP44_MOUSEExosome complex exonuclease RRP44 OS=Mus musculus OX=10090 GN=Dis3 PE=1 SV=4 | sp\|Q9CSH3\|RRP44_MOUSE Exosome complex exonuclease RRP44 OS=Mus musculus OX=10090 GN=Dis3 PE=1 SV=4 |
| A0A1W2P6U8_MOUSE | RIKEN cDNA 4932415D10 gene OS=Mus musculus OX=10090 GN=4932415D10Rik PE=4 SV=1 |
| CAP1_MOUSE | Adenylyl cyclase-associated protein 1 OS=Mus musculus OX=10090 GN=Cap1 PE=1 SV=4 |
| sp\|Q9CZS1\|AL1B1_MOUSEAldehyde dehydrogenase X, mitochondrial OS=Mus musculus OX=10090 GN=Aldh1b1 PE=1 SV=1 | sp\|Q9CZS1\|AL1B1_MOUSE Aldehyde dehydrogenase X, mitochondrial OS=Mus musculus OX=10090 GN=Aldh1b1 PE=1 SV=1 |
| Q3U1K2_MOUSE | Uncharacterized protein (Fragment) OS=Mus musculus OX=10090 GN=Atp2a3 PE=2 SV=1 |
| sp\|P27641\|XRCC5_MOUSEX-ray repair cross-complementing protein 5 OS=Mus musculus OX=10090 GN=Xrcc5 PE=1 SV=4 | sp\|P27641\|XRCC5_MOUSE X-ray repair cross-complementing protein 5 OS=Mus musculus OX=10090 GN=Xrcc5 PE=1 SV=4 |
| tr\|E9Q0B6\|E9Q0B6_MOUSE-DECOY Dynein, axonemal, heavy chain 6 OS=Mus musculus OX=10090... | tr\|E9Q0B6\|E9Q0B6_MOUSE-DECOY Dynein, axonemal, heavy chain 6 OS=Mus musculus OX=10090... |
| ARF5_MOUSE | ADP-ribosylation factor 5 OS=Mus musculus OX=10090 GN=Arf5 PE=1 SV=2 |
| NXN_MOUSE | Nucleoredoxin OS=Mus musculus OX=10090 GN=Nxn PE=1 SV=1 |
| sp\|P48678\|LMNA_MOUSE | Prelamin-A/C OS=Mus musculus OX=10090 GN=Lmna PE=1 SV=2 |
| sp\|P97313-2\|PRKDC_MOUSE Isoform 2 of DNA-dependent protein kinase catalytic subunit... | sp\|P97313-2\|PRKDC_MOUSE Isoform 2 of DNA-dependent protein kinase catalytic subunit... |
| sp\|Q6P2B1-2\|TNPO3_MOUSE | Isoform 2 of Transportin-3 OS=Mus musculus OX=10090 GN=Tnpo3 |
| tr\|E9Q8T1\|E9Q8T1_MOUSE Transforming acidic coiled-coil-containing protein 2 OS=Mus... | tr\|E9Q8T1\|E9Q8T1_MOUSE Transforming acidic coiled-coil-containing protein 2 OS=Mus... |
| tr\|A2BH46\|A2BH46_MOUSE RNA cytidine acetyltransferase (Fragment) OS=Mus musculus... | tr\|A2BH46\|A2BH46_MOUSE RNA cytidine acetyltransferase (Fragment) OS=Mus musculus... |
| sp\|P70333\|HNRH2_MOUSEHeterogeneous nuclear ribonucleoprotein H2 OS=Mus musculus OX=10090 GN=Hnrnph2 PE=1 SV=1 | sp\|P70333\|HNRH2_MOUSE Heterogeneous nuclear ribonucleoprotein H2 OS=Mus |

| | |
|---|---|
| | musculus OX=10090 GN=Hnrnph2 PE=1 SV=1 |
| VPP1_MOUSE | V-type proton ATPase 116 kDa subunit a isoform 1 OS=Mus musculus OX=10090 GN=Atp6v0a1 PE=1 SV=3 |
| B2RX66_MOUSE | MCG124812 OS=Mus musculus OX=10090 GN=Taok1 PE=1 SV=1 |
| ECHA_MOUSE | Trifunctional enzyme subunit alpha, mitochondrial OS=Mus musculus OX=10090 GN=Hadha PE=1 SV=1 |
| A0A1B0GR10_MOUSE | Predicted gene, 40460 OS=Mus musculus OX=10090 GN=Gm40460 PE=4 SV=1 |
| tr\|E9PWM3\|E9PWM3_MOUSE-DECOY Armadillo repeat-containing, X-linked 4 OS=Mus musculus... | tr\|E9PWM3\|E9PWM3_MOUSE-DECOY Armadillo repeat-containing, X-linked 4 OS=Mus musculus... |
| tr\|A0A1B0GS68\|A0A1B0GS68_MOUSE Predicted gene 45713 OS=Mus musculus OX=10090 GN=Gm45713... | tr\|A0A1B0GS68\|A0A1B0GS68_MOUSE Predicted gene 45713 OS=Mus musculus OX=10090 GN=Gm45713... |
| Q8BYP9_MOUSE | Uncharacterized protein OS=Mus musculus OX=10090 GN=Atl2 PE=2 SV=1 |
| E9Q9C6_MOUSE | Fc fragment of IgG-binding protein OS=Mus musculus OX=10090 GN=Fcgbp PE=1 SV=1 |
| sp\|Q01320\|TOP2A_MOUSEDNA topoisomerase 2-alpha OS=Mus musculus OX=10090 GN=Top2a PE=1 SV=2 | sp\|Q01320\|TOP2A_MOUSE DNA topoisomerase 2-alpha OS=Mus musculus OX=10090 GN=Top2a PE=1 SV=2 |
| E9QJU8_MOUSE | Transformation-related protein 53-binding protein 2 OS=Mus musculus OX=10090 GN=Trp53bp2 PE=1 SV=1 |
| A0A1B0GT92_MOUSE | Glycogen [starch] synthase OS=Mus musculus OX=10090 GN=Gys1 PE=1 SV=1 |
| S4R2J9_MOUSE | Protein PRRC2C OS=Mus musculus OX=10090 GN=Prrc2c PE=1 SV=1 |
| ELP1_MOUSE | Elongator complex protein 1 OS=Mus musculus OX=10090 GN=Elp1 PE=1 SV=2 |
| A0A087WQS2_MOUSE | Basic leucine zipper and W2 domain-containing protein 1 OS=Mus musculus OX=10090 GN=Bzw1 PE=1 SV=1 |
| E9QPE7_MOUSE | Myosin-11 OS=Mus musculus OX=10090 GN=Myh11 PE=1 SV=1 |
| ACACA_MOUSE | Isoform 2 of Acetyl-CoA carboxylase 1 OS=Mus musculus OX=10090 GN=Acaca |

| | |
|---|---|
| Q6PFV8_MOUSE | Cacna1g protein OS=Mus musculus OX=10090 GN=Cacna1g PE=2 SV=1 |
| sp\|P97384\|ANX11_MOUSEAnnexin A11 OS=Mus musculus OX=10090 GN=Anxa11 PE=1 SV=2 | sp\|P97384\|ANX11_MOUSE Annexin A11 OS=Mus musculus OX=10090 GN=Anxa11 PE=1 SV=2 |
| C7G3P2_MOUSE | MKIAA0536 protein (Fragment) OS=Mus musculus OX=10090 GN=Prpf4b PE=2 SV=1 |
| Q3UAY3_MOUSE | Uncharacterized protein OS=Mus musculus OX=10090 GN=Gtf3c4 PE=2 SV=1 |
| sp\|P56480\|ATPB_MOUSE ATP synthase subunit beta, mitochondrial OS=Mus musculus OX=10090... | ATP synthase subunit beta, mitochondrial OS=Mus musculus OX=10090 GN=Atp5f1b PE=1 SV=2 |
| sp\|A2AWL7-2\|MGAP_MOUSE-DECOY Isoform 2 of MAX gene-associated protein OS=Mus musculus... | sp\|A2AWL7-2\|MGAP_MOUSE-DECOY Isoform 2 of MAX gene-associated protein OS=Mus musculus... |
| sp\|Q8CGC7\|SYEP_MOUSE-DECOY Bifunctional glutamate/proline--tRNA ligase OS=Mus musculus... | sp\|Q8CGC7\|SYEP_MOUSE-DECOY Bifunctional glutamate/proline--tRNA ligase OS=Mus musculus... |
| E9QAE4_MOUSE | Histone-lysine N-methyltransferase, H3 lysine-36 and H4 lysine-20-specific OS=Mus musculus OX=10090 GN=Nsd1 PE=1 SV=1 |
| sp\|Q91XQ0-2\|DYH8_MOUSE Isoform 2 of Dynein heavy chain 8, axonemal OS=Mus musculus... | sp\|Q91XQ0-2\|DYH8_MOUSE Isoform 2 of Dynein heavy chain 8, axonemal OS=Mus musculus... |
| tr\|E9Q526\|E9Q526_MOUSE-DECOY C2 domain-containing protein 3 OS=Mus musculus OX=10090... | tr\|E9Q526\|E9Q526_MOUSE-DECOY C2 domain-containing protein 3 OS=Mus musculus OX=10090... |
| tr\|K3W4Q5\|K3W4Q5_MOUSE-DECOY Family with sequence similarity 186, member A OS=Mus... | tr\|K3W4Q5\|K3W4Q5_MOUSE-DECOY Family with sequence similarity 186, member A OS=Mus... |
| tr\|Q69Z43\|Q69Z43_MOUSE MKIAA1995 protein (Fragment) OS=Mus musculus OX=10090 GN=Nek9... | tr\|Q69Z43\|Q69Z43_MOUSE MKIAA1995 protein (Fragment) OS=Mus musculus OX=10090 GN=Nek9... |
| sp\|O54988\|SLK_MOUSE STE20-like serine/threonine-protein kinase OS=Mus musculus OX=10090... | sp\|O54988\|SLK_MOUSE STE20-like serine/threonine-protein kinase OS=Mus musculus OX=10090... |
| sp\|P68404\|KPCB_MOUSE | Protein kinase C beta type OS=Mus musculus OX=10090 GN=Prkcb PE=1 SV=4 |

| | |
|---|---|
| sp\|Q6PR54-3\|RIF1_MOUSE-DECOY Isoform 3 of Telomere-associated protein RIF1 OS=Mus... | sp\|Q6PR54-3\|RIF1_MOUSE-DECOY Isoform 3 of Telomere-associated protein RIF1 OS=Mus... |
| A0A2I3BR78_MOUSE | Exosome complex exonuclease RRP44 OS=Mus musculus OX=10090 GN=Dis3 PE=1 SV=1 |
| A2AL50_MOUSE | Alkylglycerone-phosphate synthase OS=Mus musculus OX=10090 GN=Agps PE=1 SV=1 |
| E9Q855_MOUSE | Secretory carrier-associated membrane protein OS=Mus musculus OX=10090 GN=Scamp3 PE=1 SV=1 |
| GLE1_MOUSE | Nucleoporin GLE1 OS=Mus musculus OX=10090 GN=Gle1 PE=1 SV=2 |
| sp\|A2AAE1-6\|K1109_MOUSE | Isoform 6 of Uncharacterized protein KIAA1109 OS=Mus musculus OX=10090 GN=Kiaa1109 |
| sp\|O35654\|DPOD2_MOUSEDNA polymerase delta subunit 2 OS=Mus musculus OX=10090 GN=Pold2 PE=1 SV=2 | sp\|O35654\|DPOD2_MOUSE DNA polymerase delta subunit 2 OS=Mus musculus OX=10090 GN=Pold2 PE=1 SV=2 |
| sp\|Q3TDN2-2\|FAF2_MOUSE Isoform 2 of FAS-associated factor 2 OS=Mus musculus OX=10090... | sp\|Q3TDN2-2\|FAF2_MOUSE Isoform 2 of FAS-associated factor 2 OS=Mus musculus OX=10090... |
| sp\|Q61315\|APC_MOUSE-DECOY Adenomatous polyposis coli protein OS=Mus musculus OX=10090... | sp\|Q61315\|APC_MOUSE-DECOY Adenomatous polyposis coli protein OS=Mus musculus OX=10090... |
| sp\|Q62440-2\|TLE1_MOUSE Isoform 2 of Transducin-like enhancer protein 1 OS=Mus musculus... | sp\|Q62440-2\|TLE1_MOUSE Isoform 2 of Transducin-like enhancer protein 1 OS=Mus musculus... |
| sp\|Q64521\|GPDM_MOUSE Glycerol-3-phosphate dehydrogenase, mitochondrial OS=Mus musculus... | sp\|Q64521\|GPDM_MOUSE Glycerol-3-phosphate dehydrogenase, mitochondrial OS=Mus musculus... |
| sp\|Q99J95-2\|CDK9_MOUSE Isoform 2 of Cyclin-dependent kinase 9 OS=Mus musculus OX=10090... | Isoform 2 of Cyclin-dependent kinase 9 OS=Mus musculus OX=10090 GN=Cdk9 |
| tr\|A0A0N4SVC2\|A0A0N4SVC2_MOUSE Transformer-2 protein homolog alpha OS=Mus musculus... | tr\|A0A0N4SVC2\|A0A0N4SVC2_MOUSE Transformer-2 protein homolog alpha OS=Mus musculus... |
| tr\|A0A1L1SQ51\|A0A1L1SQ51_MOUSE-DECOY Talin-2 OS=Mus musculus OX=10090 GN=Tln2 PE=1... | tr\|A0A1L1SQ51\|A0A1L1SQ51_MOUSE-DECOY Talin-2 OS=Mus musculus OX=10090 GN=Tln2 PE=1... |

| | |
|---|---|
| tr\|Q3UNM2\|Q3UNM2_MOUSE Uncharacterized protein (Fragment) OS=Mus musculus OX=10090... | tr\|Q3UNM2\|Q3UNM2_MOUSE Uncharacterized protein (Fragment) OS=Mus musculus OX=10090... |
| tr\|Q80ZU1\|Q80ZU1_MOUSE WD repeat domain 68 OS=Mus musculus OX=10090 GN=Dcaf7 PE=2... | WD repeat domain 68 OS=Mus musculus OX=10090 GN=Dcaf7 PE=2 SV=1 |
| sp\|Q921N6-2\|DDX27_MOUSE Isoform 2 of Probable ATP-dependent RNA helicase DDX27 OS=Mus... | sp\|Q921N6-2\|DDX27_MOUSE Isoform 2 of Probable ATP-dependent RNA helicase DDX27 OS=Mus... |
| tr\|F6TN03\|F6TN03_MOUSE Ribosome maturation protein SBDS (Fragment) OS=Mus musculus... | Ribosome maturation protein SBDS (Fragment) OS=Mus musculus OX=10090 GN=Sbds PE=1 SV=1 |
| COG5_MOUSE | Conserved oligomeric Golgi complex subunit 5 OS=Mus musculus OX=10090 GN=Cog5 PE=1 SV=3 |
| sp\|E9Q5F9-2\|SETD2_MOUSE Isoform 2 of Histone-lysine N-methyltransferase SETD2 OS=Mus... | Isoform 2 of Histone-lysine N-methyltransferase SETD2 OS=Mus musculus OX=10090 GN=Setd2 |
| tr\|Q8BL36\|Q8BL36_MOUSE Uncharacterized protein OS=Mus musculus OX=10090 GN=Fam98a... | tr\|Q8BL36\|Q8BL36_MOUSE Uncharacterized protein OS=Mus musculus OX=10090 GN=Fam98a... |
| tr\|Q7TNG4\|Q7TNG4_MOUSE Smc1a protein (Fragment) OS=Mus musculus OX=10090 GN=Smc1a... | tr\|Q7TNG4\|Q7TNG4_MOUSE Smc1a protein (Fragment) OS=Mus musculus OX=10090 GN=Smc1a... |
| Q3TI21_MOUSE | Uncharacterized protein OS=Mus musculus OX=10090 GN=Ddx41 PE=2 SV=1 |
| sp\|P27612\|PLAP_MOUSE Phospholipase A-2-activating protein OS=Mus musculus OX=10090... | sp\|P27612\|PLAP_MOUSE Phospholipase A-2-activating protein OS=Mus musculus OX=10090... |
| Q3UI30_MOUSE | Uncharacterized protein OS=Mus musculus OX=10090 GN=Ewsr1 PE=2 SV=1 |
| sp\|Q8VEK3-2\|HNRPU_MOUSE Isoform 2 of Heterogeneous nuclear ribonucleoprotein U OS=Mus... | Isoform 2 of Heterogeneous nuclear ribonucleoprotein U OS=Mus musculus OX=10090 GN=Hnrnpu |
| tr\|D3YWC5\|D3YWC5_MOUSE Cullin-associated NEDD8-dissociated protein 1 (Fragment) OS=Mus... | Cullin-associated NEDD8-dissociated protein 1 (Fragment) OS=Mus musculus OX=10090 GN=Cand1 PE=1 SV=1 |
| EP400_MOUSE | Isoform 4 of E1A-binding protein p400 OS=Mus musculus OX=10090 GN=Ep400 |
| tr\|Q9CX79\|Q9CX79_MOUSE Uncharacterized protein OS=Mus musculus OX=10090 GN=Ntpcr... | tr\|Q9CX79\|Q9CX79_MOUSE Uncharacterized protein OS=Mus musculus OX=10090 GN=Ntpcr... |

| | |
|---|---|
| tr\|Q6SLK2\|Q6SLK2_MOUSE Protein kinase lysine deficient 1 OS=Mus musculus OX=10090... | tr\|Q6SLK2\|Q6SLK2_MOUSE Protein kinase lysine deficient 1 OS=Mus musculus OX=10090... |
| A8DUV3_MOUSE | Alpha-globin OS=Mus musculus OX=10090 GN=Hbat1 PE=3 SV=1 |
| sp\|Q61687\|ATRX_MOUSE-DECOY Transcriptional regulator ATRX OS=Mus musculus OX=10090... | sp\|Q61687\|ATRX_MOUSE-DECOY Transcriptional regulator ATRX OS=Mus musculus OX=10090... |
| tr\|F8WGX5\|F8WGX5_MOUSE-DECOY DmX-like protein 1 OS=Mus musculus OX=10090 GN=Dmxl1... | tr\|F8WGX5\|F8WGX5_MOUSE-DECOY DmX-like protein 1 OS=Mus musculus OX=10090 GN=Dmxl1... |
| A0A0R4J1Z3_MOUSE | Transmembrane protein 33 OS=Mus musculus OX=10090 GN=Tmem33 PE=1 SV=1 |
| tr\|Q8BWM7\|Q8BWM7_MOUSE Uncharacterized protein (Fragment) OS=Mus musculus OX=10090... | tr\|Q8BWM7\|Q8BWM7_MOUSE Uncharacterized protein (Fragment) OS=Mus musculus OX=10090... |
| tr\|E9Q1M6\|E9Q1M6_MOUSE-DECOY Ankyrin repeat and KH domain-containing 1 OS=Mus musculus... | tr\|E9Q1M6\|E9Q1M6_MOUSE-DECOY Ankyrin repeat and KH domain-containing 1 OS=Mus musculus... |
| tr\|A0A1Y7VLA4\|A0A1Y7VLA4_MOUSE-DECOY Protein FAM208B (Fragment) OS=Mus musculus OX=10090... | tr\|A0A1Y7VLA4\|A0A1Y7VLA4_MOUSE-DECOY Protein FAM208B (Fragment) OS=Mus musculus OX=10090... |
| sp\|Q6IFX2\|K1C42_MOUSEKeratin, type I cytoskeletal 42 OS=Mus musculus OX=10090 GN=Krt42 PE=1 SV=1 | sp\|Q6IFX2\|K1C42_MOUSE Keratin, type I cytoskeletal 42 OS=Mus musculus OX=10090 GN=Krt42 PE=1 SV=1 |
| tr\|E9QPI5\|E9QPI5_MOUSE Sister chromatid cohesion protein PDS5 homolog A OS=Mus musculus... | Sister chromatid cohesion protein PDS5 homolog A OS=Mus musculus OX=10090 GN=Pds5a PE=1 SV=1 |
| Q3TLF6_MOUSE | Uncharacterized protein OS=Mus musculus OX=10090 GN=Mybbp1a PE=2 SV=1 |
| tr\|E9PZP8\|E9PZP8_MOUSE-DECOY HECT and RLD domain-containing E3 ubiquitin protein... | tr\|E9PZP8\|E9PZP8_MOUSE-DECOY HECT and RLD domain-containing E3 ubiquitin protein... |
| E9PWN2_MOUSE | Isoleucine--tRNA ligase, mitochondrial OS=Mus musculus OX=10090 GN=Iars2 PE=1 SV=1 |
| A0A0R4J1D0_MOUSE | Copine-2 OS=Mus musculus OX=10090 GN=Cpne2 PE=1 SV=1 |
| Q3TKX4_MOUSE | Uncharacterized protein (Fragment) OS=Mus musculus OX=10090 GN=Ddx52 PE=2 SV=1 |

| | |
|---|---|
| Q05CX1_MOUSE | Mgea5 protein (Fragment) OS=Mus musculus OX=10090 GN=Mgea5 PE=2 SV=1 |
| A2ACG7_MOUSE | Dolichyl-diphosphooligosaccharide--protein glycosyltransferase subunit 2 OS=Mus musculus OX=10090 GN=Rpn2 PE=1 SV=1 |
| TXTP_MOUSE | Tricarboxylate transport protein, mitochondrial OS=Mus musculus OX=10090 GN=Slc25a1 PE=1 SV=1 |
| A1A596_MOUSE | Pwp2 protein OS=Mus musculus OX=10090 GN=Pwp2 PE=2 SV=1 |
| Q9CS46_MOUSE | Uncharacterized protein (Fragment) OS=Mus musculus OX=10090 GN=Ilkap PE=2 SV=1 |
| Q3TJD0_MOUSE | Uncharacterized protein OS=Mus musculus OX=10090 GN=Sec61a1 PE=2 SV=1 |
| NOL8_MOUSE | Nucleolar protein 8 OS=Mus musculus OX=10090 GN=Nol8 PE=1 SV=2 |
| tr\|A2AFF6\|A2AFF6_MOUSE Cohesin subunit SA-2 OS=Mus musculus OX=10090 GN=Stag2 PE=1... | Cohesin subunit SA-2 OS=Mus musculus OX=10090 GN=Stag2 PE=1 SV=1 |
| Q3TFH8_MOUSE | Phosphoacetylglucosamine mutase OS=Mus musculus OX=10090 GN=Pgm3 PE=2 SV=1 |
| tr\|B1AQ77\|B1AQ77_MOUSE Keratin 15, isoform CRA_a OS=Mus musculus OX=10090 GN=Krt15... | Keratin 15, isoform CRA_a OS=Mus musculus OX=10090 GN=Krt15 PE=1 SV=1 |
| sp\|Q9ESW4\|AGK_MOUSE Acylglycerol kinase, mitochondrial OS=Mus musculus OX=10090 GN=Agk... | sp\|Q9ESW4\|AGK_MOUSE Acylglycerol kinase, mitochondrial OS=Mus musculus OX=10090 GN=Agk... |
| tr\|F8VPZ5\|F8VPZ5_MOUSE-DECOY Excision repair cross-complementing rodent repair deficiency,... | tr\|F8VPZ5\|F8VPZ5_MOUSE-DECOY Excision repair cross-complementing rodent repair deficiency,... |
| sp\|Q3TIX9\|SNUT2_MOUSEU4/U6.U5 tri-snRNP-associated protein 2 OS=Mus musculus OX=10090 GN=Usp39 PE=1 SV=2 | sp\|Q3TIX9\|SNUT2_MOUSE U4/U6.U5 tri-snRNP-associated protein 2 OS=Mus musculus OX=10090 GN=Usp39 PE=1 SV=2 |
| tr\|A0A0R4J0B7\|A0A0R4J0B7_MOUSE Protein SDA1 homolog OS=Mus musculus OX=10090 GN=Sdad1... | Protein SDA1 homolog OS=Mus musculus OX=10090 GN=Sdad1 PE=1 SV=1 |
| Q3TEB0_MOUSE | Uncharacterized protein OS=Mus musculus OX=10090 GN=Pspc1 PE=2 SV=1 |

| | |
|---|---|
| tr\|D3Z2X2\|D3Z2X2_MOUSE Dynein heavy chain domain 1 OS=Mus musculus OX=10090 GN=Dnhd1... | tr\|D3Z2X2\|D3Z2X2_MOUSE Dynein heavy chain domain 1 OS=Mus musculus OX=10090 GN=Dnhd1... |
| tr\|E9Q264\|E9Q264_MOUSE-DECOY Myosin, heavy chain 15 OS=Mus musculus OX=10090 GN=Myh15... | tr\|E9Q264\|E9Q264_MOUSE-DECOY Myosin, heavy chain 15 OS=Mus musculus OX=10090 GN=Myh15... |
| tr\|D6RIN1\|D6RIN1_MOUSE Protein unc-45 homolog A OS=Mus musculus OX=10090 GN=Unc45a... | tr\|D6RIN1\|D6RIN1_MOUSE Protein unc-45 homolog A OS=Mus musculus OX=10090 GN=Unc45a... |
| tr\|A0A0G2JEA5\|A0A0G2JEA5_MOUSE Predicted gene 43738 OS=Mus musculus OX=10090 GN=Gm43738... | tr\|A0A0G2JEA5\|A0A0G2JEA5_MOUSE Predicted gene 43738 OS=Mus musculus OX=10090 GN=Gm43738... |
| RICTR_MOUSE | Isoform 2 of Rapamycin-insensitive companion of mTOR OS=Mus musculus OX=10090 GN=Rictor |
| tr\|Q8C872\|Q8C872_MOUSE Transferrin receptor protein 1 OS=Mus musculus OX=10090 GN=Tfrc... | Transferrin receptor protein 1 OS=Mus musculus OX=10090 GN=Tfrc PE=1 SV=1 |
| tr\|H9KV04\|H9KV04_MOUSE Mini-chromosome maintenance complex-binding protein OS=Mus... | tr\|H9KV04\|H9KV04_MOUSE Mini-chromosome maintenance complex-binding protein OS=Mus... |
| A0A087WRK0_MOUSE | CLIP-associating protein 1 OS=Mus musculus OX=10090 GN=Clasp1 PE=1 SV=2 |
| SIN3A_MOUSE | Isoform 2 of Paired amphipathic helix protein Sin3a OS=Mus musculus OX=10090 GN=Sin3a |
| sp\|Q3UYV9\|NCBP1_MOUSENuclear cap-binding protein subunit 1 OS=Mus musculus OX=10090 GN=Ncbp1 PE=1 SV=2 | sp\|Q3UYV9\|NCBP1_MOUSE Nuclear cap-binding protein subunit 1 OS=Mus musculus OX=10090 GN=Ncbp1 PE=1 SV=2 |
| sp\|A2A6Q5-3\|CDC27_MOUSE | Isoform 3 of Cell division cycle protein 27 homolog OS=Mus musculus OX=10090 GN=Cdc27 |
| sp\|Q9D6F9\|TBB4A_MOUSETubulin beta-4A chain OS=Mus musculus OX=10090 GN=Tubb4a PE=1 SV=3 | sp\|Q9D6F9\|TBB4A_MOUSE Tubulin beta-4A chain OS=Mus musculus OX=10090 GN=Tubb4a PE=1 SV=3 |
| tr\|A0A0R4J0K2\|A0A0R4J0K2_MOUSE-DECOY Cytoskeleton-associated protein 5 OS=Mus musculus... | tr\|A0A0R4J0K2\|A0A0R4J0K2_MOUSE-DECOY Cytoskeleton-associated protein 5 OS=Mus musculus... |
| tr\|B7ZP20\|B7ZP20_MOUSE Ankyrin repeat and FYVE domain containing 1 OS=Mus musculus... | tr\|B7ZP20\|B7ZP20_MOUSE Ankyrin repeat and FYVE domain containing 1 OS=Mus musculus... |

| | |
|---|---|
| sp\|P68510\|1433F_MOUSE14-3-3 protein eta OS=Mus musculus OX=10090 GN=Ywhah PE=1 SV=2 | sp\|P68510\|1433F_MOUSE 14-3-3 protein eta OS=Mus musculus OX=10090 GN=Ywhah PE=1 SV=2 |
| sp\|Q9EQF6\|DPYL5_MOUSEDihydropyrimidinase-related protein 5 OS=Mus musculus OX=10090 GN=Dpysl5 PE=1 SV=1 | sp\|Q9EQF6\|DPYL5_MOUSE Dihydropyrimidinase-related protein 5 OS=Mus musculus OX=10090 GN=Dpysl5 PE=1 SV=1 |
| sp\|Q80TN7\|NAV3_MOUSE-DECOY Neuron navigator 3 OS=Mus musculus OX=10090 GN=Nav3 PE=1... | sp\|Q80TN7\|NAV3_MOUSE-DECOY Neuron navigator 3 OS=Mus musculus OX=10090 GN=Nav3 PE=1... |
| Q8BP60_MOUSE | Uncharacterized protein OS=Mus musculus OX=10090 PE=2 SV=1 |
| tr\|A0A1B0GT70\|A0A1B0GT70_MOUSE Erlin-2 (Fragment) OS=Mus musculus OX=10090 GN=Erlin2... | tr\|A0A1B0GT70\|A0A1B0GT70_MOUSE Erlin-2 (Fragment) OS=Mus musculus OX=10090 GN=Erlin2... |
| D4AFX7_MOUSE | DnaJ heat shock protein family (Hsp40) member C13 OS=Mus musculus OX=10090 GN=Dnajc13 PE=1 SV=1 |
| sp\|Q5SUF2-2\|LC7L3_MOUSE Isoform 2 of Luc7-like protein 3 OS=Mus musculus OX=10090... | Isoform 2 of Luc7-like protein 3 OS=Mus musculus OX=10090 GN=Luc7l3 |
| sp\|Q8K4Z5\|SF3A1_MOUSESplicing factor 3A subunit 1 OS=Mus musculus OX=10090 GN=Sf3a1 PE=1 SV=1 | sp\|Q8K4Z5\|SF3A1_MOUSE Splicing factor 3A subunit 1 OS=Mus musculus OX=10090 GN=Sf3a1 PE=1 SV=1 |
| sp\|Q9JIX8-4\|ACINU_MOUSE Isoform 4 of Apoptotic chromatin condensation inducer in... | Isoform 4 of Apoptotic chromatin condensation inducer in the nucleus OS=Mus musculus OX=10090 GN=Acin1 |
| tr\|F8VPN4\|F8VPN4_MOUSE Amylo-1,6-glucosidase, 4-alpha-glucanotransferase OS=Mus musculus... | tr\|F8VPN4\|F8VPN4_MOUSE Amylo-1,6-glucosidase, 4-alpha-glucanotransferase OS=Mus musculus... |
| APT_MOUSE | Adenine phosphoribosyltransferase OS=Mus musculus OX=10090 GN=Aprt PE=1 SV=2 |
| tr\|Q14C24\|Q14C24_MOUSE MCG14259, isoform CRA_a OS=Mus musculus OX=10090 GN=U2af1... | tr\|Q14C24\|Q14C24_MOUSE MCG14259, isoform CRA_a OS=Mus musculus OX=10090 GN=U2af1... |
| sp\|P33609\|DPOLA_MOUSEDNA polymerase alpha catalytic subunit OS=Mus musculus OX=10090 GN=Pola1 PE=1 SV=2 | sp\|P33609\|DPOLA_MOUSE DNA polymerase alpha catalytic subunit OS=Mus musculus OX=10090 GN=Pola1 PE=1 SV=2 |
| E9Q1Z0_MOUSE | Keratin 90 OS=Mus musculus OX=10090 GN=Krt90 PE=1 SV=1 |

| | |
|---|---|
| I2BP1_MOUSE | Isoform 2 of Interferon regulatory factor 2-binding protein 1 OS=Mus musculus OX=10090 GN=Irf2bp1 |
| Q5RL55_MOUSE | ATP-binding cassette, sub-family F (GCN20), member 1 OS=Mus musculus OX=10090 GN=Abcf1 PE=2 SV=1 |
| sp\|Q6ZPU9\|KBP_MOUSE KIF1-binding protein OS=Mus musculus OX=10090 GN=Kif1bp PE=1... | KIF1-binding protein OS=Mus musculus OX=10090 GN=Kif1bp PE=1 SV=2 |
| sp\|Q8BH59\|CMC1_MOUSE Calcium-binding mitochondrial carrier protein Aralar1 OS=Mus... | sp\|Q8BH59\|CMC1_MOUSE Calcium-binding mitochondrial carrier protein Aralar1 OS=Mus... |
| B1AYL0_MOUSE | Sodium channel protein OS=Mus musculus OX=10090 GN=Scn9a PE=1 SV=1 |
| tr\|Q61782\|Q61782_MOUSE Type I epidermal keratin mRNA, 3'end (Fragment) OS=Mus musculus... | tr\|Q61782\|Q61782_MOUSE Type I epidermal keratin mRNA, 3'end (Fragment) OS=Mus musculus... |
| sp\|Q8CGC6\|RBM28_MOUSERNA-binding protein 28 OS=Mus musculus OX=10090 GN=Rbm28 PE=1 SV=4 | sp\|Q8CGC6\|RBM28_MOUSE RNA-binding protein 28 OS=Mus musculus OX=10090 GN=Rbm28 PE=1 SV=4 |
| Q91YS7_MOUSE | Dual-specificity mitogen-activated protein kinase kinase 2 OS=Mus musculus OX=10090 GN=Map2k2 PE=1 SV=1 |
| Q0VGN4_MOUSE | Stx16 protein (Fragment) OS=Mus musculus OX=10090 GN=Stx16 PE=2 SV=1 |
| sp\|P62334\|PRS10_MOUSE26S proteasome regulatory subunit 10B OS=Mus musculus OX=10090 GN=Psmc6 PE=1 SV=1 | sp\|P62334\|PRS10_MOUSE 26S proteasome regulatory subunit 10B OS=Mus musculus OX=10090 GN=Psmc6 PE=1 SV=1 |
| tr\|A0A1B0GSA8\|A0A1B0GSA8_MOUSE 60S ribosomal protein L18 (Fragment) OS=Mus musculus... | tr\|A0A1B0GSA8\|A0A1B0GSA8_MOUSE 60S ribosomal protein L18 (Fragment) OS=Mus musculus... |
| B9EKB5_MOUSE | Baz2b protein OS=Mus musculus OX=10090 GN=Baz2b PE=2 SV=1 |
| Q8CIH9_MOUSE | Amidophosphoribosyltransferase OS=Mus musculus OX=10090 GN=Ppat PE=1 SV=1 |
| Q3UJX2_MOUSE | Uncharacterized protein OS=Mus musculus OX=10090 GN=Ctnnbl1 PE=2 SV=1 |
| tr\|E9Q8V6\|E9Q8V6_MOUSE-DECOY DENN/MADD domain-containing 4A OS=Mus musculus OX=10090... | tr\|E9Q8V6\|E9Q8V6_MOUSE-DECOY DENN/MADD domain-containing 4A OS=Mus musculus OX=10090... |

| | |
|---|---|
| sp\|Q91WT9\|CBS_MOUSE Cystathionine beta-synthase OS=Mus musculus OX=10090 GN=Cbs PE=1... | sp\|Q91WT9\|CBS_MOUSE Cystathionine beta-synthase OS=Mus musculus OX=10090 GN=Cbs PE=1... |
| sp\|Q3ULD5\|MCCB_MOUSE Methylcrotonoyl-CoA carboxylase beta chain, mitochondrial OS=Mus... | Methylcrotonoyl-CoA carboxylase beta chain, mitochondrial OS=Mus musculus OX=10090 GN=Mccc2 PE=1 SV=1 |
| STX4_MOUSE | Syntaxin-4 OS=Mus musculus OX=10090 GN=Stx4 PE=1 SV=1 |
| tr\|A0A1D5RMI8\|A0A1D5RMI8_MOUSE Alstrom syndrome protein 1 homolog OS=Mus musculus... | tr\|A0A1D5RMI8\|A0A1D5RMI8_MOUSE Alstrom syndrome protein 1 homolog OS=Mus musculus... |
| E9QNY8_MOUSE | Sacsin OS=Mus musculus OX=10090 GN=Sacs PE=1 SV=1 |
| F8VPZ3_MOUSE | Ubiquitin-specific peptidase 32 OS=Mus musculus OX=10090 GN=Usp32 PE=1 SV=1 |
| sp\|Q99JI4\|PSMD6_MOUSE26S proteasome non-ATPase regulatory subunit 6 OS=Mus musculus OX=10090 GN=Psmd6 PE=1 SV=1 | sp\|Q99JI4\|PSMD6_MOUSE 26S proteasome non-ATPase regulatory subunit 6 OS=Mus musculus OX=10090 GN=Psmd6 PE=1 SV=1 |
| Q6P9Q2_MOUSE | DNA-directed RNA polymerase subunit beta OS=Mus musculus OX=10090 GN=Polr1b PE=2 SV=1 |
| E0CZ42_MOUSE | Fer-1-like 6 (C. elegans) OS=Mus musculus OX=10090 GN=Fer1l6 PE=1 SV=1 |
| E9Q1R5_MOUSE | Voltage-dependent P/Q-type calcium channel subunit alpha OS=Mus musculus OX=10090 GN=Cacna1a PE=1 SV=1 |
| Q6AXF8_MOUSE | Hmgxb3 protein OS=Mus musculus OX=10090 GN=Hmgxb3 PE=2 SV=1 |
| sp\|Q3V300\|KIF22_MOUSEKinesin-like protein KIF22 OS=Mus musculus OX=10090 GN=Kif22 PE=2 SV=2 | sp\|Q3V300\|KIF22_MOUSE Kinesin-like protein KIF22 OS=Mus musculus OX=10090 GN=Kif22 PE=2 SV=2 |
| Q3TGQ4_MOUSE | Uncharacterized protein OS=Mus musculus OX=10090 GN=Secisbp2 PE=2 SV=1 |
| MT2_MOUSE | Metallothionein-2 OS=Mus musculus OX=10090 GN=Mt2 PE=1 SV=2 |
| Q6PDG0_MOUSE | Nup205 protein (Fragment) OS=Mus musculus OX=10090 GN=Nup205 PE=2 SV=1 |
| Q5M9J5_MOUSE | Ribosomal protein L9 OS=Mus musculus OX=10090 GN=Rpl9 PE=2 SV=1 |

| | |
|---|---|
| H3BKE1_MOUSE | Phosphoinositide phospholipase C (Fragment) OS=Mus musculus OX=10090 GN=Plch1 PE=1 SV=8 |
| Q91VT3_MOUSE | ATP-binding cassette, sub-family D (ALD), member 3 OS=Mus musculus OX=10090 GN=Abcd3 PE=2 SV=1 |
| JARD2_MOUSE | Isoform 2 of Protein Jumonji OS=Mus musculus OX=10090 GN=Jarid2 |
| CFA54_MOUSE | Isoform 2 of Cilia- and flagella-associated protein 54 OS=Mus musculus OX=10090 GN=Cfap54 |
| E9PV45_MOUSE | Ubiquitin carboxyl-terminal hydrolase 24 OS=Mus musculus OX=10090 GN=Usp24 PE=1 SV=1 |
| sp\|Q9D4H1\|EXOC2_MOUSEExocyst complex component 2 OS=Mus musculus OX=10090 GN=Exoc2 PE=1 SV=1 | sp\|Q9D4H1\|EXOC2_MOUSE Exocyst complex component 2 OS=Mus musculus OX=10090 GN=Exoc2 PE=1 SV=1 |
| sp\|O54824\|IL16_MOUSE | Pro-interleukin-16 OS=Mus musculus OX=10090 GN=Il16 PE=1 SV=3 |
| A0A1L1SQA8_MOUSE | 40S ribosomal protein S25 OS=Mus musculus OX=10090 GN=Rps25 PE=1 SV=1 |
| A0A1L1SU22_MOUSE | Cysteine and histidine-rich domain-containing protein 1 (Fragment) OS=Mus musculus OX=10090 GN=Chordc1 PE=1 SV=1 |
| Q921V3_MOUSE | Nitrogen fixation gene 1 (S. cerevisiae) OS=Mus musculus OX=10090 GN=Nfs1 PE=2 SV=1 |
| sp\|Q61301-3\|CTNA2_MOUSE | Isoform 3 of Catenin alpha-2 OS=Mus musculus OX=10090 GN=Ctnna2 |
| A0A0R4J0G0_MOUSE | Phosphoenolpyruvate carboxykinase [GTP], mitochondrial OS=Mus musculus OX=10090 GN=Pck2 PE=1 SV=1 |
| Q3T9L0_MOUSE | Uncharacterized protein OS=Mus musculus OX=10090 GN=Ddx39 PE=2 SV=1 |
| tr\|E9Q3N0\|E9Q3N0_MOUSE ATP-dependent DNA helicase OS=Mus musculus OX=10090 GN=Recql... | tr\|E9Q3N0\|E9Q3N0_MOUSE ATP-dependent DNA helicase OS=Mus musculus OX=10090 GN=Recql... |
| tr\|F6Y3V0\|F6Y3V0_MOUSE Receptor-type tyrosine-protein phosphatase F (Fragment) OS=Mus... | tr\|F6Y3V0\|F6Y3V0_MOUSE Receptor-type tyrosine-protein phosphatase F (Fragment) OS=Mus... |

| | |
|---|---|
| tr\|F8VQE9\|F8VQE9_MOUSE Arf-GAP with GTPase, ANK repeat and PH domain-containing protein... | Arf-GAP with GTPase, ANK repeat and PH domain-containing protein 3 OS=Mus musculus OX=10090 GN=Agap3 PE=1 SV=1 |
| tr\|Q3T9Y9\|Q3T9Y9_MOUSE Uncharacterized protein (Fragment) OS=Mus musculus OX=10090... | tr\|Q3T9Y9\|Q3T9Y9_MOUSE Uncharacterized protein (Fragment) OS=Mus musculus OX=10090... |
| AGO2_MOUSE | Protein argonaute-2 OS=Mus musculus OX=10090 GN=Ago2 PE=1 SV=3 |
| tr\|B1AR51\|B1AR51_MOUSE Dynein, axonemal, heavy chain 9 OS=Mus musculus OX=10090 GN=Dnah9... | tr\|B1AR51\|B1AR51_MOUSE Dynein, axonemal, heavy chain 9 OS=Mus musculus OX=10090 GN=Dnah9... |
| tr\|Q3TU36\|Q3TU36_MOUSE RAP1, GTP-GDP dissociation stimulator 1 OS=Mus musculus OX=10090... | RAP1, GTP-GDP dissociation stimulator 1 OS=Mus musculus OX=10090 GN=Rap1gds1 PE=1 SV=1 |
| sp\|E9Q6J5-3\|BD1L1_MOUSE Isoform New of Biorientation of chromosomes in cell division... | Isoform New of Biorientation of chromosomes in cell division protein 1-like 1 OS=Mus musculus OX=10090 GN=Bod1l |
| D3YYT1_MOUSE | Putative oxidoreductase GLYR1 OS=Mus musculus OX=10090 GN=Glyr1 PE=1 SV=1 |
| tr\|A0A1B0GRY3\|A0A1B0GRY3_MOUSE Predicted gene 45861 OS=Mus musculus OX=10090 GN=Gm45861... | tr\|A0A1B0GRY3\|A0A1B0GRY3_MOUSE Predicted gene 45861 OS=Mus musculus OX=10090 GN=Gm45861... |
| tr\|A0A140LIJ4\|A0A140LIJ4_MOUSE-DECOY Dynein, axonemal, heavy chain 14 OS=Mus musculus... | tr\|A0A140LIJ4\|A0A140LIJ4_MOUSE-DECOY Dynein, axonemal, heavy chain 14 OS=Mus musculus... |
| Q571M5_MOUSE | MKIAA4011 protein (Fragment) OS=Mus musculus OX=10090 GN=Itch PE=2 SV=1 |
| J3QN19_MOUSE | DNA primase OS=Mus musculus OX=10090 GN=Prim1 PE=1 SV=1 |
| sp\|Q9CY64\|BIEA_MOUSE Biliverdin reductase A OS=Mus musculus OX=10090 GN=Blvra PE=1... | sp\|Q9CY64\|BIEA_MOUSE Biliverdin reductase A OS=Mus musculus OX=10090 GN=Blvra PE=1... |
| sp\|Q62388\|ATM_MOUSE Serine-protein kinase ATM OS=Mus musculus OX=10090 GN=Atm PE=1... | sp\|Q62388\|ATM_MOUSE Serine-protein kinase ATM OS=Mus musculus OX=10090 GN=Atm PE=1... |
| sp\|Q922H2\|PDK3_MOUSE [Pyruvate dehydrogenase (acetyl-transferring)] kinase isozyme... | sp\|Q922H2\|PDK3_MOUSE [Pyruvate dehydrogenase (acetyl-transferring)] kinase isozyme... |
| Q3UGS1_MOUSE | Uncharacterized protein OS=Mus musculus OX=10090 GN=Nipsnap1 PE=2 SV=1 |

| | |
|---|---|
| tr\|V9GX23\|V9GX23_MOUSE Vacuolar protein sorting 13D (Fragment) OS=Mus musculus OX=10090... | Vacuolar protein sorting 13D (Fragment) OS=Mus musculus OX=10090 GN=Vps13d PE=1 SV=1 |
| sp\|Q9QZQ1-2\|AFAD_MOUSE-DECOY Isoform 1 of Afadin OS=Mus musculus OX=10090 GN=Afdn | sp\|Q9QZQ1-2\|AFAD_MOUSE-DECOY Isoform 1 of Afadin OS=Mus musculus OX=10090 GN=Afdn |
| sp\|P01027\|CO3_MOUSE-DECOY Complement C3 OS=Mus musculus OX=10090 GN=C3 PE=1 SV=3 | sp\|P01027\|CO3_MOUSE-DECOY Complement C3 OS=Mus musculus OX=10090 GN=C3 PE=1 SV=3 |
| sp\|Q9WTI7-2\|MYO1C_MOUSE | Isoform 2 of Unconventional myosin-Ic OS=Mus musculus OX=10090 GN=Myo1c |
| sp\|Q8C1B1-2\|CAMP2_MOUSE Isoform 2 of Calmodulin-regulated spectrin-associated protein... | sp\|Q8C1B1-2\|CAMP2_MOUSE Isoform 2 of Calmodulin-regulated spectrin-associated protein... |
| sp\|Q3TEA8-2\|HP1B3_MOUSE | Isoform 2 of Heterochromatin protein 1-binding protein 3 OS=Mus musculus OX=10090 GN=Hp1bp3 |
| tr\|B9EKC3\|B9EKC3_MOUSE-DECOY Rho GTPase activating protein 5 OS=Mus musculus OX=10090... | tr\|B9EKC3\|B9EKC3_MOUSE-DECOY Rho GTPase activating protein 5 OS=Mus musculus OX=10090... |
| tr\|A0A0G2JG52\|A0A0G2JG52_MOUSE Crystallin beta-gamma domain-containing 1 OS=Mus musculus... | tr\|A0A0G2JG52\|A0A0G2JG52_MOUSE Crystallin beta-gamma domain-containing 1 OS=Mus musculus... |
| sp\|Q11011\|PSA_MOUSE Puromycin-sensitive aminopeptidase OS=Mus musculus OX=10090 GN=Npepps... | Puromycin-sensitive aminopeptidase OS=Mus musculus OX=10090 GN=Npepps PE=1 SV=2 |
| sp\|Q6NXI6-2\|RPRD2_MOUSE Isoform 2 of Regulation of nuclear pre-mRNA domain-containing... | sp\|Q6NXI6-2\|RPRD2_MOUSE Isoform 2 of Regulation of nuclear pre-mRNA domain-containing... |
| tr\|Q8K205\|Q8K205_MOUSE Processing of 1, ribonuclease P/MRP family, (S. cerevisiae)... | Processing of 1, ribonuclease P/MRP family, (S. cerevisiae) OS=Mus musculus OX=10090 GN=Pop1 PE=1 SV=1 |
| sp\|A2CG49-7\|KALRN_MOUSE-DECOY Isoform 7 of Kalirin OS=Mus musculus OX=10090 GN=Kalrn | sp\|A2CG49-7\|KALRN_MOUSE-DECOY Isoform 7 of Kalirin OS=Mus musculus OX=10090 GN=Kalrn |
| sp\|Q9JLC8-3\|SACS_MOUSE-DECOY Isoform 3 of Sacsin OS=Mus musculus OX=10090 GN=Sacs | sp\|Q9JLC8-3\|SACS_MOUSE-DECOY Isoform 3 of Sacsin OS=Mus musculus OX=10090 GN=Sacs |
| tr\|A0A1D5RLL4\|A0A1D5RLL4_MOUSE-DECOY Transformation/transcription domain-associated... | tr\|A0A1D5RLL4\|A0A1D5RLL4_MOUSE-DECOY Transformation/transcription domain-associated... |
| NLE1_MOUSE | Notchless protein homolog 1 OS=Mus musculus OX=10090 GN=Nle1 PE=1 SV=4 |

| | |
|---|---|
| tr\|Q3TKP3\|Q3TKP3_MOUSE Uncharacterized protein OS=Mus musculus OX=10090 GN=Atxn10... | tr\|Q3TKP3\|Q3TKP3_MOUSE Uncharacterized protein OS=Mus musculus OX=10090 GN=Atxn10... |
| D3Z487_MOUSE | Translation initiation factor eIF-2B subunit beta (Fragment) OS=Mus musculus OX=10090 GN=Eif2b2 PE=1 SV=1 |
| tr\|A0A1S6GWH6\|A0A1S6GWH6_MOUSE-DECOY Uncharacterized protein OS=Mus musculus OX=10090... | tr\|A0A1S6GWH6\|A0A1S6GWH6_MOUSE-DECOY Uncharacterized protein OS=Mus musculus OX=10090... |
| A2AK42_MOUSE | Predicted gene 13691 OS=Mus musculus OX=10090 GN=Gm13697 PE=4 SV=1 |
| sp\|Q3UHX0\|NOL8_MOUSE-DECOY Nucleolar protein 8 OS=Mus musculus OX=10090 GN=Nol8 PE=1... | sp\|Q3UHX0\|NOL8_MOUSE-DECOY Nucleolar protein 8 OS=Mus musculus OX=10090 GN=Nol8 PE=1... |
| sp\|Q4VA53-3\|PDS5B_MOUSE-DECOY Isoform 3 of Sister chromatid cohesion protein PDS5... | sp\|Q4VA53-3\|PDS5B_MOUSE-DECOY Isoform 3 of Sister chromatid cohesion protein PDS5... |
| tr\|A0A140LJJ5\|A0A140LJJ5_MOUSE-DECOY A-kinase anchor protein 13 OS=Mus musculus OX=10090... | tr\|A0A140LJJ5\|A0A140LJJ5_MOUSE-DECOY A-kinase anchor protein 13 OS=Mus musculus OX=10090... |
| sp\|P70298\|CUX2_MOUSE-DECOY Homeobox protein cut-like 2 OS=Mus musculus OX=10090 GN=Cux2... | sp\|P70298\|CUX2_MOUSE-DECOY Homeobox protein cut-like 2 OS=Mus musculus OX=10090 GN=Cux2... |
| tr\|A7TU71\|A7TU71_MOUSE-DECOY Protein Shroom2 OS=Mus musculus OX=10090 GN=Shroom2... | tr\|A7TU71\|A7TU71_MOUSE-DECOY Protein Shroom2 OS=Mus musculus OX=10090 GN=Shroom2... |
| tr\|D3Z5K8\|D3Z5K8_MOUSE SH3 and multiple ankyrin repeat domains protein 2 OS=Mus musculus... | SH3 and multiple ankyrin repeat domains protein 2 OS=Mus musculus OX=10090 GN=Shank2 PE=1 SV=1 |
| tr\|E9PWT1\|E9PWT1_MOUSE Transformation/transcription domain-associated protein OS=Mus... | Transformation/transcription domain-associated protein OS=Mus musculus OX=10090 GN=Trrap PE=1 SV=1 |
| tr\|H7BX26\|H7BX26_MOUSE Centrosomal protein of 170 kDa OS=Mus musculus OX=10090 GN=Cep170... | tr\|H7BX26\|H7BX26_MOUSE Centrosomal protein of 170 kDa OS=Mus musculus OX=10090 GN=Cep170... |
| tr\|Q3UYH9\|Q3UYH9_MOUSE Alpha-1,4 glucan phosphorylase OS=Mus musculus OX=10090 GN=Pygb... | Alpha-1,4 glucan phosphorylase OS=Mus musculus OX=10090 GN=Pygb PE=2 SV=1 |
| sp\|Q8QZS1\|HIBCH_MOUSE3-hydroxyisobutyryl-CoA hydrolase, mitochondrial OS=Mus musculus OX=10090 GN=Hibch PE=1 SV=1 | sp\|Q8QZS1\|HIBCH_MOUSE 3-hydroxyisobutyryl-CoA hydrolase, mitochondrial OS=Mus musculus OX=10090 GN=Hibch PE=1 SV=1 |

| | |
|---|---|
| B1AUN3_MOUSE | Eukaryotic translation initiation factor 2B, subunit 3 OS=Mus musculus OX=10090 GN=Eif2b3 PE=1 SV=1 |
| RFC4_MOUSE | Replication factor C subunit 4 OS=Mus musculus OX=10090 GN=Rfc4 PE=1 SV=1 |
| tr\|F8WIE5\|F8WIE5_MOUSE E3 ubiquitin-protein ligase HECTD1 OS=Mus musculus OX=10090... | tr\|F8WIE5\|F8WIE5_MOUSE E3 ubiquitin-protein ligase HECTD1 OS=Mus musculus OX=10090... |
| tr\|A0A0R4J0D3\|A0A0R4J0D3_MOUSE Dolichyl-diphosphooligosaccharide--protein glycosyltransferase... | tr\|A0A0R4J0D3\|A0A0R4J0D3_MOUSE Dolichyl-diphosphooligosaccharide--protein glycosyltransferase... |
| IF2B_MOUSE | Eukaryotic translation initiation factor 2 subunit 2 OS=Mus musculus OX=10090 GN=Eif2s2 PE=1 SV=1 |
| WRIP1_MOUSE | Isoform 2 of ATPase WRNIP1 OS=Mus musculus OX=10090 GN=Wrnip1 |
| K7Q751_MOUSE | Focal ashension kinase 1 OS=Mus musculus OX=10090 GN=Ptk2 PE=2 SV=1 |
| G3UZA0_MOUSE | KRR1 small subunit processome component OS=Mus musculus OX=10090 GN=Krr1 PE=1 SV=1 |
| A2AS98_MOUSE | Nck-associated protein 1 OS=Mus musculus OX=10090 GN=Nckap1 PE=1 SV=1 |
| SUCA_MOUSE | Succinate--CoA ligase [ADP/GDP-forming] subunit alpha, mitochondrial OS=Mus musculus OX=10090 GN=Suclg1 PE=1 SV=4 |
| tr\|Q3U0F8\|Q3U0F8_MOUSE Uncharacterized protein (Fragment) OS=Mus musculus OX=10090... | tr\|Q3U0F8\|Q3U0F8_MOUSE Uncharacterized protein (Fragment) OS=Mus musculus OX=10090... |
| A0A0G2JFB4_MOUSE | Eukaryotic translation initiation factor 4E OS=Mus musculus OX=10090 GN=Eif4e PE=1 SV=1 |
| UCHL5_MOUSE | Isoform 2 of Ubiquitin carboxyl-terminal hydrolase isozyme L5 OS=Mus musculus OX=10090 GN=Uchl5 |
| G3X9F7_MOUSE | DEAD (Asp-Glu-Ala-Asp) box polypeptide 60 OS=Mus musculus OX=10090 GN=Ddx60 PE=1 SV=1 |
| B2RY79_MOUSE | Dock9 protein OS=Mus musculus OX=10090 GN=Dock9 PE=2 SV=1 |

| | |
|---|---|
| Q68FC6_MOUSE | Wrd43 protein (Fragment) OS=Mus musculus OX=10090 GN=Wdr43 PE=2 SV=1 |
| tr\|Q922G9\|Q922G9_MOUSE Copg2 protein (Fragment) OS=Mus musculus OX=10090 GN=Copg2... | tr\|Q922G9\|Q922G9_MOUSE Copg2 protein (Fragment) OS=Mus musculus OX=10090 GN=Copg2... |
| A0A1W2P7C7_MOUSE | Plasma membrane calcium-transporting ATPase 1 (Fragment) OS=Mus musculus OX=10090 GN=Atp2b1 PE=1 SV=1 |
| sp\|P26516\|PSMD7_MOUSE26S proteasome non-ATPase regulatory subunit 7 OS=Mus musculus OX=10090 GN=Psmd7 PE=1 SV=2 | sp\|P26516\|PSMD7_MOUSE 26S proteasome non-ATPase regulatory subunit 7 OS=Mus musculus OX=10090 GN=Psmd7 PE=1 SV=2 |
| Q3UXM3_MOUSE | Uncharacterized protein OS=Mus musculus OX=10090 GN=Maged1 PE=2 SV=1 |
| Q6NZD2_MOUSE | Sorting nexin-1 OS=Mus musculus OX=10090 GN=Snx1 PE=1 SV=1 |
| Q31143_MOUSE | Histocompatibility 2, O region beta locus OS=Mus musculus OX=10090 GN=H2-Ob PE=3 SV=1 |
| sp\|P04117\|FABP4_MOUSEFatty acid-binding protein, adipocyte OS=Mus musculus OX=10090 GN=Fabp4 PE=1 SV=3 | sp\|P04117\|FABP4_MOUSE Fatty acid-binding protein, adipocyte OS=Mus musculus OX=10090 GN=Fabp4 PE=1 SV=3 |
| tr\|A0A2I3BQ03\|A0A2I3BQ03_MOUSE 14-3-3 protein zeta/delta (Fragment) OS=Mus musculus... | tr\|A0A2I3BQ03\|A0A2I3BQ03_MOUSE 14-3-3 protein zeta/delta (Fragment) OS=Mus musculus... |
| Q3TT06_MOUSE | Uncharacterized protein OS=Mus musculus OX=10090 GN=Nup107 PE=2 SV=1 |
| FRY_MOUSE | Protein furry homolog OS=Mus musculus OX=10090 GN=Fry PE=1 SV=1 |
| ARMC3_MOUSE | Isoform 2 of Armadillo repeat-containing protein 3 OS=Mus musculus OX=10090 GN=Armc3 |
| Q149V4_MOUSE | Histone H2A OS=Mus musculus OX=10090 GN=Hist2h2ac PE=2 SV=1 |
| A0A0U1RNK9_MOUSE | Electron transfer flavoprotein subunit beta (Fragment) OS=Mus musculus OX=10090 GN=Etfb PE=1 SV=2 |
| A0A1S6GWJ0_MOUSE | Oxysterol-binding protein OS=Mus musculus OX=10090 GN=Osbpl9 PE=2 SV=1 |

| | |
|---|---|
| A2ALR8_MOUSE | Ubiquitin carboxyl-terminal hydrolase 48 (Fragment) OS=Mus musculus OX=10090 GN=Usp48 PE=1 SV=1 |
| NU160_MOUSE | Isoform 2 of Nuclear pore complex protein Nup160 OS=Mus musculus OX=10090 GN=Nup160 |
| Q7TNL5_MOUSE | Serine/threonine-protein phosphatase 2A 56 kDa regulatory subunit OS=Mus musculus OX=10090 GN=Ppp2r5d PE=2 SV=1 |
| TSN_MOUSE | Translin OS=Mus musculus OX=10090 GN=Tsn PE=1 SV=1 |
| sp\|E9PVA8\|GCN1_MOUSE-DECOY eIF-2-alpha kinase activator GCN1 OS=Mus musculus OX=10090... | sp\|E9PVA8\|GCN1_MOUSE-DECOY eIF-2-alpha kinase activator GCN1 OS=Mus musculus OX=10090... |
| sp\|P14152\|MDHC_MOUSE Malate dehydrogenase, cytoplasmic OS=Mus musculus OX=10090 GN=Mdh1... | sp\|P14152\|MDHC_MOUSE Malate dehydrogenase, cytoplasmic OS=Mus musculus OX=10090 GN=Mdh1... |
| sp\|Q8C196\|CPSM_MOUSE Carbamoyl-phosphate synthase [ammonia], mitochondrial OS=Mus... | sp\|Q8C196\|CPSM_MOUSE Carbamoyl-phosphate synthase [ammonia], mitochondrial OS=Mus... |
| sp\|Q8K0C9\|GMDS_MOUSE GDP-mannose 4,6 dehydratase OS=Mus musculus OX=10090 GN=Gmds... | GDP-mannose 4,6 dehydratase OS=Mus musculus OX=10090 GN=Gmds PE=1 SV=1 |
| sp\|Q9JJV2-2\|PROF2_MOUSE | Isoform 2 of Profilin-2 OS=Mus musculus OX=10090 GN=Pfn2 |
| tr\|A0A0A0MQM0\|A0A0A0MQM0_MOUSE Eukaryotic translation initiation factor 5A (Fragment)... | tr\|A0A0A0MQM0\|A0A0A0MQM0_MOUSE Eukaryotic translation initiation factor 5A (Fragment)... |
| tr\|A0A0R4J1L9\|A0A0R4J1L9_MOUSE Microtubule-associated tumor suppressor 1 homolog... | tr\|A0A0R4J1L9\|A0A0R4J1L9_MOUSE Microtubule-associated tumor suppressor 1 homolog... |
| tr\|A2ARP8\|A2ARP8_MOUSE Microtubule-associated protein 1A OS=Mus musculus OX=10090... | Microtubule-associated protein 1A OS=Mus musculus OX=10090 GN=Map1a PE=1 SV=1 |
| tr\|B2RQQ5\|B2RQQ5_MOUSE Microtubule-associated protein 1B OS=Mus musculus OX=10090... | tr\|B2RQQ5\|B2RQQ5_MOUSE Microtubule-associated protein 1B OS=Mus musculus OX=10090... |
| tr\|D6RHT6\|D6RHT6_MOUSE E3 ubiquitin-protein ligase TTC3 (Fragment) OS=Mus musculus... | E3 ubiquitin-protein ligase TTC3 (Fragment) OS=Mus musculus OX=10090 GN=Ttc3 PE=1 SV=1 |
| tr\|E9Q0N0\|E9Q0N0_MOUSE-DECOY Intersectin-1 OS=Mus musculus OX=10090 GN=Itsn1 PE=1... | tr\|E9Q0N0\|E9Q0N0_MOUSE-DECOY Intersectin-1 OS=Mus musculus OX=10090 GN=Itsn1 PE=1... |

| | |
|---|---|
| sp\|Q5SVQ0-4\|KAT7_MOUSE | Isoform 4 of Histone acetyltransferase KAT7 OS=Mus musculus OX=10090 GN=Kat7 |
| sp\|Q9D2G2\|ODO2_MOUSE Dihydrolipoyllysine-residue succinyltransferase component of... | sp\|Q9D2G2\|ODO2_MOUSE Dihydrolipoyllysine-residue succinyltransferase component of... |
| sp\|Q9ERD7\|TBB3_MOUSE Tubulin beta-3 chain OS=Mus musculus OX=10090 GN=Tubb3 PE=1... | Tubulin beta-3 chain OS=Mus musculus OX=10090 GN=Tubb3 PE=1 SV=1 |
| sp\|Q9CZX0-2\|ELP3_MOUSE Isoform 2 of Elongator complex protein 3 OS=Mus musculus OX=10090... | Isoform 2 of Elongator complex protein 3 OS=Mus musculus OX=10090 GN=Elp3 |
| I3PQW3_MOUSE | LARP7 OS=Mus musculus OX=10090 GN=Larp7 PE=4 SV=1 |
| W6PPR4_MOUSE | 480-kDa ankyrinG OS=Mus musculus OX=10090 GN=Ank3 PE=4 SV=1 |
| sp\|O88492-2\|PLIN4_MOUSE-DECOY Isoform 2 of Perilipin-4 OS=Mus musculus OX=10090 GN=Plin4 | sp\|O88492-2\|PLIN4_MOUSE-DECOY Isoform 2 of Perilipin-4 OS=Mus musculus OX=10090 GN=Plin4 |
| sp\|Q8BKX6\|SMG1_MOUSE Serine/threonine-protein kinase SMG1 OS=Mus musculus OX=10090... | Serine/threonine-protein kinase SMG1 OS=Mus musculus OX=10090 GN=Smg1 PE=1 SV=3 |
| sp\|Q99LH1\|NOG2_MOUSE Nucleolar GTP-binding protein 2 OS=Mus musculus OX=10090 GN=Gnl2... | sp\|Q99LH1\|NOG2_MOUSE Nucleolar GTP-binding protein 2 OS=Mus musculus OX=10090 GN=Gnl2... |
| tr\|A0A0A0MQ73\|A0A0A0MQ73_MOUSE-DECOY Histone-lysine N-methyltransferase 2D OS=Mus... | tr\|A0A0A0MQ73\|A0A0A0MQ73_MOUSE-DECOY Histone-lysine N-methyltransferase 2D OS=Mus... |
| tr\|H3BLK8\|H3BLK8_MOUSE-DECOY Zinc finger homeobox protein 4 OS=Mus musculus OX=10090... | tr\|H3BLK8\|H3BLK8_MOUSE-DECOY Zinc finger homeobox protein 4 OS=Mus musculus OX=10090... |
| tr\|Q8BVX3\|Q8BVX3_MOUSE Uncharacterized protein (Fragment) OS=Mus musculus OX=10090... | tr\|Q8BVX3\|Q8BVX3_MOUSE Uncharacterized protein (Fragment) OS=Mus musculus OX=10090... |
| sp\|Q80Y81\|RNZ2_MOUSE | Zinc phosphodiesterase ELAC protein 2 OS=Mus musculus OX=10090 GN=Elac2 PE=1 SV=1 |
| A0A068BFS4_MOUSE | Ring finger protein 1 OS=Mus musculus OX=10090 GN=Ring1 PE=2 SV=1 |
| O70349_MOUSE | MCG15924, isoform CRA_a OS=Mus musculus OX=10090 GN=SKI PE=4 SV=1 |

| | |
|---|---|
| Q8BSH9_MOUSE | Nucleosome assembly protein 1-like 1 OS=Mus musculus OX=10090 GN=Nap1l1 PE=1 SV=1 |
| Q3TST1_MOUSE | Gremlin OS=Mus musculus OX=10090 GN=Grem2 PE=2 SV=1 |
| NIPBL_MOUSE | Isoform 2 of Nipped-B-like protein OS=Mus musculus OX=10090 GN=Nipbl |
| tr\|G3UW82\|G3UW82_MOUSE MCG140437, isoform CRA_d OS=Mus musculus OX=10090 GN=Myh2... | MCG140437, isoform CRA_d OS=Mus musculus OX=10090 GN=Myh2 PE=1 SV=1 |
| sp\|Q45VK7-2\|DYHC2_MOUSE-DECOY Isoform 2 of Cytoplasmic dynein 2 heavy chain 1 OS=Mus... | sp\|Q45VK7-2\|DYHC2_MOUSE-DECOY Isoform 2 of Cytoplasmic dynein 2 heavy chain 1 OS=Mus... |
| sp\|Q8VBZ3\|CLPT1_MOUSECleft lip and palate transmembrane protein 1 homolog OS=Mus musculus OX=10090 GN=Clptm1 PE=1 SV=1 | sp\|Q8VBZ3\|CLPT1_MOUSE Cleft lip and palate transmembrane protein 1 homolog OS=Mus musculus OX=10090 GN=Clptm1 PE=1 SV=1 |
| IMA5_MOUSE | Importin subunit alpha-5 OS=Mus musculus OX=10090 GN=Kpna1 PE=1 SV=2 |
| sp\|Q9QY06-2\|MYO9B_MOUSE-DECOY Isoform 2 of Unconventional myosin-IXb OS=Mus musculus... | sp\|Q9QY06-2\|MYO9B_MOUSE-DECOY Isoform 2 of Unconventional myosin-IXb OS=Mus musculus... |
| Q3THX4_MOUSE | Uncharacterized protein OS=Mus musculus OX=10090 GN=Vps45 PE=2 SV=1 |
| E9Q8K8_MOUSE | Zinc finger CCCH domain-containing protein 4 OS=Mus musculus OX=10090 GN=Zc3h4 PE=1 SV=1 |
| SORL_MOUSE | Sortilin-related receptor OS=Mus musculus OX=10090 GN=Sorl1 PE=1 SV=3 |
| sp\|Q9Z0P5-2\|TWF2_MOUSE | Isoform 2 of Twinfilin-2 OS=Mus musculus OX=10090 GN=Twf2 |
| sp\|Q62315-2\|JARD2_MOUSE-DECOY Isoform 2 of Protein Jumonji OS=Mus musculus OX=10090... | sp\|Q62315-2\|JARD2_MOUSE-DECOY Isoform 2 of Protein Jumonji OS=Mus musculus OX=10090... |
| Q66X24_MOUSE | NALP-beta (Fragment) OS=Mus musculus OX=10090 GN=Nlrp4d PE=2 SV=1 |
| Q3TCH2_MOUSE | Ubiquitin carboxyl-terminal hydrolase (Fragment) OS=Mus musculus OX=10090 GN=Uchl1 PE=2 SV=1 |
| tr\|A0A1W2P7W3\|A0A1W2P7W3_MOUSE-DECOY Protein tyrosine phosphatase, receptor type,... | tr\|A0A1W2P7W3\|A0A1W2P7W3_MOUSE-DECOY Protein tyrosine phosphatase, receptor type,... |

| | |
|---|---|
| sp\|Q64028-2\|PHC1_MOUSE-DECOY Isoform 2 of Polyhomeotic-like protein 1 OS=Mus musculus... | sp\|Q64028-2\|PHC1_MOUSE-DECOY Isoform 2 of Polyhomeotic-like protein 1 OS=Mus musculus... |
| A0A1W2P7A1_MOUSE | 40S ribosomal protein S12 OS=Mus musculus OX=10090 GN=Rps12 PE=1 SV=1 |
| A0A1B0GT06_MOUSE | Predicted gene, 29735 OS=Mus musculus OX=10090 GN=Gm29735 PE=4 SV=1 |
| sp\|Q6PE01\|SNR40_MOUSEU5 small nuclear ribonucleoprotein 40 kDa protein OS=Mus musculus OX=10090 GN=Snrnp40 PE=1 SV=1 | sp\|Q6PE01\|SNR40_MOUSE U5 small nuclear ribonucleoprotein 40 kDa protein OS=Mus musculus OX=10090 GN=Snrnp40 PE=1 SV=1 |
| SRSF3_MOUSE | Isoform Short of Serine/arginine-rich splicing factor 3 OS=Mus musculus OX=10090 GN=Srsf3 |
| PGM2_MOUSE | Phosphoglucomutase-2 OS=Mus musculus OX=10090 GN=Pgm2 PE=1 SV=1 |
| sp\|Q8BKX6\|SMG1_MOUSE-DECOY Serine/threonine-protein kinase SMG1 OS=Mus musculus OX=10090... | sp\|Q8BKX6\|SMG1_MOUSE-DECOY Serine/threonine-protein kinase SMG1 OS=Mus musculus OX=10090... |
| ARAF_MOUSE | Serine/threonine-protein kinase A-Raf OS=Mus musculus OX=10090 GN=Araf PE=1 SV=2 |
| Q3UG68_MOUSE | Dolichyl-diphosphooligosaccharide--protein glycosyltransferase 48 kDa subunit OS=Mus musculus OX=10090 GN=Ddost PE=2 SV=1 |
| Q3TX38_MOUSE | Uncharacterized protein OS=Mus musculus OX=10090 GN=Vdac3 PE=1 SV=1 |
| tr\|Q3TIV2\|Q3TIV2_MOUSE Uncharacterized protein OS=Mus musculus OX=10090 GN=Zc3h11a... | Uncharacterized protein OS=Mus musculus OX=10090 GN=Zc3h11a PE=2 SV=1 |
| Q3UZR8_MOUSE | Eukaryotic translation initiation factor 2B, subunit 1 (Alpha) OS=Mus musculus OX=10090 GN=Eif2b1 PE=1 SV=1 |
| B2RY11_MOUSE | Abca12 protein OS=Mus musculus OX=10090 GN=Abca12 PE=2 SV=1 |
| G3X9F2_MOUSE | Histone demethylase UTY OS=Mus musculus OX=10090 GN=Uty PE=1 SV=1 |
| A0A1B0GSA0_MOUSE | Predicted gene 45618 OS=Mus musculus OX=10090 GN=Gm45618 PE=4 SV=1 |

| | |
|---|---|
| A0A0R4J150_MOUSE | Oxysterol-binding protein OS=Mus musculus OX=10090 GN=Osbpl8 PE=1 SV=1 |
| PLSI_MOUSE | Plastin-1 OS=Mus musculus OX=10090 GN=Pls1 PE=1 SV=1 |
| Q6A0C9_MOUSE | MKIAA0109 protein (Fragment) OS=Mus musculus OX=10090 GN=Ap2m1 PE=2 SV=1 |
| sp\|A2AKX3-2\|SETX_MOUSE | Isoform 2 of Probable helicase senataxin OS=Mus musculus OX=10090 GN=Setx |
| sp\|Q641K1-2\|CBPC1_MOUSE | Isoform 2 of Cytosolic carboxypeptidase 1 OS=Mus musculus OX=10090 GN=Agtpbp1 |
| MUG1_MOUSE | Murinoglobulin-1 OS=Mus musculus OX=10090 GN=Mug1 PE=1 SV=3 |
| E9Q3I4_MOUSE | Intersectin-1 OS=Mus musculus OX=10090 GN=Itsn1 PE=1 SV=1 |
| A0A0R4J198_MOUSE | Succinyl-CoA:3-ketoacid-coenzyme A transferase OS=Mus musculus OX=10090 GN=Oxct2b PE=3 SV=1 |
| tr\|Q3TI71\|Q3TI71_MOUSE Uncharacterized protein OS=Mus musculus OX=10090 GN=Prpf8... | tr\|Q3TI71\|Q3TI71_MOUSE Uncharacterized protein OS=Mus musculus OX=10090 GN=Prpf8... |
| sp\|Q3B7Z2\|OSBP1_MOUSEOxysterol-binding protein 1 OS=Mus musculus OX=10090 GN=Osbp PE=1 SV=3 | sp\|Q3B7Z2\|OSBP1_MOUSE Oxysterol-binding protein 1 OS=Mus musculus OX=10090 GN=Osbp PE=1 SV=3 |
| sp\|Q3UDE2\|TTL12_MOUSETubulin--tyrosine ligase-like protein 12 OS=Mus musculus OX=10090 GN=Ttll12 PE=1 SV=1 | sp\|Q3UDE2\|TTL12_MOUSE Tubulin--tyrosine ligase-like protein 12 OS=Mus musculus OX=10090 GN=Ttll12 PE=1 SV=1 |
| Q8C7E6_MOUSE | Uncharacterized protein OS=Mus musculus OX=10090 PE=2 SV=1 |
| B1AR28_MOUSE | Very long-chain-specific acyl-CoA dehydrogenase, mitochondrial OS=Mus musculus OX=10090 GN=Acadvl PE=1 SV=1 |
| CO3_MOUSE | Complement C3 OS=Mus musculus OX=10090 GN=C3 PE=1 SV=3 |
| Q3TNP4_MOUSE | Uncharacterized protein OS=Mus musculus OX=10090 GN=Fam208b PE=2 SV=1 |
| GRM3_MOUSE | Metabotropic glutamate receptor 3 OS=Mus musculus OX=10090 GN=Grm3 PE=1 SV=1 |

| | |
|---|---|
| D3Z5G8_MOUSE | Disco-interacting protein 2 homolog B OS=Mus musculus OX=10090 GN=Dip2b PE=1 SV=1 |
| E9Q7H6_MOUSE | Threonine--tRNA ligase, mitochondrial OS=Mus musculus OX=10090 GN=Tars2 PE=1 SV=1 |
| R4H4Y7_MOUSE | SCY1-like 1 splice variant 13 OS=Mus musculus OX=10090 GN=Scyl1 PE=2 SV=1 |
| B7ZMT5_MOUSE | Mmrn1 protein OS=Mus musculus OX=10090 GN=Mmrn1 PE=2 SV=1 |
| sp\|Q8K411-2\|PREP_MOUSE | Isoform 2 of Presequence protease, mitochondrial OS=Mus musculus OX=10090 GN=Pitrm1 |
| EDC3_MOUSE | Enhancer of mRNA-decapping protein 3 OS=Mus musculus OX=10090 GN=Edc3 PE=1 SV=1 |
| sp\|Q9Z2I0\|LETM1_MOUSEMitochondrial proton/calcium exchanger protein OS=Mus musculus OX=10090 GN=Letm1 PE=1 SV=1 | sp\|Q9Z2I0\|LETM1_MOUSE Mitochondrial proton/calcium exchanger protein OS=Mus musculus OX=10090 GN=Letm1 PE=1 SV=1 |
| PSA4_MOUSE | Proteasome subunit alpha type-4 OS=Mus musculus OX=10090 GN=Psma4 PE=1 SV=1 |
| Q52L97_MOUSE | Importin subunit alpha OS=Mus musculus OX=10090 GN=Kpna2 PE=1 SV=1 |
| G3X9H5_MOUSE | Huntingtin OS=Mus musculus OX=10090 GN=Htt PE=1 SV=1 |
| Q3UD86_MOUSE | Uncharacterized protein OS=Mus musculus OX=10090 GN=Mov10 PE=2 SV=1 |
| HELLS_MOUSE | Isoform 2 of Lymphocyte-specific helicase OS=Mus musculus OX=10090 GN=Hells |
| A0A0J9YU62_MOUSE | C-terminal-binding protein 1 OS=Mus musculus OX=10090 GN=Ctbp1 PE=1 SV=1 |
| Q3TF25_MOUSE | Uncharacterized protein OS=Mus musculus OX=10090 GN=Atp5o PE=2 SV=1 |
| sp\|Q9JK48-2\|SHLB1_MOUSE | Isoform 2 of Endophilin-B1 OS=Mus musculus OX=10090 GN=Sh3glb1 |
| tr\|A2AAN2\|A2AAN2_MOUSE-DECOY Signal recognition particle subunit SRP68 OS=Mus musculus... | tr\|A2AAN2\|A2AAN2_MOUSE-DECOY Signal recognition particle subunit SRP68 OS=Mus musculus... |

| | |
|---|---|
| tr\|A8IP69\|A8IP69_MOUSE 14-3-3 protein gamma subtype OS=Mus musculus OX=10090 GN=Ywhag... | tr\|A8IP69\|A8IP69_MOUSE 14-3-3 protein gamma subtype OS=Mus musculus OX=10090 GN=Ywhag... |
| tr\|B2RWU2\|B2RWU2_MOUSE-DECOY Immunodeficiency virus type I enhancer binding protein... | tr\|B2RWU2\|B2RWU2_MOUSE-DECOY Immunodeficiency virus type I enhancer binding protein... |
| tr\|Q3TXN6\|Q3TXN6_MOUSE Uncharacterized protein (Fragment) OS=Mus musculus OX=10090... | Uncharacterized protein (Fragment) OS=Mus musculus OX=10090 GN=Cstf3 PE=2 SV=1 |
| tr\|Q3U9P7\|Q3U9P7_MOUSE Succinyl-CoA:3-ketoacid-coenzyme A transferase OS=Mus musculus... | Succinyl-CoA:3-ketoacid-coenzyme A transferase OS=Mus musculus OX=10090 GN=Oxct1 PE=2 SV=1 |
| tr\|A0A1B0GS20\|A0A1B0GS20_MOUSE-DECOY Predicted gene 45861 OS=Mus musculus OX=10090... | tr\|A0A1B0GS20\|A0A1B0GS20_MOUSE-DECOY Predicted gene 45861 OS=Mus musculus OX=10090... |
| tr\|Q3TVV7\|Q3TVV7_MOUSE Uncharacterized protein OS=Mus musculus OX=10090 GN=Umps PE=2... | Uncharacterized protein OS=Mus musculus OX=10090 GN=Umps PE=2 SV=1 |
| sp\|P35922-2\|FMR1_MOUSE-DECOY Isoform 2 of Synaptic functional regulator FMR1 OS=Mus... | sp\|P35922-2\|FMR1_MOUSE-DECOY Isoform 2 of Synaptic functional regulator FMR1 OS=Mus... |
| tr\|A0A087WQ94\|A0A087WQ94_MOUSE-DECOY Tensin 1 (Fragment) OS=Mus musculus OX=10090... | tr\|A0A087WQ94\|A0A087WQ94_MOUSE-DECOY Tensin 1 (Fragment) OS=Mus musculus OX=10090... |
| tr\|E9Q1M1\|E9Q1M1_MOUSE-DECOY PDZ domain-containing 2 OS=Mus musculus OX=10090 GN=Pdzd2... | tr\|E9Q1M1\|E9Q1M1_MOUSE-DECOY PDZ domain-containing 2 OS=Mus musculus OX=10090 GN=Pdzd2... |
| tr\|Q9CPN9\|Q9CPN9_MOUSE RIKEN cDNA 2210010C04 gene OS=Mus musculus OX=10090 GN=2210010C04Rik... | tr\|Q9CPN9\|Q9CPN9_MOUSE RIKEN cDNA 2210010C04 gene OS=Mus musculus OX=10090 GN=2210010C04Rik... |
| sp\|Q8CI03-2\|FWCH1_MOUSE-DECOY Isoform 2 of FLYWCH-type zinc finger-containing protein... | sp\|Q8CI03-2\|FWCH1_MOUSE-DECOY Isoform 2 of FLYWCH-type zinc finger-containing protein... |
| sp\|Q9QXS1-6\|PLEC_MOUSE-DECOY Isoform PLEC-0,1C of Plectin OS=Mus musculus OX=10090... | sp\|Q9QXS1-6\|PLEC_MOUSE-DECOY Isoform PLEC-0,1C of Plectin OS=Mus musculus OX=10090... |
| sp\|O88196-2\|TTC3_MOUSE-DECOY Isoform 2 of E3 ubiquitin-protein ligase TTC3 OS=Mus... | sp\|O88196-2\|TTC3_MOUSE-DECOY Isoform 2 of E3 ubiquitin-protein ligase TTC3 OS=Mus... |
| sp\|Q8BI29-2\|SARG_MOUSE Isoform 2 of Specifically androgen-regulated gene protein... | sp\|Q8BI29-2\|SARG_MOUSE Isoform 2 of Specifically androgen-regulated gene protein... |

| | |
|---|---|
| tr\|D3Z794\|D3Z794_MOUSE Small ubiquitin-related modifier 2 OS=Mus musculus OX=10090... | tr\|D3Z794\|D3Z794_MOUSE Small ubiquitin-related modifier 2 OS=Mus musculus OX=10090... |
| tr\|D3YYK9\|D3YYK9_MOUSE-DECOY Brefeldin A-inhibited guanine nucleotide-exchange protein... | tr\|D3YYK9\|D3YYK9_MOUSE-DECOY Brefeldin A-inhibited guanine nucleotide-exchange protein... |
| tr\|A0A158SIT8\|A0A158SIT8_MOUSE Proline, glutamic acid and leucine rich protein 1... | tr\|A0A158SIT8\|A0A158SIT8_MOUSE Proline, glutamic acid and leucine rich protein 1... |
| tr\|B2RUJ4\|B2RUJ4_MOUSE-DECOY Atad5 protein OS=Mus musculus OX=10090 GN=Atad5 PE=2... | tr\|B2RUJ4\|B2RUJ4_MOUSE-DECOY Atad5 protein OS=Mus musculus OX=10090 GN=Atad5 PE=2... |
| tr\|F8VPK0\|F8VPK0_MOUSE Tetratricopeptide repeat domain 37 OS=Mus musculus OX=10090... | tr\|F8VPK0\|F8VPK0_MOUSE Tetratricopeptide repeat domain 37 OS=Mus musculus OX=10090... |
| sp\|P53349\|M3K1_MOUSE-DECOY Mitogen-activated protein kinase kinase kinase 1 OS=Mus... | sp\|P53349\|M3K1_MOUSE-DECOY Mitogen-activated protein kinase kinase kinase 1 OS=Mus... |
| Q8VED4_MOUSE | Serine/threonine-protein phosphatase OS=Mus musculus OX=10090 GN=Ppp2cb PE=2 SV=1 |
| sp\|Q05860-3\|FMN1_MOUSE-DECOY Isoform 3 of Formin-1 OS=Mus musculus OX=10090 GN=Fmn1 | sp\|Q05860-3\|FMN1_MOUSE-DECOY Isoform 3 of Formin-1 OS=Mus musculus OX=10090 GN=Fmn1 |
| tr\|B2RX10\|B2RX10_MOUSE-DECOY Fanconi anemia, complementation group M OS=Mus musculus... | tr\|B2RX10\|B2RX10_MOUSE-DECOY Fanconi anemia, complementation group M OS=Mus musculus... |
| sp\|Q69ZA1-2\|CDK13_MOUSE Isoform 2 of Cyclin-dependent kinase 13 OS=Mus musculus OX=10090... | Isoform 2 of Cyclin-dependent kinase 13 OS=Mus musculus OX=10090 GN=Cdk13 |
| tr\|A0A087WSS1\|A0A087WSS1_MOUSE RIKEN cDNA 2310035C23 gene OS=Mus musculus OX=10090... | tr\|A0A087WSS1\|A0A087WSS1_MOUSE RIKEN cDNA 2310035C23 gene OS=Mus musculus OX=10090... |
| tr\|Q3TEK2\|Q3TEK2_MOUSE-DECOY Uncharacterized protein OS=Mus musculus OX=10090 GN=Hspa8... | tr\|Q3TEK2\|Q3TEK2_MOUSE-DECOY Uncharacterized protein OS=Mus musculus OX=10090 GN=Hspa8... |
| sp\|O88491\|NSD1_MOUSE-DECOY Histone-lysine N-methyltransferase, H3 lysine-36 and H4... | sp\|O88491\|NSD1_MOUSE-DECOY Histone-lysine N-methyltransferase, H3 lysine-36 and H4... |
| sp\|A0JP43-3\|EFCB5_MOUSE-DECOY Isoform 3 of EF-hand calcium-binding domain-containing... | sp\|A0JP43-3\|EFCB5_MOUSE-DECOY Isoform 3 of EF-hand calcium-binding domain-containing... |

| | |
|---|---|
| sp\|Q9DB77\|QCR2_MOUSE Cytochrome b-c1 complex subunit 2, mitochondrial OS=Mus musculus... | sp\|Q9DB77\|QCR2_MOUSE Cytochrome b-c1 complex subunit 2, mitochondrial OS=Mus musculus... |
| sp\|Q9D0T1\|NH2L1_MOUSENHP2-like protein 1 OS=Mus musculus OX=10090 GN=Snu13 PE=1 SV=4 | sp\|Q9D0T1\|NH2L1_MOUSE NHP2-like protein 1 OS=Mus musculus OX=10090 GN=Snu13 PE=1 SV=4 |
| tr\|Q3UG63\|Q3UG63_MOUSE Uncharacterized protein OS=Mus musculus OX=10090 GN=Ascc2... | tr\|Q3UG63\|Q3UG63_MOUSE Uncharacterized protein OS=Mus musculus OX=10090 GN=Ascc2... |
| tr\|Q545N7\|Q545N7_MOUSE Creatine kinase, mitochondrial 1, ubiquitous, isoform CRA_a... | tr\|Q545N7\|Q545N7_MOUSE Creatine kinase, mitochondrial 1, ubiquitous, isoform CRA_a... |
| sp\|Q05512-2\|MARK2_MOUSE-DECOY Isoform 2 of Serine/threonine-protein kinase MARK2... | sp\|Q05512-2\|MARK2_MOUSE-DECOY Isoform 2 of Serine/threonine-protein kinase MARK2... |
| sp\|Q99N50-2\|SYTL2_MOUSE Isoform 2 of Synaptotagmin-like protein 2 OS=Mus musculus... | sp\|Q99N50-2\|SYTL2_MOUSE Isoform 2 of Synaptotagmin-like protein 2 OS=Mus musculus... |
| tr\|Q3U0X6\|Q3U0X6_MOUSE-DECOY Zinc finger protein 217 OS=Mus musculus OX=10090 GN=Zfp217... | tr\|Q3U0X6\|Q3U0X6_MOUSE-DECOY Zinc finger protein 217 OS=Mus musculus OX=10090 GN=Zfp217... |
| A2AKU9_MOUSE | ATP synthase subunit gamma OS=Mus musculus OX=10090 GN=Atp5c1 PE=1 SV=1 |
| Q8K366_MOUSE | Exosc10 protein OS=Mus musculus OX=10090 GN=Exosc10 PE=1 SV=1 |
| ENOB_MOUSE | Beta-enolase OS=Mus musculus OX=10090 GN=Eno3 PE=1 SV=3 |
| sp\|Q9CXF7-2\|CHD1L_MOUSE Isoform 2 of Chromodomain-helicase-DNA-binding protein 1-like... | Isoform 2 of Chromodomain-helicase-DNA-binding protein 1-like OS=Mus musculus OX=10090 GN=Chd1l |
| sp\|Q99K01-2\|PDXD1_MOUSE Isoform 2 of Pyridoxal-dependent decarboxylase domain-containing... | sp\|Q99K01-2\|PDXD1_MOUSE Isoform 2 of Pyridoxal-dependent decarboxylase domain-containing... |
| tr\|Q3TTI6\|Q3TTI6_MOUSE Uncharacterized protein (Fragment) OS=Mus musculus OX=10090... | tr\|Q3TTI6\|Q3TTI6_MOUSE Uncharacterized protein (Fragment) OS=Mus musculus OX=10090... |
| Q3U2B4_MOUSE | Uncharacterized protein OS=Mus musculus OX=10090 GN=Xrcc1 PE=2 SV=1 |
| E9Q4D0_MOUSE | Proprotein convertase subtilisin/kexin type 6 OS=Mus musculus OX=10090 GN=Pcsk6 PE=1 SV=2 |

| | |
|---|---|
| sp\|Q3TZZ7\|ESYT2_MOUSEExtended synaptotagmin-2 OS=Mus musculus OX=10090 GN=Esyt2 PE=1 SV=1 | sp\|Q3TZZ7\|ESYT2_MOUSE Extended synaptotagmin-2 OS=Mus musculus OX=10090 GN=Esyt2 PE=1 SV=1 |
| sp\|Q61037-2\|TSC2_MOUSE | Isoform A of Tuberin OS=Mus musculus OX=10090 GN=Tsc2 |
| tr\|A3KGJ7\|A3KGJ7_MOUSE-DECOY Centrosome-associated protein CEP250 OS=Mus musculus... | tr\|A3KGJ7\|A3KGJ7_MOUSE-DECOY Centrosome-associated protein CEP250 OS=Mus musculus... |
| A2AFK7_MOUSE | Eukaryotic initiation factor 4A-III (Fragment) OS=Mus musculus OX=10090 GN=Eif4a3 PE=1 SV=1 |
| E0CXV8_MOUSE | Polyhomeotic-like protein 1 OS=Mus musculus OX=10090 GN=Phc1 PE=1 SV=1 |
| Q3UDM0_MOUSE | MOB kinase activator 1B OS=Mus musculus OX=10090 GN=Mob1b PE=1 SV=1 |
| sp\|P30416\|FKBP4_MOUSEPeptidyl-prolyl cis-trans isomerase FKBP4 OS=Mus musculus OX=10090 GN=Fkbp4 PE=1 SV=5 | sp\|P30416\|FKBP4_MOUSE Peptidyl-prolyl cis-trans isomerase FKBP4 OS=Mus musculus OX=10090 GN=Fkbp4 PE=1 SV=5 |
| A2A4N7_MOUSE | Next to BRCA1 gene 1 protein OS=Mus musculus OX=10090 GN=Nbr1 PE=1 SV=1 |
| tr\|B1AUX2\|B1AUX2_MOUSE-DECOY Host cell factor 1 OS=Mus musculus OX=10090 GN=Hcfc1... | tr\|B1AUX2\|B1AUX2_MOUSE-DECOY Host cell factor 1 OS=Mus musculus OX=10090 GN=Hcfc1... |
| tr\|F8WIE5\|F8WIE5_MOUSE-DECOY E3 ubiquitin-protein ligase HECTD1 OS=Mus musculus OX=10090... | tr\|F8WIE5\|F8WIE5_MOUSE-DECOY E3 ubiquitin-protein ligase HECTD1 OS=Mus musculus OX=10090... |
| E9Q805_MOUSE | FERM, ARHGEF and pleckstrin domain-containing protein 1 (Fragment) OS=Mus musculus OX=10090 GN=Farp1 PE=1 SV=1 |
| sp\|Q8CH77-2\|NAV1_MOUSE | Isoform 2 of Neuron navigator 1 OS=Mus musculus OX=10090 GN=Nav1 |
| tr\|E9QPI5\|E9QPI5_MOUSE-DECOY Sister chromatid cohesion protein PDS5 homolog A OS=Mus... | tr\|E9QPI5\|E9QPI5_MOUSE-DECOY Sister chromatid cohesion protein PDS5 homolog A OS=Mus... |
| sp\|E9Q8I9\|FRY_MOUSE-DECOY Protein furry homolog OS=Mus musculus OX=10090 GN=Fry PE=1... | sp\|E9Q8I9\|FRY_MOUSE-DECOY Protein furry homolog OS=Mus musculus OX=10090 GN=Fry PE=1... |
| tr\|F8WJD4\|F8WJD4_MOUSE-DECOY Symplekin OS=Mus musculus OX=10090 GN=Sympk PE=1 SV=1 | tr\|F8WJD4\|F8WJD4_MOUSE-DECOY Symplekin OS=Mus musculus OX=10090 GN=Sympk PE=1 SV=1 |

| | |
|---|---|
| Q8C1V4_MOUSE | Lysine--tRNA ligase OS=Mus musculus OX=10090 GN=Kars PE=2 SV=1 |
| tr\|B2RXX9\|B2RXX9_MOUSE-DECOY Myosin, heavy polypeptide 7, cardiac muscle, beta OS=Mus... | tr\|B2RXX9\|B2RXX9_MOUSE-DECOY Myosin, heavy polypeptide 7, cardiac muscle, beta OS=Mus... |
| sp\|Q5SSL4\|ABR_MOUSE Active breakpoint cluster region-related protein OS=Mus musculus... | sp\|Q5SSL4\|ABR_MOUSE Active breakpoint cluster region-related protein OS=Mus musculus... |
| Q3U2K1_MOUSE | Uncharacterized protein (Fragment) OS=Mus musculus OX=10090 GN=Prmt3 PE=2 SV=1 |
| sp\|Q91WJ8-2\|FUBP1_MOUSE Isoform 2 of Far upstream element-binding protein 1 OS=Mus... | Isoform 2 of Far upstream element-binding protein 1 OS=Mus musculus OX=10090 GN=Fubp1 |
| A2A601_MOUSE | Importin subunit alpha-1 (Fragment) OS=Mus musculus OX=10090 GN=Kpna2 PE=1 SV=1 |
| DPP9_MOUSE | Dipeptidyl peptidase 9 OS=Mus musculus OX=10090 GN=Dpp9 PE=1 SV=2 |
| UFL1_MOUSE | E3 UFM1-protein ligase 1 OS=Mus musculus OX=10090 GN=Ufl1 PE=1 SV=2 |
| sp\|Q4JK59\|TET2_MOUSE-DECOY Methylcytosine dioxygenase TET2 OS=Mus musculus OX=10090... | sp\|Q4JK59\|TET2_MOUSE-DECOY Methylcytosine dioxygenase TET2 OS=Mus musculus OX=10090... |
| sp\|Q6PH08-3\|ERC2_MOUSE | Isoform 3 of ERC protein 2 OS=Mus musculus OX=10090 GN=Erc2 |
| sp\|Q9CXT8\|MPPB_MOUSE Mitochondrial-processing peptidase subunit beta OS=Mus musculus... | Mitochondrial-processing peptidase subunit beta OS=Mus musculus OX=10090 GN=Pmpcb PE=1 SV=1 |
| tr\|A0A0U1RPD8\|A0A0U1RPD8_MOUSE General transcription factor 3C polypeptide 1 OS=Mus... | tr\|A0A0U1RPD8\|A0A0U1RPD8_MOUSE General transcription factor 3C polypeptide 1 OS=Mus... |
| tr\|A0A1D5RMI8\|A0A1D5RMI8_MOUSE-DECOY Alstrom syndrome protein 1 homolog OS=Mus musculus... | tr\|A0A1D5RMI8\|A0A1D5RMI8_MOUSE-DECOY Alstrom syndrome protein 1 homolog OS=Mus musculus... |
| tr\|Q3TM70\|Q3TM70_MOUSE EH-domain containing 4 OS=Mus musculus OX=10090 GN=Ehd4 PE=1... | EH-domain containing 4 OS=Mus musculus OX=10090 GN=Ehd4 PE=1 SV=1 |
| tr\|Q8BL63\|Q8BL63_MOUSE GPI-anchor transamidase OS=Mus musculus OX=10090 GN=Pigk PE=1... | tr\|Q8BL63\|Q8BL63_MOUSE GPI-anchor transamidase OS=Mus musculus OX=10090 GN=Pigk PE=1... |
| MECP2_MOUSE | Isoform B of Methyl-CpG-binding protein 2 OS=Mus musculus OX=10090 GN=Mecp2 |

| | |
|---|---|
| sp\|Q91XQ0-2\|DYH8_MOUSE-DECOY Isoform 2 of Dynein heavy chain 8, axonemal OS=Mus musculus... | sp\|Q91XQ0-2\|DYH8_MOUSE-DECOY Isoform 2 of Dynein heavy chain 8, axonemal OS=Mus musculus... |
| tr\|E9Q718\|E9Q718_MOUSE Procollagen-lysine,2-oxoglutarate 5-dioxygenase 2 OS=Mus musculus... | Procollagen-lysine,2-oxoglutarate 5-dioxygenase 2 OS=Mus musculus OX=10090 GN=Plod2 PE=1 SV=1 |
| tr\|F6WI02\|F6WI02_MOUSE Stomatin-like protein 2, mitochondrial (Fragment) OS=Mus musculus... | Stomatin-like protein 2, mitochondrial (Fragment) OS=Mus musculus OX=10090 GN=Stoml2 PE=1 SV=1 |
| tr\|Q3TPN9\|Q3TPN9_MOUSE Uncharacterized protein OS=Mus musculus OX=10090 GN=Chuk PE=2... | tr\|Q3TPN9\|Q3TPN9_MOUSE Uncharacterized protein OS=Mus musculus OX=10090 GN=Chuk PE=2... |
| tr\|E9Q526\|E9Q526_MOUSE C2 domain-containing protein 3 OS=Mus musculus OX=10090 GN=C2cd3... | tr\|E9Q526\|E9Q526_MOUSE C2 domain-containing protein 3 OS=Mus musculus OX=10090 GN=C2cd3... |
| E9Q0R3_MOUSE | Gamma-tubulin complex component OS=Mus musculus OX=10090 GN=Tubgcp3 PE=1 SV=1 |
| GPC4_MOUSE | Glypican-4 OS=Mus musculus OX=10090 GN=Gpc4 PE=1 SV=2 |
| sp\|Q99JY0\|ECHB_MOUSE Trifunctional enzyme subunit beta, mitochondrial OS=Mus musculus... | sp\|Q99JY0\|ECHB_MOUSE Trifunctional enzyme subunit beta, mitochondrial OS=Mus musculus... |
| Q80Y25_MOUSE | Capn1 protein OS=Mus musculus OX=10090 GN=Capn1 PE=2 SV=1 |
| Q8CGI3_MOUSE | Sfrs11 protein (Fragment) OS=Mus musculus OX=10090 GN=Srsf11 PE=2 SV=1 |
| Q9CXZ9_MOUSE | Uncharacterized protein OS=Mus musculus OX=10090 GN=Galk1 PE=2 SV=1 |
| RL32_MOUSE | 60S ribosomal protein L32 OS=Mus musculus OX=10090 GN=Rpl32 PE=1 SV=2 |
| sp\|Q9D4H8\|CUL2_MOUSE | Cullin-2 OS=Mus musculus OX=10090 GN=Cul2 PE=1 SV=2 |
| sp\|Q9Z110-2\|P5CS_MOUSE Isoform Short of Delta-1-pyrroline-5-carboxylate synthase... | Isoform Short of Delta-1-pyrroline-5-carboxylate synthase OS=Mus musculus OX=10090 GN=Aldh18a1 |
| RT34_MOUSE | 28S ribosomal protein S34, mitochondrial OS=Mus musculus OX=10090 GN=Mrps34 PE=1 SV=1 |

| | |
|---|---|
| tr\|E9Q1M1\|E9Q1M1_MOUSE PDZ domain-containing 2 OS=Mus musculus OX=10090 GN=Pdzd2... | tr\|E9Q1M1\|E9Q1M1_MOUSE PDZ domain-containing 2 OS=Mus musculus OX=10090 GN=Pdzd2... |
| sp\|Q059U7\|INTU_MOUSE | Protein inturned OS=Mus musculus OX=10090 GN=Intu PE=1 SV=1 |
| tr\|A0A2I3BQ39\|A0A2I3BQ39_MOUSE Ubiquitin carboxyl-terminal hydrolase (Fragment) OS=Mus... | tr\|A0A2I3BQ39\|A0A2I3BQ39_MOUSE Ubiquitin carboxyl-terminal hydrolase (Fragment) OS=Mus... |
| tr\|Q3UFZ4\|Q3UFZ4_MOUSE Uncharacterized protein OS=Mus musculus OX=10090 GN=Nol10... | tr\|Q3UFZ4\|Q3UFZ4_MOUSE Uncharacterized protein OS=Mus musculus OX=10090 GN=Nol10... |
| tr\|Q3U7E6\|Q3U7E6_MOUSE Uncharacterized protein OS=Mus musculus OX=10090 GN=Surf4... | tr\|Q3U7E6\|Q3U7E6_MOUSE Uncharacterized protein OS=Mus musculus OX=10090 GN=Surf4... |
| G3X9G2_MOUSE | Misshapen-like kinase 1 OS=Mus musculus OX=10090 GN=Mink1 PE=1 SV=1 |
| G3X981_MOUSE | Peripherin OS=Mus musculus OX=10090 GN=Prph PE=1 SV=1 |
| tr\|B2RT41\|B2RT41_MOUSE-DECOY Zinc finger, C3H1-type-containing OS=Mus musculus OX=10090... | tr\|B2RT41\|B2RT41_MOUSE-DECOY Zinc finger, C3H1-type-containing OS=Mus musculus OX=10090... |
| sp\|Q8BTM8\|FLNA_MOUSE-DECOY Filamin-A OS=Mus musculus OX=10090 GN=Flna PE=1 SV=5 | sp\|Q8BTM8\|FLNA_MOUSE-DECOY Filamin-A OS=Mus musculus OX=10090 GN=Flna PE=1 SV=5 |
| sp\|Q80X41-2\|VRK1_MOUSE Isoform 2 of Serine/threonine-protein kinase VRK1 OS=Mus musculus... | Isoform 2 of Serine/threonine-protein kinase VRK1 OS=Mus musculus OX=10090 GN=Vrk1 |
| A2AA71_MOUSE | Protein transport protein Sec24A OS=Mus musculus OX=10090 GN=Sec24a PE=1 SV=1 |
| ARL1_MOUSE | ADP-ribosylation factor-like protein 1 OS=Mus musculus OX=10090 GN=Arl1 PE=1 SV=1 |
| tr\|Q14DP3\|Q14DP3_MOUSE Leucyl-tRNA synthetase, mitochondrial OS=Mus musculus OX=10090... | Leucyl-tRNA synthetase, mitochondrial OS=Mus musculus OX=10090 GN=Lars2 PE=2 SV=1 |
| B2RXS1_MOUSE | Nbeal2 protein OS=Mus musculus OX=10090 GN=Nbeal2 PE=2 SV=1 |
| sp\|A2AN08-3\|UBR4_MOUSE-DECOY Isoform 3 of E3 ubiquitin-protein ligase UBR4 OS=Mus... | sp\|A2AN08-3\|UBR4_MOUSE-DECOY Isoform 3 of E3 ubiquitin-protein ligase UBR4 OS=Mus... |

| | |
|---|---|
| tr\|Q3TJX6\|Q3TJX6_MOUSE Uncharacterized protein OS=Mus musculus OX=10090 GN=Utp18... | tr\|Q3TJX6\|Q3TJX6_MOUSE Uncharacterized protein OS=Mus musculus OX=10090 GN=Utp18... |
| tr\|Q3UQJ9\|Q3UQJ9_MOUSE Uncharacterized protein OS=Mus musculus OX=10090 GN=Trps1... | tr\|Q3UQJ9\|Q3UQJ9_MOUSE Uncharacterized protein OS=Mus musculus OX=10090 GN=Trps1... |
| tr\|A0A286YDA2\|A0A286YDA2_MOUSE Nucleolar and coiled-body phosphoprotein 1 OS=Mus... | tr\|A0A286YDA2\|A0A286YDA2_MOUSE Nucleolar and coiled-body phosphoprotein 1 OS=Mus... |
| sp\|P08551\|NFL_MOUSE Neurofilament light polypeptide OS=Mus musculus OX=10090 GN=Nefl... | Neurofilament light polypeptide OS=Mus musculus OX=10090 GN=Nefl PE=1 SV=5 |
| sp\|Q8BH58\|TIPRL_MOUSETIP41-like protein OS=Mus musculus OX=10090 GN=Tiprl PE=1 SV=1 | sp\|Q8BH58\|TIPRL_MOUSE TIP41-like protein OS=Mus musculus OX=10090 GN=Tiprl PE=1 SV=1 |
| tr\|A0A0R4J1W7\|A0A0R4J1W7_MOUSE CDC23 (Cell division cycle 23, yeast, homolog), isoform... | tr\|A0A0R4J1W7\|A0A0R4J1W7_MOUSE CDC23 (Cell division cycle 23, yeast, homolog), isoform... |
| tr\|Q6PET9\|Q6PET9_MOUSE DNA topoisomerase 2 (Fragment) OS=Mus musculus OX=10090 GN=Top2a... | tr\|Q6PET9\|Q6PET9_MOUSE DNA topoisomerase 2 (Fragment) OS=Mus musculus OX=10090 GN=Top2a... |
| Q5U4B1_MOUSE | Replication factor C subunit 1 OS=Mus musculus OX=10090 GN=Rfc1 PE=1 SV=1 |
| Q3US09_MOUSE | Uncharacterized protein (Fragment) OS=Mus musculus OX=10090 GN=Thoc2 PE=2 SV=1 |
| tr\|Q3UGZ4\|Q3UGZ4_MOUSE-DECOY Spectrin beta chain OS=Mus musculus OX=10090 GN=Sptbn2... | tr\|Q3UGZ4\|Q3UGZ4_MOUSE-DECOY Spectrin beta chain OS=Mus musculus OX=10090 GN=Sptbn2... |
| B2RUG9_MOUSE | Adenomatosis polyposis coli OS=Mus musculus OX=10090 GN=Apc PE=1 SV=1 |
| sp\|Q9Z0H8-2\|CLIP2_MOUSE Isoform 2 of CAP-Gly domain-containing linker protein 2 OS=Mus... | sp\|Q9Z0H8-2\|CLIP2_MOUSE Isoform 2 of CAP-Gly domain-containing linker protein 2 OS=Mus... |
| sp\|Q8C1B7-3\|SEP11_MOUSE | Isoform 3 of Septin-11 OS=Mus musculus OX=10090 GN=Sept11 |
| sp\|F8VPN2-2\|TEX15_MOUSE Isoform 2 of Testis-expressed protein 15 OS=Mus musculus... | Isoform 2 of Testis-expressed protein 15 OS=Mus musculus OX=10090 GN=Tex15 |
| tr\|Q4FE56\|Q4FE56_MOUSE Probable ubiquitin carboxyl-terminal hydrolase FAF-X OS=Mus... | tr\|Q4FE56\|Q4FE56_MOUSE Probable ubiquitin carboxyl-terminal hydrolase FAF-X OS=Mus... |

| | |
|---|---|
| tr\|A0A1D5RLM8\|A0A1D5RLM8_MOUSE-DECOY Predicted gene 11639 OS=Mus musculus OX=10090... | tr\|A0A1D5RLM8\|A0A1D5RLM8_MOUSE-DECOY Predicted gene 11639 OS=Mus musculus OX=10090... |
| tr\|F6RGI3\|F6RGI3_MOUSE-DECOY Predicted gene 340 OS=Mus musculus OX=10090 GN=Gm340... | tr\|F6RGI3\|F6RGI3_MOUSE-DECOY Predicted gene 340 OS=Mus musculus OX=10090 GN=Gm340... |
| sp\|Q60749\|KHDR1_MOUSEKH domain-containing, RNA-binding, signal transduction-associated protein 1 OS=Mus musculus OX=10090 GN=Khdrbs1 PE=1 SV=2 | sp\|Q60749\|KHDR1_MOUSE KH domain-containing, RNA-binding, signal transduction-associated protein 1 OS=Mus musculus OX=10090 GN=Khdrbs1 PE=1 SV=2 |
| tr\|A2A9H6\|A2A9H6_MOUSE-DECOY Predicted gene 884 OS=Mus musculus OX=10090 GN=Gm884... | tr\|A2A9H6\|A2A9H6_MOUSE-DECOY Predicted gene 884 OS=Mus musculus OX=10090 GN=Gm884... |
| tr\|E9Q4K7\|E9Q4K7_MOUSE Kinesin family member 13B OS=Mus musculus OX=10090 GN=Kif13b... | tr\|E9Q4K7\|E9Q4K7_MOUSE Kinesin family member 13B OS=Mus musculus OX=10090 GN=Kif13b... |
| tr\|A0A087WS83\|A0A087WS83_MOUSE Voltage-dependent R-type calcium channel subunit alpha... | tr\|A0A087WS83\|A0A087WS83_MOUSE Voltage-dependent R-type calcium channel subunit alpha... |
| sp\|P27048\|RSMB_MOUSE Small nuclear ribonucleoprotein-associated protein B OS=Mus... | sp\|P27048\|RSMB_MOUSE Small nuclear ribonucleoprotein-associated protein B OS=Mus... |
| sp\|Q8K2Q7\|BROX_MOUSE BRO1 domain-containing protein BROX OS=Mus musculus OX=10090... | sp\|Q8K2Q7\|BROX_MOUSE BRO1 domain-containing protein BROX OS=Mus musculus OX=10090... |
| tr\|B9EKS2\|B9EKS2_MOUSE-DECOY Jumonji domain containing 1B OS=Mus musculus OX=10090... | tr\|B9EKS2\|B9EKS2_MOUSE-DECOY Jumonji domain containing 1B OS=Mus musculus OX=10090... |
| Q9D3C4_MOUSE | Actin-related protein 2/3 complex subunit 4 OS=Mus musculus OX=10090 GN=Arpc4 PE=2 SV=1 |
| sp\|Q0KL02-3\|TRIO_MOUSE | Isoform 3 of Triple functional domain protein OS=Mus musculus OX=10090 GN=Trio |
| Q3KR45_MOUSE | DNA (cytosine-5)-methyltransferase 3B OS=Mus musculus OX=10090 GN=Dnmt3b PE=1 SV=1 |
| sp\|Q8R5H1-5\|UBP15_MOUSE Isoform 5 of Ubiquitin carboxyl-terminal hydrolase 15 OS=Mus... | Isoform 5 of Ubiquitin carboxyl-terminal hydrolase 15 OS=Mus musculus OX=10090 GN=Usp15 |
| A9JR55_MOUSE | Mybpc3 protein OS=Mus musculus OX=10090 GN=Mybpc3 PE=2 SV=1 |

| | |
|---|---|
| BIG2_MOUSE | Brefeldin A-inhibited guanine nucleotide-exchange protein 2 OS=Mus musculus OX=10090 GN=Arfgef2 PE=1 SV=1 |
| A0A1B0GRJ4_MOUSE | Predicted gene 7579 OS=Mus musculus OX=10090 GN=Gm7579 PE=4 SV=1 |
| Q3TZU7_MOUSE | Sorting nexin OS=Mus musculus OX=10090 GN=Snx9 PE=2 SV=1 |
| sp\|Q9WTU3-3\|SCN8A_MOUSE-DECOY Isoform 3 of Sodium channel protein type 8 subunit... | sp\|Q9WTU3-3\|SCN8A_MOUSE-DECOY Isoform 3 of Sodium channel protein type 8 subunit... |
| Q8CGG9_MOUSE | Fes protein (Fragment) OS=Mus musculus OX=10090 GN=Fes PE=2 SV=1 |
| B1ARD8_MOUSE | Schlafen 8 OS=Mus musculus OX=10090 GN=Slfn8 PE=1 SV=1 |
| Q7TPY0_MOUSE | Gtf2h1 protein OS=Mus musculus OX=10090 GN=Gtf2h1 PE=2 SV=1 |
| KC1E_MOUSE | Casein kinase I isoform epsilon OS=Mus musculus OX=10090 GN=Csnk1e PE=1 SV=2 |
| B7ZMR9_MOUSE | MCG12507, isoform CRA_a OS=Mus musculus OX=10090 GN=Zfp800 PE=1 SV=1 |
| sp\|Q6P5B0\|RRP12_MOUSERRP12-like protein OS=Mus musculus OX=10090 GN=Rrp12 PE=1 SV=1 | sp\|Q6P5B0\|RRP12_MOUSE RRP12-like protein OS=Mus musculus OX=10090 GN=Rrp12 PE=1 SV=1 |
| DVL2_MOUSE | Segment polarity protein dishevelled homolog DVL-2 OS=Mus musculus OX=10090 GN=Dvl2 PE=1 SV=2 |
| ADT1_MOUSE | ADP/ATP translocase 1 OS=Mus musculus OX=10090 GN=Slc25a4 PE=1 SV=4 |
| sp\|Q9R269\|PEPL_MOUSE-DECOY Periplakin OS=Mus musculus OX=10090 GN=Ppl PE=1 SV=1 | sp\|Q9R269\|PEPL_MOUSE-DECOY Periplakin OS=Mus musculus OX=10090 GN=Ppl PE=1 SV=1 |
| sp\|Q61102\|ABCB7_MOUSEATP-binding cassette sub-family B member 7, mitochondrial OS=Mus musculus OX=10090 GN=Abcb7 PE=1 SV=3 | sp\|Q61102\|ABCB7_MOUSE ATP-binding cassette sub-family B member 7, mitochondrial OS=Mus musculus OX=10090 GN=Abcb7 PE=1 SV=3 |
| Q3TL71_MOUSE | Uncharacterized protein OS=Mus musculus OX=10090 GN=Hnrnpk PE=2 SV=1 |
| A0A286YD72_MOUSE | Voltage-dependent L-type calcium channel subunit alpha OS=Mus musculus OX=10090 GN=Cacna1d PE=1 SV=1 |

| | |
|---|---|
| B1B1A8_MOUSE | Myosin light chain kinase, smooth muscle OS=Mus musculus OX=10090 GN=Mylk PE=1 SV=1 |
| Q7TQR2_MOUSE | Olfactory receptor OS=Mus musculus OX=10090 GN=Olfr1465 PE=3 SV=1 |
| sp\|Q3UYI5\|RGL3_MOUSE | Ral guanine nucleotide dissociation stimulator-like 3 OS=Mus musculus OX=10090 GN=Rgl3 PE=1 SV=2 |
| Q6A049_MOUSE | MKIAA0546 protein (Fragment) OS=Mus musculus OX=10090 GN=Zfc3h1 PE=2 SV=1 |
| Q3TXH6_MOUSE | Uncharacterized protein OS=Mus musculus OX=10090 GN=Smarcd2 PE=2 SV=1 |
| A2A9H6_MOUSE | Predicted gene 884 OS=Mus musculus OX=10090 GN=Gm884 PE=1 SV=1 |
| Q9D122_MOUSE | Uncharacterized protein OS=Mus musculus OX=10090 GN=Krtap4-2 PE=2 SV=1 |
| Q9JJD8_MOUSE | Brain cDNA, clone MNCb-1272, similar to Mus musculus chaperonin subunit 2 (beta) (Cct2), mRNA OS=Mus musculus OX=10090 GN=Cct2 PE=2 SV=1 |
| SMRCD_MOUSE | Isoform 2 of SWI/SNF-related matrix-associated actin-dependent regulator of chromatin subfamily A containing DEAD/H box 1 OS=Mus musculus OX=10090 GN=Smarcad1 |
| E9PWI6_MOUSE | GRIP and coiled-coil domain-containing protein 2 OS=Mus musculus OX=10090 GN=Gcc2 PE=1 SV=1 |
| O88348_MOUSE | Deubiquitinating enzyme (Fragment) OS=Mus musculus OX=10090 GN=Usp12 PE=2 SV=1 |
| S4R1W5_MOUSE | RNA-binding motif protein 6 OS=Mus musculus OX=10090 GN=Rbm6 PE=1 SV=1 |
| sp\|P97480-2\|EYA3_MOUSE | Isoform 2 of Eyes absent homolog 3 OS=Mus musculus OX=10090 GN=Eya3 |
| E9QP54_MOUSE | Jouberin OS=Mus musculus OX=10090 GN=Ahi1 PE=1 SV=1 |
| B2KFD2_MOUSE | 2810474O19Rik protein OS=Mus musculus OX=10090 GN=2810474O19Rik PE=2 SV=1 |

| | |
|---|---|
| sp\|Q8BW94-2\|DYH3_MOUSE | Isoform 2 of Dynein heavy chain 3, axonemal OS=Mus musculus OX=10090 GN=Dnah3 |
| E9QKE4_MOUSE | Rab3 GTPase-activating protein non-catalytic subunit OS=Mus musculus OX=10090 GN=Rab3gap2 PE=1 SV=1 |
| DDX6_MOUSE | Probable ATP-dependent RNA helicase DDX6 OS=Mus musculus OX=10090 GN=Ddx6 PE=1 SV=1 |
| MSH6_MOUSE | DNA mismatch repair protein Msh6 OS=Mus musculus OX=10090 GN=Msh6 PE=1 SV=3 |
| A0A0R4J1Y3_MOUSE | Probable tRNA N6-adenosine threonylcarbamoyltransferase OS=Mus musculus OX=10090 GN=Osgep PE=1 SV=1 |
| D3YZF6_MOUSE | Family with sequence similarity 205, member A1 OS=Mus musculus OX=10090 GN=Fam205a1 PE=4 SV=1 |
| Q3UPA1_MOUSE | Guanine nucleotide binding protein, alpha 11 OS=Mus musculus OX=10090 GN=Gna11 PE=1 SV=1 |
| sp\|P70451\|FER_MOUSE Tyrosine-protein kinase Fer OS=Mus musculus OX=10090 GN=Fer PE=1... | sp\|P70451\|FER_MOUSE Tyrosine-protein kinase Fer OS=Mus musculus OX=10090 GN=Fer PE=1... |
| sp\|Q3UH68-3\|LIMC1_MOUSE Isoform 3 of LIM and calponin homology domains-containing... | sp\|Q3UH68-3\|LIMC1_MOUSE Isoform 3 of LIM and calponin homology domains-containing... |
| sp\|Q8CF89\|TAB1_MOUSE TGF-beta-activated kinase 1 and MAP3K7-binding protein 1 OS=Mus... | sp\|Q8CF89\|TAB1_MOUSE TGF-beta-activated kinase 1 and MAP3K7-binding protein 1 OS=Mus... |
| sp\|Q8R2N2\|UTP4_MOUSE U3 small nucleolar RNA-associated protein 4 homolog OS=Mus musculus... | U3 small nucleolar RNA-associated protein 4 homolog OS=Mus musculus OX=10090 GN=Utp4 PE=2 SV=3 |
| sp\|Q8VI56-2\|LRP4_MOUSE-DECOY Isoform 2 of Low-density lipoprotein receptor-related... | sp\|Q8VI56-2\|LRP4_MOUSE-DECOY Isoform 2 of Low-density lipoprotein receptor-related... |
| sp\|Q9Z0R6-2\|ITSN2_MOUSE-DECOY Isoform 2 of Intersectin-2 OS=Mus musculus OX=10090... | sp\|Q9Z0R6-2\|ITSN2_MOUSE-DECOY Isoform 2 of Intersectin-2 OS=Mus musculus OX=10090... |
| tr\|B2RY21\|B2RY21_MOUSE-DECOY Abca12 protein OS=Mus musculus OX=10090 GN=Abca12 PE=2... | tr\|B2RY21\|B2RY21_MOUSE-DECOY Abca12 protein OS=Mus musculus OX=10090 GN=Abca12 PE=2... |

| | |
|---|---|
| tr\|F6VUT6\|F6VUT6_MOUSE-DECOY Predicted gene 8251 OS=Mus musculus OX=10090 GN=Gm8251... | tr\|F6VUT6\|F6VUT6_MOUSE-DECOY Predicted gene 8251 OS=Mus musculus OX=10090 GN=Gm8251... |
| tr\|G5E8B1\|G5E8B1_MOUSE Protein tyrosine phosphatase, non-receptor type 13 OS=Mus... | tr\|G5E8B1\|G5E8B1_MOUSE Protein tyrosine phosphatase, non-receptor type 13 OS=Mus... |
| tr\|Q6NV50\|Q6NV50_MOUSE-DECOY Catenin (Cadherin associated protein), alpha 1 OS=Mus... | tr\|Q6NV50\|Q6NV50_MOUSE-DECOY Catenin (Cadherin associated protein), alpha 1 OS=Mus... |
| tr\|Q8C1T2\|Q8C1T2_MOUSE Uncharacterized protein OS=Mus musculus OX=10090 GN=Psmd6... | tr\|Q8C1T2\|Q8C1T2_MOUSE Uncharacterized protein OS=Mus musculus OX=10090 GN=Psmd6... |
| tr\|Q9CTM9\|Q9CTM9_MOUSE Uncharacterized protein (Fragment) OS=Mus musculus OX=10090... | tr\|Q9CTM9\|Q9CTM9_MOUSE Uncharacterized protein (Fragment) OS=Mus musculus OX=10090... |
| tr\|Q9D3V6\|Q9D3V6_MOUSE Core histone macro-H2A OS=Mus musculus OX=10090 GN=H2afy3... | Core histone macro-H2A OS=Mus musculus OX=10090 GN=H2afy3 PE=2 SV=1 |
| Q3UYK8_MOUSE | Uncharacterized protein OS=Mus musculus OX=10090 GN=Rab3gap1 PE=2 SV=1 |
| tr\|A2ALL9\|A2ALL9_MOUSE Calcium-transporting ATPase OS=Mus musculus OX=10090 GN=Atp2b3... | Calcium-transporting ATPase OS=Mus musculus OX=10090 GN=Atp2b3 PE=1 SV=1 |
| E9QAQ7_MOUSE | AT-rich interactive domain-containing protein 1A OS=Mus musculus OX=10090 GN=Arid1a PE=1 SV=1 |
| ATLA3_MOUSE | Isoform 2 of Atlastin-3 OS=Mus musculus OX=10090 GN=Atl3 |
| tr\|B2RRE0\|B2RRE0_MOUSE-DECOY A kinase (PRKA) anchor protein (Gravin) 12 OS=Mus musculus... | tr\|B2RRE0\|B2RRE0_MOUSE-DECOY A kinase (PRKA) anchor protein (Gravin) 12 OS=Mus musculus... |
| tr\|Q6DFZ1\|Q6DFZ1_MOUSE-DECOY Golgi-specific brefeldin A-resistance factor 1 OS=Mus... | tr\|Q6DFZ1\|Q6DFZ1_MOUSE-DECOY Golgi-specific brefeldin A-resistance factor 1 OS=Mus... |
| B2RY30_MOUSE | Exosc9 protein OS=Mus musculus OX=10090 GN=Exosc9 PE=2 SV=1 |
| sp\|Q8BGQ7\|SYAC_MOUSE-DECOY Alanine--tRNA ligase, cytoplasmic OS=Mus musculus OX=10090... | sp\|Q8BGQ7\|SYAC_MOUSE-DECOY Alanine--tRNA ligase, cytoplasmic OS=Mus musculus OX=10090... |
| tr\|A0A140T8T2\|A0A140T8T2_MOUSE-DECOY Slit homolog 2 protein OS=Mus musculus OX=10090... | tr\|A0A140T8T2\|A0A140T8T2_MOUSE-DECOY Slit homolog 2 protein OS=Mus musculus OX=10090... |

| | |
|---|---|
| sp\|Q9QY15-2\|DDX25_MOUSE Isoform 2 of ATP-dependent RNA helicase DDX25 OS=Mus musculus... | sp\|Q9QY15-2\|DDX25_MOUSE Isoform 2 of ATP-dependent RNA helicase DDX25 OS=Mus musculus... |
| sp\|Q8BQM9-2\|MD12L_MOUSE-DECOY Isoform 2 of Mediator of RNA polymerase II transcription... | sp\|Q8BQM9-2\|MD12L_MOUSE-DECOY Isoform 2 of Mediator of RNA polymerase II transcription... |
| sp\|Q80V86\|INT8_MOUSE Integrator complex subunit 8 OS=Mus musculus OX=10090 GN=Ints8... | sp\|Q80V86\|INT8_MOUSE Integrator complex subunit 8 OS=Mus musculus OX=10090 GN=Ints8... |
| tr\|A0A140LIZ7\|A0A140LIZ7_MOUSE-DECOY NHS-like protein 1 OS=Mus musculus OX=10090... | tr\|A0A140LIZ7\|A0A140LIZ7_MOUSE-DECOY NHS-like protein 1 OS=Mus musculus OX=10090... |
| sp\|Q9Z1K5\|ARI1_MOUSE E3 ubiquitin-protein ligase ARIH1 OS=Mus musculus OX=10090 GN=Arih1... | sp\|Q9Z1K5\|ARI1_MOUSE E3 ubiquitin-protein ligase ARIH1 OS=Mus musculus OX=10090 GN=Arih1... |
| tr\|A0A087WPT7\|A0A087WPT7_MOUSE Phosphatidylinositol 3,4,5-trisphosphate 5-phosphatase... | tr\|A0A087WPT7\|A0A087WPT7_MOUSE Phosphatidylinositol 3,4,5-trisphosphate 5-phosphatase... |
| sp\|Q8BYA0\|TBCD_MOUSE Tubulin-specific chaperone D OS=Mus musculus OX=10090 GN=Tbcd... | Tubulin-specific chaperone D OS=Mus musculus OX=10090 GN=Tbcd PE=1 SV=1 |
| D3Z0Y2_MOUSE | Peroxiredoxin-6 OS=Mus musculus OX=10090 GN=Prdx6 PE=1 SV=1 |
| tr\|Q3UH59\|Q3UH59_MOUSE-DECOY Myosin-10 OS=Mus musculus OX=10090 GN=Myh10 PE=1 SV=1 | tr\|Q3UH59\|Q3UH59_MOUSE-DECOY Myosin-10 OS=Mus musculus OX=10090 GN=Myh10 PE=1 SV=1 |
| PPCE_MOUSE | Prolyl endopeptidase OS=Mus musculus OX=10090 GN=Prep PE=1 SV=1 |
| sp\|Q9D4H8-2\|CUL2_MOUSE-DECOY Isoform 2 of Cullin-2 OS=Mus musculus OX=10090 GN=Cul2 | sp\|Q9D4H8-2\|CUL2_MOUSE-DECOY Isoform 2 of Cullin-2 OS=Mus musculus OX=10090 GN=Cul2 |
| tr\|Q3U048\|Q3U048_MOUSE-DECOY Uncharacterized protein (Fragment) OS=Mus musculus OX=10090... | tr\|Q3U048\|Q3U048_MOUSE-DECOY Uncharacterized protein (Fragment) OS=Mus musculus OX=10090... |
| Q8C1L7_MOUSE | 40S ribosomal protein S21 OS=Mus musculus OX=10090 GN=mCG_6739 PE=2 SV=1 |
| sp\|Q04859\|MAK_MOUSE Serine/threonine-protein kinase MAK OS=Mus musculus OX=10090... | sp\|Q04859\|MAK_MOUSE Serine/threonine-protein kinase MAK OS=Mus musculus OX=10090... |
| tr\|Q3T9Y8\|Q3T9Y8_MOUSE Eukaryotic translation initiation factor 3 subunit I OS=Mus... | tr\|Q3T9Y8\|Q3T9Y8_MOUSE Eukaryotic translation initiation factor 3 subunit I OS=Mus... |

| | |
|---|---|
| tr\|F6T9C3\|F6T9C3_MOUSE Translation initiation factor eIF-2B subunit epsilon (Fragment)... | tr\|F6T9C3\|F6T9C3_MOUSE Translation initiation factor eIF-2B subunit epsilon (Fragment)... |
| sp\|Q924W5\|SMC6_MOUSE Structural maintenance of chromosomes protein 6 OS=Mus musculus... | sp\|Q924W5\|SMC6_MOUSE Structural maintenance of chromosomes protein 6 OS=Mus musculus... |
| Q3UGX2_MOUSE | Spectrin beta chain OS=Mus musculus OX=10090 GN=Sptb PE=1 SV=1 |
| A2A702_MOUSE | Eukaryotic translation initiation factor 3 subunit M OS=Mus musculus OX=10090 GN=Eif3m PE=1 SV=1 |
| PURA_MOUSE | Transcriptional activator protein Pur-alpha OS=Mus musculus OX=10090 GN=Pura PE=1 SV=1 |
| tr\|E0CZ42\|E0CZ42_MOUSE-DECOY Fer-1-like 6 (C. elegans) OS=Mus musculus OX=10090 GN=Fer1l6... | tr\|E0CZ42\|E0CZ42_MOUSE-DECOY Fer-1-like 6 (C. elegans) OS=Mus musculus OX=10090 GN=Fer1l6... |
| B2RT32_MOUSE | Predicted gene, OTTMUSG00000010105 OS=Mus musculus OX=10090 GN=Gm13034 PE=2 SV=1 |
| sp\|Q8BI72\|CARF_MOUSE CDKN2A-interacting protein OS=Mus musculus OX=10090 GN=Cdkn2aip... | CDKN2A-interacting protein OS=Mus musculus OX=10090 GN=Cdkn2aip PE=1 SV=1 |
| sp\|Q8CJ27\|ASPM_MOUSE-DECOY Abnormal spindle-like microcephaly-associated protein... | sp\|Q8CJ27\|ASPM_MOUSE-DECOY Abnormal spindle-like microcephaly-associated protein... |
| tr\|Q3TWH2\|Q3TWH2_MOUSE-DECOY Uncharacterized protein OS=Mus musculus OX=10090 GN=Nnt... | tr\|Q3TWH2\|Q3TWH2_MOUSE-DECOY Uncharacterized protein OS=Mus musculus OX=10090 GN=Nnt... |
| tr\|Q3UX80\|Q3UX80_MOUSE Uncharacterized protein OS=Mus musculus OX=10090 GN=Snrpf... | tr\|Q3UX80\|Q3UX80_MOUSE Uncharacterized protein OS=Mus musculus OX=10090 GN=Snrpf... |
| tr\|B7ZCC3\|B7ZCC3_MOUSE-DECOY Ataxin 7-like 2 OS=Mus musculus OX=10090 GN=Atxn7l2... | tr\|B7ZCC3\|B7ZCC3_MOUSE-DECOY Ataxin 7-like 2 OS=Mus musculus OX=10090 GN=Atxn7l2... |
| A0A0B4J1L8_MOUSE | Immunoglobulin superfamily member 2 OS=Mus musculus OX=10090 GN=Cd101 PE=1 SV=1 |
| tr\|Q3T9H2\|Q3T9H2_MOUSE-DECOY Uncharacterized protein (Fragment) OS=Mus musculus OX=10090... | tr\|Q3T9H2\|Q3T9H2_MOUSE-DECOY Uncharacterized protein (Fragment) OS=Mus musculus OX=10090... |

| | |
|---|---|
| sp\|Q9WUU9\|GANP_MOUSE-DECOY Germinal-center associated nuclear protein OS=Mus musculus... | sp\|Q9WUU9\|GANP_MOUSE-DECOY Germinal-center associated nuclear protein OS=Mus musculus... |
| NKTR_MOUSE | NK-tumor recognition protein OS=Mus musculus OX=10090 GN=Nktr PE=1 SV=4 |
| tr\|E9Q9A3\|E9Q9A3_MOUSE-DECOY Clathrin coat assembly protein AP180 OS=Mus musculus... | tr\|E9Q9A3\|E9Q9A3_MOUSE-DECOY Clathrin coat assembly protein AP180 OS=Mus musculus... |
| D3Z0B3_MOUSE | Acetyl-CoA carboxylase 2 (Fragment) OS=Mus musculus OX=10090 GN=Acacb PE=1 SV=1 |
| A0A0R4J0U8_MOUSE | RIKEN cDNA 1190005F20, isoform CRA_a OS=Mus musculus OX=10090 GN=Trmt1l PE=1 SV=1 |
| sp\|Q9D2I5-5\|ARMC9_MOUSE | Isoform 5 of LisH domain-containing protein ARMC9 OS=Mus musculus OX=10090 GN=Armc9 |
| ECHM_MOUSE | Enoyl-CoA hydratase, mitochondrial OS=Mus musculus OX=10090 GN=Echs1 PE=1 SV=1 |
| tr\|A0A0G2JDZ9\|A0A0G2JDZ9_MOUSE-DECOY Ankyrin repeat domain-containing protein 17... | tr\|A0A0G2JDZ9\|A0A0G2JDZ9_MOUSE-DECOY Ankyrin repeat domain-containing protein 17... |
| Q3UIJ0_MOUSE | Uncharacterized protein OS=Mus musculus OX=10090 GN=Cct7 PE=2 SV=1 |
| D3Z131_MOUSE | THO complex subunit 6 homolog OS=Mus musculus OX=10090 GN=Thoc6 PE=1 SV=1 |
| tr\|B4YB44\|B4YB44_MOUSE Retrotransposon-like 1 OS=Mus musculus OX=10090 GN=Rtl1 PE=2... | tr\|B4YB44\|B4YB44_MOUSE Retrotransposon-like 1 OS=Mus musculus OX=10090 GN=Rtl1 PE=2... |
| A0A140LJ59_MOUSE | Eukaryotic translation initiation factor 3 subunit K OS=Mus musculus OX=10090 GN=Eif3k PE=1 SV=1 |
| tr\|A0A096P6K7\|A0A096P6K7_MOUSE Transcription termination factor 1b, mitochondrial... | tr\|A0A096P6K7\|A0A096P6K7_MOUSE Transcription termination factor 1b, mitochondrial... |
| tr\|Q8K314\|Q8K314_MOUSE Atp2b1 protein (Fragment) OS=Mus musculus OX=10090 GN=Atp2b1... | tr\|Q8K314\|Q8K314_MOUSE Atp2b1 protein (Fragment) OS=Mus musculus OX=10090 GN=Atp2b1... |
| PTN11_MOUSE | Isoform 2 of Tyrosine-protein phosphatase non-receptor type 11 OS=Mus musculus OX=10090 GN=Ptpn11 |

| | |
|---|---|
| Q8BQD9_MOUSE | Uncharacterized protein OS=Mus musculus OX=10090 PE=2 SV=1 |
| tr\|Q5SXR4\|Q5SXR4_MOUSE Probable ATP-dependent RNA helicase DHX40 (Fragment) OS=Mus... | tr\|Q5SXR4\|Q5SXR4_MOUSE Probable ATP-dependent RNA helicase DHX40 (Fragment) OS=Mus... |
| D3YZ98_MOUSE | Phosphatidylinositol 3-kinase catalytic subunit type 3 OS=Mus musculus OX=10090 GN=Pik3c3 PE=1 SV=1 |
| tr\|Q05BH3\|Q05BH3_MOUSE DEAH (Asp-Glu-Ala-His) box polypeptide 16 OS=Mus musculus... | tr\|Q05BH3\|Q05BH3_MOUSE DEAH (Asp-Glu-Ala-His) box polypeptide 16 OS=Mus musculus... |
| sp\|A6H603\|NWD1_MOUSE-DECOY NACHT domain- and WD repeat-containing protein 1 OS=Mus... | sp\|A6H603\|NWD1_MOUSE-DECOY NACHT domain- and WD repeat-containing protein 1 OS=Mus... |
| sp\|O54990-2\|PROM1_MOUSE-DECOY Isoform 2 of Prominin-1 OS=Mus musculus OX=10090 GN=Prom1 | sp\|O54990-2\|PROM1_MOUSE-DECOY Isoform 2 of Prominin-1 OS=Mus musculus OX=10090 GN=Prom1 |
| SDHA_MOUSE | Succinate dehydrogenase [ubiquinone] flavoprotein subunit, mitochondrial OS=Mus musculus OX=10090 GN=Sdha PE=1 SV=1 |
| sp\|Q08122-1\|TLE3_MOUSE-DECOY Isoform 1 of Transducin-like enhancer protein 3 OS=Mus... | sp\|Q08122-1\|TLE3_MOUSE-DECOY Isoform 1 of Transducin-like enhancer protein 3 OS=Mus... |
| Q9D050_MOUSE | Mitochondrial carrier homolog 2 OS=Mus musculus OX=10090 GN=Mtch2 PE=1 SV=1 |
| tr\|D3Z7E6\|D3Z7E6_MOUSE Platelet-activating factor acetylhydrolase IB subunit gamma... | tr\|D3Z7E6\|D3Z7E6_MOUSE Platelet-activating factor acetylhydrolase IB subunit gamma... |
| IPO13_MOUSE | Isoform 2 of Importin-13 OS=Mus musculus OX=10090 GN=Ipo13 |
| sp\|P11881-2\|ITPR1_MOUSE-DECOY Isoform 2 of Inositol 1,4,5-trisphosphate receptor... | sp\|P11881-2\|ITPR1_MOUSE-DECOY Isoform 2 of Inositol 1,4,5-trisphosphate receptor... |
| tr\|F8VQ05\|F8VQ05_MOUSE FRY-like transcription coactivator OS=Mus musculus OX=10090... | tr\|F8VQ05\|F8VQ05_MOUSE FRY-like transcription coactivator OS=Mus musculus OX=10090... |
| sp\|P35569\|IRS1_MOUSE-DECOY Insulin receptor substrate 1 OS=Mus musculus OX=10090... | sp\|P35569\|IRS1_MOUSE-DECOY Insulin receptor substrate 1 OS=Mus musculus OX=10090... |

| | |
|---|---|
| sp\|Q8CH18-3\|CCAR1_MOUSE-DECOY Isoform 3 of Cell division cycle and apoptosis regulator... | sp\|Q8CH18-3\|CCAR1_MOUSE-DECOY Isoform 3 of Cell division cycle and apoptosis regulator... |
| Q9DBU8_MOUSE | Uncharacterized protein OS=Mus musculus OX=10090 GN=Ap3m1 PE=2 SV=1 |
| sp\|Q99LI9\|CLP1_MOUSE Polyribonucleotide 5'-hydroxyl-kinase Clp1 OS=Mus musculus OX=10090... | sp\|Q99LI9\|CLP1_MOUSE Polyribonucleotide 5'-hydroxyl-kinase Clp1 OS=Mus musculus OX=10090... |
| tr\|Q9CRE9\|Q9CRE9_MOUSE Uncharacterized protein (Fragment) OS=Mus musculus OX=10090... | Uncharacterized protein (Fragment) OS=Mus musculus OX=10090 GN=Eif2s3x PE=2 SV=1 |
| tr\|A0A087WPJ1\|A0A087WPJ1_MOUSE-DECOY RIKEN cDNA 1700017N19 gene OS=Mus musculus OX=10090... | tr\|A0A087WPJ1\|A0A087WPJ1_MOUSE-DECOY RIKEN cDNA 1700017N19 gene OS=Mus musculus OX=10090... |
| sp\|O54692\|ZW10_MOUSE Centromere/kinetochore protein zw10 homolog OS=Mus musculus... | sp\|O54692\|ZW10_MOUSE Centromere/kinetochore protein zw10 homolog OS=Mus musculus... |
| NFH_MOUSE | Neurofilament heavy polypeptide OS=Mus musculus OX=10090 GN=Nefh PE=1 SV=3 |
| SAE1_MOUSE | SUMO-activating enzyme subunit 1 OS=Mus musculus OX=10090 GN=Sae1 PE=1 SV=1 |
| sp\|Q6PHU5-2\|SORT_MOUSE | Isoform 2 of Sortilin OS=Mus musculus OX=10090 GN=Sort1 |
| tr\|E9PZP8\|E9PZP8_MOUSE HECT and RLD domain-containing E3 ubiquitin protein ligase... | HECT and RLD domain-containing E3 ubiquitin protein ligase family member 1 OS=Mus musculus OX=10090 GN=Herc1 PE=1 SV=1 |
| sp\|Q8CG48\|SMC2_MOUSE-DECOY Structural maintenance of chromosomes protein 2 OS=Mus... | sp\|Q8CG48\|SMC2_MOUSE-DECOY Structural maintenance of chromosomes protein 2 OS=Mus... |
| sp\|Q7TMB8-2\|CYFP1_MOUSE-DECOY Isoform 2 of Cytoplasmic FMR1-interacting protein 1... | sp\|Q7TMB8-2\|CYFP1_MOUSE-DECOY Isoform 2 of Cytoplasmic FMR1-interacting protein 1... |
| tr\|Q3TYX0\|Q3TYX0_MOUSE-DECOY Uncharacterized protein OS=Mus musculus OX=10090 GN=St5... | tr\|Q3TYX0\|Q3TYX0_MOUSE-DECOY Uncharacterized protein OS=Mus musculus OX=10090 GN=St5... |
| tr\|Q8BR43\|Q8BR43_MOUSE-DECOY Uncharacterized protein OS=Mus musculus OX=10090 GN=Jph3... | tr\|Q8BR43\|Q8BR43_MOUSE-DECOY Uncharacterized protein OS=Mus musculus OX=10090 GN=Jph3... |
| tr\|Q80TX5\|Q80TX5_MOUSE MKIAA0572 protein (Fragment) OS=Mus musculus OX=10090 GN=Mcm3ap... | tr\|Q80TX5\|Q80TX5_MOUSE MKIAA0572 protein (Fragment) OS=Mus musculus OX=10090 GN=Mcm3ap... |

| | |
|---|---|
| sp\|Q8BJE2-2\|BTNL9_MOUSE-DECOY Isoform 2 of Butyrophilin-like protein 9 OS=Mus musculus... | sp\|Q8BJE2-2\|BTNL9_MOUSE-DECOY Isoform 2 of Butyrophilin-like protein 9 OS=Mus musculus... |
| sp\|O88307\|SORL_MOUSE-DECOY Sortilin-related receptor OS=Mus musculus OX=10090 GN=Sorl1... | sp\|O88307\|SORL_MOUSE-DECOY Sortilin-related receptor OS=Mus musculus OX=10090 GN=Sorl1... |
| tr\|E9Q7R9\|E9Q7R9_MOUSE Cilia and flagella-associated protein 43 OS=Mus musculus OX=10090... | tr\|E9Q7R9\|E9Q7R9_MOUSE Cilia and flagella-associated protein 43 OS=Mus musculus OX=10090... |
| tr\|A0A1D5RMD1\|A0A1D5RMD1_MOUSE-DECOY Predicted gene 16486 OS=Mus musculus OX=10090... | tr\|A0A1D5RMD1\|A0A1D5RMD1_MOUSE-DECOY Predicted gene 16486 OS=Mus musculus OX=10090... |
| sp\|P62274\|RS29_MOUSE 40S ribosomal protein S29 OS=Mus musculus OX=10090 GN=Rps29... | sp\|P62274\|RS29_MOUSE 40S ribosomal protein S29 OS=Mus musculus OX=10090 GN=Rps29... |
| sp\|P16092-6\|FGFR1_MOUSE | Isoform 6 of Fibroblast growth factor receptor 1 OS=Mus musculus OX=10090 GN=Fgfr1 |
| tr\|A0A1B0GRL1\|A0A1B0GRL1_MOUSE-DECOY Predicted gene 10153 OS=Mus musculus OX=10090... | tr\|A0A1B0GRL1\|A0A1B0GRL1_MOUSE-DECOY Predicted gene 10153 OS=Mus musculus OX=10090... |
| B2RX77_MOUSE | MCG14870 OS=Mus musculus OX=10090 GN=Polr3c PE=2 SV=1 |
| TTK_MOUSE | Dual specificity protein kinase TTK OS=Mus musculus OX=10090 GN=Ttk PE=1 SV=1 |
| sp\|P52480-2\|KPYM_MOUSE-DECOY Isoform M1 of Pyruvate kinase PKM OS=Mus musculus OX=10090... | sp\|P52480-2\|KPYM_MOUSE-DECOY Isoform M1 of Pyruvate kinase PKM OS=Mus musculus OX=10090... |
| tr\|B2RY79\|B2RY79_MOUSE-DECOY Dock9 protein OS=Mus musculus OX=10090 GN=Dock9 PE=2... | tr\|B2RY79\|B2RY79_MOUSE-DECOY Dock9 protein OS=Mus musculus OX=10090 GN=Dock9 PE=2... |
| sp\|Q8R5H1-2\|UBP15_MOUSE-DECOY Isoform 2 of Ubiquitin carboxyl-terminal hydrolase... | sp\|Q8R5H1-2\|UBP15_MOUSE-DECOY Isoform 2 of Ubiquitin carboxyl-terminal hydrolase... |
| sp\|Q6QI06-2\|RICTR_MOUSE-DECOY Isoform 2 of Rapamycin-insensitive companion of mTOR... | sp\|Q6QI06-2\|RICTR_MOUSE-DECOY Isoform 2 of Rapamycin-insensitive companion of mTOR... |
| tr\|A2AAU3\|A2AAU3_MOUSE Probable helicase with zinc finger domain OS=Mus musculus... | tr\|A2AAU3\|A2AAU3_MOUSE Probable helicase with zinc finger domain OS=Mus musculus... |

| | |
|---|---|
| tr\|Q8R562\|Q8R562_MOUSE-DECOY Ribonucleoprotein OS=Mus musculus OX=10090 GN=Trove2... | tr\|Q8R562\|Q8R562_MOUSE-DECOY Ribonucleoprotein OS=Mus musculus OX=10090 GN=Trove2... |
| sp\|Q8CC88\|VWA8_MOUSE-DECOY von Willebrand factor A domain-containing protein 8 OS=Mus... | sp\|Q8CC88\|VWA8_MOUSE-DECOY von Willebrand factor A domain-containing protein 8 OS=Mus... |
| sp\|A2A8L1\|CHD5_MOUSE-DECOY Chromodomain-helicase-DNA-binding protein 5 OS=Mus musculus... | sp\|A2A8L1\|CHD5_MOUSE-DECOY Chromodomain-helicase-DNA-binding protein 5 OS=Mus musculus... |
| tr\|Q3TZI5\|Q3TZI5_MOUSE-DECOY DNA mismatch repair protein OS=Mus musculus OX=10090... | tr\|Q3TZI5\|Q3TZI5_MOUSE-DECOY DNA mismatch repair protein OS=Mus musculus OX=10090... |
| tr\|Q6PCP0\|Q6PCP0_MOUSE-DECOY DEAD (Asp-Glu-Ala-Asp) box polypeptide 21 OS=Mus musculus... | tr\|Q6PCP0\|Q6PCP0_MOUSE-DECOY DEAD (Asp-Glu-Ala-Asp) box polypeptide 21 OS=Mus musculus... |
| E9Q419_MOUSE | Beta-adrenergic receptor kinase 1 OS=Mus musculus OX=10090 GN=Grk2 PE=1 SV=1 |
| sp\|Q8R5K2-2\|UBP33_MOUSE-DECOY Isoform 2 of Ubiquitin carboxyl-terminal hydrolase... | sp\|Q8R5K2-2\|UBP33_MOUSE-DECOY Isoform 2 of Ubiquitin carboxyl-terminal hydrolase... |
| tr\|B2RQL1\|B2RQL1_MOUSE-DECOY 4932413O14Rik protein OS=Mus musculus OX=10090 GN=Ccdc158... | tr\|B2RQL1\|B2RQL1_MOUSE-DECOY 4932413O14Rik protein OS=Mus musculus OX=10090 GN=Ccdc158... |
| sp\|Q9D4H4-2\|AMOL1_MOUSE-DECOY Isoform 2 of Angiomotin-like protein 1 OS=Mus musculus... | sp\|Q9D4H4-2\|AMOL1_MOUSE-DECOY Isoform 2 of Angiomotin-like protein 1 OS=Mus musculus... |
| tr\|A0A1L1SQF0\|A0A1L1SQF0_MOUSE Uncharacterized protein (Fragment) OS=Mus musculus... | Uncharacterized protein (Fragment) OS=Mus musculus OX=10090 PE=4 SV=1 |
| sp\|E9Q6J5-3\|BD1L1_MOUSE-DECOY Isoform New of Biorientation of chromosomes in cell... | sp\|E9Q6J5-3\|BD1L1_MOUSE-DECOY Isoform New of Biorientation of chromosomes in cell... |
| sp\|Q8CG72-2\|ARHL2_MOUSE Isoform 2 of Poly(ADP-ribose) glycohydrolase ARH3 OS=Mus... | sp\|Q8CG72-2\|ARHL2_MOUSE Isoform 2 of Poly(ADP-ribose) glycohydrolase ARH3 OS=Mus... |
| sp\|Q9QYX7-2\|PCLO_MOUSE-DECOY Isoform 2 of Protein piccolo OS=Mus musculus OX=10090... | sp\|Q9QYX7-2\|PCLO_MOUSE-DECOY Isoform 2 of Protein piccolo OS=Mus musculus OX=10090... |
| sp\|E9Q286\|ICE1_MOUSE-DECOY Little elongation complex subunit 1 OS=Mus musculus OX=10090... | sp\|E9Q286\|ICE1_MOUSE-DECOY Little elongation complex subunit 1 OS=Mus musculus OX=10090... |

| | |
|---|---|
| sp\|Q9ESZ8-2\|GTF2I_MOUSE-DECOY Isoform 2 of General transcription factor II-I OS=Mus... | sp\|Q9ESZ8-2\|GTF2I_MOUSE-DECOY Isoform 2 of General transcription factor II-I OS=Mus... |
| sp\|Q8CIN4\|PAK2_MOUSE Serine/threonine-protein kinase PAK 2 OS=Mus musculus OX=10090... | sp\|Q8CIN4\|PAK2_MOUSE Serine/threonine-protein kinase PAK 2 OS=Mus musculus OX=10090... |
| sp\|A2A5R2\|BIG2_MOUSE-DECOY Brefeldin A-inhibited guanine nucleotide-exchange protein... | sp\|A2A5R2\|BIG2_MOUSE-DECOY Brefeldin A-inhibited guanine nucleotide-exchange protein... |
| Q8BKH5_MOUSE | RIKEN cDNA C330023M02, isoform CRA_b OS=Mus musculus OX=10090 GN=Naa25 PE=2 SV=1 |
| GSH1_MOUSE | Glutamate--cysteine ligase catalytic subunit OS=Mus musculus OX=10090 GN=Gclc PE=1 SV=4 |
| Q8BSN6_MOUSE | Vesicle-associated membrane protein 4 OS=Mus musculus OX=10090 GN=Vamp4 PE=1 SV=1 |
| sp\|Q8CIE6\|COPA_MOUSE Coatomer subunit alpha OS=Mus musculus OX=10090 GN=Copa PE=1... | sp\|Q8CIE6\|COPA_MOUSE Coatomer subunit alpha OS=Mus musculus OX=10090 GN=Copa PE=1... |
| sp\|P53349\|M3K1_MOUSE Mitogen-activated protein kinase kinase kinase 1 OS=Mus musculus... | sp\|P53349\|M3K1_MOUSE Mitogen-activated protein kinase kinase kinase 1 OS=Mus musculus... |
| sp\|Q64735\|CR1L_MOUSE | Complement component receptor 1-like protein OS=Mus musculus OX=10090 GN=Cr1l PE=1 SV=1 |
| sp\|Q9QXS6-2\|DREB_MOUSE-DECOY Isoform A2 of Drebrin OS=Mus musculus OX=10090 GN=Dbn1 | sp\|Q9QXS6-2\|DREB_MOUSE-DECOY Isoform A2 of Drebrin OS=Mus musculus OX=10090 GN=Dbn1 |
| D3YX61_MOUSE | Rho-related GTP-binding protein RhoJ OS=Mus musculus OX=10090 GN=Rhoj PE=1 SV=1 |
| Q91YQ6_MOUSE | Acot5 protein OS=Mus musculus OX=10090 GN=Acot5 PE=2 SV=1 |
| Q9D3H4_MOUSE | Keratin-associated protein 2-4 OS=Mus musculus OX=10090 GN=Gm11938 PE=2 SV=1 |
| sp\|Q8R326\|PSPC1_MOUSEParaspeckle component 1 OS=Mus musculus OX=10090 GN=Pspc1 PE=1 SV=1 | sp\|Q8R326\|PSPC1_MOUSE Paraspeckle component 1 OS=Mus musculus OX=10090 GN=Pspc1 PE=1 SV=1 |

| | |
|---|---|
| sp\|Q7M6Z4-2\|KIF27_MOUSE Isoform 2 of Kinesin-like protein KIF27 OS=Mus musculus OX=10090... | Isoform 2 of Kinesin-like protein KIF27 OS=Mus musculus OX=10090 GN=Kif27 |
| A0A0A6YW33_MOUSE | MCG18090 OS=Mus musculus OX=10090 GN=Gm6525 PE=4 SV=1 |
| K3W4M7_MOUSE | Putative ATP-dependent RNA helicase DHX33 (Fragment) OS=Mus musculus OX=10090 GN=Dhx33 PE=1 SV=1 |
| B2RQR1_MOUSE | Notch gene homolog 4 (Drosophila) OS=Mus musculus OX=10090 GN=Notch4 PE=2 SV=1 |
| A2AP88_MOUSE | Zinc finger CCCH domain-containing protein 6 OS=Mus musculus OX=10090 GN=Zc3h6 PE=1 SV=1 |
| Q3UGI2_MOUSE | Uncharacterized protein OS=Mus musculus OX=10090 GN=Gpatch2l PE=2 SV=1 |
| E9PV91_MOUSE | A disintegrin and metalloproteinase with thrombospondin motifs 4 OS=Mus musculus OX=10090 GN=Adamts4 PE=1 SV=1 |
| D3Z2F2_MOUSE | 60 kDa heat shock protein, mitochondrial (Fragment) OS=Mus musculus OX=10090 GN=Hspd1 PE=1 SV=1 |
| tr\|Q3TUG2\|Q3TUG2_MOUSE DNA-directed RNA polymerase subunit (Fragment) OS=Mus musculus... | DNA-directed RNA polymerase subunit (Fragment) OS=Mus musculus OX=10090 GN=Polr1a PE=2 SV=1 |
| Q8CDE0_MOUSE | Uncharacterized protein OS=Mus musculus OX=10090 GN=Flot2 PE=2 SV=1 |
| Q8R268_MOUSE | Vomeronasal type-1 receptor OS=Mus musculus OX=10090 GN=Vmn1r212 PE=2 SV=1 |
| Q8BM06_MOUSE | Uncharacterized protein OS=Mus musculus OX=10090 GN=Nell2 PE=2 SV=1 |
| sp\|E9Q8T7\|DYH1_MOUSE-DECOY Dynein heavy chain 1, axonemal OS=Mus musculus OX=10090... | sp\|E9Q8T7\|DYH1_MOUSE-DECOY Dynein heavy chain 1, axonemal OS=Mus musculus OX=10090... |
| DRG2_MOUSE | Developmentally-regulated GTP-binding protein 2 OS=Mus musculus OX=10090 GN=Drg2 PE=1 SV=1 |
| A0A286YE31_MOUSE | Ubiquitin-conjugating enzyme E2 E1 (Fragment) OS=Mus musculus OX=10090 GN=Ube2e1 PE=1 SV=1 |

| | |
|---|---|
| D6RG95_MOUSE | YTH domain-containing protein 1 OS=Mus musculus OX=10090 GN=Ythdc1 PE=1 SV=1 |
| B7ZNS2_MOUSE | Disks large-associated protein 4 OS=Mus musculus OX=10090 GN=Dlgap4 PE=1 SV=1 |
| tr\|Q8BP21\|Q8BP21_MOUSE Uncharacterized protein OS=Mus musculus OX=10090 GN=Cdk4 PE=2... | tr\|Q8BP21\|Q8BP21_MOUSE Uncharacterized protein OS=Mus musculus OX=10090 GN=Cdk4 PE=2... |
| B2RWZ7_MOUSE | Phospholipase A2 OS=Mus musculus OX=10090 GN=Pla2g4f PE=2 SV=1 |
| sp\|Q9QYF9\|NDRG3_MOUSEProtein NDRG3 OS=Mus musculus OX=10090 GN=Ndrg3 PE=1 SV=1 | sp\|Q9QYF9\|NDRG3_MOUSE Protein NDRG3 OS=Mus musculus OX=10090 GN=Ndrg3 PE=1 SV=1 |
| E9QK04_MOUSE | Neogenin OS=Mus musculus OX=10090 GN=Neo1 PE=1 SV=1 |
| Q3TYL7_MOUSE | DnaJ homolog subfamily B member 1 OS=Mus musculus OX=10090 GN=Dnajb1 PE=1 SV=1 |
| sp\|Q80TL7-2\|MON2_MOUSE | Isoform 2 of Protein MON2 homolog OS=Mus musculus OX=10090 GN=Mon2 |
| A0A0U1RP06_MOUSE | DNA repair protein-complementing XP-C cells homolog (Fragment) OS=Mus musculus OX=10090 GN=Xpc PE=1 SV=1 |
| B9EKC3_MOUSE | Rho GTPase activating protein 5 OS=Mus musculus OX=10090 GN=Arhgap5 PE=2 SV=1 |
| A0A1Y7VLS3_MOUSE | TOG array regulator of axonemal microtubules protein 1 OS=Mus musculus OX=10090 GN=Togaram1 PE=1 SV=1 |
| TWF1_MOUSE | Twinfilin-1 OS=Mus musculus OX=10090 GN=Twf1 PE=1 SV=2 |
| G3X8V0_MOUSE | Leucine-rich repeat-containing protein 63 OS=Mus musculus OX=10090 GN=Lrrc63 PE=4 SV=1 |
| tr\|Q3V117\|Q3V117_MOUSE ATP-citrate synthase OS=Mus musculus OX=10090 GN=Acly PE=1... | ATP-citrate synthase OS=Mus musculus OX=10090 GN=Acly PE=1 SV=1 |
| LSHR_MOUSE | Lutropin-choriogonadotropic hormone receptor OS=Mus musculus OX=10090 GN=Lhcgr PE=2 SV=1 |
| K0BWC3_MOUSE | Palladin isoform OS=Mus musculus OX=10090 GN=Palld PE=2 SV=1 |

| | |
|---|---|
| Q3U261_MOUSE | Uncharacterized protein (Fragment) OS=Mus musculus OX=10090 GN=Tacstd2 PE=2 SV=1 |
| BROMI_MOUSE | Isoform 2 of Protein broad-minded OS=Mus musculus OX=10090 GN=Tbc1d32 |
| Q3U2X1_MOUSE | Uncharacterized protein OS=Mus musculus OX=10090 GN=Nup62 PE=2 SV=1 |
| tr\|E9Q6A7\|E9Q6A7_MOUSE-DECOY Bromodomain PHD finger transcription factor OS=Mus musculus... | tr\|E9Q6A7\|E9Q6A7_MOUSE-DECOY Bromodomain PHD finger transcription factor OS=Mus musculus... |
| E9Q3A0_MOUSE | E3 ubiquitin-protein ligase Trim36 OS=Mus musculus OX=10090 GN=Trim36 PE=1 SV=1 |
| D3Z4I0_MOUSE | Phosphatidylinositol glycan anchor biosynthesis, class G OS=Mus musculus OX=10090 GN=Pigg PE=1 SV=1 |
| sp\|P08101-3\|FCGR2_MOUSE | Isoform IIB1' of Low affinity immunoglobulin gamma Fc region receptor II OS=Mus musculus OX=10090 GN=Fcgr2 |
| sp\|Q6PCM2-2\|INT6_MOUSE | Isoform 2 of Integrator complex subunit 6 OS=Mus musculus OX=10090 GN=Ints6 |
| tr\|A0A1Y7VM39\|A0A1Y7VM39_MOUSE Extended synaptotagmin-2 OS=Mus musculus OX=10090... | tr\|A0A1Y7VM39\|A0A1Y7VM39_MOUSE Extended synaptotagmin-2 OS=Mus musculus OX=10090... |
| sp\|A2AFR3-2\|FRPD4_MOUSE-DECOY Isoform 2 of FERM and PDZ domain-containing protein... | sp\|A2AFR3-2\|FRPD4_MOUSE-DECOY Isoform 2 of FERM and PDZ domain-containing protein... |
| sp\|Q8BGF7-4\|PAN2_MOUSE | Isoform 4 of PAN2-PAN3 deadenylation complex catalytic subunit Pan2 OS=Mus musculus OX=10090 GN=Pan2 |
| D3YU89_MOUSE | Citron Rho-interacting kinase OS=Mus musculus OX=10090 GN=Cit PE=1 SV=1 |
| A0A023T778_MOUSE | Mago nashi protein OS=Mus musculus OX=10090 GN=Magohb PE=1 SV=1 |
| A0A1W2P7W3_MOUSE | Protein tyrosine phosphatase, receptor type, B, isoform CRA_b OS=Mus musculus OX=10090 GN=Ptprb PE=1 SV=1 |
| A0A2I3BRX9_MOUSE | Ral GTPase-activating protein subunit alpha-1 OS=Mus musculus OX=10090 GN=Ralgapa1 PE=1 SV=1 |
| sp\|P53564-3\|CUX1_MOUSE | Isoform 3 of Homeobox protein cut-like 1 OS=Mus musculus OX=10090 GN=Cux1 |

| | |
|---|---|
| ZO2_MOUSE | Tight junction protein ZO-2 OS=Mus musculus OX=10090 GN=Tjp2 PE=1 SV=2 |
| sp\|Q9D4J7\|PHF6_MOUSE | PHD finger protein 6 OS=Mus musculus OX=10090 GN=Phf6 PE=1 SV=1 |

Supplementary Data 2. Proteomics results, neurons.
